# Modular small RNA drives pathogen emergence

**DOI:** 10.1101/2023.03.17.533193

**Authors:** Deepak Balasubramanian, Salvador Almagro-Moreno

## Abstract

Pathogen emergence is a poorly understood complex phenomenon. To date, the molecular mechanisms that allow strains within a bacterial population to emerge as human pathogens remain mostly enigmatic. We recently uncovered that toxigenic *Vibrio cholerae* encode preadaptations to host colonization, what we term virulence adaptive polymorphisms (VAPs), however, the molecular mechanisms driving them are not known. *ompU* is a VAP-encoding gene that is associated with the production of the major outer membrane porin OmpU. Here, we show that the *ompU* ORF also encodes a modular small RNA overlapping its 3’ terminus that plays a major role in *V. cholerae* physiology. We determined that the OmpU-encoded sRNA (OueS) strongly suppresses biofilm formation, a phenotype that is essential for host intestinal colonization, via repression of iron uptake. OueS controls over 84% of the genes regulated by ToxR, a major virulence regulator, and plays an integral role during the infection process. We demonstrate that OueS is critical for intestinal colonization and its bimodular nature dictates the virulence potential of *V. cholerae*. Overall, our study reveals specific molecular mechanisms leading to the emergence of pathogenic traits in bacteria unveiling the hidden genetics associated with this process. We propose a scenario where a limited number of modular genes could explain the emergence of novel phenotypic traits in biological systems.

## INTRODUCTION

The emergence of human pathogens is one of the most pressing public health concerns of modern times. Pathogen emergence is a complex and multifactorial phenomenon that culminates with the organism acquiring the ability to colonize and harm the human host^1–3^. Understanding the specific elements associated with this process is critical for successful disease management and control^4^. Even though recent insights have begun to elucidate some of the mechanisms and forces driving pathogen emergence, to date, the molecular pathways that facilitate certain strains within a species to emerge as pathogens remain largely unknown^4–8^.

*Vibrio cholerae* O1 is the etiological agent of the severe diarrheal disease cholera and remains a major scourge in places with limited access to clean drinking water and proper sanitation practices^9,10^. Following ingestion in the form of planktonic cells or as part of a biofilm, *V. cholerae* must survive adverse conditions such as stomach acidity, bile, or host antimicrobial peptides to colonize the small intestine^11–14^. Successful colonization requires stringent downregulation of biofilm formation, as it triggers a strong immune response, and upregulation of flagella-mediated motility to swim through the mucus layer towards the epithelium^15–17^. A complex regulatory cascade mediated by the virulence regulator ToxR facilitates adaptation to the harsh conditions within the host and, along with other regulators, the timely expression of virulence genes^18–21^.

ToxR is a transmembrane protein with a cytoplasmic DNA-binding domain that controls the expression of hundreds of genes^19, 22–24^. Among them, ToxR directly regulates the expression of the cholera toxin (CT), the source of the watery diarrhea associated with cholera, and, together with TcpP, the expression of ToxT, a master virulence regulator^22, 25^. ToxR also reciprocally modulates the expression of two outer membrane porins: OmpU and OmpT. Specifically, ToxR activates transcription of *ompU* whereas it represses *ompT* expression^23, 26, 27^. OmpU is highly expressed during the infection process of *V. cholerae*, making up to 60% of the outer membrane and is essential for host survival, virulence, and intestinal colonization^23, 28, 29^. The porin OmpU is associated with numerous virulence-related phenotypes and confers bile and anionic detergent resistance, and tolerance to organic acids and cationic peptides, such as the host derived peptide P2 and the antibiotic polymyxin B^11, 12, 23, 29, 30^. Beyond resistance to cell damaging agents, OmpU has also been associated with several more complex phenotypes. For instance, OmpU exerts immunomodulatory effects such as induction of host tolerance to LPS and suppression of the host immune response by inducing M1 polarization of macrophages^31, 32^. Furthermore, Provenzano *et al*. showed that a Δ*ompU* mutant exhibits altered production of CT and TCP^29^. However, the mechanisms by which an outer membrane porin like OmpU can be involved in and modulate such diverse virulence-related phenotypes remain poorly understood.

Despite the presence of over 200 serogroups, only *V. cholerae* strains from serogroups O1 and the now extinct O139 can cause cholera in humans^33–35^. Strains from these two serogroups are phylogenetically related and confined to a single clade containing all toxigenic *V. cholerae* called the pandemic cholera group^36^. We previously developed a comparative genomics framework to elucidate the evolutionary origins and emergence of toxigenic *V. cholerae* strains^37^. Our analyses of environmental and clinical strains of *V. cholerae* identified the presence of allelic variations in core genes that circulate in non-toxigenic environmental populations and confer preadaptations to host colonization, enhancing their pandemic potential. The presence of these virulence adaptive polymorphisms (VAPs) is associated with the emergence of toxigenic strains of *V. cholerae*, explaining their limited distribution to specific clades^37^. Interestingly, we determined that *ompU* in toxigenic *V. cholerae* strains encode VAPs and some of the virulence traits associated with the porin are allelic-dependent^37, 38^. Furthermore, VAP-encoding environmental alleles of *ompU* confer phenotypes associated with clinical outcomes such as antimicrobial resistance^37, 38^. These phenotypes are absent from mutant strains with *ompU* alleles that do not encode VAPs, indicating that these adaptations to virulence are present in their genomic background prior to host colonization^37, 38^. Overall, our data indicates that OmpU-associated traits are dependent on the specific allele encoded by a given strain and is directly associated with the emergence of toxigenic *V. cholerae*. To date, the molecular mechanisms controlling these allelic-dependent phenotypes remain unaddressed.

In this study, we determined that the toxigenic allele of OmpU stringently represses biofilm formation and modulates the transcriptome of *V. cholerae*. We identified a novel sRNA encoded overlapping the 3’ terminus and downstream region of the *ompU* ORF that is responsible for this behavior, which we termed *<u>o</u>mp<u>U</u>*-<u>e</u>ncoded <u>s</u>RNA (OueS). We demonstrate that OueS represses biofilm formation in toxigenic *V. cholerae* by repressing iron uptake mechanisms. Transcriptome analysis reveals that OueS modulates 84.85% of the ToxR virulence regulon, suggesting an integral role of OueS in *V. cholerae* pathogenesis. Finally, we show that OueS is critical for intestinal colonization and the modular nature of the sRNA among *V. cholerae* strains dictates the virulence potential of the bacterium. These findings provide major insights into the emergence of pandemic *V. cholerae* and reveals specific molecular mechanisms that lead to the emergence of pathogenic traits in bacteria.

## RESULTS

### OmpU represses biofilm formation in *V. cholerae* and modulates its transcriptome

OmpU plays a major role in *V. cholerae* pathogenesis^15, 16–19, 23^, nonetheless, it remains unclear how an outer membrane porin can modulate disparate critical virulence-related phenotypes. For instance, we previously demonstrated that an *ompU* deletion mutant in *V. cholerae* exhibits a 3-fold increase in biofilm formation^37^. To dissect this phenotype, we ectopically expressed *ompU* from an inducible plasmid in an in-frame isogenic *ompU* deletion mutant in *V. cholerae* C6706, an El Tor strain from the seventh cholera pandemic. Expression of *ompU* decreases biofilm formation in the Δ*ompU* mutant back to wild-type (WT) levels, indicating that OmpU represses biofilm formation in *V. cholerae* (**Fig. 1A**). On the other hand, expression of an environmental *ompU* allele from a strain isolated from the Great Bay Estuary in New Hampshire, *V. cholerae* GBE1114^39^, either ectopically or through allelic exchange in the isogenic C6706 background, does not repress biofilm formation, suggesting this phenotype is allelic-dependent (**Fig. 1A**)^37^. Interestingly, biofilm suppression is critical in the early stages of intestinal colonization as biofilm components elicit a strong immune response leading to clearance of *V. cholerae* strains that do not timely repress the process^40^. Thus, we found it fascinating and quite puzzling that expression of a porin localized in the outer membrane can suppress a phenotype that is central to the infection process.

**Figure 1.**
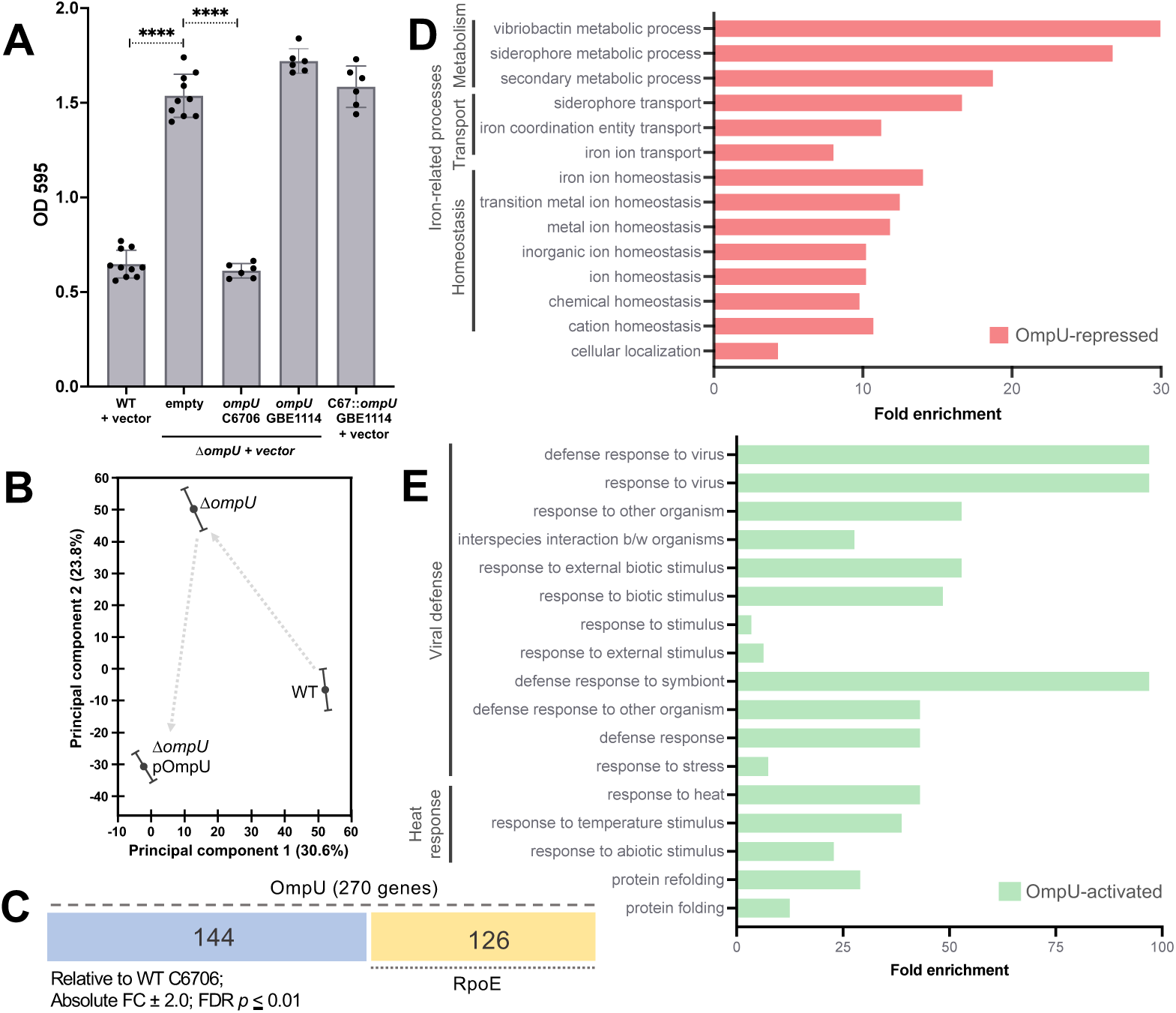
OmpU represses biofilm formation and modulates gene expression. (A) Biofilm formation of *V. cholerae* C6706 WT, Δ*ompU* and Δ*ompU* strains expressing OmpU^C6706^ or OmpU^GBE1114^, either in *cis* or *trans*. n ≥ 6. Two-tailed T-test *p*-value **** ≤ 0.0001. (B) Transcriptomes of biofilm-derived cells were visualized by principal component analysis and reveal an OmpU-mediated shift in the datasets. (C) Total number of differentially regulated genes (absolute fold change 2.0; FDR *p* ≤ 0.01) from the transcriptomes of the wild-type C6706 were compared against mutant strains of *ompU* and the alternative sigma factor-encoding gene *rpoE* from biofilm-grown cells. Functional enrichment of the OmpU-regulated genes was performed by the Panther Overrepresentation test (GO Biological process)^85, 86^ to classify the OmpU-repressed (D) and activated (E) datasets.

To gain insights into the mechanisms behind this marked phenotype, we examined the transcriptional changes associated with OmpU-mediated biofilm repression. We performed RNA-Seq analyses of *V. cholerae* biofilms produced by WT, Δ*ompU* and Δ*ompU*-pOmpU^C6706^ strains. Our data reveals a drastic shift in the transcriptome of the Δ*ompU* mutant strain compared to WT, with differential regulation of 270 genes of which 181 genes are repressed by OmpU and 89 are activated (absolute FC ≥ 2.0, FDR *p* ≤ 0.01) (**Fig 1B; Table S1**). Ectopic expression of *ompU*^C6706^ in the Δ*ompU* background results in a shift of the transcriptome, closer to WT, and the recovery in expression of the dysregulated genes (**Table S1**). GO enrichment analysis reveals that OmpU primarily represses genes involved in iron metabolism and acquisition, specifically in vibriobactin metabolism, siderophore transport, and iron ion homeostasis (FDR *p* ≤ 0.05) (**Fig 1D; Table S1**). On the other hand, OmpU activates genes involved in the defense response mechanisms to phages and symbionts, as well as those involved in heat response and protein folding (**Fig 1E**).

Upon misfolding, OmpU activates the cell envelope stress response via the release of sigma factor RpoE in the cytosol^41^. To address the potential role of RpoE in the transcriptomic changes associated with OmpU, we examined the biofilm transcriptome of a Δ*rpoE* mutant strain **(Fig 1C)**. Comparison of the transcriptome datasets of Δ*ompU* and Δ*rpoE* strains, relative to WT, demonstrates that although there is overlap between the two datasets, a large percentage of the transcriptomic changes associated with *ompU* expression appear to be independent of RpoE activation (**Fig 1C; Table S2**). Specifically, OmpU modulates the expression of 270 genes during biofilm formation and 144 of those genes (53.3%) are not associated with RpoE (**Fig 1C; Table S2**). Overall, our data indicates that OmpU represses biofilm formation in an allele-dependent manner and has a significant impact on the *V. cholerae* transcriptome.

### The *ompU* ORF encodes a small RNA

Given the large transcriptomic changes associated with OmpU extending beyond the activation of RpoE, we considered the possibility of the *ompU* gene coding for regulatory sequences within its ORF besides the porin. A genome-wide sRNA screening in *V. cholerae* indicated, among thousands of other sRNAs, potential sRNAs within the 3’ end of *ompU*^42^. To examine whether the *ompU* ORF encodes an internal sRNA, we performed northern blot analysis with probes targeting the 3’ termini of *ompU* and the 5’ termini as a control. We reasoned that a probe targeting the sRNA locus within *ompU* should result in two bands corresponding to both the *ompU* mRNA and the potential sRNA. The northern blot data shows that a probe specific for the 5’ end of the *ompU* ORF (674,890–675,054 ntd) results in a ∼1100 ntd band corresponding to the expected size of the mRNA transcript (**Fig 2A)**. On the other hand, in addition to the ∼1.1 kb mRNA band, a smaller size band appears with a probe targeting the 3’ end of the *ompU* ORF (674,837–674,876 ntd) potentially corresponding to a sRNA (∼160 ntd) (**Fig 2A**).

**Figure 2.**
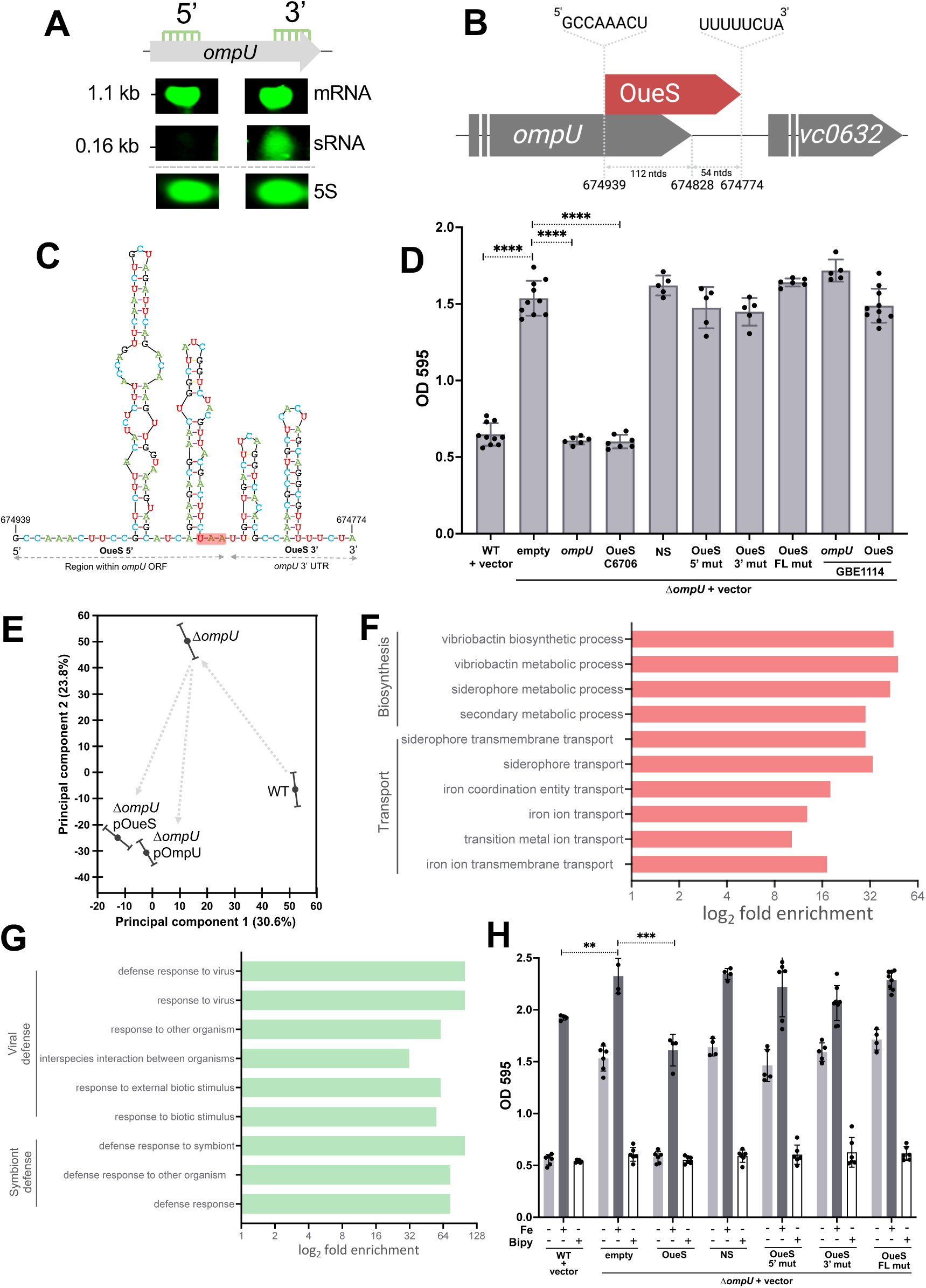
OmpU encodes a small RNA in its 3’ terminus. (A) Northern blot analyses. Total RNA was obtained at the early stationary phase (OD_600_ ∼2.0) from the WT cells and subjected to northern blot analysis using probes specific for the 5’ and 3’ ends of the *ompU* ORF. 5S rRNA-specific probes were used as a loading control. (B) Schematic representation of the genomic location, coordinates, and termini of the newly identified *ompU*-encoded sRNA, OueS. (C) Predicted secondary structure of OueS from toxigenic strains. The 5’ region of OueS that overlaps with the *ompU* ORF and the 3’ region that lies immediately downstream are indicated. The stop codon of the *ompU* ORF is highlighted in red (D) Expression of OueS from toxigenic strain *V. cholerae* C6706 specifically suppresses biofilm formation. n ≥ 5. (E) Expression of OueS, visualized as a PCA plot, reveals a shift in the transcriptome similar to ectopic OmpU expression during biofilm formation. Functional gene ontology (GO) categorization of the genes (F) repressed and (G) activated by OueS during biofilm formation was performed by the Panther Overrepresentation Test^85, 86^. OueS represses iron uptake-related categories and activates genes involved in viral and symbiont defense. (H) Biofilm formation of strains ectopically expressing either the WT or mutant versions of OueS was examined under iron-replete (dark grey bars) and iron-depleted (white bars) conditions by exogenous addition of 100 µM FeCl_3_ or 100 µM 2,2’-bipyridyl, respectively, compared to normal iron conditions (light grey bars). Two-tailed T-test *p* values for D and H: ** < 0.001, *** < 0.0001, **** < 0.00001.

To define the 5’ and 3’ termini of the potential sRNA encoded within the *ompU* ORF, we performed rapid amplification of cDNA ends (RACE) assays. Given the overabundance of the *ompU* transcript and the overlapping nature of the mRNA and the potential sRNA, our initial 5’ RACE studies yielded only the termini of ∼1100 bp *ompU* mRNA transcript. To circumvent this, total RNA from LB-grown wild-type strain was size fractionated on a MOPS agarose gel and the 100-500 ntd fraction, encompassing the potential sRNA region was extracted and used as template for RACE studies. Assays on this size-fractionated RNA led to the identification of a sRNA corresponding to 166 ntds that we termed *<u>o</u>mp*U-<u>e</u>ncoded <u>s</u>mall RNA (OueS) (**Fig 2B)**. Surprisingly, 112 ntds of the sRNA lies within the 3’ terminus of the *ompU* ORF and 54 ntds lie downstream of the stop codon (674,939 – 674,774 on chromosome I). *In silico* secondary structure prediction reveal a four stem-loop structure, with the terminal stem-loop ending in a poly-U tail typical of a Rho-independent terminator^43, 44^ (**Fig 2C**). Interestingly, the *ompU* allele GBE1114, which does not lead to biofilm suppression, encodes a distinct allele of OueS (**Fig S1**). Even though the length of both alleles is 166 ntds, their predicted secondary structures drastically differ in the 5’ region encoded within the *ompU* ORF but is identical in the OueS region downstream of *ompU* suggesting a modular structure of OueS (**Fig S1B, F**). Overall, our results indicate that the *ompU* ORF in *V. cholerae* encodes both the OmpU porin and the 5’ terminus of a modular 166 ntd sRNA that we termed OueS.

### OueS represses biofilm formation in *V. cholerae*

Next, we examined whether the *ompU*-mediated biofilm repression seen in **Fig 1A** is mediated by OueS. We ectopically expressed OueS in the Δ*ompU* mutant (Δ*ompU*-pOueS) and determined that the strain recovers the WT phenotype and displays reduced biofilm formation when compared to Δ*ompU* (**Fig 2D**). This finding indicates that OueS is responsible for the *ompU*-mediated biofilm repression in *V. cholerae* (**Fig 2D**). To test the specificity of OueS-mediated biofilm repression, first we ectopically expressed a non-specific (NS) region outside of the OueS locus located towards the 5’ end of the *ompU* ORF (675,307–675,521). The NS region had no effect on biofilm formation compared to the Δ*ompU* strain (**Fig 2D**). Subsequently, we planned on generating mutations within OueS to potentially nullify its effect. However, the 5’ end of OueS is encoded within the *ompU* ORF and thus, mutations affecting OueS could also affect the OmpU coding region. To ensure that the mutations within the sRNA do not disrupt the protein coding sequence, we took advantage of the codon wobble and mutated every third base in the OueS locus, disrupting the sRNA while preserving the OmpU amino acid sequence. Three different OueS mutants were generated: OueS 5’-mut harboring mutations exclusively in the OueS region that lies within the *ompU* ORF, OueS 3’-mut, with mutations only in the OueS region downstream of *ompU*, and OueS containing mutations across the full length of OueS, FL-mut. The mutations within OueS in all three cases disrupted their predicted secondary structures while retaining the OmpU protein coding sequence (**Fig S1**). Ectopic expression of either of the three OueS mutant sRNAs (OueS 5’-mut, 3’-mut and FL-mut) in the Δ*ompU* strain do not lead to suppression of biofilm formation (**Fig 2D**). Furthermore, unlike the OueS allele from clinical strain C6706, ectopic expression of OueS^GBE1114^ does not lead to biofilm suppression either (**Fig 2D**). Overall, our results demonstrate that **a)** OueS is responsible for biofilm repression in *V. cholerae*, **b)** this phenotype is dependent on the OueS allele and **c)** the sRNA has a modular structure that divides its *ompU*-encoded 5’ end from its 3’ terminus.

### OueS is responsible for *ompU-*dependent transcriptomic changes during biofilm formation

OmpU modifies the transcriptome of *V. cholerae* during biofilm formation (**Fig 1B and C**). We hypothesized that expression of OueS would be responsible for this altered transcriptional profile. To address this, we examined the transcriptomes of Δ*ompU*-pOueS cells harvested from biofilms. Comparison of the biofilm transcriptomes of Δ*ompU*-pOueS to those of the WT and Δ*ompU* strains reveal that OueS expression restores the shift in the transcriptome similar to ectopic *ompU* expression (**Fig 2E**). We compared the differentially regulated genes (absolute FC ≥ 2.0, FDR *p* ≤ 0.01) of two pairwise comparisons: WT Vs Δ*ompU* (270 genes) and Δ*ompU*-pOueS Vs Δ*ompU* (267 genes). There are 132 genes (48.89%) in the overlap region whose expression is modulated by OueS, 101 of which are repressed and 31 activated (**Table S3)**. Analysis of the expression values of these 132 genes between the two pair-wise comparisons reveals that the direction of the fold change (activation Vs repression) as well as the expression levels are identical in 131 of the 132 genes between the two groups (Pearson Correlation Coefficient = 0.9998; **Table S3**). GO enrichment analysis on the genes activated and repressed by OueS reveal that 63 of the 101 OueS repressed genes are enriched in the GO Biological process database and 17 of the 31 activated ones are enriched. Similar to the OmpU transcriptome, among the OueS-repressed genes there was a significant positive enrichment (FDR *p* < 0.05) of genes belonging to the vibriobactin biosynthetic processes and siderophore transmembrane transport (**Fig 2F**). Furthermore, the OueS activated genes are primarily involved in the ‘defense response to virus’ and ‘defense response to symbionts’ categories (**Fig 2G**). Overall, our data indicates that OueS is responsible for RpoE-independent changes in the biofilm transcriptome by OmpU.

### OueS suppresses biofilm formation by repressing iron uptake

We analyzed the OueS-regulated genes for insights into the molecular basis behind the stringent biofilm repression associated with the sRNA. Iron is important for biofilm formation and OueS downregulates the expression of numerous iron uptake genes^45, 46^ (**Fig 2F**; **Table S3**). Next, we examined the interplay between iron availability and biofilm formation and their association with OueS. Addition of exogenous iron to the media results in increased biofilm formation across all the strains virtually negating the role of OueS in biofilm repression (**Fig 2H**). On the other hand, depletion of the native iron in the media using the iron chelator 2,2’-bipyridyl (100 µM), results in drastically reduced biofilm formation in all strains, closely resembling the phenotype associated with OueS production (**Fig 2H**). Furthermore, expression of OueS during planktonic growth does not affect growth patterns irrespective of iron levels in the media (**Fig S2**). Overall, our data strongly indicates that OueS suppresses biofilm formation by repressing iron uptake in *V. cholerae*.

### OueS is associated with ToxR-mediated phenotypes

ToxR directly regulates the expression of *ompU*^47^. Consequently, next, we investigated the potential regulatory relationships between ToxR and OueS. Deletion of ToxR leads to an increase in biofilm production similar to a Δ*ompU* mutant (**Fig 3A**). Ectopic expression of either the *ompU* ORF or OueS in the Δ*toxR* strain restores biofilm repression to WT levels (**Fig 3A**). This OueS-mediated repression is specific as the WT phenotype could not be rescued by ectopic expression of neither the OueS 5’-, 3’- or FL-mut alleles nor the NS region within the *ompU* ORF (**Fig 3A**). Furthermore, pairwise comparisons of WT and Δ*toxR*-pOueS relative to Δ*toxR* reveal that 42.69% of the ToxR regulated genes in biofilms overlap with the OueS transcriptome (155 of 363 genes; FC ± 2.0, FDR *p* ≤ 0.01; **Table S6**). The directionality of expression (activation Vs repression) in 149 of these 155 OueS-regulated genes (96%) is identical between the WT and Δ*toxR*-pOueS strains, strongly suggesting that OueS restores a significant portion of the regulatory changes observed with the loss of ToxR function in biofilms (**Table S6**). Overall, these findings indicate that the ToxR-mediated biofilm repression is due to downstream OueS expression and highlight a connection between the sRNA and ToxR regulome.

**Figure 3.**
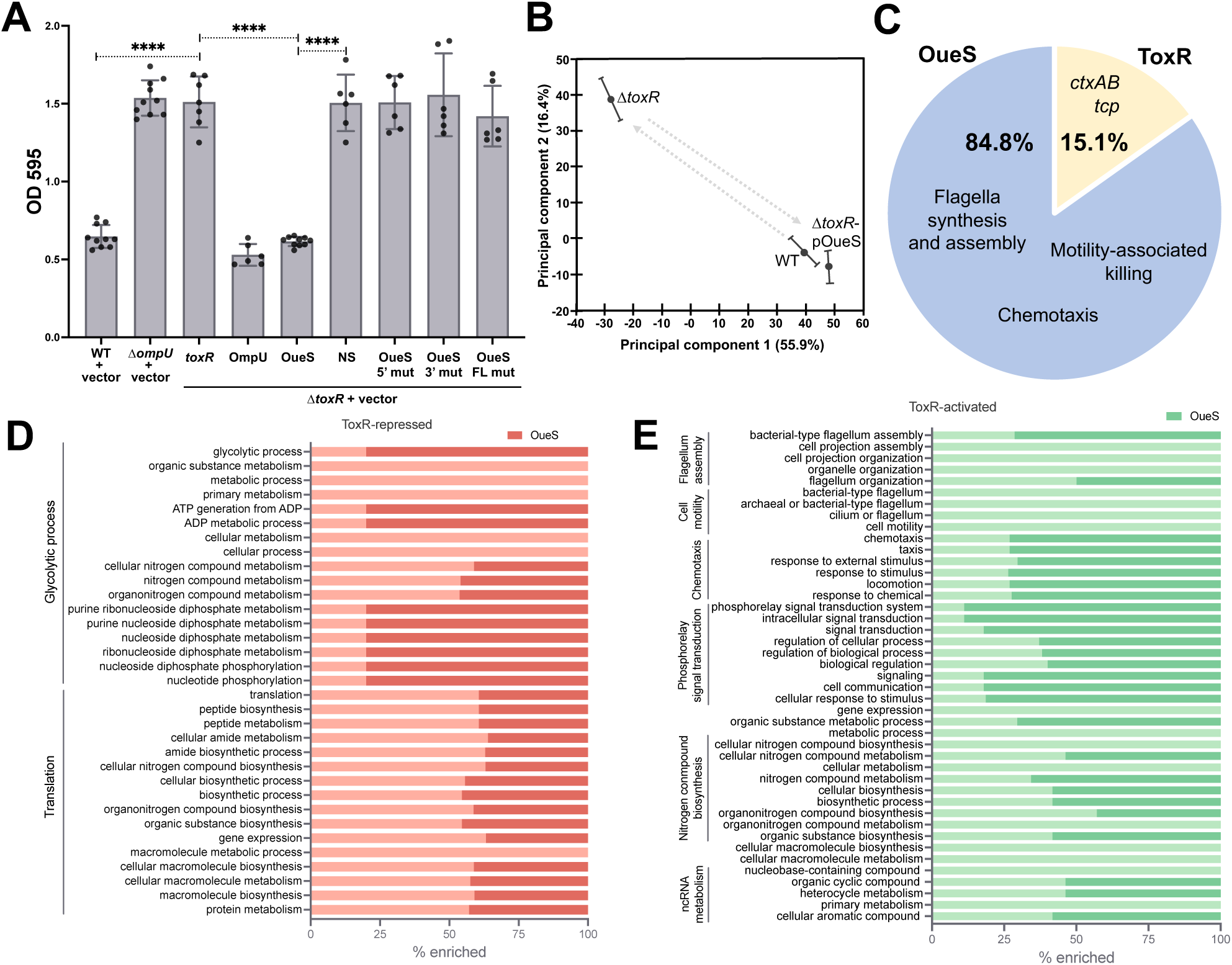
OueS acts as a functional surrogate of ToxR. (A) Ectopic expression of OueS represses biofilm formation in a Δ*toxR* mutant. n ≥ 6. Two-tailed T-test *p*-value **** ≤ 0.0001. (B-E) OueS modulates the ToxR transcriptome under virulence-inducing conditions (AKI)^48, 80^. (B) A Δ*toxR* strain ectopically expressing OueS exhibit a similar expression profile to the WT as visualized by PCA plot. (C) Analysis of the significantly regulated genes (absolute FC 2.0; FDR *p* ≤ 0.01) between the datasets reveals that OueS (blue slice) modulates the expression of 84.85% (493 genes) of the ToxR virulence regulon (581 genes). These do not include *ctxAB* or the TCP operon. Functional enrichment analysis^85, 86^ of the ToxR regulome under AKI conditions reveals that several of the functional categories that are (D) repressed (red) and (E) activated (green) by ToxR are primarily modulated by OueS (dark red, D, and dark green, E).

### OueS acts as a functional surrogate of ToxR

OmpU is highly expressed during the infection process and is critical for colonization^11, 23, 27^. Next, we examined the role of OmpU and OueS in the context of virulence-associated conditions. In order to do that, first, we analyzed the transcriptomes of the WT, Δ*ompU* and Δ*ompU*-pOmpU strains under virulence-inducing (AKI) conditions ^48^. Our analyses reveal that OmpU regulates a much larger proportion of genes under AKI conditions that during biofilm formation modulating the expression of 947 genes (560 activated and 379 repressed), compared to 270 genes (181 repressed, 89 activated) in biofilms (**Table S4**). Ectopic expression of OmpU in Δ*ompU* decreases the number of differentially expressed genes to 170 from the 947 genes in Δ*ompU* (82.04% recovery of the Δ*ompU* transcriptome to WT levels) (**Table S4**). Furthermore, most of the OmpU-modulated genes under AKI conditions are RpoE-independent as only 158 of the 947 (16.68%) are shared with the RpoE regulon under AKI conditions, suggesting a much lesser role of the envelope stress response during infection (**Table S4**). Additionally, OueS modulates the expression of 455 of the 947 genes (48%) of the OmpU transcriptome under AKI conditions suggesting an integral role of OueS in this process (**Table S5**).

Subsequently, we investigated the potential relationship of OueS and ToxR under AKI conditions. Principal component analysis (PCA) of the RNA-Seq based transcriptomic data shows a change in the transcriptome of the Δ*toxR* strain compared to WT (**Fig 3B**). Strikingly, ectopic expression of OueS in the Δ*toxR* mutant (Δ*toxR*-pOueS) under AKI conditions results in a drastic shift in the transcriptome closely resembling the WT (**Fig 3B**). To examine the extent of OueS complementation, we determined the ratio of the fold change between the ToxR and OueS regulons [(*toxR*/WT) / (*toxR*-pOueS/WT)]. Our analysis reveals that OueS modulates the expression of at least 493 of the 581 genes within the ToxR regulon (84.85%) under AKI conditions (**Fig 3C**; **Table S7**). Importantly, OueS does not modulate expression of *ctxAB* or the TCP operon (**Fig 3C**; **Table S7**). Finally, we performed a functional enrichment analysis of the 581 genes of the ToxR regulon to determine the specific genes that OueS regulates based on functional categories. OueS is responsible for the control of most of the ToxR-downregulated genes involved in glycolysis and translation (**Fig 3D; Table S7**). On the other hand, the ToxR upregulation of genes associated with chemotaxis, phosphorelay signaling as well as genes involved in nitrogen metabolism appear to be OueS-dependent (**Fig 3E; Table S7**). Overall, OueS acts as a functional surrogate of ToxR under virulence-inducing conditions controlling at least 84.85% of the ToxR-regulated genes.

### OueS is expressed from the *ompU* promoter

To explore the regulation of OueS expression, first we generated mutant strains where we cloned OueS together with varying lengths of its upstream region to identify a potential promoter within *ompU*. We did so in 10 bp increments (P_0_, P_10_, P_20_, … P_100_) from zero bp upstream of OueS (P_0_) to 100 bp upstream (P_100_) (**Fig S3**). We monitored biofilm formation of these strains and found that all repress biofilm formation similar to WT, indicating that OueS does not have a native promoter within *ompU* but appears to be driven by the *ompU* promoter (P*_ompU_*) (**Fig S3)**. To verify this, we generated an *ompU* promoter deletion mutant. We deleted the 300 bp region immediately upstream of the *ompU* start codon, which encodes both the promoter and transcription start site^49^, generating the strain ΔP*_ompU_*, which retains an intact *ompU* and OueS coding region. Biofilm formation in ΔP*_ompU_* is similar to a Δ*ompU* strain indicating lack of OueS expression (**Fig S3)**. Thus, the expression of OueS is driven from P*_ompU_* and not from an internal promoter, suggesting it is controlled by ToxR.

### OueS is required for intestinal colonization

OueS is essential for phenotypes associated with successful host colonization such as biofilm repression and controls a large percentage of the ToxR regulon. In order to examine whether OueS plays a direct role in intestinal colonization, we performed competition assays using the infant mouse model of infection^50, 51^. Inducer-dependent ectopic expression of the sRNA is not feasible in animal models. However, our data demonstrates that the strain expressing OueS directly from P*_ompU_* in the Δ*ompU* background (Δ*ompU*-P*_ompU_*OueS) exhibits an identical phenotype as the strain ectopically expressing the sRNA making it suitable for *in vivo* intestinal colonization assays (**Fig S3**). We also generated isogenic mutant strains encoding the OueS variants (5’-, 3’- and FL-mut) directly driven by the *ompU* promoter in the Δ*ompU* background. The competitive fitness during intestinal colonization of these strains was determined against a *ΔlacZ* strain^37, 51, 52^. Further, to determine potential differences in spatial colonization dynamics of the strains, the small intestine was separated into proximal (P), medial (M), and distal (D) sections from each infected animal^51^. The ability of the various mutants to colonize the intestine relative to the WT strain was calculated by determining their competitive indices (CI). As expected, Δ*ompU* demonstrates a ∼1-log decrease in the CI across the entire length of the small intestine (CI 0.08 - 0.14; **Fig 4A**). Interestingly, the strain expressing OueS exhibits a CI similar to WT across the length of the intestine (CI 0.98 - 1.47) (**Fig 4A**). These changes in colonization are specific to OueS as strains expressing either OueS 5’-mut (CI 0.12 - 0.43), 3’-mut (CI 0.26 - 0.33) or FL-mut (P 0.16 - 0.24) are not capable of restoring WT CI (**Fig 4A**). Finally, the mutant with a deletion in the *ompU* promoter (ΔP*_ompU_*) shows a decrease in CI across the different sections of the small intestine similar to the Δ*ompU* strain (CI 0.08 - 0.11; **Fig 4A**). Taken together, our results reveal that OueS is essential for intestinal colonization of toxigenic *V. cholerae*.

**Figure 4.**
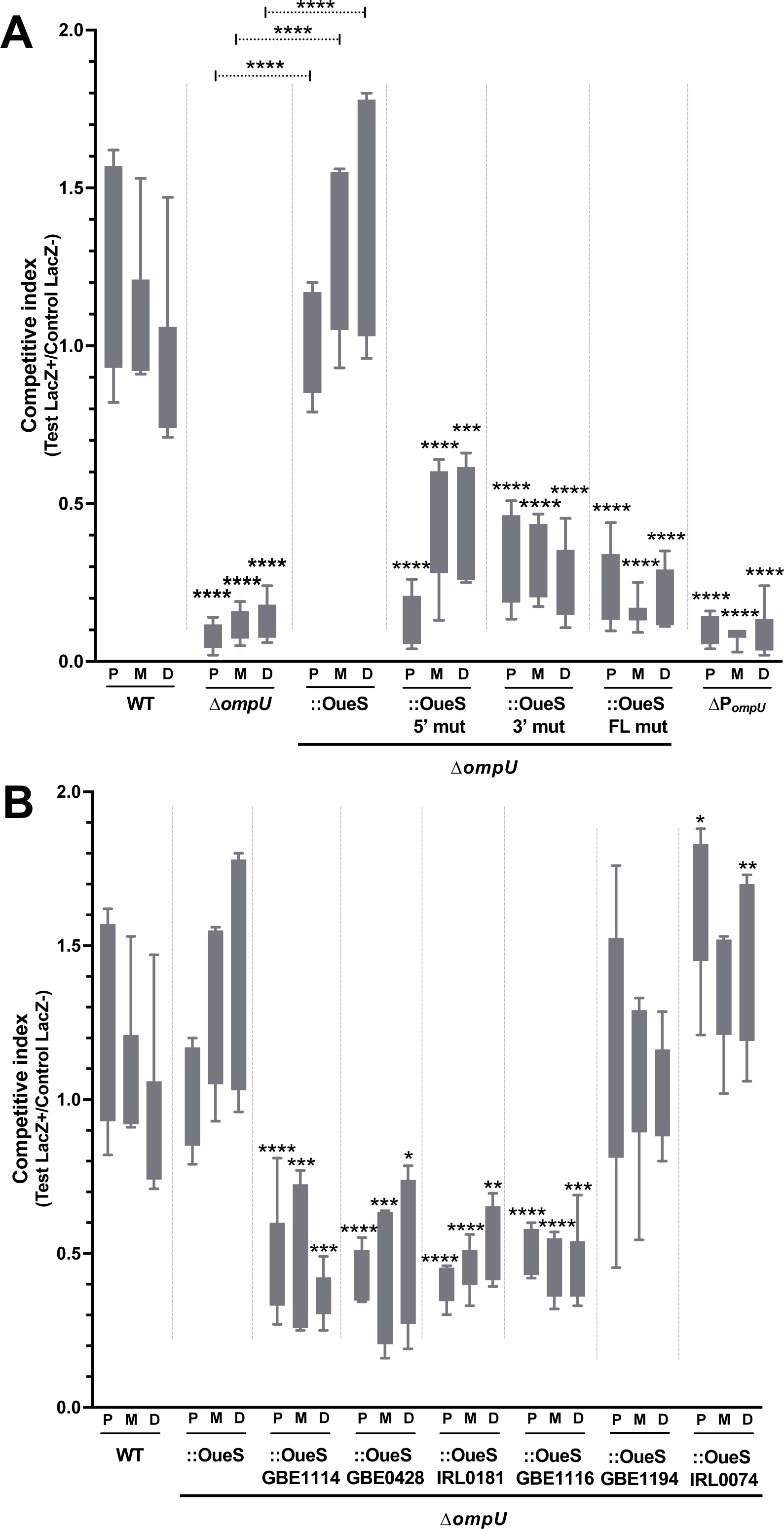
OueS is essential for intestinal colonization and the emergence of toxigenic *V. cholerae*. *V. cholerae* strains were orogastrically inoculated in 3-5-day old CD-1 mice to perform competition assays. Strains were monitored for intestinal colonization across the proximal (P), medial (M) and distal (D) sections of the small intestine. The data is represented as competitive indices (CI). (A) Expression of the toxigenic allele of OueS is required for intestinal colonization of *V. cholerae*. (B) OueS alleles from environmental strains lead to differential intestinal colonization of *V. cholerae* isogenic mutants. n ≥ 6 per strain. *p*-values indicate significance compared to the WT strain, unless otherwise specified, and are calculated by 2way ANOVA in Šidák’s multiple comparisons test; * <0.01, ** < 0.001, *** < 0.0001, **** < 0.00001.

### Modular heterogeneity in OueS drives emergence of toxigenic strains

Our data demonstrates that specific allelic variations of OueS are essential for virulence-associated phenotypes such as biofilm repression to emerge (**Fig 2D**). Recently, we examined the natural genetic variability of *ompU* and identified 41 alleles of the gene in over 1600 sequences analyzed^38^. We modeled the secondary structures of OueS from several of those representative clades and identified a diverse set of conserved OueS structures, including the one from toxigenic strains (**Fig S4A**). The 3’ region downstream of *ompU* is remarkably conserved, further corroborating the modular nature of this sRNA (**Fig S4B**). We constructed six isogenic mutant strains encoding different OueS alleles based on their structural diversity and examined their ability for intestinal colonization. Competition assays reveal a heterogenous colonization landscape mediated by the various OueS alleles. For instance, mutants expressing the OueS allele from the environmental strains GBE1194 and IRL0074 colonize the small intestine similar to WT (CI 1.05 – 1.61; **Fig 4B**). On the other hand, isogenic mutants encoding OueS from strains GBE1114, GBE0428, IRL0081, and GBE1116 exhibit a decrease in colonization (CI 0.36 – 0.54; **Fig 4B**). Overall, our data indicates that **a)** the OueS allele encoded by toxigenic strains evolved as a preadaptation to virulence since some environmental strains (e.g., GBE1194) encode alleles that also lead to successful colonization, and **b)** OueS is critical for the emergence of pathogenic potential in *V. cholerae*, as strains like GBE1114 encode alleles that lead to impaired intestinal colonization. Small RNAs modulate gene expression by direct binding to their target mRNAs^53, 54^. However, we did not find a direct correlation between the CI associated with a specific OueS allele and either its length (166-196 ntds) or its predicted secondary structures (**Fig S4**). Nonetheless, sequence variation analysis unequivocally reveals that OueS is bimodular, comprising variable *ompU*-encoded 5’ modules and a conserved 3’ one, the former directly contributing to the emergence of pathogenic *V. cholerae* from environmental populations.

## DISCUSSION

Emerging and re-emerging infectious diseases pose immense health and economic risks^55^. To date, the evolutionary drivers that facilitate certain strains within a population to emerge as pathogens remain largely unknown. This critically hinders our ability to develop effective strategies (e.g., policies, therapeutics, vaccines, etc.) against facultative pathogens in a timely fashion. For numerous bacterial pathogens, only strains confined to a single clade have emerged as pathogens, a pattern termed clonal offshoot. Pathogens that exhibit this pattern include *Vibrio parahaemolyticus, Yersinia enterocolitica*, *Hemophilus influenzae* and *Enterococcus faecalis*^56–59^, among others. The Pandemic Cholera Group in *V. cholerae* represents a quintessential example of clonal offshoot^36, 37, 60–63^. Recently, we proposed that the presence of allelic variations in their core genome might explain the confined nature of numerous facultative bacterial pathogens, including toxigenic *V. cholerae* isolates^37^.

The gene coding for the major outer membrane protein OmpU encodes these allelic variations in the form of virulence adaptive polymorphisms and plays a critical role in pathogen emergence^6^. Porins are known to harbor moonlighting functions in various bacteria, including adhesion and signaling^64, 65^, activating host signal transduction pathways^66^, or modulating immune responses^67^. Nonetheless, there is a gap in the mechanistic understanding of how porins facilitate such diverse and complex phenotypes. In this study, we discovered that OmpU encodes a sRNA in its 3’ terminus, called OueS, that explains the disparate phenotypes associated with the porin and sheds light on its association with pathogen emergence. OueS suppresses biofilm formation by limiting iron uptake and acts as a functional surrogate of the virulence regulator ToxR (**Fig 5A**). sRNAs have been previously associated with diverse processes in *V. cholerae* including virulence regulation^68–72^. Interestingly, allelic variability dictates the phenotypes associated with OueS and its role during the infection process. We propose a scenario where OueS variants circulate in environmental *V. cholerae* populations with some determining the emergence of virulence potential. The ecological drivers affecting the selection and distribution of the *ompU* alleles encoding the diverse OueS variants within *V. cholerae* environmental gene pools remain enigmatic. Defining these will shed critical light on the fundamental forces that drive the assembly and emergence of strains with pathogenic potential in a bacterial population. Furthermore, despite the vast sequence and structural divergences between the OueS alleles, two convergent colonization phenotypes emerge from them (**Fig 5B**). Determining the specific mechanisms resulting in this functional convergence will provide critical insights into the molecular drivers controlling the emergence of virulence-adaptive traits in human pathogens.

**Figure 5.**
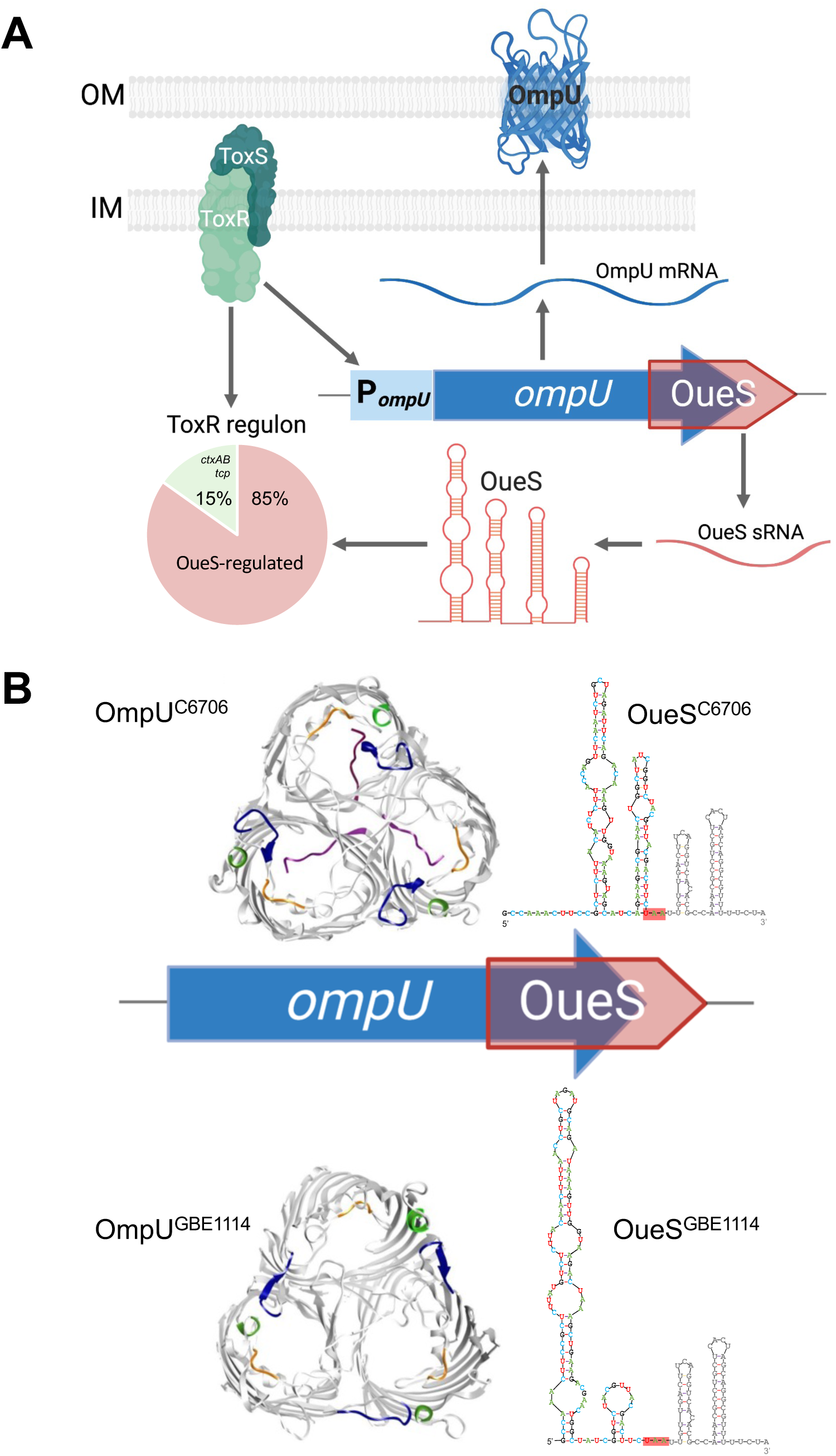
The *ompU* ORF encodes a modular porin and sRNA. (A) The virulence regulator ToxR transcriptionally modulates hundreds of genes including activating the expression of the *ompU* ORF via P*_ompU_*. The *ompU* ORF codes for the OmpU porin (blue) as well as the OueS sRNA (red) overlapping its 3’ terminus. OueS acts as a functional surrogate of ToxR by modulating the expression of 63% of its regulon. (B) *V. cholerae* encodes several alleles of *ompU*, which code for structurally and functionally distinct modular versions of OmpU and OueS. Modularity in the porin and sRNA dictates the emergence of differential virulence-associated phenotypes and, ultimately, host intestinal colonization.

Modularity has become a central concept in biology, stemming from the idea that biological systems are intricately linked networks that are typically divided into sets of strongly interacting modular subsystems^73^. Modularity spans many scales in biological systems from genes, to cells, to ecosystems and provides a cohesive mechanistic framework to explain biological complexity^74–77^. Recently, we determined that the porin OmpU contains modular protein domains that uniquely contribute to the emergence of antimicrobial resistance in certain alleles^38^. Here, we unveil a new layer in the complex modular nature of the *ompU* ORF and demonstrate that OueS also displays modular characteristics. OueS is a bimodular sRNA comprising of **a)** a variable 5’ module encoded within *ompU* that contributes to diversity and emergence potential and **b)** a highly conserved 3’ module immediately downstream from the *ompU* ORF with unknown specific function but critical for colonization (**Fig 5B**). Taken together, our results indicate that the *ompU* ORF encodes both a porin and a sRNA, and their modular nature enables the emergence of phenotypes essential for host colonization such as antimicrobial resistance or stringent biofilm regulation.

The wealth of information embedded within this ∼1 kb region of DNA is astonishing and suggests the presence of what we term “nested fractal compression” (NFC) of genetic information, which might be critical for the emergence of complex phenotypes. We contend this information is **a)** nested and fractal, as increasing the resolution **within** the unit ORF (nested) unveils successive layers of complexity (fractal), and **b)** compressed, as the overlapping of the coregulated modular porin gene with that of the modular sRNA confers a non-linear, finetuned regulation that is greater than the individual contributions of either of the two. It is tempting to think that the intricate arrangement of components within this ORF might have evolved as a modular recombinational unit that allows for the precise acquisition of different traits encoded within one quantum-like element. We propose a conceptual framework where the interactions between modular ORFs functioning as recombinational units of NFC information form hubs of robust regulatory networks driving the emergence of complex phenotypes. Overall, our findings provide mechanistic insights into the emergence of pathogenic traits in bacteria unraveling the hidden genetics associated with this process and suggest a scenario in which a limited number of modular units could explain the emergence of novel phenotypic traits and complexity in biological systems.

## MATERIALS AND METHODS

### Bacterial strains, media, and growth conditions

All *V. cholerae* strains used in this study are derivatives of the seventh pandemic El Tor biotype strain C6706, first isolated from a patient in Peru in 1961^78^. *E. coli* S17 λ-*pir* strains harboring a chromosomally integrated RP4 plasmid and *tra* genes^79^ were used for conjugation of the deletion constructs into *V. cholerae* as well as for routine cloning purposes.

The following media were used for bacterial cultivation: Luria Bertani (LB) broth (10 g l^-1^ tryptone, 5 g l^-1^ yeast extract, 5 g l^-1^ NaCl). Biofilm assays were performed in tryptone broth (tryptone 10 g l^-1^, NaCl 5 g l^-1^). AKI broth was used for virulence-inducing conditions ^48, 80^ (15 g l^-1^ Bacto^TM^ peptone, 4 g l^-1^ yeast extract, 5 g l^-1^ NaCl, 0.03% w/v sodium bicarbonate). Plates were used at a final concentration of 1.5% agar. Where necessary, the following antibiotics or supplements were used: streptomycin 1000 µg ml^-1^, carbenicillin 50 µg ml^-1^, 2,2’-bipyridyl 100 µg ml^-1^, FeCl_3_ 100 µg ml^-1^, 5-bromo-4-chloro-3-indolyl-ß-D-galactopyranoside (X-gal) 40 µg ml^-1^, and arabinose (0.01%). Reagents were procured from Sigma.

For routine culture, *E. coli* and *V. cholerae* were grown in LB broth at 37 °C with aeration (200 rpm) in a rotary shaker. Biofilm and growth curve assays were performed in 96-well flat bottom, tissue culture-treated plates (CytoOne) incubated at 37 °C.

### Cloning, vectors, and strain construction

(a) In-frame deletion mutants of *ompU*, *toxR* and *lacZ* were constructed via homologous recombination^81^. Briefly, about 500 bp fragments flanking the gene of interest to be deleted were PCR amplified, cloned as *Bgl*II-*Not*I (upstream fragment) or *Not*I-*Eco*RI (downstream fragment) in tandem into the *V. cholerae* suicide vector pKAS154^81^, and electroporated into *E. coli* S17λ-*pir*^79^. *E. coli* strains harboring the deletion constructs were then conjugated with *V. cholerae* C6706, and allelic exchange was carried out with appropriate antibiotic selection, as described^81^. Positive clones were first screened by colony PCR and then confirmed by sequencing the deletion loci from genomic DNA using flanking primers. The *ompU* promoter mutant ΔP*_ompU_,* was generated in a similar manner by amplifying the regions immediately flanking the 300 bp locus right before the *ompU* start codon, generating a deletion construct in pKAS154, followed by allelic exchange into C6706, as described above.

(b) Ectopic OueS-expressing clones and non-specific *ompU* locus clones were generated by PCR amplification from the appropriate genomic DNA template using specific primers listed in **Table S8**. The amplicons were then cloned as *Eco*RI-*Hin*dIII fragments into pBAD22. The OueS modular mutants were synthesized commercially (Twist Biosciences) and then cloned into pBAD22 as above. After introduction into *E. coli* by electroporation and sequence confirmation of the inserts, the plasmids were electroporated into the appropriate *V. cholerae* strains, followed by selection on carbenicillin-containing LB plates. For strains harboring the empty vector, plasmid pBAD22 was electroporated into the appropriate *V. cholerae* strain and selected as above. Ectopic *ompU*-expressing clones were generated in a similar manner.

(c) Chromosomal promoter fusions were constructed as described above for the deletion constructs. The ∼500 bp regions upstream of the *ompU* ORF including P*_ompU_ was* first cloned as a *Xba*I-*Sac*I fragment into pKAS154 to generate plasmid pDBS467. Subsequently, the various OueS constructs: OueS 5’-mut, 3’-mut, FL-mut fragments (commercially synthesized; Twist Biosciences), and the various OueS alleles (amplified from the respective genomic DNA) were cloned into pDBS467 downstream of P*_ompU_* as a *Sac*I-*Eco*RI fragment. Allelic exchange to replace the *ompU* ORF was then performed as described above.

All deletion and complementation constructs in *E. coli* were Sanger sequenced before introduction into *V. cholerae*. Genomic DNA was isolated from all *V. cholerae* deletion and chromosomal fusion strains, and the region(s) of interest amplified using flaking primers for confirmation by Sanger sequencing.

### RNA isolation and transcriptome studies

For Northern blots and RACE studies, RNA was isolated using the hot-phenol extraction method^82–84^. Briefly, the cell pellets harvested under appropriate growth conditions were resuspended in Solution A (0.5% SDS, 20 mM sodium acetate, 10 mM EDTA), followed immediately by the addition of 500 µL of acid-phenol:chloroform (Ambion) preheated to 65 °C and vortexed. After incubation at 65 °C for 10 minutes, the samples were centrifuged at 21,000 x g for 10 minutes and the aqueous layer reextracted twice with acid-phenol:chloroform. The aqueous phase was then extracted twice with chloroform and precipitated overnight at -80 °C with 2.5 volumes of ice-cold absolute ethanol. The total RNA was then pelleted and washed with cold 70% ethanol, pellets dried and resuspended in 100 µL RNase-free water containing 40 U murine RNase inhibitor (NEB). Following quantification by Nanodrop, 50 µg of the total RNA was DNase-treated (Ambion RNase-free DNase) following manufacturer protocols. The RNA was reextracted as above and precipitated with 1/10^th^ volume 3M sodium acetate and absolute ethanol followed by a 70% ethanol wash. Final resuspension of the dried RNA pellets was in 50 µL of RNase-free water containing RNase inhibitor. Size fractionation for RACE studies was performed by running 10 µg of hot phenol-isolated RNA on a 1.25% MOPS agarose gel (Lonza) in 1x MOPS buffer with the Riboruler low range RNA ladder (Thermo) for size determination. Gel extraction was performed using the Zymoclean gel RNA recovery kit (Zymoresearch).

For transcriptome studies, RNA was isolated using a commercial kit (DirectZol RNA Miniprep kit, Zymo Research) following manufacturer instructions. DNase treatment, rRNA depletion, library preparation and Illumina Hi-Seq sequencing was performed at SeqCenter, LLC (Pittsburgh, PA). CLC Genomics Workbench (Qiagen) was used for transcriptome data processing. Briefly, paired Illumina reads were mapped to the El Tor reference genome (GenBank AE003852.1- Chromosome I; AE003853.1- chromosome II) using the following settings to generate the transcripts per million bases (TPM) values : mismatch cost 2, insertion / deletion cost 3, length / similarity fraction 0.8, global strand specific alignment. Subsequent pairwise analysis between two datasets of interest (absolute fold change ≥ 2.0 filtered on average expression for FDR correction ≤ 0.01) were used for downstream analysis. Functional enrichment analysis of the transcriptome data was performed by the Panther Overrepresentation Test against the Biological processes dataset of the GO Ontology Database (Released 07-01-2022)^85, 86^.

### Biofilm assays

The ability of the strains to form biofilms on surfaces was determined using the crystal violet-based assays in 96-well polystyrene plates, optimized from published protocols^87^. Briefly, the strains from LB plates were grown in tryptone broth with carbenicillin for 14-16 hours at 37 °C with aeration. The cultures were then diluted 1:500 in fresh tryptone broth supplemented with carbenicillin and arabinose and 200 µL dispensed in wells of the 96-well plates. Ferric chloride (100 µM) or 2, 2’-bipyridyl (100 µM) were supplemented, as needed, before dispensing the cultures. After static incubation for 24 hours at 37 °C, the cultures were poured off and plates gently washed three times with water to remove unattached cells. The surface attached cells were then stained with 225 µL per well of crystal violet (0.1% in water) for 20 minutes at room temperature and washed thoroughly with water. The attached stain was subsequently eluted with 225 µL per well of 50% acetic acid for 15 minutes and quantified by reading the absorbance in a Tecan Sunrise plate reader at 595 nm.

To examine the effect of iron on biofilm formation, TB media was supplemented with either 100 µM FeCl_3_ (MP Biomedicals) for iron repletion or 100 µM 2, 2’-bipyridyl (Sigma) to deplete residual iron. Growth and processing of biofilms was performed as described above.

### Northern blot assays

For northern blot assays, total RNA isolated from LB-grown cultures (OD ∼2.0) using the hot-phenol method were used. Downstream processing was performed by Tribioscience Inc. (Sunnyvale, CA). Briefly, 8 µg RNA per lane was first resolved on a 1.2% agarose gel (Lonza) and transferred for 20 hours to a nylon membrane by capillary action. Airdried membranes were then UV cross-linked (254 nm; 120mJ/cm^2^) and hybridized to biotin-conjugated probes using the NorthernMax-Gly system following manufacturer protocols (Invitrogen). The following 5’-biotinylated probes were synthesized commercially (IDT): *ompU* 5’ end (DB156: tcacttcaccgtaagtgccaccgataccagcgtag), *ompU* 3’ end (DB246: gtaacgtagaccgatagccagttcgtcttctgatgctact), 5S rRNA gene (DB158: ttcgtttcacttctgagttcgg). Probe detection was performed using the IRDye 800cw Streptavidin (LI-COR, Inc.) and imaged on LI-COR Odyssey digital imager.

### Rapid Amplification of cDNA Ends (RACE)

The 5’ and 3’ termini of the sRNA OueS were determined by the RACE assay (SMARTer RACE 5’/3’ kit, Takara Bio USA), adapted for non-poly-A-tailed RNA. Kit components were used, unless otherwise mentioned. Briefly, total RNA was separated on a 1% MOPS agarose gel and the 100-500 ntd region was gel extracted (Zymoclean Gel RNA recovery kit). This size fractionated RNA was used as the template for both 5’ and 3’ RACE studies.

For 5’ RACE, first strand cDNA synthesis was performed by random priming followed by annealing the SMARTer IIA oligonucleotide to the 5’ ends of the newly synthesized cDNA fragments. This was followed by PCR amplification using a forward primer specific to the 5’-attached oligonucleotide and a reverse primer within the *ompU* gene (DB196: 5’ GATTACGCCAAGCTTtcgtaacgtagaccgatagccagttcgtct 3’). The fragments were then cloned non-directionally using the A-overhangs generated during the final PCR step and electroporated into *E. coli* Stellar Competent cells (Clontech) and selected on carbenicillin plates. Plasmids from positive clones were then sequenced to identify the 5’ terminus of OueS. For 3’ RACE, the size-fractionated total RNA was poly-A-tailed using the *E. coli* Poly(A) polymerase (New England Biolabs). cDNA was then synthesized using a primer containing a complementary poly-T sequence and a 5’ overhang (DB164). The newly synthesized cDNA was then used as template for a second strand synthesis using an *ompU*-specific primer that binds inside the 3’ end of the *ompU* ORF (DB195: 5’ GATTACGCCAAGCTTccgctcttacatctcttaccagttcaatctgctag 3’). The amplicon was then purified, and PCR amplified using the 3’ *ompU*-specific primer and a primer targeting the 5’ overhang of the poly-T primer (DB164) used in the first step of the process. The resulting amplicons were then cloned non-directionally into vectors with a T-overhang and electroporated into *E. coli*. Subsequently, plasmids from positive clones were sequenced to define the 3’ terminus of OueS.

### Ethics statement

Three-to-five-day-old CD-1 ® IGS mice (Strain code 022; Charles River Laboratories) of both genders were randomly selected from mixed litters for all animal experiments, in accordance with the guidelines and protocols approved by the Institutional Animal Care and Use Committee (IACUC) at the University of Central Florida (#PROTO202000124). Infant mice were housed with the mothers, monitored under the care of full-time staff, and weaned 2 hours prior to infection.

### Infant mouse competition assays

Intestinal colonization competition assays in 3-5-day-old infant mice were performed with LB-grown *lacZ*^+^ cells of the WT, deletion mutants and OueS alleles competed against an isogenic WT strain harboring a *lacZ* deletion, essentially as described^37, 51, 52^. The colonization efficiencies of the different strains in the proximal, medial, and distal portions of the small intestine harvested at ∼22 hours post infection are represented as competitive indices^51^. Briefly, strains were inoculated from fresh LB plates into LB broth, incubated for 12 hours at 37 °C with aeration, and diluted 1:1000 fold in fresh LB broth. The test (*lacZ*^+^) and control (*lacZ*^-^) strains were mixed 1:1 to obtain ∼10^6^ CFU/mL. Fifty microliters of the bacterial mixture, corresponding to ∼10^5^ CFUs, were then used to intragastrically inoculate 3-5-day-old infant mice and maintained at 30 °C for 22 hours. The exact input CFU numbers were determined by plating dilutions on LB-Sm-X-Gal plates. Approximately 22 hours post infection, the mice were euthanized, their small intestines divided into the proximal, medial, and distal sections and transferred separately into 4 mL LB-10% glycerol stored on ice. The samples were then processed for CFU by homogenizing the tissues, serially diluting in 1x PBS (pH 7.4) and plating appropriate dilutions on LB-Sm-X-Gal plates. *In vitro* competition experiments were performed in parallel by inoculating 50 µL of the input CFU in LB-Sm broth and incubating for 22 hours at 37 °C with aeration. Samples were then serially diluted and plated as for the *in vivo* samples above. Results are presented as the competitive index (CI), which is the ratio of the intestinal *lacZ*^+^ CFU (blue colonies) to *lacZ*^-^ CFU (white colonies) normalized to the input CFU ratio and *in vitro* competition CFU ratio.

### Growth Kinetics Under Iron Limitation

Overnight LB-grown cultures were diluted 1:1000 in LB broth with or without supplementation with FeCl_3_ (iron-replete) or 2,2’-bipyridyl (iron-deplete). 200 µL aliquots of the cultures dispensed in flat-bottom 96-well plates (tissue culture-treated, CytoOne) were then monitored for A_595_ every 30 minutes for 24 hours at 37 °C in a Tecan Sunrise plate reader. Each biological replicate had at least six technical replicates and the data for each condition were averaged across technical and biological replicates corrected to the baseline (LB) A_595_ values.

### *In silico* analyses

The secondary structures of the WT and mutant versions and alleles of OueS were determined using mFold on the UNAFold web server (RNA Folding Form V2.3) using default parameters.

### Statistical analyses

All data were analyzed for statistical significance using the two-tailed unpaired T-test on Graphpad Prism 9 (V 9.5.1), except for the transcriptome data that was analyzed within the CLC Genomics Workbench (Qiagen). Unless otherwise indicated, all experiments were performed with at least five independent biological replicates (n ≥ 5) and the aggregate data (mean ± s.d.) are shown.

## ACKNOWLEDGEMENTS

This work was supported by a National Science Foundation (NSF) CAREER award (#2045671) and a Burroughs Wellcome Investigator in the Pathogenesis of Infectious Disease (#1021977) to SAM.

## AUTHOR CONTRIBUTIONS

SAM designed research. DB performed research. DB and SAM analyzed data and wrote the manuscript.

## SUPPLEMENTARY MATERIAL

**Supplementary Figure 1: OueS mutants and alleles**. (A) Sequence conservation of the *ompU* ORF encompassing the OueS 5’ locus before (WT) and after codon wobble mutations (OueS-5’ mut, OueS-3’ mut and OueS-FL mut) are shown to demonstrate an unaltered protein coding sequence. The 166 ntd OueS sequence from (B) C6706 (ΔG -40.0), (C) OueS 5’-mut (ΔG -40.7), (D) OueS 3’-mut (ΔG -24.6), and (E) OueS full length-mut (ΔG -26.5) were used to generate their predicted secondary structures on mFold (unafold.org). The structures requiring the least free energy (dG) were selected for comparison. Mutations significantly alter the structures of the OueS variants (C, D, E), compared to the WT OueS^C6706^ (B) from Fig 2C, shown here for comparison. (F) Predicted secondary structure of OueS from the environmental strain GBE1114 and (G) pairwise alignment with OueS^C6706^ highlight variations arising from sequence differences in the sRNA region within the *ompU* ORF, resulting in a bimodular sRNA.

**Supplementary Figure 2: OueS regulates iron uptake in biofilms, but not planktonic growth**. The planktonic growth patterns of the WT, Δ*ompU* cells ectopically expressing either the unmutated or mutated versions of OueS in (A) tryptone broth and under (B) iron-replete (100 µM FeCl_3_) or (C) iron-depleted (100 µM 2,2’-bipyridyl) conditions. Growth patterns are similar between the strains in each condition irrespective of iron concentration.

**Supplementary Figure 3: OueS is expressed from the *ompU* promoter.** Mutant strains encoding varying lengths of the region upstream of OueS were monitored for biofilm formation. All strains repressed biofilm formation similar to the ectopic OueS expressing clone (Fig 2D), with the exception of a deletion strain for P*_ompU_*. *p-*value: *** ≤ 0.0001.

**Supplementary Figure 4: Variation within the OueS alleles.** *ompU* alleles were analyzed for sequence variation^38^ and OueS sequences were extracted from representative clades for secondary structure prediction (www.unafold.org). (A) The alleles match one of the eight predicted secondary structures. (B) Sequence alignment of the OueS loci from representative alleles underlines the variability that contributes to the secondary structure variation seen in A.

## Supplementary Tables

**Table S1.** Differentially regulated genes by OmpU compared to wild-type C6706 and OmpU complementation from biofilm growth.

**Table S2.** Differentially expressed genes in biofilms in ompU and rpoE mutants compared to WT C6706.

**Table S3:** OueS-regulated biofilm genes obtained by comparing expression values of WT and Δ*ompU*-pOueS biofilm grown cells to their expression in the ompU mutant.

**Table S4:** Differentially expressed genes in the ompU and rpoE mutants compared to WT C6706 when grown in virulence inducing AKI conditions.

**Table S5:** Differentially expressed 453 genes in the *ompU* mutant whose expression is rescued by OueS compared to WT C6706 when grown in virulence inducing AKI conditions.

**Table S6:** Gene expression in WT and ΔtoxR-pOueS strains from biofilm growth. Absolute fold changes (> 2.0) are relative to expression in the ΔtoxR strain.

**Table S7:** Overlap between the ToxR and OueS virulence regulons. Subset of the ToxR AKI regulon that comprises the OueS regulated genes (yellow highlight) identified by an absolute ratio of ≥ 2.0 between the two datasets. Expression values are relative to the WT strain.

**Table S8:** Strains, vectors and primers used in this study.

