## Supplementary figures and images for "Modular small RNA drives pathogen emergence"

### Fig S1

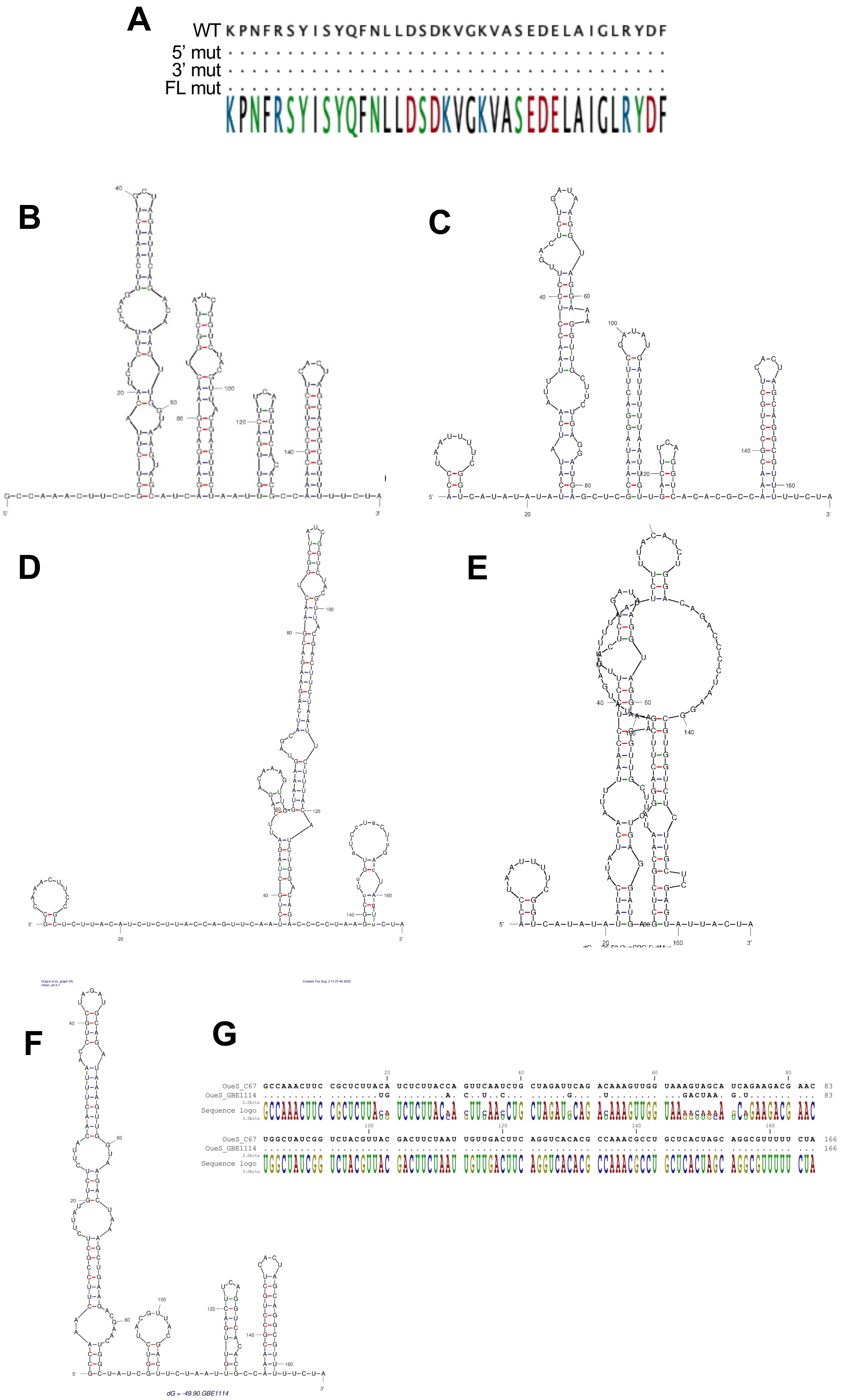

### Fig S2

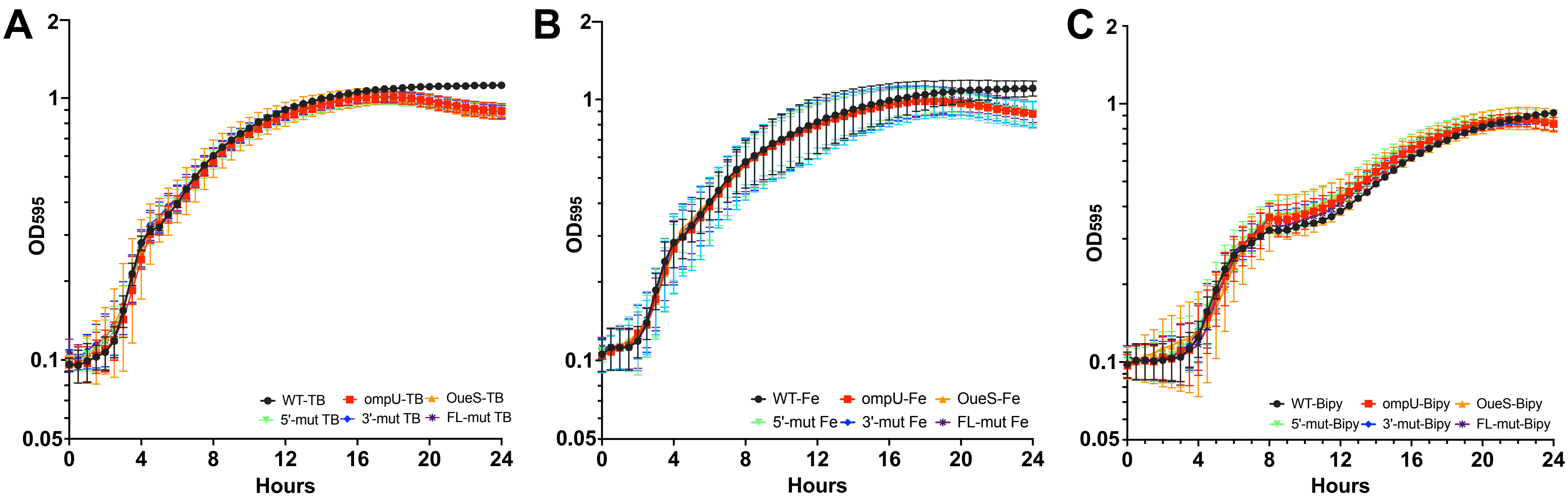

### Fig S3

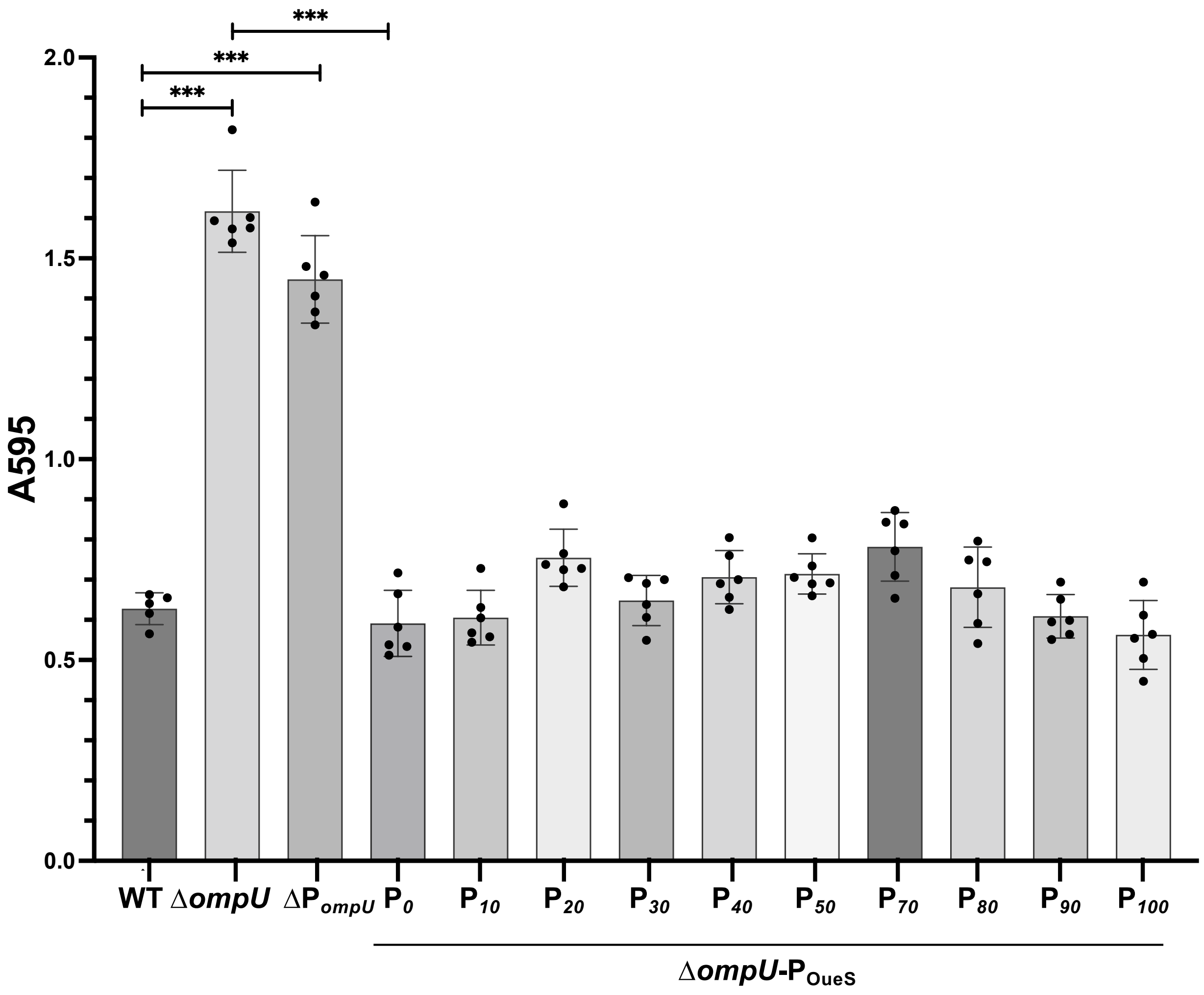

**Supplementary Figure 5**

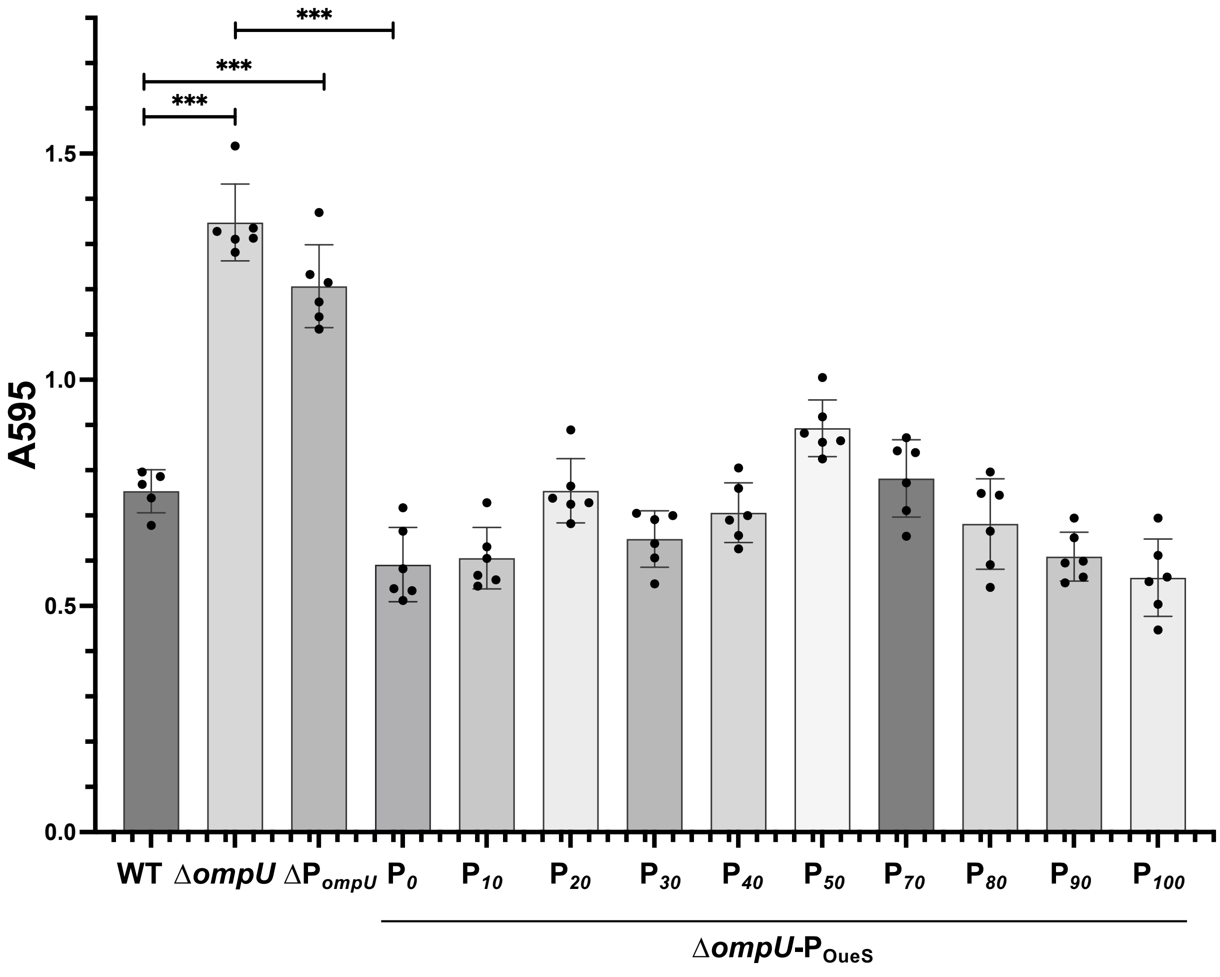

### Fig S4

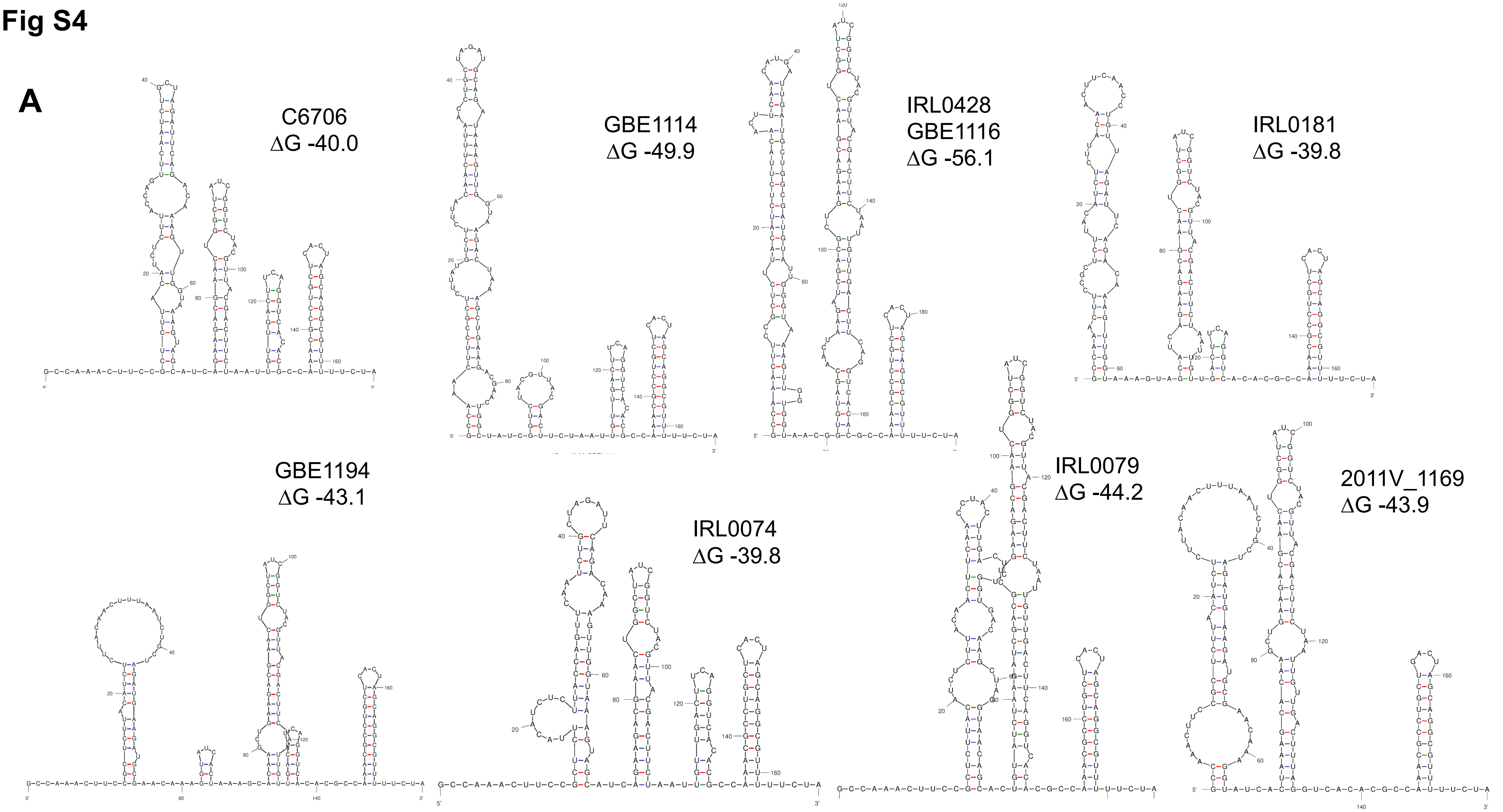

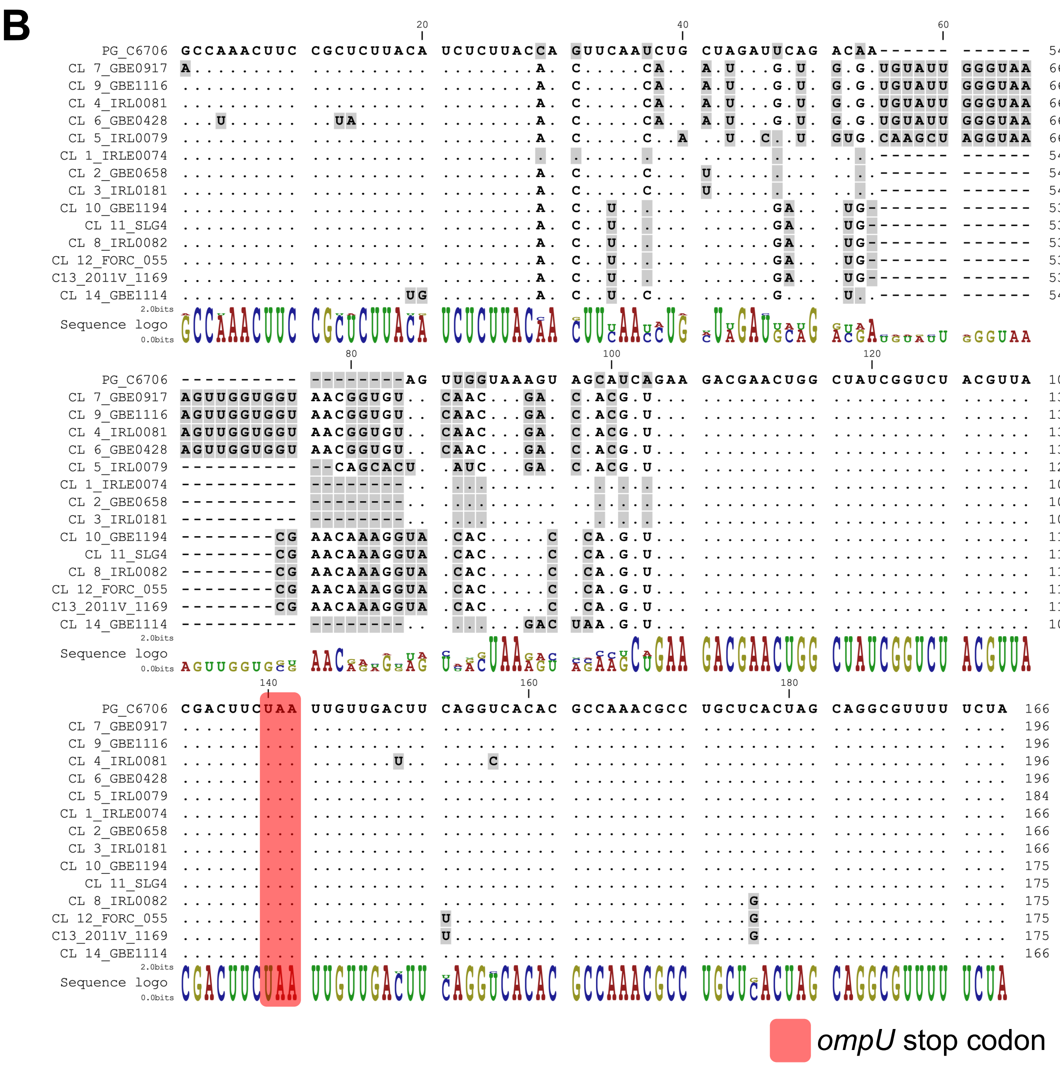
