## Supplementary material for "Modular small RNA drives pathogen emergence": Table S1

**Table S1.** Differentially regulated genes by OmpU compared to wild-type C6706 and OmpU complementation from biofilm growth.

| Genes activated by OmpU | | | | | |
|---|---|---|---|---|---|
| **Locus tag** | **Function** | **Δ*ompU*/WT** | | **Δ*ompU*-pOmpU/WT** | |
| | | **FC** | **FDR *p*-value** | **FC** | **FDR *p*-value** |
| VC0176 | transcriptional regulator, putative | -2.8 | 7.1E-03 | -1.0 | 9.3E-01 |
| VC0177 | hypothetical protein | -2.6 | 9.1E-05 | -1.0 | 9.9E-01 |
| VC0178 | patatin-related protein | -2.2 | 1.7E-04 | -1.1 | 8.7E-01 |
| VC0179 | hypothetical protein | -2.5 | 9.0E-06 | -1.2 | 5.0E-01 |
| VC0180 | conserved hypothetical protein | -2.3 | 1.2E-06 | -1.3 | 1.2E-01 |
| VC0181 | conserved hypothetical protein | -2.2 | 4.4E-03 | -1.2 | 2.9E-01 |
| VC0379a | phage shock protein G | -2.2 | 4.0E-05 | -2.1 | 1.4E-04 |
| VC0494 | conserved hypothetical protein | -2.9 | 1.7E-04 | 1.4 | 1.6E-01 |
| VC0495 | conserved hypothetical protein | -2.1 | 9.6E-03 | -1.2 | 5.2E-01 |
| VC0502 | type IV pilin, putative | -2.7 | 1.1E-06 | 1.1 | 7.7E-01 |
| VC0503 | conserved hypothetical protein | -3.3 | 7.9E-12 | -1.3 | 1.1E-01 |
| VC0504 | hypothetical protein | -2.5 | 7.7E-05 | -1.7 | 2.8E-03 |
| VC0505 | hypothetical protein | -2.9 | 1.6E-04 | -1.7 | 1.0E-02 |
| VC0506 | hypothetical protein | -2.6 | 5.0E-04 | -1.2 | 4.6E-01 |
| VC0507 | hypothetical protein | -4.8 | 2.2E-04 | -1.1 | 8.1E-01 |
| VC0566 | protease DO | -2.3 | 9.7E-06 | -1.7 | 2.4E-03 |
| VC0633 | outer membrane protein OmpU | -15766.6 | 0.0E+00 | -615.6 | 0.0E+00 |
| VC0711 | clpB protein | -2.0 | 5.2E-04 | -2.0 | 3.2E-05 |
| VC0817 | transposase, putative | -2.1 | 2.4E-03 | 1.3 | 2.5E-01 |
| VC0818 | pseudogene; within VPI-I; unknown function | -2.2 | 5.2E-03 | 1.1 | 7.1E-01 |
| VC0825 | toxin co-regulated pilus biosynthesis protein I | -2.1 | 4.5E-03 | 1.3 | 1.2E-01 |
| VC0856 | dnaJ protein | -2.1 | 2.4E-05 | -1.6 | 4.7E-03 |
| VC0886 | hypothetical protein | -2.3 | 2.7E-05 | -1.7 | 2.6E-07 |
| VC0985 | heat shock protein HtpG | -2.1 | 6.5E-05 | -2.1 | 2.1E-05 |
| VC1217 | conserved hypothetical protein | -2.2 | 3.5E-03 | -1.3 | 4.0E-01 |
| VC1316 | chemotaxis protein CheY, putative | -2.3 | 1.1E-04 | -1.8 | 1.6E-03 |
| VC1422 | sodium/alanine symporter | -2.2 | 2.5E-05 | -1.1 | 7.1E-01 |
| VC1472 | hypothetical protein | -4.3 | 1.5E-04 | 3.1 | 9.0E-05 |
| VC1562 | beta-lactamase-relat ed protein | -2.2 | 4.2E-03 | -1.3 | 3.4E-01 |
| VC1675 | multidrug resistance protein, putative | -2.0 | 1.1E-03 | -1.5 | 2.0E-02 |
| VC1676 | phage shock protein C | -2.3 | 3.4E-05 | -1.8 | 7.8E-05 |
| VC1677 | phage shock protein B | -2.2 | 6.7E-04 | -2.4 | 3.1E-09 |
| VC1678 | phage shock protein A | -2.6 | 4.5E-06 | -1.9 | 6.0E-05 |
| VC1762 | hypothetical protein | -2.1 | 1.1E-03 | -1.3 | 1.3E-01 |
| VC1763 | chemotaxis protein MotB-related protein | -2.8 | 1.1E-05 | -1.1 | 7.8E-01 |
| VC1774 | conserved hypothetical protein | -2.0 | 3.7E-03 | 1.9 | 1.0E-03 |
| VC1786 | DNA repair protein RadC, putative | -3.5 | 1.3E-03 | -1.0 | 9.9E-01 |
| VC1793 | hypothetical protein | -2.9 | 5.3E-03 | 1.2 | 5.9E-01 |
| VC1802 | hypothetical protein | -3.5 | 3.2E-03 | -1.6 | 2.7E-01 |
| VC2149 | hypothetical protein | -2.0 | 8.1E-04 | -2.0 | 4.2E-06 |
| VC2663 | molecular chaperone groEL_1 | -2.1 | 8.1E-04 | -2.1 | 2.8E-05 |
| VC2664 | chaperonin, 60 Kd subunit, groES_1 | -2.0 | 1.1E-03 | -1.5 | 4.0E-02 |
| VCA0275 | IS1004 transposase | -5.0 | 8.7E-03 | 1.3 | 6.6E-01 |
| VCA0298 | hypothetical protein | -2.2 | 9.0E-03 | -1.5 | 7.1E-03 |
| VCA0308 | deoxyguanosinetripho sphatetriphosphohydrolase-rela ted protein | -2.1 | 9.0E-05 | -1.6 | 3.9E-06 |
| VCA0310 | hypothetical protein | -2.1 | 1.1E-03 | -1.8 | 8.6E-03 |
| VCA0316 | acetyltransferase, putative | -2.5 | 4.0E-04 | -2.0 | 6.0E-03 |
| VCA0336 | hypothetical protein | -3.2 | 9.5E-06 | -2.2 | 5.9E-05 |
| VCA0339 | hypothetical protein | -3.1 | 4.0E-03 | -1.9 | 1.6E-01 |
| VCA0344 | hypothetical protein | -2.1 | 1.6E-04 | -1.2 | 3.0E-01 |
| VCA0346 | H-REV 107-related protein | -3.1 | 2.4E-10 | -1.8 | 5.6E-04 |
| VCA0354 | hypothetical protein | -2.0 | 2.1E-03 | -1.3 | 3.2E-01 |
| VCA0355 | conserved hypothetical protein | -2.5 | 7.8E-06 | -1.5 | 6.0E-02 |
| VCA0356 | hypothetical protein | -2.1 | 3.5E-04 | -1.3 | 5.0E-02 |
| VCA0356a | hypothetical protein | -2.3 | 6.4E-03 | -1.5 | 5.0E-02 |
| VCA0357 | hypothetical protein | -3.0 | 3.0E-03 | -2.2 | 6.9E-03 |
| VCA0358 | hypothetical protein | -2.0 | 4.5E-03 | -1.2 | 4.0E-01 |
| VCA0367 | hypothetical protein | -2.1 | 2.8E-04 | -1.1 | 6.8E-01 |
| VCA0382a | membrane protein | -2.0 | 1.0E-02 | -1.7 | 2.0E-02 |
| VCA0386 | hypothetical protein | -2.7 | 1.5E-06 | -1.9 | 6.7E-06 |
| VCA0397 | hypothetical protein | -2.4 | 1.7E-04 | -1.4 | 1.0E-01 |
| VCA0398 | hypothetical protein | -2.7 | 1.2E-04 | -2.4 | 1.0E-05 |
| VCA0409 | hypothetical protein | -2.2 | 6.2E-04 | -1.7 | 1.0E-02 |
| VCA0410 | hypothetical protein | -2.0 | 5.3E-03 | -1.5 | 8.0E-02 |
| VCA0415 | conserved hypothetical protein | -3.1 | 1.5E-03 | -1.7 | 6.0E-02 |
| VCA0419 | hypothetical protein | -2.6 | 2.0E-05 | -2.1 | 1.0E-03 |
| VCA0420 | hypothetical protein | -2.5 | 6.1E-04 | -1.5 | 1.9E-01 |
| VCA0428 | hypothetical protein | -2.2 | 1.6E-05 | -1.4 | 7.0E-02 |
| VCA0431 | hypothetical protein | -3.0 | 4.7E-08 | -1.7 | 5.7E-03 |
| VCA0443 | lipoprotein Blc | -2.4 | 1.0E-04 | -1.2 | 3.3E-01 |
| VCA0446 | haemagglutinin | -3.6 | 6.6E-11 | -1.6 | 2.4E-03 |
| VCA0447 | haemagglutinin associated protein | -2.4 | 3.2E-05 | -1.5 | 2.0E-02 |
| VCA0448 | hypothetical protein | -3.2 | 2.9E-07 | -2.2 | 1.2E-05 |
| VCA0449 | hypothetical protein | -4.7 | 9.7E-06 | -1.2 | 5.4E-01 |
| VCA0450 | hypothetical protein | -2.7 | 8.3E-03 | -1.2 | 5.6E-01 |
| VCA0451 | hypothetical protein | -2.7 | 9.4E-03 | -1.4 | 2.9E-01 |
| VCA0455 | conserved hypothetical protein | -2.1 | 1.8E-04 | -1.1 | 6.1E-01 |
| VCA0465 | hypothetical protein | -2.1 | 5.4E-03 | 1.2 | 4.2E-01 |
| VCA0480 | hypothetical protein | -2.2 | 2.5E-03 | -1.7 | 7.4E-03 |

| | | | | | |
|---|---|---|---|---|---|
| VCA0508 | transposase OrfAB, subunit B | -2.8 | 4.6E-07 | -1.0 | 9.0E-01 |
| VCA0519 | fructose repressor | -2.0 | 9.4E-04 | -1.6 | 1.0E-03 |
| VCA0690 | acetyl-CoA acetyltransferase | -2.1 | 1.5E-04 | -2.7 | 3.1E-10 |
| VCA0691 | acetoacetyl-CoA reductase | -2.2 | 6.0E-06 | -2.5 | 5.1E-13 |
| VCA0697 | sensory box/GGDEF family protein | -2.0 | 4.8E-04 | -1.8 | 2.3E-05 |
| VCA1115 | ParA family protein | -2.0 | 2.5E-04 | -1.3 | 2.0E-01 |
| 16Sh | 16S ribosomal RNA | -2.3 | 2.1E-05 | 1.3 | 1.8E-01 |
| tRNA-Arg-1 | tRNA biosynthesis | -2.4 | 6.4E-03 | 2.6 | 2.2E-09 |
| tRNA-Gly-8 | tRNA biosynthesis | -2.1 | 2.4E-03 | 1.6 | 8.5E-03 |
| tRNA-Met-8 | tRNA biosynthesis | -2.9 | 1.8E-04 | 1.1 | 7.0E-01 |
| **Genes repressed by OmpU** | | | | | |
| **Locus tag** | **Function** | **Δ*ompU*/WT** | | **Δ*ompU*-pOmpU/WT** | |
| | | **FC** | **FDR *p*-value** | **FC** | **FDR *p*-value** |
| VC0017 | hypothetical protein | 2.7 | 1.1E-05 | -1.5 | 2.0E-02 |
| VC0062 | thiamin-phosphate pyrophosphorylase | 2.9 | 4.3E-03 | 4.8 | 2.6E-26 |
| VC0063 | thiF protein | 3.1 | 1.0E-04 | 4.8 | 2.1E-27 |
| VC0065 | thiG protein | 3.3 | 1.2E-06 | 4.4 | 3.2E-26 |
| VC0066 | thiH protein | 2.5 | 4.6E-04 | 3.2 | 7.8E-17 |
| VC0091 | O-methyltransferase- related protein | 3.7 | 4.1E-07 | 1.2 | 3.0E-01 |
| VC0107 | hypothetical protein | 8.4 | 1.6E-16 | 1.8 | 4.5E-04 |
| VC0157 | alkaline serine protease | 4.5 | 2.0E-12 | 2.2 | 1.2E-05 |
| VC0199 | hemolysin secretion ATP-binding protein,putative | 3.4 | 3.1E-05 | 2.0 | 2.7E-03 |
| VC0200 | iron(III) compound receptor | 6.5 | 6.9E-11 | 3.4 | 7.4E-11 |
| VC0202 | iron(III) ABC transporter, periplasmiciron-compound-bindi ng protein | 2.5 | 4.1E-03 | 2.4 | 8.1E-05 |
| VC0284 | putative outer membrane receptor | 3.1 | 1.3E-08 | 2.5 | 6.7E-09 |
| VC0364 | bacterioferritin-ass ociated ferredoxin | 3.3 | 8.9E-06 | 1.9 | 9.6E-05 |
| VC0474 | iron-regulated virulence regulatory proteinIrgB | 3.3 | 4.5E-05 | 1.3 | 3.8E-01 |
| VC0475 | iron-regulated outer membrane virulence protein,TonB receptor family | 8.2 | 2.5E-37 | 6.5 | 2.0E-38 |
| VC0608 | iron(III) ABC transporter, periplasmiciron-compound-binding protein | 4.8 | 1.2E-12 | 2.3 | 1.5E-06 |
| VC0676 | nptA protein | 4.5 | 2.7E-15 | 4.0 | 1.8E-18 |
| VC0719 | DNA-binding response regulator PhoB | 2.4 | 3.0E-05 | 2.8 | 8.3E-16 |
| VC0747 | conserved hypothetical protein | 2.2 | 2.7E-04 | 1.1 | 6.0E-01 |
| VC0748 | aminotransferase NifS, class V | 2.1 | 5.6E-03 | 1.2 | 3.7E-01 |
| VC0749 | NifU-related protein | 2.3 | 6.0E-04 | 1.4 | 4.0E-02 |
| VC0754 | conserved hypothetical protein | 2.2 | 8.3E-04 | 1.7 | 2.7E-03 |
| VC0771 | vibriobactin-specifi c isochorismatase | 8.0 | 3.6E-13 | 2.8 | 1.4E-10 |
| VC0773 | vibriobactin-specifi c isochorismate synthase | 5.0 | 1.5E-05 | 2.9 | 8.0E-04 |
| VC0774 | vibriobactin-specifi c2,3-dihydro-2,3-dihydroxybenz oate dehydrogenase | 5.8 | 1.6E-14 | 2.0 | 5.5E-03 |
| VC0775 | vibriobactin synthesis protein, putative | 2.7 | 6.6E-05 | 1.4 | 1.0E-01 |
| VC0776 | ferric vibriobactin ABC transporter, periplasmicferric vibriobactin-binding protein | 4.2 | 2.6E-11 | 2.0 | 8.8E-04 |
| VC0777 | ferric vibriobactin ABC transporter, permeaseprotein | 3.5 | 1.1E-05 | 1.6 | 1.4E-01 |
| VC0778 | ferric vibriobactin ABC transporter, permeaseprotein | 3.1 | 1.2E-05 | 1.2 | 5.6E-01 |
| VC0820 | ToxR-activated gene A protein | 4.0 | 5.9E-09 | 1.2 | 4.1E-01 |
| VC0821 | hypothetical protein | 3.6 | 3.5E-13 | 1.2 | 3.7E-01 |
| VC0823 | hypothetical protein | 2.8 | 9.0E-09 | 1.5 | 1.0E-02 |
| VC0834 | toxin co-regulated pilus biosynthesis protein S | 2.9 | 7.6E-04 | 1.3 | 4.3E-01 |
| VC0835 | toxin co-regulated pilus biosynthesis protein T | 2.8 | 1.4E-06 | 1.1 | 7.1E-01 |
| VC0838 | TCP pilus virulence regulatory protein | 4.6 | 2.4E-05 | 2.1 | 1.3E-01 |
| VC0928 | hypothetical protein | 2.2 | 2.3E-03 | 6.7 | 8.2E-23 |
| VC0929 | hypothetical protein | 5.8 | 6.0E-27 | 1.9 | 1.4E-04 |
| VC0930 | hemolysin-related protein | 2.7 | 6.8E-03 | 3.1 | 1.9E-04 |
| VC0931 | conserved hypothetical protein | 3.1 | 3.1E-04 | 2.3 | 3.6E-04 |
| VC0932 | hypothetical protein | 4.8 | 4.4E-07 | 4.6 | 1.1E-07 |
| VC0935 | hypothetical protein | 3.2 | 1.2E-03 | 7.2 | 6.0E-12 |
| VC0936 | polysaccharide export-related protein | 2.4 | 9.5E-03 | 4.8 | 3.0E-10 |
| VC0937 | exopolysaccharide biosynthesis protein,putative | 2.3 | 3.9E-03 | 3.9 | 3.6E-12 |
| VC0938 | hypothetical protein | 2.5 | 4.5E-04 | 3.6 | 4.2E-12 |
| VC0940 | conserved hypothetical protein | 2.1 | 3.7E-03 | 1.2 | 2.8E-01 |
| VC0972 | porin, putative | 2.3 | 3.2E-04 | 1.0 | 9.8E-01 |
| VC1029 | GGDEF family protein | 2.3 | 5.5E-03 | 1.4 | 1.0E-01 |
| VC1051 | hypothetical protein | 2.1 | 9.6E-03 | 3.5 | 9.7E-18 |
| VC1061 | cysteine synthase/cystathionine beta-synthasefamily protein | 3.4 | 7.6E-09 | 2.0 | 1.9E-06 |
| VC1154 | hypothetical protein | 2.0 | 2.7E-04 | 1.1 | 8.1E-01 |
| VC1204 | formiminoglutamase | 2.8 | 9.0E-05 | 1.8 | 2.0E-02 |
| VC1264 | iron-regulated protein A, putative | 2.2 | 1.6E-04 | 1.9 | 5.5E-05 |
| VC1265 | hypothetical protein | 2.4 | 3.6E-05 | 1.4 | 9.6E-03 |
| VC1268 | conserved hypothetical protein | 2.4 | 1.0E-03 | 1.4 | 1.9E-01 |
| VC1313 | methyl-accepting chemotaxis protein | 2.6 | 9.0E-06 | 2.1 | 2.7E-06 |
| VC1318 | outer membrane protein OmpV | 7.9 | 1.8E-21 | 16.1 | 1.0E-74 |
| VC1329 | opacity protein-related protein | 13.6 | 2.5E-37 | 2.7 | 4.9E-06 |
| VC1330 | hypothetical protein | 4.3 | 1.9E-12 | 1.5 | 1.8E-01 |
| VC1344 | 4-hydroxyphenylpyruv ate dioxygenase | 3.9 | 2.5E-12 | 4.9 | 6.5E-26 |
| VC1345 | oxidoreductase, putative | 5.2 | 4.1E-21 | 5.9 | 1.3E-43 |
| VC1346 | conserved hypothetical protein | 4.9 | 2.5E-21 | 4.3 | 9.1E-24 |
| VC1347 | glutathione S-transferase, putative | 4.4 | 2.9E-13 | 5.3 | 3.7E-31 |
| VC1418 | hypothetical protein | 2.4 | 1.1E-05 | 2.5 | 1.3E-10 |
| VC1419 | hypothetical protein | 3.0 | 2.4E-07 | 2.1 | 2.2E-03 |
| VC1420 | hypothetical protein | 2.2 | 1.6E-03 | 1.7 | 8.4E-03 |
| VC1448 | RTX toxin transporter | 2.3 | 1.0E-03 | 1.4 | 1.2E-01 |
| VC1456 | cholera enterotoxin, B subunit | 6.2 | 1.1E-09 | 2.4 | 2.0E-02 |
| VC1457 | cholera enterotoxin, A subunit | 3.6 | 2.2E-07 | 1.2 | 6.0E-01 |
| VC1543 | hypothetical protein | 2.6 | 8.0E-08 | 1.6 | 1.5E-03 |
| VC1544 | tonB2 protein | 2.8 | 1.3E-06 | 1.6 | 6.0E-02 |

| | | | | | |
|---|---|---|---|---|---|
| VC1545 | TonB system transport protein ExbD2 | 2.8 | 1.1E-04 | -1.0 | 9.8E-01 |
| VC1546 | TonB system transport protein ExbB2 | 3.7 | 5.5E-09 | 1.8 | 4.3E-03 |
| VC1547 | biopolymer transport protein ExbB-relatedprotein | 3.6 | 2.1E-11 | 1.6 | 3.9E-03 |
| VC1548 | hypothetical protein | 3.6 | 4.3E-06 | 1.7 | 3.0E-02 |
| VC1570 | quinol oxidase, subunit II | 3.9 | 1.9E-07 | 1.7 | 2.5E-03 |
| VC1571 | quinol oxidase, subunit I | 3.3 | 8.0E-08 | 1.7 | 2.9E-04 |
| VC1572 | hypothetical protein | 2.7 | 1.3E-04 | 1.0 | 9.4E-01 |
| VC1573 | fumarate hydratase, class II | 4.4 | 2.7E-07 | 1.8 | 7.7E-03 |
| VC1577 | hypothetical protein | 5.0 | 5.9E-08 | 6.8 | 7.2E-26 |
| VC1578 | hypothetical protein | 5.8 | 4.7E-10 | 8.6 | 6.7E-19 |
| VC1579 | enterobactin synthetase component F-relatedprotein | 5.8 | 7.1E-15 | 8.7 | 3.5E-28 |
| VC1585 | catalase | 5.7 | 2.5E-10 | 4.3 | 4.9E-19 |
| VC1633 | hypothetical protein | 2.6 | 2.1E-05 | 1.7 | 3.5E-03 |
| VC1634 | multidrug resistance protein | 2.2 | 2.4E-03 | 1.4 | 2.9E-03 |
| VC1644 | hypothetical protein | 2.3 | 2.4E-05 | 1.9 | 1.5E-04 |
| VC1688 | hypothetical protein | 9.3 | 7.6E-12 | 1.7 | 7.0E-02 |
| VC1776 | N-acetylneuraminate lyase, putative | 2.2 | 6.8E-04 | 1.8 | 1.4E-04 |
| VC1807 | pseudogene | 3.0 | 2.0E-03 | 1.1 | 9.2E-01 |
| VC1808 | hypothetical protein | 3.2 | 5.5E-12 | 1.4 | 1.2E-01 |
| VC1819 | aldehyde dehydrogenase | 5.7 | 7.2E-22 | 3.9 | 1.5E-24 |
| VC1888 | hemolysin-related protein | 4.1 | 3.5E-12 | 4.3 | 2.3E-17 |
| VC1928 | C4-dicarboxylate transport protein DctQ,putative | 2.2 | 1.1E-05 | 1.2 | 3.1E-01 |
| VC1945 | FAD monooxygenase, PheA/TfdB family | 2.6 | 2.2E-06 | -1.2 | 5.4E-01 |
| VC1947 | transcriptional regulator, LysR family | 3.2 | 4.8E-06 | -1.2 | 6.7E-01 |
| VC1948 | hypothetical protein | 52.7 | 1.1E-06 | 2.1 | 5.9E-01 |
| VC1949 | pvcA protein | 6.4 | 3.5E-13 | -1.1 | 9.0E-01 |
| VC1962 | lipoprotein | 2.1 | 3.7E-05 | 2.4 | 1.3E-08 |
| VC2004 | conserved hypothetical protein | 3.4 | 1.5E-10 | 1.4 | 2.7E-01 |
| VC2069 | flagellar biosynthetic protein FlhA | 2.2 | 7.7E-05 | -1.1 | 8.2E-01 |
| VC2209 | nonribosomal peptide synthetase VibF | 4.5 | 1.5E-13 | 1.5 | 9.8E-04 |
| VC2210 | vibriobactin utilization protein ViuB | 7.1 | 1.2E-28 | 3.1 | 1.2E-11 |
| VC2211 | ferric vibriobactin receptor | 5.5 | 4.5E-11 | 3.2 | 4.1E-16 |
| VC2212 | hypothetical protein | 3.1 | 5.8E-09 | 3.5 | 3.3E-18 |
| VC2385 | RNA-directed DNA polymerase | 2.4 | 8.1E-07 | 1.5 | 3.1E-03 |
| VC2386 | conserved hypothetical protein | 2.1 | 5.0E-04 | 1.2 | 4.8E-01 |
| VC2387 | conserved hypothetical protein | 3.0 | 8.7E-06 | 1.5 | 2.0E-02 |
| VC2416 | 2`,3`-cyclic-nucleot ide 2`-phosphodiesterase,putative | 2.4 | 1.2E-05 | 2.4 | 2.7E-08 |
| VC2539 | thiamin ABC transporter, periplasmicthiamin-binding protein | 2.7 | 2.8E-05 | 4.6 | 2.3E-38 |
| VC2667 | hypothetical protein | 6.7 | 2.8E-12 | 4.7 | 2.7E-24 |
| VC2691 | periplasmic protein cpxP, putative | 2.3 | 8.7E-04 | 3.5 | 1.3E-14 |
| VC2694 | superoxide dismutase, Mn | 3.5 | 8.2E-14 | 1.7 | 2.0E-04 |
| VC2704 | hypothetical protein | 2.1 | 3.2E-05 | 1.2 | 4.5E-01 |
| VC2705 | sodium/solute symporter, putative | 2.0 | 1.5E-04 | 1.6 | 8.8E-03 |
| VCA0063 | protease II | 2.8 | 2.9E-05 | 2.1 | 2.5E-05 |
| VCA0070 | phosphate ABC transporter, periplasmicphosphate-binding protein | 3.0 | 5.6E-07 | 2.6 | 3.3E-08 |
| VCA0071 | phosphate ABC transporter, permease protein | 2.8 | 4.1E-06 | 2.2 | 9.0E-04 |
| VCA0072 | phosphate ABC transporter, permease protein | 2.5 | 1.8E-03 | 1.6 | 6.0E-02 |
| VCA0073 | phosphate ABC transporter, ATP-binding protein | 2.7 | 8.7E-06 | 1.5 | 6.0E-02 |
| VCA0083 | multidrug resistance protein D | 4.8 | 1.4E-05 | 1.5 | 6.0E-02 |
| VCA0084 | soxR protein | 2.2 | 4.2E-03 | 1.5 | 2.0E-02 |
| VCA0094 | conserved hypothetical protein | 14.1 | 2.5E-23 | 2.0 | 6.0E-02 |
| VCA0095 | hypothetical protein | 5.8 | 5.5E-05 | 1.5 | 6.8E-01 |
| VCA0140 | spindolin-related protein | 4.7 | 1.1E-14 | 2.3 | 4.9E-08 |
| VCA0144 | immunogenic protein | 3.0 | 3.1E-03 | 3.2 | 5.8E-16 |
| VCA0146 | conserved hypothetical protein | 2.9 | 1.6E-03 | 4.7 | 4.6E-14 |
| VCA0147 | transcriptional regulator, putative | 3.6 | 1.3E-04 | 3.1 | 1.2E-06 |
| VCA0152 | conserved hypothetical protein | 2.0 | 3.5E-03 | 1.1 | 6.5E-01 |
| VCA0160 | tryptophan-specific transport protein | 5.2 | 7.8E-13 | 1.6 | 1.0E-02 |
| VCA0161 | tryptophanase | 6.0 | 5.3E-22 | 2.2 | 8.7E-07 |
| VCA0163 | conserved hypothetical protein | 2.7 | 5.5E-05 | 1.8 | 3.7E-03 |
| VCA0218 | thermolabile hemolysin | 3.4 | 1.2E-07 | 5.6 | 5.6E-36 |
| VCA0221 | lactonizing lipase | 5.7 | 3.0E-15 | 1.5 | 1.1E-01 |
| VCA0222 | lipase activator protein, putative | 3.4 | 6.9E-05 | 2.4 | 7.4E-04 |
| VCA0227 | iron(III) ABC transporter, periplasmiciron-compound-binding protein | 4.3 | 1.1E-13 | 2.7 | 8.8E-11 |
| VCA0228 | iron(III) ABC transporter, permease protein | 2.9 | 1.6E-05 | 1.5 | 9.0E-02 |
| VCA0229 | iron(III) ABC transporter, permease protein | 2.2 | 3.4E-03 | 1.2 | 5.3E-01 |
| VCA0230 | iron(III) ABC transporter, ATP-binding protein | 2.9 | 5.7E-06 | 2.1 | 1.6E-04 |
| VCA0231 | transcriptional regulator, AraC/XylS family | 7.9 | 9.6E-20 | 2.4 | 3.2E-06 |
| VCA0232 | enterobactin receptor, VctA | 6.9 | 2.9E-13 | 1.1 | 7.8E-01 |
| VCA0233 | hypothetical protein | 4.6 | 9.3E-08 | 1.8 | 5.0E-02 |
| VCA0271 | hypothetical protein | 2.2 | 2.0E-05 | 1.7 | 1.3E-03 |
| VCA0276 | Glycine cleavage system P protein GcvP, authentic frameshift | 2.8 | 5.9E-09 | 2.7 | 3.8E-12 |
| VCA0277 | glycine cleavage system H protein | 2.2 | 5.4E-05 | 2.2 | 2.8E-08 |
| VCA0278 | serine hydroxymethyltransferase | 2.4 | 2.8E-06 | 2.1 | 5.1E-07 |
| VCA0280 | glycine cleavage system protein GcvT; contains authentic frameshift | 2.1 | 8.5E-05 | 1.8 | 1.6E-04 |
| VCA0312 | hypothetical protein | 18.9 | 7.5E-20 | 20.0 | 2.0E-34 |
| VCA0454 | sulfate-binding protein | 2.2 | 6.2E-03 | 1.1 | 7.8E-01 |
| VCA0576 | heme transport protein HutA | 6.6 | 2.9E-14 | 9.7 | 3.5E-28 |
| VCA0682 | transcriptional regulator UhpA | 2.4 | 2.5E-04 | -1.1 | 6.4E-01 |
| VCA0683 | sensor protein UhpB | 2.1 | 3.6E-03 | -1.2 | 2.8E-01 |
| VCA0827 | pterin-4-alpha-carbi nolamine dehydratase | 2.5 | 3.7E-06 | 4.4 | 3.5E-28 |
| VCA0828 | phenylalanine-4-hydr oxylase | 2.5 | 7.1E-05 | 2.8 | 1.0E-09 |
| VCA0849 | hypothetical protein | 2.3 | 4.6E-04 | 3.1 | 3.3E-12 |

| | | | | | |
|---|---|---|---|---|---|
| VCA0864 | methyl-accepting chemotaxis protein | 3.0 | 8.1E-06 | 4.3 | 3.5E-21 |
| VCA0865 | hemagglutinin/protea se | 2.4 | 7.4E-03 | -1.6 | 7.0E-02 |
| VCA0866 | hypothetical protein | 2.4 | 5.8E-03 | -1.3 | 4.9E-01 |
| VCA0874 | hypothetical protein | 2.3 | 3.1E-03 | 4.6 | 3.6E-11 |
| VCA0885 | threonine 3-dehydrogenase | 3.0 | 2.1E-11 | 1.8 | 1.1E-04 |
| VCA0886 | 2-amino-3-ketobutyra te coenzyme A ligase | 3.1 | 2.1E-08 | 2.4 | 9.8E-13 |
| VCA0907 | conserved hypothetical protein | 5.9 | 1.8E-21 | 3.1 | 1.3E-14 |
| VCA0908 | conserved hypothetical protein | 7.5 | 2.6E-28 | 3.4 | 1.3E-15 |
| VCA0909 | oxygen-independent coproporphyrinogen IIIoxidase, putative | 9.8 | 1.3E-15 | 2.6 | 2.0E-04 |
| VCA0910 | tonB1 protein | 9.8 | 7.3E-20 | 3.9 | 2.2E-09 |
| VCA0911 | TonB system transport protein ExbB1 | 10.3 | 1.5E-17 | 3.3 | 2.4E-09 |
| VCA0912 | TonB system transport protein ExbD1 | 7.3 | 2.2E-11 | 3.0 | 6.0E-09 |
| VCA0913 | hemin ABC transporter, periplasmic hemin-bindingprotein HutB | 6.6 | 8.0E-10 | 2.5 | 2.9E-08 |
| VCA0914 | hemin ABC transporter, permease protein,putative | 5.6 | 6.5E-11 | 2.6 | 1.1E-07 |
| VCA0928 | hypothetical protein | 2.4 | 1.2E-03 | -1.6 | 1.2E-01 |
| VCA0946 | maltose/maltodextrin ABC transporter,ATP-binding protein | 2.3 | 3.5E-05 | -1.3 | 2.4E-01 |
| VCA0952 | transcriptional regulator, LuxR family | 3.5 | 6.7E-04 | 3.6 | 3.8E-05 |
| VCA0962 | hypothetical protein | 2.3 | 1.1E-05 | 1.7 | 1.7E-03 |
| VCA0976 | hypothetical protein | 7.5 | 2.1E-13 | 2.2 | 2.9E-04 |
| VCA0977 | ABC transporter, ATP-binding protein | 5.4 | 1.4E-21 | 2.8 | 4.0E-22 |
| VCA1031 | putative methyl accepting chemotaxis protein; authentic frameshift mutation | 2.3 | 1.7E-04 | 1.5 | 4.0E-02 |
| VCA1041 | phosphomannomutase, putative | 2.2 | 2.2E-04 | 1.1 | 7.2E-01 |
| 5Sb | 5S ribosomal RNA | 4.9 | 1.4E-05 | 1.7 | 3.2E-01 |
| 5Sc | 5S ribosomal RNA | 16.0 | 1.5E-03 | -359.6 | 3.0E-02 |
| 5Sd | 5S ribosomal RNA | 9.0 | 4.7E-03 | -61.6 | 2.5E-03 |
| 5Sf | 5S ribosomal RNA | 7.1 | 1.1E-03 | 3.6 | 1.2E-07 |
| 5Sh | 5S ribosomal RNA | 3.1 | 9.0E-03 | 1.7 | 3.6E-03 |
| VCr025 | 5S ribosomal RNA | 4.7 | 3.0E-04 | 1.5 | 1.3E-01 |
