## Supplementary material for "Modular small RNA drives pathogen emergence": Table S2

**Table S2.** Differentially expressed genes in biofilms in *ompU* and *rpoE* mutants compared to WT C6706.

| 144 differentially-expressed genes exclusively in Δ*ompU* and not Δ*rpoE*, compared to the WT | | | |
|---|---|---|---|
| **Name** | **Function** | **Fold change** | **FDR *p*-value** |
| VC0176 | transcriptional regulator, putative | -2.75 | 6.55E-04 |
| VC0177 | hypothetical protein | -2.63 | 3.84E-06 |
| VC0178 | patatin-related protein | -2.22 | 8.01E-06 |
| VC0179 | hypothetical protein | -2.45 | 2.59E-07 |
| VC0180 | conserved hypothetical protein | -2.30 | 2.75E-08 |
| VC0495 | conserved hypothetical protein | -2.06 | 9.36E-04 |
| VC0502 | type IV pilin, putative | -2.69 | 2.63E-08 |
| VC0504 | hypothetical protein | -2.46 | 3.14E-06 |
| VC0505 | hypothetical protein | -2.86 | 7.32E-06 |
| VC0506 | hypothetical protein | -2.63 | 2.69E-05 |
| VC0507 | hypothetical protein | -4.79 | 1.07E-05 |
| VC0633 | outer membrane protein OmpU | -15766.55 | 0.00E+00 |
| VC0711 | clpB protein | -2.00 | 2.84E-05 |
| VC0817 | transposase, putative | -2.08 | 1.67E-04 |
| VC0856 | dnaJ protein | -2.12 | 8.37E-07 |
| VC0886 | hypothetical protein | -2.27 | 9.61E-07 |
| VC0985 | heat shock protein HtpG | -2.09 | 2.60E-06 |
| VC1217 | conserved hypothetical protein | -2.15 | 2.56E-04 |
| VC1316 | chemotaxis protein CheY, putative | -2.28 | 4.53E-06 |
| VC1562 | beta-lactamase-relat ed protein | -2.19 | 3.31E-04 |
| VC1676 | phage shock protein C | -2.33 | 1.27E-06 |
| VC1677 | phage shock protein B, pspB | -2.15 | 3.76E-05 |
| VC1762 | hypothetical protein | -2.09 | 6.59E-05 |
| VC1763 | chemotaxis protein MotB-related protein | -2.84 | 3.22E-07 |
| VC1774 | conserved hypothetical protein | -2.01 | 2.78E-04 |
| VC1793 | hypothetical protein | -2.92 | 4.44E-04 |
| VC2149 | hypothetical protein | -2.03 | 4.67E-05 |
| VC2663 | molecular chaperone groEL_1 | -2.12 | 4.68E-05 |
| VC2664 | chaperonin, 60 Kd subunit, groES_1 | -2.00 | 6.87E-05 |
| VCA0275 | IS1004 transposase | -5.01 | 8.16E-04 |
| VCA0298 | hypothetical protein | -2.22 | 8.51E-04 |
| VCA0336 | hypothetical protein | -3.17 | 2.75E-07 |
| VCA0344 | hypothetical protein | -2.06 | 7.21E-06 |
| VCA0354 | hypothetical protein | -2.02 | 1.45E-04 |
| VCA0357 | hypothetical protein | -2.97 | 2.16E-04 |
| VCA0358 | hypothetical protein | -2.04 | 3.62E-04 |
| VCA0367 | hypothetical protein | -2.05 | 1.39E-05 |
| VCA0382a | membrane protein | -2.00 | 9.91E-04 |
| VCA0397 | hypothetical protein | -2.43 | 7.81E-06 |
| VCA0398 | hypothetical protein | -2.74 | 5.30E-06 |
| VCA0409 | hypothetical protein | -2.16 | 3.44E-05 |
| VCA0410 | hypothetical protein | -2.03 | 4.45E-04 |
| VCA0415 | conserved hypothetical protein | -3.06 | 9.79E-05 |
| VCA0420 | hypothetical protein | -2.45 | 3.34E-05 |
| VCA0428 | hypothetical protein | -2.17 | 5.40E-07 |
| VCA0431 | hypothetical protein | -2.95 | 8.75E-10 |
| VCA0443 | lipoprotein Blc | -2.42 | 4.27E-06 |
| VCA0446 | haemagglutinin | -3.57 | 9.29E-13 |
| VCA0447 | haemagglutinin associated protein | -2.40 | 1.17E-06 |
| VCA0448 | hypothetical protein | -3.19 | 6.07E-09 |
| VCA0449 | hypothetical protein | -4.72 | 2.87E-07 |
| VCA0450 | hypothetical protein | -2.69 | 7.69E-04 |
| VCA0451 | hypothetical protein | -2.65 | 8.98E-04 |
| VCA0455 | conserved hypothetical protein | -2.12 | 8.65E-06 |
| VCA0508 | transposase OrfAB, subunit B | -2.81 | 1.01E-08 |
| VCA0697 | sensory box/GGDEF family protein | -2.01 | 2.57E-05 |
| VC0065 | thiG protein | 3.25 | 2.71E-08 |
| VC0066 | thiH protein | 2.48 | 2.44E-05 |
| VC0199 | hemolysin secretion ATP-binding protein,putative | 3.43 | 1.12E-06 |
| VC0200 | iron(III) compound receptor | 6.52 | 9.92E-13 |
| VC0202 | iron(III) ABC transporter, periplasmiciron-compound-binding protein | 2.47 | 3.19E-04 |
| VC0284 | putative outer membrane receptor | 3.05 | 2.37E-10 |
| VC0474 | iron-regulated virulence regulatory proteinIrgB | 3.29 | 1.74E-06 |

| | | | |
|---|---|---|---|
| VC0719 | DNA-binding response regulator PhoB | 2.39 | 1.09E-06 |
| VC0773 | vibriobactin-specifi c isochorismate synthase | 4.96 | 4.85E-07 |
| VC0775 | vibriobactin synthesis protein, putative | 2.70 | 2.65E-06 |
| VC0777 | ferric vibriobactin ABC transporter, permeaseprotein | 3.51 | 3.53E-07 |
| VC0778 | ferric vibriobactin ABC transporter, permeaseprotein | 3.12 | 3.68E-07 |
| VC0820 | ToxR-activated gene A protein | 4.04 | 1.01E-10 |
| VC0821 | hypothetical protein | 3.56 | 3.27E-15 |
| VC0823 | hypothetical protein | 2.76 | 1.59E-10 |
| VC0929 | hypothetical protein | 5.80 | 9.83E-30 |
| VC0931 | conserved hypothetical protein | 3.07 | 1.60E-05 |
| VC0932 | hypothetical protein | 4.84 | 9.46E-09 |
| VC0935 | hypothetical protein | 3.19 | 7.66E-05 |
| VC0937 | exopolysaccharide biosynthesis protein,putative | 2.29 | 3.01E-04 |
| VC0940 | conserved hypothetical protein | 2.08 | 2.76E-04 |
| VC1154 | hypothetical protein | 2.04 | 1.37E-05 |
| VC1264 | iron-regulated protein A, putative | 2.22 | 7.50E-06 |
| VC1265 | hypothetical protein | 2.35 | 1.36E-06 |
| VC1268 | conserved hypothetical protein | 2.35 | 6.02E-05 |
| VC1313 | methyl-accepting chemotaxis protein | 2.55 | 2.57E-07 |
| VC1330 | hypothetical protein | 4.29 | 1.94E-14 |
| VC1347 | glutathione S-transferase, putative | 4.40 | 2.47E-15 |
| VC1418 | hypothetical protein | 2.44 | 3.36E-07 |
| VC1419 | hypothetical protein | 2.97 | 5.04E-09 |
| VC1420 | hypothetical protein | 2.18 | 1.03E-04 |
| VC1448 | RTX toxin transporter | 2.34 | 6.17E-05 |
| VC1543 | hypothetical protein | 2.61 | 1.53E-09 |
| VC1544 | tonB2 protein | 2.82 | 3.21E-08 |
| VC1545 | TonB system transport protein ExbD2 | 2.84 | 4.65E-06 |
| VC1546 | TonB system transport protein ExbB2 | 3.67 | 8.91E-11 |
| VC1547 | biopolymer transport protein ExbB-relatedprotein | 3.61 | 2.63E-13 |
| VC1573 | fumarate hydratase, class II, fumC | 4.43 | 5.59E-09 |
| VC1585 | catalase | 5.65 | 3.81E-12 |
| VC1634 | multidrug resistance protein | 2.18 | 1.66E-04 |
| VC1688 | hypothetical protein | 9.33 | 9.07E-14 |
| VC1776 | N-acetylneuraminate lyase, putative | 2.18 | 3.80E-05 |
| VC1808 | hypothetical protein | 3.19 | 6.45E-14 |
| VC1945 | FAD monooxygenase, PheA/TfdB family | 2.58 | 5.40E-08 |
| VC1947 | transcriptional regulator, LysR family | 3.24 | 1.25E-07 |
| VC1948 | hypothetical protein | 52.72 | 2.54E-08 |
| VC1949 | pvcA protein | 6.42 | 3.26E-15 |
| VC2004 | conserved hypothetical protein | 3.36 | 2.22E-12 |
| VC2069 | flagellar biosynthetic protein FlhA | 2.19 | 3.16E-06 |
| VC2209 | nonribosomal peptide synthetase VibF | 4.50 | 1.20E-15 |
| VC2210 | vibriobactin utilization protein ViuB | 7.05 | 1.28E-31 |
| VC2211 | ferric vibriobactin receptor | 5.50 | 6.13E-13 |
| VC2212 | hypothetical protein | 3.05 | 9.59E-11 |
| VC2385 | RNA-directed DNA polymerase | 2.44 | 1.82E-08 |
| VC2386 | conserved hypothetical protein | 2.08 | 2.69E-05 |
| VC2387 | conserved hypothetical protein | 2.96 | 2.44E-07 |
| VC2416 | 2`,3`-cyclic-nucleot ide 2`-phosphodiesterase,putative | 2.35 | 3.80E-07 |
| VC2667 | hypothetical protein | 6.70 | 3.09E-14 |
| VC2694 | superoxide dismutase, Mn | 3.54 | 6.01E-16 |
| VC2704 | hypothetical protein | 2.10 | 1.18E-06 |
| VC2705 | sodium/solute symporter, putative | 2.02 | 6.89E-06 |
| VCA0073 | phosphate ABC transporter, ATP-binding protein | 2.71 | 2.44E-07 |
| VCA0084 | soxR protein | 2.17 | 3.34E-04 |
| VCA0095 | hypothetical protein | 5.84 | 2.18E-06 |
| VCA0140 | spindolin-related protein | 4.73 | 7.10E-17 |
| VCA0144 | immunogenic protein | 2.98 | 2.24E-04 |
| VCA0147 | transcriptional regulator, putative | 3.56 | 6.02E-06 |
| VCA0160 | tryptophan-specific transport protein | 5.17 | 7.59E-15 |
| VCA0218 | thermolabile hemolysin | 3.35 | 2.36E-09 |
| VCA0222 | lipase activator protein, putative | 3.41 | 2.80E-06 |
| VCA0228 | iron(III) ABC transporter, permease protein | 2.85 | 5.14E-07 |
| VCA0229 | iron(III) ABC transporter, permease protein | 2.24 | 2.48E-04 |
| VCA0230 | iron(III) ABC transporter, ATP-binding protein | 2.91 | 1.52E-07 |
| VCA0232 | enterobactin receptor, VctA | 6.91 | 2.52E-15 |
| VCA0233 | hypothetical protein | 4.59 | 1.82E-09 |
| VCA0277 | glycine cleavage system H protein | 2.15 | 2.12E-06 |
| VCA0312 | hypothetical protein | 18.88 | 3.27E-22 |

| | | | |
|---|---|---|---|
| VCA0454 | sulfate-binding protein | 2.20 | 5.46E-04 |
| VCA0682 | transcriptional regulator UhpA | 2.37 | 1.21E-05 |
| VCA0865 | hemagglutinin/protea se | 2.37 | 6.83E-04 |
| VCA0866 | hypothetical protein | 2.37 | 5.08E-04 |
| VCA0908 | conserved hypothetical protein | 7.45 | 3.55E-31 |
| VCA0912 | TonB system transport protein ExbD1 | 7.31 | 2.80E-13 |
| VCA0913 | hemin ABC transporter, periplasmic hemin-bindingprotein HutB | 6.55 | 1.27E-11 |
| VCA0914 | hemin ABC transporter, permease protein,putative | 5.58 | 8.99E-13 |
| VCA0928 | hypothetical protein | 2.37 | 7.18E-05 |
| VCA0946 | maltose/maltodextrin ABC transporter,ATP-binding protein | 2.32 | 1.33E-06 |
| VCA0962 | hypothetical protein | 2.29 | 3.20E-07 |

| 1288 differentially-expressed genes exclusively in ∆*rpoE* and not ∆*ompU* , compared to the WT | | | |
|---|---|---|---|
| **Locus Tag** | **Function** | **Fold change** | **FDR *p* -value** |
| 16Sa | 16S ribosomal RNA | -36.78 | 1.29531E-47 |
| 16Sb | 16S ribosomal RNA | -28.41 | 1.16083E-37 |
| 16Sc | 16S ribosomal RNA | -26.06 | 7.26317E-41 |
| 16Sd | 16S ribosomal RNA | -4.18 | 7.97962E-10 |
| 16Se | 16S ribosomal RNA | -17.77 | 3.85367E-32 |
| 16Sg | 16S ribosomal RNA | -37.00 | 8.65305E-43 |
| 23Sa | 23S ribosomal RNA | -10.75 | 1.09283E-22 |
| 23Sc | 23S ribosomal RNA | -9.75 | 1.49741E-08 |
| 23Sd | 23S ribosomal RNA | -9.86 | 6.53025E-24 |
| 23Se | 23S ribosomal RNA | -166.20 | 1.0355E-104 |
| 23Sf | 23S ribosomal RNA | -5.76 | 9.09732E-15 |
| 23Sg | 23S ribosomal RNA | -28.39 | 1.48846E-41 |
| 23Sh | 23S ribosomal RNA | -2.74 | 1.45461E-05 |
| 5Sa | 5S ribosomal RNA | -87.55 | 1.01597E-32 |
| 5Se | 5S ribosomal RNA | -48.28 | 3.09435E-38 |
| 5Sg | 5S ribosomal RNA | -19110.69 | 5.183E-06 |
| VC0008 | amino acid ABC transporter, ATP-binding protein | -2.31 | 6.66E-09 |
| VC0026 | zinc-binding alcohol dehydrogenase | -2.22 | 1.01E-12 |
| VC0027 | threonine dehydratase | -5.38 | 6.61E-78 |
| VC0028 | dihydroxy-acid dehydratase | -3.87 | 6.59E-35 |
| VC0030 | acetolactate synthase II, small subunit | -2.67 | 4.91E-07 |
| VC0032 | ComM-related protein | -2.05 | 3.72E-05 |
| VC0036 | FixG-related protein | -2.82 | 4.20E-24 |
| VC0048 | smf protein | -2.43 | 6.46E-11 |
| VC0049 | smg protein | -3.10 | 3.89E-29 |
| VC0050 | DNA topoisomerase I-related protein | -3.73 | 1.53E-22 |
| VC0075 | MadN protein | -5.69 | 6.52E-47 |
| VC0076 | universal stress protein A | -7.29 | 5.00E-20 |
| VC0079 | conserved hypothetical protein | -3.77 | 8.23E-31 |
| VC0089 | cytochrome c551 peroxidase | -11.01 | 2.78E-62 |
| VC0098 | methyl-accepting chemotaxis protein | -3.64 | 1.68E-25 |
| VC0131 | conserved hypothetical protein | -2.59 | 9.19E-12 |
| VC0139 | DPS family protein | -2.12 | 2.39E-09 |
| VC0159 | RNA-binding protein | -2.02 | 1.19E-09 |
| VC0164 | multidrug resistance protein, putative | -2.57 | 6.38E-13 |
| VC0174 | hypothetical protein | -4.59 | 8.79E-41 |
| VC0185 | transposase, putative | -2.32 | 2.50E-10 |
| VC0204 | conserved hypothetical protein | -5.16 | 4.30E-22 |
| VC0216 | methyl-accepting chemotaxis protein | -3.90 | 2.32E-32 |
| VC0217 | DNA repair protein RadC | -4.77 | 1.00E-19 |
| VC0229 | hypothetical protein | -2.53 | 9.68E-22 |
| VC0235 | lipopolysaccharide biosynthesis protein,putative | -2.39 | 7.92E-18 |
| VC0245 | rfbG protein | -2.27 | 1.18E-07 |
| VC0246 | lipopolysaccharide/O -antigen transport protein | -2.16 | 7.35E-13 |
| VC0247 | lipopolysaccharide/O -antigen transport protein | -2.71 | 3.25E-26 |
| VC0248 | acyl carrier protein, putative | -2.57 | 1.79E-26 |
| VC0249 | rfbL protein | -2.92 | 9.84E-14 |
| VC0250 | iron-containing alcohol dehydrogenase familyprotein RfbM | -3.35 | 2.61E-20 |
| VC0251 | acyl protein synthase/acyl-CoA reductase RfbN | -2.96 | 2.28E-08 |
| VC0252 | acetyltransferase RfbO, CysE/LacA/LpxA/NodLfamily | -2.11 | 3.22E-10 |
| VC0254 | conserved hypothetical protein | -2.50 | 2.00E-12 |
| VC0266 | conserved hypothetical protein | -2.05 | 5.44E-05 |
| VC0274 | hypothetical protein | -2.03 | 1.42E-11 |
| VC0280 | cadaverine/lysine antiporter CadB, putative | -2.88 | 1.05E-06 |
| VC0281 | lysine decarboxylase, inducible | -2.02 | 2.89E-04 |

| | | | |
|---|---|---|---|
| VC0282 | methyl-accepting chemotaxis protein | -3.39 | 6.66E-28 |
| VC0285 | 4-hydroxy-2-oxogluta ratealdolase/2-deydro-3-deoxyp hosphogluco | -2.25 | 1.92E-09 |
| VC0286 | gluconate permease, putative | -3.94 | 2.35E-22 |
| VC0287 | thermoresistant gluconokinase | -2.76 | 7.42E-10 |
| VC0288 | phosphogluconate dehydratase | -3.41 | 6.32E-11 |
| VC0295 | acetyl-CoA carboxylase, biotin carboxylase | -2.91 | 3.64E-21 |
| VC0296 | acetyl-CoA carboxylase, biotin carboxyl carrierprotein | -2.10 | 1.13E-11 |
| VC0298 | acetyl-CoA synthase | -2.59 | 8.76E-12 |
| VC0329 | DNA-directed RNA polymerase, beta | -2.29 | 3.47E-05 |
| VC0330 | regulator of sigma D | -2.91 | 1.58E-16 |
| VC0338 | transporter, putative | -4.26 | 3.61E-33 |
| VC0353 | conserved hypothetical protein | -2.73 | 6.55E-18 |
| VC0357 | conserved hypothetical protein | -2.56 | 1.03E-12 |
| VC0377 | conserved hypothetical protein | -2.36 | 9.24E-13 |
| VC0389 | Na+/H+ antiporter, putative | -2.81 | 5.15E-19 |
| VC0400 | MSHA biogenesis protein MshJ | -2.43 | 2.46E-12 |
| VC0401 | MSHA biogenesis protein MshK | -2.97 | 2.02E-12 |
| VC0402 | MSHA biogenesis protein MshL | -4.06 | 1.81E-25 |
| VC0403 | MSHA biogenesis protein MshM | -5.95 | 7.61E-51 |
| VC0404 | MSHA biogenesis protein MshN | -6.21 | 9.01E-61 |
| VC0405 | MSHA biogenesis protein MshE | -5.30 | 5.04E-55 |
| VC0406 | MSHA biogenesis protein MshG | -4.31 | 1.89E-53 |
| VC0410 | MSHA pilin protein MshC | -2.40 | 5.41E-23 |
| VC0411 | MSHA pilin protein MshD | -2.65 | 4.40E-16 |
| VC0412 | hypothetical protein | -3.50 | 3.33E-40 |
| VC0413 | hypothetical protein | -3.32 | 1.19E-24 |
| VC0414 | hypothetical protein | -4.50 | 1.51E-49 |
| VC0419 | cytoplasmic axial filament protein | -2.06 | 2.55E-11 |
| VC0420 | conserved hypothetical protein | -2.25 | 2.54E-16 |
| VC0423 | arginine deiminase | -4.29 | 1.69E-34 |
| VC0441 | bis(5`-nucleosyl)-te traphosphatase | -2.81 | 3.17E-28 |
| VC0442 | ApaG protein | -2.38 | 1.41E-19 |
| VC0443 | 16S rRNA (adenine(1518)-N(6)/adenine(1519)-N(6))- dimethyltrans | -2.63 | 8.59E-27 |
| VC0445 | survival protein SurA | -2.04 | 1.76E-14 |
| VC0462 | twitching motility protein PilT | -3.94 | 1.72E-27 |
| VC0463 | twitching motility protein PilT | -5.05 | 5.50E-36 |
| VC0488 | extracellular solute-binding protein, putative | -2.58 | 3.01E-22 |
| VC0490 | conserved hypothetical protein | -2.72 | 6.90E-13 |
| VC0491 | hypothetical protein | -3.58 | 5.04E-46 |
| VC0492 | hypothetical protein | -2.67 | 1.37E-11 |
| VC0534 | RNA polymerase sigma-38 factor | -3.13 | 7.30E-15 |
| VC0550 | oxaloacetate decarboxylase, alpha subunit | -3.11 | 2.68E-22 |
| VC0551 | oxaloacetate decarboxylase, beta subunit | -2.86 | 3.11E-19 |
| VC0552 | quinone oxidoreductase | -4.52 | 2.50E-36 |
| VC0553 | conserved hypothetical protein | -4.48 | 2.79E-24 |
| VC0554 | protease, insulinase family/protease, insulinasefamily | -4.19 | 1.75E-38 |
| VC0653 | c-di-GMP phosphodiesterase A-related protein | -2.27 | 1.39E-10 |
| VC0658 | c-di-GMP phosphodiesterase A-related protein | -3.92 | 4.46E-21 |
| VC0661 | conserved hypothetical protein | -3.65 | 4.98E-17 |
| VC0665 | sigma-54 dependent transcriptional regulator | -3.19 | 5.27E-22 |
| VC0691 | anhydro-N-acetylmuramic acid kinase | -2.14 | 4.55E-16 |
| VC0708 | conserved hypothetical protein | -2.20 | 1.15E-12 |
| VC0730 | copper homeostasis protein | -3.78 | 2.09E-16 |
| VC0737 | acetoin utilization protein AcuB, putative | -2.70 | 7.81E-24 |
| VC0762 | conserved hypothetical protein | -2.83 | 3.60E-20 |
| VC0770 | conserved hypothetical protein | -2.01 | 5.69E-04 |
| VC0788 | DOPA-dioxygenase-rel ated protein | -2.07 | 7.53E-07 |
| VC0802 | hypothetical protein | -3.60 | 2.79E-14 |
| VC0824 | 2-Cys peroxiredoxin | -3.34 | 1.82E-12 |
| VC0828 | toxin co-regulated pilin | -41.51 | 1.65E-47 |
| VC0829 | toxin co-regulated pilus biosynthesis protein B | -2.92 | 1.21E-07 |
| VC0830 | toxin co-regulated pilus biosynthesis protein Q | -2.55 | 2.00E-04 |
| VC0831 | toxin co-regulated pilus biosynthesis outermembrane protein C | -4.38 | 1.60E-17 |
| VC0832 | toxin co-regulated pilus biosynthesis protein R | -7.24 | 8.28E-12 |
| VC0833 | toxin co-regulated pilus biosynthesis protein D | -7.42 | 1.40E-11 |
| VC0836 | toxin co-regulated pilus biosynthesis protein E | -4.02 | 1.38E-08 |
| VC0837 | toxin co-regulated pilus biosynthesis protein F | -5.34 | 4.26E-18 |
| VC0839 | leader peptidase TcpJ | -5.10 | 1.37E-06 |
| VC0841 | accessory colonization factor AcfC | -2.12 | 1.25E-04 |
| VC0844 | accessory colonization factor AcfA | -5.63 | 1.56E-16 |

| | | | |
|---|---|---|---|
| VC0845 | conserved hypothetical protein | -2.42 | 3.22E-08 |
| VC0863 | conserved hypothetical protein | -2.18 | 1.15E-18 |
| VC0893 | chemotaxis protein PomB | -2.25 | 4.51E-20 |
| VC0896 | transcriptional regulator, LysR family | -2.17 | 6.01E-07 |
| VC0899 | conserved hypothetical protein | -8.39 | 6.82E-56 |
| VC0910 | PTS system, trehalose-specific IIBC component | -3.47 | 5.66E-19 |
| VC0911 | trehalose-6-phosphat e hydrolase | -2.95 | 2.52E-11 |
| VC0925 | polysaccharide biosynthesis protein, putative | -2.06 | 7.89E-06 |
| VC0926 | hypothetical protein | -2.09 | 2.08E-08 |
| VC0927 | UDP-N-acetyl-D-manno samine transferase | -2.46 | 3.60E-10 |
| VC0957 | conserved hypothetical protein | -2.13 | 1.19E-08 |
| VC0958 | apolipoprotein N-acyltransferase | -2.13 | 5.61E-16 |
| VC0975 | conserved hypothetical protein | -2.00 | 3.89E-13 |
| VC0976 | conserved hypothetical protein | -2.39 | 2.10E-13 |
| VC0979 | oxidoreductase, short-chaindehydrogenase/reduc tase family | -2.95 | 7.86E-09 |
| VC0980 | epimerase | -3.22 | 5.97E-17 |
| VC0995 | PTS system, N-acetylglucosamine-specific IIABCcomponent | -2.73 | 1.51E-14 |
| VC0998 | hypothetical protein | -2.58 | 4.78E-13 |
| VC1008 | sodium-type flagellar protein MotY | -2.06 | 3.75E-14 |
| VC1022 | phosphorelay protein | -2.22 | 7.16E-09 |
| VC1026 | molybdenum cofactor biosynthesis protein MoaC | -2.48 | 3.07E-05 |
| VC1028 | molybdenum cofactor biosynthesis protein E | -2.40 | 2.22E-06 |
| VC1031 | inosine monophosphate dehydrogenase-relatedprotein | -6.90 | 1.52E-94 |
| VC1032 | zinc/cadmium/mercury/lead-transporting ATPase | -2.21 | 5.37E-03 |
| VC1049 | transcriptional regulator, LysR family | -3.09 | 1.51E-20 |
| VC1050 | response regulator | -4.16 | 6.22E-25 |
| VC1062 | transcriptional regulator, AsnC family | -2.29 | 1.20E-05 |
| VC1063 | acyl-CoA thioesterase II | -2.20 | 2.98E-16 |
| VC1074 | hypothetical protein | -3.29 | 5.09E-14 |
| VC1080 | hypothetical protein | -4.52 | 6.31E-40 |
| VC1081 | response regulator | -7.60 | 1.41E-53 |
| VC1082 | response regulator | -15.06 | 1.19E-78 |
| VC1083 | hypothetical protein | -12.71 | 5.94E-81 |
| VC1084 | sensory box sensor histidine kinase | -14.82 | 3.96E-103 |
| VC1085 | sensor histidine kinase | -10.04 | 3.20E-78 |
| VC1086 | response regulator | -8.65 | 1.41E-68 |
| VC1087 | response regulator | -6.46 | 3.88E-43 |
| VC1088 | sensor histidine kinase | -5.72 | 4.06E-37 |
| VC1089 | periplasmic binding protein-related protein | -10.26 | 2.76E-57 |
| VC1103 | ABC transporter, ATP-binding protein | -2.01 | 1.08E-07 |
| VC1130 | DNA-binding protein H-NS | -2.61 | 2.76E-19 |
| VC1137 | phosphoribosylformim ino-5-aminoimidazolecarboxamid e ribotide is | -2.07 | 7.55E-07 |
| VC1138 | imidazoleglycerol-ph osphate synthase, cyclasesubunit | -2.84 | 3.23E-13 |
| VC1143 | conserved hypothetical protein | -2.06 | 2.63E-07 |
| VC1144 | ATP-dependent Clp protease, ATP-binding subunitClpA | -3.07 | 1.31E-14 |
| VC1147 | iron-containing alcohol dehydrogenase | -6.99 | 6.32E-71 |
| VC1150 | hypothetical protein | -2.02 | 2.00E-07 |
| VC1153 | conserved hypothetical protein | -2.18 | 2.42E-11 |
| VC1155 | response regulator | -4.40 | 1.54E-32 |
| VC1156 | sensor histidine kinase | -4.38 | 2.11E-29 |
| VC1158 | hypothetical protein | -2.41 | 1.23E-11 |
| VC1187 | hypothetical protein | -5.18 | 9.45E-17 |
| VC1188 | malate oxidoreductase | -11.77 | 2.17E-72 |
| VC1201 | hypothetical protein | -4.17 | 5.76E-33 |
| VC1202 | histidine ammonia-lyase | -5.35 | 1.80E-46 |
| VC1203 | urocanate hydratase | -6.96 | 6.50E-47 |
| VC1205 | imidazolonepropionas e | -4.04 | 1.00E-18 |
| VC1206 | histidine utilization repressor | -4.14 | 2.00E-20 |
| VC1207 | hypothetical protein | -2.59 | 1.83E-11 |
| VC1222 | integration host factor, alpha subunit | -3.47 | 2.08E-21 |
| VC1223 | hypothetical protein | -2.40 | 2.08E-09 |
| VC1235 | sodium/dicarboxylate symporter | -2.43 | 4.62E-08 |
| VC1236 | PilB-related protein | -2.28 | 1.16E-11 |
| VC1248 | methyl-accepting chemotaxis protein | -10.14 | 1.43E-50 |
| VC1267 | hypothetical protein | -2.41 | 1.56E-08 |
| VC1283 | PTS system, cellobiose-specific IIA component | -2.01 | 6.77E-04 |
| VC1284 | 6-phospho-beta-gluco sidase | -2.33 | 3.41E-11 |
| VC1287 | periplasmic glucans biosynthesis protein MdoH | -2.18 | 2.97E-12 |
| VC1295 | conserved hypothetical protein | -2.74 | 1.41E-09 |
| VC1298 | methyl-accepting chemotaxis protein | -2.04 | 3.01E-10 |

| | | | |
|---|---|---|---|
| VC1312 | alanine racemase, putative | -3.17 | 9.70E-14 |
| VC1317 | conserved hypothetical protein | -3.47 | 3.15E-26 |
| VC1319 | sensor histidine kinase | -4.70 | 1.81E-25 |
| VC1320 | DNA-binding response regulator | -5.04 | 1.55E-33 |
| VC1322 | conserved hypothetical protein | -6.92 | 2.62E-42 |
| VC1323 | hypothetical protein | -5.95 | 4.85E-40 |
| VC1325 | galactoside ABC transporter, periplasmicD-galactose/D-gluco se-bi | -89.16 | 9.08E-109 |
| VC1327 | galactoside ABC transporter, ATP-bindingprotein | -67.63 | 1.89E-182 |
| VC1328 | galactoside ABC transporter, permease protein | -46.24 | 1.92E-169 |
| VC1332 | conserved hypothetical protein | -2.06 | 1.69E-09 |
| VC1339 | conserved hypothetical protein | -2.09 | 8.47E-06 |
| VC1340 | prpE protein | -2.35 | 1.36E-11 |
| VC1348 | response regulator | -3.37 | 1.99E-26 |
| VC1349 | sensory box sensor histidine kinase/responseregulator | -2.63 | 7.80E-12 |
| VC1353 | GGDEF family protein | -2.41 | 9.11E-10 |
| VC1359 | amino acid ABC transporter, ATP-binding protein | -4.62 | 3.86E-45 |
| VC1360 | amino acid ABC transporter, permease protein | -5.94 | 7.82E-36 |
| VC1361 | amino acid ABC transporter, permease protein | -6.67 | 2.32E-35 |
| VC1362 | amino acid ABC transporter, periplasmic aminoacid-binding protein | -8.05 | 5.24E-40 |
| VC1367 | GGDEF family protein | -3.74 | 7.48E-32 |
| VC1376 | GGDEF family protein | -2.87 | 3.97E-17 |
| VC1394 | methyl-accepting chemotaxis protein | -6.14 | 2.73E-48 |
| VC1395 | response regulator CheY1 | -9.46 | 1.30E-78 |
| VC1396 | hypothetical protein | -12.14 | 5.03E-61 |
| VC1397 | chemotaxis protein CheA | -17.12 | 2.74E-103 |
| VC1398 | chemotaxis protein CheY | -19.30 | 6.90E-66 |
| VC1399 | chemotaxis protein methyltransferase CheR | -19.21 | 3.61E-85 |
| VC1400 | hypothetical protein | -10.21 | 1.92E-52 |
| VC1401 | protein-glutamate methylesterase CheB | -9.36 | 8.25E-49 |
| VC1402 | purine-binding chemotaxis protein Chew,putative | -9.80 | 2.22E-48 |
| VC1403 | methyl-accepting chemotaxis protein | -12.04 | 1.81E-69 |
| VC1406 | methyl-accepting chemotaxis protein | -4.82 | 4.38E-40 |
| VC1409 | multidrug resistance protein, putative | -4.64 | 1.55E-19 |
| VC1415 | hcp protein | -2.16 | 1.14E-05 |
| VC1433 | conserved hypothetical protein | -3.23 | 2.02E-15 |
| VC1437 | cation transport ATPase, E1-E2 family | -2.33 | 5.57E-22 |
| VC1444 | hypothetical protein | -2.25 | 7.37E-13 |
| VC1445 | sensor histidine kinase/response regulator | -3.50 | 6.41E-24 |
| VC1468 | conserved hypothetical protein | -2.34 | 1.06E-05 |
| VC1470 | XRE family transcriptional regulator | -2.38 | 3.22E-03 |
| VC1474 | conserved hypothetical protein | -2.60 | 1.23E-05 |
| VC1524 | ABC transporter, permease protein | -3.05 | 3.34E-12 |
| VC1539 | conserved hypothetical protein | -2.21 | 2.01E-16 |
| VC1560 | catalase/peroxidase | -4.78 | 1.21E-26 |
| VC1563 | conserved hypothetical protein | -4.71 | 1.41E-16 |
| VC1565 | outer membrane protein TolC, putative | -7.31 | 1.13E-21 |
| VC1566 | conserved hypothetical protein | -3.19 | 2.20E-10 |
| VC1567 | hypothetical protein | -2.09 | 5.06E-08 |
| VC1568 | ABC transporter, ATP-binding protein | -3.37 | 7.73E-11 |
| VC1583 | superoxide dismutase, Cu-Zn | -2.12 | 2.47E-05 |
| VC1590 | acetolactate synthase | -6.14 | 2.76E-27 |
| VC1591 | oxidoreductase, short-chaindehydrogenase/reduc tase family | -6.96 | 5.22E-22 |
| VC1593 | GGDEF family protein | -4.35 | 1.66E-28 |
| VC1594 | aldose 1-epimerase | -41.74 | 1.37E-180 |
| VC1595 | galactokinase | -101.86 | 0.00E+00 |
| VC1596 | galactose-1-phosphat e uridylyltransferase | -80.75 | 0.00E+00 |
| VC1598 | ABC transporter, periplasmic substrate-bindingprotein-relat ed prote | -3.43 | 2.47E-20 |
| VC1602 | chemotaxis protein CheV | -2.51 | 3.46E-12 |
| VC1603 | hypothetical protein | -2.37 | 3.05E-15 |
| VC1606 | conserved hypothetical protein | -2.04 | 1.21E-09 |
| VC1608 | conserved hypothetical protein | -2.13 | 1.33E-07 |
| VC1618 | multidrug resistance protein, putative | -2.25 | 9.81E-11 |
| VC1620 | c-di-GMP-regulated adhesin | -2.34 | 9.24E-06 |
| VC1623 | carboxynorspermidine decarboxylase | -3.20 | 3.55E-18 |
| VC1624 | conserved hypothetical protein | -2.61 | 3.57E-25 |
| VC1627 | Na+/H+ antiporter protein | -2.26 | 9.17E-19 |
| VC1641 | conserved hypothetical protein | -2.84 | 3.41E-19 |
| VC1643 | methyl-accepting chemotaxis protein | -11.04 | 2.84E-80 |
| VC1651 | response regulator VieB | -4.73 | 1.17E-26 |
| VC1652 | response regulator VieA | -3.53 | 2.34E-20 |

| | | | |
|---|---|---|---|
| VC1653 | sensory box sensor histidine kinase/responseregulator VieS | -2.67 | 2.85E-19 |
| VC1659 | conserved hypothetical protein | -2.03 | 1.97E-13 |
| VC1660a | ABC transporter permease | -3.09 | 9.50E-17 |
| VC1662 | conserved hypothetical protein | -2.19 | 6.15E-10 |
| VC1672 | DNA-3-methyladenine glycosidase I | -2.36 | 2.18E-05 |
| VC1673 | transporter, AcrB/D/F family | -4.64 | 9.34E-32 |
| VC1674 | periplasmic linker protein, putative | -5.73 | 2.69E-35 |
| VC1679 | psp operon transcriptional activator | -2.11 | 4.09E-06 |
| VC1689 | hypothetical protein | -2.19 | 1.84E-12 |
| VC1695 | formate transporter 1, putative | -2.28 | 8.13E-08 |
| VC1710 | conserved hypothetical protein | -2.47 | 2.80E-12 |
| VC1726 | glycogen synthase | -3.19 | 1.89E-34 |
| VC1727 | glucose-1-phosphate adenylyltransferase | -3.02 | 1.67E-16 |
| VC1736 | arginyl-tRNA-protein transferase-relatedprotein | -4.31 | 5.05E-26 |
| VC1746 | transcriptional regulator, TetR family | -2.40 | 1.36E-09 |
| VC1750 | hypothetical protein | -2.08 | 9.96E-06 |
| VC1756 | periplasmic linker protein, putative | -3.89 | 2.48E-31 |
| VC1757 | transporter, AcrB/D/F family | -3.76 | 2.21E-33 |
| VC1777 | conserved hypothetical protein | -2.21 | 3.51E-09 |
| VC1821 | PTS system, fructose-specific IIBC component | -2.08 | 4.90E-07 |
| VC1831 | sensor histidine kinase | -2.20 | 2.73E-10 |
| VC1841 | conserved hypothetical protein | -2.07 | 4.83E-10 |
| VC1842 | conserved hypothetical protein | -2.74 | 1.29E-05 |
| VC1843 | cytochrome d ubiquinol oxidase, subunit II | -3.72 | 1.92E-22 |
| VC1844 | cytochrome d ubiquinol oxidase, subunit I | -3.92 | 3.11E-20 |
| VC1851 | conserved hypothetical protein | -4.04 | 4.77E-25 |
| VC1867 | conserved hypothetical protein | -2.03 | 1.62E-11 |
| VC1868 | methyl-accepting chemotaxis protein | -6.30 | 3.06E-40 |
| VC1872 | conserved hypothetical protein | -8.71 | 8.50E-25 |
| VC1873 | conserved hypothetical protein | -12.76 | 3.48E-99 |
| VC1874 | conserved hypothetical protein | -6.23 | 1.64E-31 |
| VC1933 | hypothetical protein | -5.21 | 3.17E-59 |
| VC1934 | GGDEF family protein | -4.81 | 2.03E-27 |
| VC1950 | biotin sulfoxide reductase | -3.16 | 2.93E-18 |
| VC1952 | chitinase | -2.52 | 1.66E-12 |
| VC1979 | deoxyguanosinetripho sphate triphosphohydrolase | -2.89 | 6.91E-28 |
| VC1987 | outer membrane lipoprotein Slp, putative | -2.02 | 3.06E-14 |
| VC1991 | hypothetical protein | -3.48 | 1.87E-28 |
| VC2000 | glyceraldehyde 3-phosphate dehydrogenase | -2.29 | 2.71E-05 |
| VC2001 | conserved hypothetical protein | -3.87 | 6.61E-21 |
| VC2007 | transcriptional regulator, ROK family | -3.04 | 1.18E-29 |
| VC2008 | pyruvate kinase II | -2.85 | 5.99E-32 |
| VC2009 | conserved hypothetical protein | -2.21 | 4.49E-08 |
| VC2013 | PTS system, glucose-specific IIBC component | -6.28 | 2.99E-36 |
| VC2032 | hypothetical protein | -2.26 | 8.72E-09 |
| VC2058 | hypothetical protein | -2.79 | 6.19E-22 |
| VC2059 | purine-binding chemotaxis protein CheW | -2.16 | 6.27E-09 |
| VC2060 | conserved hypothetical protein | -4.48 | 3.75E-36 |
| VC2061 | ParA family protein | -2.58 | 2.17E-16 |
| VC2062 | protein-glutamate methylesterase CheB | -2.56 | 4.66E-16 |
| VC2063 | chemotaxis protein CheA | -2.90 | 6.15E-11 |
| VC2064 | chemotaxis protein CheZ | -2.52 | 4.12E-13 |
| VC2085 | succinyl-CoA synthetase subunit beta | -2.20 | 9.44E-07 |
| VC2120 | flagellar biosynthetic protein FlhB | -2.24 | 2.84E-10 |
| VC2121 | flagellar biosynthetic protein FliR | -2.70 | 3.40E-13 |
| VC2123 | flagellar biosynthetic protein FliP | -3.61 | 5.50E-21 |
| VC2124 | flagellar protein FliO | -4.26 | 2.17E-32 |
| VC2125 | flagellar motor switch protein FliN | -3.22 | 1.64E-25 |
| VC2126 | flagellar motor switch protein FliM | -3.29 | 3.23E-20 |
| VC2127 | flagellar protein FliL, putative | -2.68 | 2.87E-16 |
| VC2128 | flagellar hook-length control protein FliK,putative | -3.29 | 9.33E-20 |
| VC2130 | flagellum-specific ATP synthase FliI | -2.31 | 2.70E-11 |
| VC2142 | flagellin FlaB | -2.42 | 4.07E-05 |
| VC2143 | flagellin FlaD | -2.34 | 1.17E-04 |
| VC2161 | methyl-accepting chemotaxis protein | -4.03 | 5.86E-24 |
| VC2166 | Trp repressor-binding protein | -2.12 | 9.64E-09 |
| VC2167 | hypothetical protein | -2.14 | 2.11E-05 |
| VC2188 | flagellin core protein A | -2.07 | 2.84E-04 |
| VC2190 | flagellar hook-associated protein FlgL | -2.49 | 2.23E-13 |

| | | | |
|---|---|---|---|
| VC2191 | flagellar hook-associated protein FlgM | -2.33 | 5.65E-11 |
| VC2192 | flagellar protein FlgJ | -2.20 | 1.18E-09 |
| VC2193 | flagellar P-ring protein FlgI | -3.30 | 1.92E-25 |
| VC2195 | flagellar basal-body rod protein FlgG | -2.12 | 4.19E-09 |
| VC2196 | flagellar basal-body rod protein FlgF | -2.25 | 1.86E-10 |
| VC2197 | flagellar hook protein FlgE | -2.02 | 9.99E-06 |
| VC2198 | basal-body rod modification protein FlgD | -2.10 | 1.88E-07 |
| VC2201 | chemotaxis protein methyltransferase CheR | -4.29 | 8.95E-23 |
| VC2202 | chemotaxis protein CheV | -3.54 | 3.24E-20 |
| VC2206 | conserved hypothetical protein | -2.86 | 2.57E-16 |
| VC2207 | hypothetical protein | -2.95 | 2.24E-18 |
| VC2208 | hypothetical protein | -2.30 | 1.03E-13 |
| VC2240 | decarboxylase | -8.45 | 7.67E-45 |
| VC2241 | cytochrome c554 | -2.61 | 1.89E-15 |
| VC2246 | ribonuclease HII | -2.52 | 8.60E-09 |
| VC2247 | lipid-A-disaccharide synthase | -2.24 | 1.03E-08 |
| VC2248 | acyl-(acyl-carrier-p rotein)--UDP-N-acetylglucosami ne O-ac | -2.90 | 1.24E-20 |
| VC2249 | (3R)-hydroxymyristoyl-ACP dehydratase | -3.43 | 1.24E-27 |
| VC2250 | UDP-3-O-[3-hydroxymyristoyl] glucosamine N-acyltransferas | -2.99 | 1.60E-20 |
| VC2264 | conserved hypothetical protein | -2.75 | 5.62E-20 |
| VC2275 | transcriptional regulator Crl | -4.45 | 4.01E-31 |
| VC2285 | GGDEF family protein | -2.48 | 1.06E-13 |
| VC2314 | hypothetical protein | -3.17 | 1.47E-07 |
| VC2320 | exodeoxyribonuclease V, 135 kDa subunit | -2.73 | 8.75E-29 |
| VC2333 | ribosomal protein S6 modificationprotein-related protein | -2.01 | 1.15E-13 |
| VC2336 | methionyl-tRNA synthetase-related protein | -2.51 | 2.23E-08 |
| VC2337 | transcriptional regulator, LacI family | -2.63 | 9.04E-28 |
| VC2338 | ß-lactamase lacZ | -24.31 | 2.47E-129 |
| VC2340 | conserved hypothetical protein | -4.09 | 7.89E-32 |
| VC2344 | hypothetical protein | -2.33 | 1.02E-11 |
| VC2348 | phosphopentomutase | -3.10 | 1.12E-24 |
| VC2362 | threonine synthase | -4.03 | 4.91E-30 |
| VC2363 | homoserine kinase | -4.30 | 7.69E-53 |
| VC2364 | bifunctional aspartokinase I/homoserine dehydrogenase I | -2.89 | 3.87E-28 |
| VC2366 | s-adenosylmethionine :2-demethylmenaquinonemethyltr ansferase | -4.50 | 1.33E-43 |
| VC2367 | hypothetical protein | -2.27 | 1.09E-11 |
| VC2370 | sensory box/GGDEF family protein | -3.78 | 8.35E-19 |
| VC2371 | conserved hypothetical protein | -2.85 | 2.05E-07 |
| VC2374 | glutamate synthase subunit beta | -4.66 | 5.34E-26 |
| VC2378 | conserved hypothetical protein | -4.06 | 7.71E-32 |
| VC2384 | conserved hypothetical protein | -2.72 | 3.42E-12 |
| VC2413 | dihydrolipoamide acetyltransferase | -2.38 | 6.22E-17 |
| VC2429 | conserved hypothetical protein | -2.87 | 2.69E-06 |
| VC2436 | outer membrane channel protein | -2.08 | 2.20E-08 |
| VC2438 | glutamate-ammonia-li gase adenylyltransferase | -2.25 | 1.62E-18 |
| VC2454 | GGDEF family protein | -3.04 | 5.23E-22 |
| VC2455 | hypothetical protein | -5.83 | 2.97E-29 |
| VC2456 | hypothetical protein | -5.41 | 7.94E-59 |
| VC2464 | sigma-E factor regulatory protein RseC | -11.06 | 1.60E-118 |
| VC2465 | sigma-E factor regulatory protein RseB | -10.98 | 1.52E-116 |
| VC2466 | sigma-E factor negative regulatory protein RseA | -17.25 | 1.45E-202 |
| VC2467 | RNA polymerase sigma-E factor | -78682.69 | 1.92E-07 |
| VC2469 | l-aspartate oxidase | -2.71 | 1.76E-19 |
| VC2487 | conserved hypothetical protein | -2.89 | 3.81E-13 |
| VC2491 | 3-isopropylmalate dehydrogenase | -2.46 | 1.03E-16 |
| VC2492 | 3-isopropylmalate dehydratase, large subunit | -3.47 | 1.14E-27 |
| VC2493 | 3-isopropylmalate dehydratase small subunit | -3.96 | 1.31E-46 |
| VC2494 | hypothetical protein | -3.74 | 8.42E-23 |
| VC2497 | conserved hypothetical protein | -2.09 | 6.80E-17 |
| VC2507 | conserved hypothetical protein | -2.77 | 2.15E-19 |
| VC2529 | RNA polymerase sigma-54 factor | -3.13 | 3.95E-13 |
| VC2530 | sigma-54 modulation protein, putative | -3.36 | 1.37E-44 |
| VC2550 | conserved hypothetical protein | -2.41 | 3.33E-14 |
| VC2565 | elaA protein | -10.52 | 4.34E-96 |
| VC2566 | conserved hypothetical protein | -6.24 | 8.02E-48 |
| VC2605 | hypothetical protein | -2.15 | 8.09E-09 |
| VC2613 | phosphoribulokinase | -4.69 | 8.49E-59 |

| | | | |
|---|---|---|---|
| VC2615 | conserved hypothetical protein | -2.60 | 6.67E-10 |
| VC2622 | hypothetical protein | -3.64 | 4.30E-20 |
| VC2630 | fimbrial assembly protein | -2.34 | 6.34E-15 |
| VC2632 | fimbrial assembly protein PilO, putative | -2.21 | 5.71E-03 |
| VC2677 | transcriptional repressor, LacI family | -4.94 | 1.08E-47 |
| VC2697 | GGDEF family protein | -2.64 | 4.07E-12 |
| VC2702 | transcriptional regulator, LuxR family | -4.99 | 8.51E-39 |
| VC2703 | conserved hypothetical protein | -2.44 | 5.54E-22 |
| VC2711 | ATP-dependent DNA helicase RecG | -2.21 | 1.54E-09 |
| VC2722 | cysQ protein | -2.48 | 2.17E-11 |
| VC2725 | general secretion pathway protein L | -2.02 | 2.69E-08 |
| VC2726 | general secretion pathway protein K | -3.05 | 5.86E-20 |
| VC2727 | general secretion pathway protein J | -3.02 | 5.77E-08 |
| VC2728 | general secretion pathway protein I | -3.89 | 7.83E-15 |
| VC2729 | general secretion pathway protein H | -2.82 | 9.78E-29 |
| VC2730 | general secretion pathway protein G | -2.64 | 2.61E-26 |
| VC2738 | phosphoenolpyruvate carboxykinase | -2.98 | 2.20E-07 |
| VC2746 | glutamine synthetase | -2.10 | 4.18E-08 |
| VC2750 | GGDEF family protein | -2.17 | 3.98E-08 |
| VC2751 | adenosine deaminase | -3.22 | 6.13E-36 |
| VCA0008 | methyl-accepting chemotaxis protein | -7.91 | 2.92E-42 |
| VCA0009 | hypothetical protein | -5.82 | 2.46E-19 |
| VCA0015 | hypothetical protein | -2.33 | 1.52E-17 |
| VCA0018 | vgrG protein | -2.14 | 5.78E-06 |
| VCA0025 | transporter, NadC family | -2.19 | 1.52E-15 |
| VCA0030 | hypothetical protein | -2.11 | 6.80E-09 |
| VCA0031 | methyl-accepting chemotaxis protein | -10.20 | 2.69E-64 |
| VCA0032 | hypothetical protein | -8.72 | 1.62E-66 |
| VCA0033 | hypothetical protein | -2.30 | 1.21E-10 |
| VCA0034 | conserved hypothetical protein | -15.84 | 2.27E-93 |
| VCA0047 | conserved hypothetical protein | -3.11 | 1.63E-21 |
| VCA0048 | conserved hypothetical protein | -2.90 | 1.76E-15 |
| VCA0049 | GGDEF family protein | -2.21 | 2.38E-10 |
| VCA0051 | hypothetical protein | -2.30 | 1.07E-15 |
| VCA0052 | hypothetical protein | -2.81 | 2.26E-16 |
| VCA0074 | GGDEF family protein | -2.35 | 3.95E-11 |
| VCA0080 | GGDEF family protein | -2.39 | 5.04E-13 |
| VCA0101 | conserved hypothetical protein | -2.09 | 1.10E-08 |
| VCA0105 | conserved hypothetical protein | -2.83 | 3.11E-08 |
| VCA0106 | hypothetical protein | -2.50 | 4.49E-09 |
| VCA0109 | hypothetical protein | -2.15 | 1.21E-07 |
| VCA0111 | hypothetical protein | -2.35 | 8.06E-08 |
| VCA0112 | hypothetical protein | -3.34 | 1.81E-20 |
| VCA0113 | hypothetical protein | -5.15 | 3.10E-27 |
| VCA0114 | hypothetical protein | -4.44 | 7.33E-21 |
| VCA0115 | hypothetical protein | -4.99 | 4.84E-20 |
| VCA0116 | clpB protein | -3.98 | 2.19E-34 |
| VCA0117 | sigma-54 dependent transcriptional regulator | -3.66 | 2.65E-23 |
| VCA0118 | hypothetical protein | -4.74 | 1.52E-14 |
| VCA0119 | hypothetical protein | -4.41 | 1.38E-19 |
| VCA0120 | IcmF-related protein | -2.84 | 8.76E-22 |
| VCA0121 | hypothetical protein | -3.64 | 7.67E-23 |
| VCA0122 | hypothetical protein | -2.89 | 8.93E-06 |
| VCA0129 | ribose ABC transporter permease | -2.16 | 7.31E-07 |
| VCA0130 | ribose ABC transporter, periplasmicD-ribose-binding protein | -3.58 | 1.13E-21 |
| VCA0131 | ribokinase | -2.21 | 1.75E-07 |
| VCA0136 | glycerophosphoryl diester phosphodiesterase | -2.45 | 7.26E-13 |
| VCA0137 | glycerol-3-phosphate transporter | -3.30 | 2.15E-18 |
| VCA0153 | conserved hypothetical protein | -4.32 | 7.50E-20 |
| VCA0154 | conserved hypothetical protein | -3.22 | 4.65E-15 |
| VCA0155 | NADH dehydrogenase, putative | -4.23 | 3.06E-27 |
| VCA0156 | conserved hypothetical protein | -3.55 | 1.60E-19 |
| VCA0157 | NADH dehydrogenase, putative | -3.21 | 1.65E-20 |
| VCA0165 | GGDEF family protein | -4.21 | 1.03E-29 |
| VCA0168 | hypothetical protein | -4.44 | 2.02E-54 |
| VCA0171 | conserved hypothetical protein | -2.15 | 3.34E-09 |
| VCA0172 | conserved hypothetical protein | -3.48 | 1.71E-12 |
| VCA0173 | hypothetical protein | -3.80 | 2.12E-20 |
| VCA0174 | conserved hypothetical protein | -2.14 | 7.05E-05 |

| | | | |
|---|---|---|---|
| VCA0175 | MoxR-related protein | -4.72 | 3.14E-20 |
| VCA0189 | response regulator | -2.16 | 1.04E-11 |
| VCA0190 | hypothetical protein | -2.22 | 5.55E-09 |
| VCA0191 | conserved hypothetical protein | -3.11 | 2.93E-18 |
| VCA0200 | hypothetical protein | -2.17 | 1.25E-11 |
| VCA0201 | hypothetical protein | -2.59 | 6.99E-15 |
| VCA0205 | C4-dicarboxylate transporter, anaerobic | -5.23 | 3.86E-29 |
| VCA0210 | response regulator, putative | -5.39 | 2.21E-38 |
| VCA0211 | sensory box sensor histidine kinase | -5.30 | 3.47E-40 |
| VCA0219 | haemolysin | -4.94 | 1.28E-36 |
| VCA0220 | hemolysin secretion protein HylB | -2.58 | 6.81E-19 |
| VCA0223 | protease | -2.85 | 1.16E-16 |
| VCA0224 | hypothetical protein | -2.19 | 1.40E-05 |
| VCA0265 | hypothetical protein | -4.36 | 3.14E-26 |
| VCA0268 | methyl-accepting chemotaxis protein | -3.76 | 2.36E-15 |
| VCA0274a | hypothetical protein | -2.42 | 1.38E-10 |
| VCA0281 | integrase, putative | -2.72 | 6.35E-17 |
| VCA0282 | IS5 transposase | -3.61 | 7.72E-22 |
| VCA0283 | hypothetical protein | -2.42 | 1.09E-10 |
| VCA0301 | oxidoreductase, short-chaindehydrogenase/reduc tase family | -2.07 | 1.34E-11 |
| VCA0302 | hypothetical protein | -2.95 | 5.39E-13 |
| VCA0303 | hypothetical protein | -4.41 | 5.55E-54 |
| VCA0304 | hypothetical protein | -2.86 | 4.07E-10 |
| VCA0306 | hypothetical protein | -2.58 | 3.79E-10 |
| VCA0307 | hypothetical protein | -2.82 | 7.53E-18 |
| VCA0309 | conserved hypothetical protein | -3.19 | 6.68E-19 |
| VCA0315 | hypothetical protein | -2.44 | 8.31E-08 |
| VCA0317 | lipoprotein Blc | -2.58 | 1.85E-13 |
| VCA0328 | biphenyl-2,3-diol 1,2-dioxygenase III-relatedprotein | -3.97 | 5.12E-36 |
| VCA0329 | hypothetical protein | -2.02 | 3.93E-06 |
| VCA0329b | hypothetical protein | -2.07 | 1.94E-04 |
| VCA0331 | hypothetical protein | -2.35 | 1.55E-11 |
| VCA0337 | microcin immunity protein MccF | -3.31 | 2.33E-23 |
| VCA0338 | conserved hypothetical protein | -3.28 | 2.16E-18 |
| VCA0340 | hypothetical protein | -3.09 | 2.79E-17 |
| VCA0341 | biphenyl-2,3-diol 1,2-dioxygenase III-relatedprotein | -2.31 | 9.27E-10 |
| VCA0353 | hypothetical protein | -2.71 | 5.75E-03 |
| VCA0365 | hypothetical protein | -10.53 | 2.60E-04 |
| VCA0366 | hypothetical protein | -2.53 | 5.14E-13 |
| VCA0381 | hypothetical protein | -2.47 | 9.42E-07 |
| VCA0416 | hypothetical protein | -2.01 | 1.18E-04 |
| VCA0417 | acetyltransferase, putative | -2.93 | 9.66E-16 |
| VCA0439 | microcin immunity protein MccF | -2.83 | 2.20E-20 |
| VCA0440 | hypothetical protein | -2.29 | 2.78E-06 |
| VCA0442 | acetyltransferase, putative | -2.89 | 9.49E-12 |
| VCA0511 | anaerobic ribonucleoside-triphosphate reductase | -2.89 | 2.14E-21 |
| VCA0512 | anaerobic ribonucleoside-triphosphate reductaseactivating protein | -2.34 | 1.18E-07 |
| VCA0516 | PTS system, fructose-specific IIBC component | -2.13 | 3.65E-15 |
| VCA0526 | conserved hypothetical protein | -3.16 | 1.03E-11 |
| VCA0536 | conserved hypothetical protein | -2.06 | 3.19E-16 |
| VCA0544 | conserved hypothetical protein | -5.68 | 1.69E-56 |
| VCA0563 | NAD(P) transhydrogenase, alpha subunit | -2.05 | 3.93E-16 |
| VCA0570 | Sui1 family protein | -4.11 | 5.21E-30 |
| VCA0571 | hypothetical protein | -2.97 | 7.34E-18 |
| VCA0573 | DamX-related protein | -2.28 | 8.59E-21 |
| VCA0574 | conserved hypothetical protein | -3.05 | 5.92E-38 |
| VCA0583 | hypothetical protein | -4.76 | 3.87E-27 |
| VCA0584 | glutathione S-transferase, putative | -3.07 | 1.59E-30 |
| VCA0588 | peptide ABC transporter, ATP-binding protein,putative | -2.41 | 7.71E-11 |
| VCA0589 | peptide ABC transporter, permease protein,putative | -3.66 | 2.79E-11 |
| VCA0590 | peptide ABC transporter, permease protein,putative | -4.33 | 3.60E-16 |
| VCA0591 | peptide ABC transporter, periplasmicpeptide-binding protein, putative | -3.95 | 1.42E-43 |
| VCA0592 | MutT/nudix family protein | -2.44 | 5.79E-22 |
| VCA0593 | hypothetical protein | -4.26 | 6.16E-26 |
| VCA0607 | regulator of nucleoside diphosphate kinase | -2.11 | 8.41E-16 |
| VCA0620 | thiosulfate sulfurtransferase SseA, putative | -2.23 | 1.14E-17 |
| VCA0624 | transketolase 1 | -2.31 | 3.04E-12 |
| VCA0637 | oxygen-insensitive NAD(P)H nitroreductase | -2.50 | 3.00E-07 |
| VCA0638 | transporter, AcrB/D/F family | -6.26 | 1.23E-52 |
| VCA0639 | AcrA/AcrE family protein | -5.56 | 8.46E-64 |

| | | | |
|---|---|---|---|
| VCA0644 | NADH oxidase, putative | -2.02 | 1.11E-07 |
| VCA0646 | conserved hypothetical protein/hemolysin,putative | -5.01 | 4.16E-26 |
| VCA0647 | hypothetical protein | -6.07 | 1.36E-19 |
| VCA0648 | hypothetical protein | -3.54 | 9.82E-08 |
| VCA0649 | hypothetical protein | -2.79 | 1.30E-05 |
| VCA0650 | hypothetical protein | -5.35 | 2.05E-13 |
| VCA0651 | conserved hypothetical protein | -4.73 | 1.72E-05 |
| VCA0653 | PTS system, sucrose-specific IIBC component | -2.85 | 1.04E-08 |
| VCA0666 | L-serine dehydratase 1 | -3.95 | 4.19E-30 |
| VCA0669 | sugar transporter family protein | -2.41 | 1.68E-07 |
| VCA0676 | iron-sulfur cluster-binding protein NapF | -3.11 | 7.72E-13 |
| VCA0677 | napD protein | -2.30 | 7.17E-05 |
| VCA0678 | periplasmic nitrate reductase | -2.96 | 2.02E-13 |
| VCA0679 | periplasmic nitrate reductase, cytochrome c-typeprotein | -2.70 | 9.13E-06 |
| VCA0680 | periplasmic nitrate reductase, cytochrome c-typeprotein | -2.22 | 1.89E-05 |
| VCA0681 | conserved hypothetical protein | -4.36 | 3.46E-36 |
| VCA0684 | regulatory protein UhpC | -2.30 | 8.38E-14 |
| VCA0686 | iron(III) ABC transporter, permease protein | -2.24 | 2.79E-18 |
| VCA0687 | ferric transporter ATP-binding subunit | -2.28 | 7.45E-14 |
| VCA0688 | polyhydroxyalkanoic acid synthase | -4.59 | 6.57E-23 |
| VCA0689 | conserved hypothetical protein | -2.22 | 2.42E-04 |
| VCA0698 | hypothetical protein | -5.51 | 1.08E-38 |
| VCA0704 | phosphoglycerate transport systemtranscriptional regulatory protein | -2.40 | 3.12E-10 |
| VCA0719 | sensor histide kinase | -7.03 | 2.41E-47 |
| VCA0720 | guanylate cyclase-related protein | -6.48 | 9.43E-79 |
| VCA0728 | hypothetical protein | -3.01 | 4.84E-17 |
| VCA0729 | hypothetical protein | -3.35 | 4.29E-29 |
| VCA0731 | hypothetical protein | -2.90 | 1.78E-09 |
| VCA0736 | sensor histidine kinase LuxQ | -3.78 | 1.20E-27 |
| VCA0737 | luxP protein | -2.77 | 1.08E-28 |
| VCA0738 | hypothetical protein | -7.06 | 7.02E-45 |
| VCA0743 | conserved hypothetical protein | -3.89 | 9.25E-26 |
| VCA0744 | glycerol kinase | -4.50 | 5.04E-27 |
| VCA0762 | hypothetical | -2.86 | 1.21E-14 |
| VCA0763 | conserved hypothetical protein | -2.45 | 2.15E-21 |
| VCA0764 | MutT/nudix family protein | -2.61 | 8.80E-12 |
| VCA0798 | CbbY family protein | -2.17 | 6.82E-17 |
| VCA0802 | conserved hypothetical protein | -3.43 | 3.57E-43 |
| VCA0803 | serine protease, putative | -5.29 | 5.57E-39 |
| VCA0822 | aspartokinase, putative | -2.04 | 2.57E-09 |
| VCA0829 | acetyl-CoA synthase | -3.20 | 1.48E-21 |
| VCA0851 | conserved hypothetical protein | -2.61 | 1.06E-08 |
| VCA0870 | D-alanyl-D-alanine endopeptidase | -2.60 | 8.34E-17 |
| VCA0877 | hydrolase, putative | -5.18 | 8.45E-44 |
| VCA0878 | integrase-related protein | -8.11 | 4.38E-71 |
| VCA0880 | hypothetical protein | -7.70 | 6.63E-53 |
| VCA0881 | hypothetical protein | -5.39 | 7.30E-31 |
| VCA0882 | hypothetical protein | -6.43 | 4.05E-30 |
| VCA0883 | hypothetical protein | -3.40 | 2.07E-12 |
| VCA0884 | hypothetical protein | -6.27 | 3.10E-40 |
| VCA0895 | chemotactic transducer-related protein | -6.15 | 1.19E-55 |
| VCA0900 | hypothetical protein | -3.98 | 1.43E-29 |
| VCA0901 | hypothetical protein | -2.16 | 2.96E-09 |
| VCA0906 | methyl-accepting chemotaxis protein | -8.73 | 1.12E-75 |
| VCA0915 | hemin importer ATP-binding protein | -3.67 | 1.88E-24 |
| VCA0917 | transcriptional regulator, TetR family | -2.66 | 2.94E-14 |
| VCA0920 | hypothetical protein | -2.06 | 1.09E-09 |
| VCA0923 | methyl-accepting chemotaxis protein | -4.94 | 7.94E-61 |
| VCA0931 | conserved hypothetical protein | -2.53 | 1.76E-14 |
| VCA0936 | hypothetical protein | -2.83 | 2.77E-10 |
| VCA0951 | conserved hypothetical protein | -9.24 | 1.06E-28 |
| VCA0965 | GGDEF family protein | -7.93 | 8.88E-45 |
| VCA0966 | hypothetical protein | -5.70 | 8.07E-29 |
| VCA0975 | ATP-dependent protease LA-related protein | -7.10 | 7.15E-64 |
| VCA0978 | amino acid ABC transporter, periplasmic aminoacid-binding protein, | -14.38 | 5.27E-96 |
| VCA0979 | methyl-accepting chemotaxis protein | -7.77 | 2.89E-58 |
| VCA0981 | hypothetical protein | -11.43 | 2.01E-79 |
| VCA0984 | L-lactate dehydrogenase | -4.59 | 1.39E-16 |
| VCA0985 | oxidoreductase/iron- sulfur cluster-bindingprotein | -6.31 | 2.65E-39 |
| VCA0986 | conserved hypothetical protein | -2.65 | 9.09E-14 |

| | | | |
|---|---|---|---|
| VCA0987 | phosphoenolpyruvate synthase | -2.91 | 3.49E-19 |
| VCA0988 | methyl-accepting chemotaxis protein | -5.67 | 9.43E-33 |
| VCA0993 | N-ethylmaleimide reductase | -2.56 | 5.78E-12 |
| VCA0994 | hypothetical protein | -2.25 | 2.51E-13 |
| VCA0998 | NADH-dependent flavin oxidoreductase, Oyefamily | -3.20 | 1.16E-10 |
| VCA1011 | glutaredoxin-related protein | -2.04 | 8.45E-07 |
| VCA1015 | Na+/H+ antiporter | -6.55 | 1.09E-94 |
| VCA1016 | hypothetical protein | -3.81 | 9.08E-29 |
| VCA1017 | methylated-DNA--prot ein-cysteineS-methyltransferas e | -5.12 | 2.93E-36 |
| VCA1023 | hypothetical protein | -2.23 | 7.65E-11 |
| VCA1024 | hypothetical protein | -3.59 | 3.42E-22 |
| VCA1025 | glucosamine-6-phosphate deaminase | -2.38 | 3.24E-11 |
| VCA1029 | glycogen operon protein GlgX | -3.55 | 1.90E-28 |
| VCA1030 | hypothetical protein | -3.08 | 6.32E-27 |
| VCA1033 | extracellular solute-binding protein, putative | -6.68 | 2.09E-42 |
| VCA1034 | methyl-accepting chemotaxis protein | -9.98 | 6.86E-75 |
| VCA1036 | pseudogene | -2.10 | 1.46E-13 |
| VCA1038 | amino acid ABC transporter, permease protein | -2.87 | 1.27E-16 |
| VCA1039 | amino acid ABC transporter, periplasmic aminoacid-binding protein | -2.65 | 3.50E-27 |
| VCA1040 | amino acid ABC transporter, permease protein | -2.93 | 1.74E-23 |
| VCA1042 | Ccm2-related protein | -2.43 | 7.19E-13 |
| VCA1045 | PTS system, mannitol-specific IIABC component | -2.02 | 1.37E-09 |
| VCA1054 | conserved hypothetical protein | -4.20 | 1.84E-51 |
| VCA1056 | methyl-accepting chemotaxis protein | -5.07 | 9.89E-37 |
| VCA1067 | aldehyde dehydrogenase | -3.43 | 1.11E-20 |
| VCA1069 | methyl-accepting chemotaxis protein | -2.36 | 8.73E-15 |
| VCA1071 | sodium/proline symporter | -2.82 | 1.11E-16 |
| VCA1080 | secretion protein, HlyD family | -2.20 | 1.39E-13 |
| VCA1082 | hypothetical protein | -2.91 | 6.26E-24 |
| VCA1084 | toxin secretion ATP-binding protein | -2.35 | 1.52E-17 |
| VCA1086 | response regulator | -3.88 | 1.74E-20 |
| VCA1087 | anti-sigma F factor antagonist, putative | -2.72 | 2.10E-14 |
| VCA1088 | methyl-accepting chemotaxis protein | -3.15 | 6.83E-21 |
| VCA1089 | cheB3 methylesterase | -8.91 | 7.17E-63 |
| VCA1090 | chemotaxis protein CheD, putative | -7.52 | 1.14E-46 |
| VCA1091 | chemotaxis protein methyltransferase CheR | -5.22 | 2.24E-32 |
| VCA1092 | methyl-accepting chemotaxis protein | -5.87 | 9.07E-37 |
| VCA1093 | purine-binding chemotaxis protein CheW | -6.63 | 1.83E-37 |
| VCA1094 | purine-binding chemotaxis protein CheW | -5.99 | 9.54E-33 |
| VCA1095 | chemotaxis protein CheA | -5.70 | 2.79E-44 |
| VCA1096 | chemotaxis protein CheY | -6.75 | 8.92E-48 |
| VCA1097 | conserved hypothetical protein | -6.41 | 1.30E-72 |
| VCA1104 | phoP homolog | -2.96 | 9.29E-18 |
| VCA1105 | DNA-binding response regulator | -3.54 | 2.32E-22 |
| VCA1106 | hypothetical protein | -2.60 | 3.77E-15 |
| VCA1113 | spermidine/putrescin e ABC transporter,periplasmic spermidine/putr | -2.60 | 4.85E-10 |
| 23Sb | 23Sb | 2.65 | 9.10419E-06 |
| tRNA-Ala-2 | tRNA-Ala-2 | 2.37 | 1.25E-03 |
| tRNA-Ala-3 | tRNA-Ala-3 | 2.57 | 4.27E-03 |
| tRNA-Ala-5 | tRNA-Ala-5 | 4.69 | 6.08E-11 |
| tRNA-Arg- | tRNA-Arg-4 | 4.85 | 4.28E-24 |
| tRNA-Arg- | tRNA-Arg-5 | 5.59 | 5.30E-28 |
| tRNA-Arg- | tRNA-Arg-6 | 7.33 | 1.65E-08 |
| tRNA-Arg- | tRNA-Arg-7 | 5.10 | 8.07E-04 |
| tRNA-Arg- | tRNA-Arg-8 | 8.08 | 1.11E-07 |
| tRNA-Asn- | tRNA-Asn-1 | 6.11 | 2.00E-06 |
| tRNA-Asn- | tRNA-Asn-2 | 3.23 | 4.48E-05 |
| tRNA-Asn- | tRNA-Asn-3 | 3.67 | 7.49E-10 |
| tRNA-Asn- | tRNA-Asn-4 | 5.08 | 5.57E-04 |
| tRNA-Asp- | tRNA-Asp-4 | 2.98 | 9.36E-15 |
| tRNA-Asp- | tRNA-Asp-5 | 9.07 | 2.02E-12 |
| tRNA-Cys- | tRNA-Cys-1 | 5.10 | 4.09E-20 |
| tRNA-Gln- | tRNA-Gln-1 | 2.27 | 6.96E-04 |
| tRNA-Gln- | tRNA-Gln-3 | 2.46 | 1.82E-04 |
| tRNA-Gln- | tRNA-Gln-5 | 2.64 | 3.08E-04 |
| tRNA-Glu- | tRNA-Glu-2 | 4.41 | 3.99E-43 |
| tRNA-Glu- | tRNA-Glu-3 | 5.52 | 1.58E-27 |
| tRNA-Gly- | tRNA-Gly-1 | 8.36 | 5.15E-17 |
| tRNA-Gly- | tRNA-Gly-3 | 2.80 | 1.82E-17 |
| tRNA-Gly- | tRNA-Gly-4 | 2.92 | 1.51E-13 |

| | | | |
|---|---|---|---|
| tRNA-Gly- | tRNA-Gly-5 | 3.11 | 6.67E-11 |
| tRNA-Gly- | tRNA-Gly-6 | 5.16 | 1.63E-06 |
| tRNA-Gly- | tRNA-Gly-7 | 3.33 | 6.33E-06 |
| tRNA-His-1 | tRNA-His-1 | 3.67 | 8.56E-27 |
| tRNA-His-2 | tRNA-His-2 | 2.21 | 2.61E-05 |
| tRNA-Ile-1 | tRNA-Ile-1 | 7.64 | 2.50E-19 |
| tRNA-Ile-2 | tRNA-Ile-2 | 6.67 | 5.21E-17 |
| tRNA-Leu- | tRNA-Leu-1 | 5.09 | 5.71E-13 |
| tRNA-Leu- | tRNA-Leu-12 | 5.85 | 1.30E-07 |
| tRNA-Leu- | tRNA-Leu-2 | 2.56 | 5.61E-06 |
| tRNA-Leu- | tRNA-Leu-3 | 2.09 | 2.40E-04 |
| tRNA-Leu- | tRNA-Leu-4 | 4.94 | 3.32E-10 |
| tRNA-Leu- | tRNA-Leu-5 | 2.18 | 1.53E-10 |
| tRNA-Leu- | tRNA-Leu-6 | 2.19 | 3.39E-07 |
| tRNA-Leu- | tRNA-Leu-7 | 3.35 | 1.25E-09 |
| tRNA-Leu- | tRNA-Leu-8 | 5.53 | 4.61E-24 |
| tRNA-Leu- | tRNA-Leu-9 | 2.85 | 8.54E-04 |
| tRNA-Lys- | tRNA-Lys-1 | 3.64 | 7.05E-21 |
| tRNA-Lys- | tRNA-Lys-2 | 4.39 | 5.68E-19 |
| tRNA-Met- | tRNA-Met-2 | 3.56 | 2.05E-21 |
| tRNA-Met- | tRNA-Met-3 | 4.29 | 1.42E-23 |
| tRNA-Met- | tRNA-Met-4 | 4.91 | 2.19E-17 |
| tRNA-Met- | tRNA-Met-5 | 8.41 | 6.48E-21 |
| tRNA-Met- | tRNA-Met-6 | 6.98 | 2.81E-15 |
| tRNA-Met- | tRNA-Met-7 | 3.02 | 3.87E-04 |
| tRNA-Met- | tRNA-Met-9 | 2.17 | 2.12E-07 |
| tRNA-Phe- | tRNA-Phe-1 | 5.09 | 2.06E-38 |
| tRNA-Phe- | tRNA-Phe-2 | 5.79 | 3.29E-16 |
| tRNA-Phe- | tRNA-Phe-3 | 2.39 | 9.10E-19 |
| tRNA-Pro- | tRNA-Pro-2 | 4.08 | 5.37E-14 |
| tRNA-Ser- | tRNA-Ser-4 | 8.45 | 3.19E-43 |
| tRNA-Ser- | tRNA-Ser-5 | 7.49 | 6.15E-67 |
| tRNA-Thr- | tRNA-Thr-2 | 2.21 | 4.40E-13 |
| tRNA-Thr- | tRNA-Thr-3 | 2.22 | 1.07E-07 |
| tRNA-Thr- | tRNA-Thr-4 | 4.46 | 9.80E-14 |
| tRNA-Thr- | tRNA-Thr-5 | 7.09 | 6.36E-31 |
| tRNA-Thr- | tRNA-Thr-6 | 4.93 | 1.70E-17 |
| tRNA-Trp- | tRNA-Trp-1 | 3.26 | 1.23E-04 |
| tRNA-Tyr- | tRNA-Tyr-1 | 2.27 | 1.95E-14 |
| tRNA-Tyr- | tRNA-Tyr-2 | 2.90 | 2.79E-04 |
| tRNA-Tyr- | tRNA-Tyr-3 | 5.41 | 2.49E-14 |
| tRNA-Tyr- | tRNA-Tyr-4 | 4.66 | 9.34E-25 |
| tRNA-Tyr- | tRNA-Tyr-5 | 5.92 | 6.72E-25 |
| tRNA-Val-2 | tRNA-Val-2 | 2.81 | 1.71E-04 |
| tRNA-Val-3 | tRNA-Val-3 | 3.79 | 9.26E-20 |
| tRNA-Val-4 | tRNA-Val-4 | 3.36 | 3.40E-19 |
| VC0006 | ribonuclease P | 2.07 | 1.03E-10 |
| VC0007 | ribosomal protein L34 | 2.25 | 3.27E-09 |
| VC0011 | chromosome replication initiator DnaA | 2.30 | 2.46E-16 |
| VC0015a | hypothetical protein | 52.55 | 3.29E-87 |
| VC0018 | 16 kDa heat shock protein A | 3.30 | 4.73E-17 |
| VC0022 | conserved hypothetical protein | 2.65 | 3.35E-20 |
| VC0025 | hypothetical protein | 8.66 | 5.50E-51 |
| VC0038 | hypothetical protein | 2.10 | 1.06E-06 |
| VC0040 | hemolysin, putative | 3.14 | 6.65E-25 |
| VC0051 | phosphoribosylaminoi midazole carboxylase, ATPasesubunit | 2.02 | 4.16E-14 |
| VC0052 | phosphoribosylaminoi midazole carboxylase,catalytic subunit | 2.93 | 9.32E-29 |
| VC0059 | hypothetical protein | 5.79 | 8.37E-34 |
| VC0061 | thiamin biosynthesis protein ThiC | 3.71 | 3.40E-38 |
| VC0064 | thiS protein | 3.44 | 2.16E-03 |
| VC0069 | multidrug resistance protein, putative | 2.99 | 4.61E-18 |
| VC0070 | hypothetical protein | 3.37 | 6.71E-30 |
| VC0074 | hypothetical protein | 2.38 | 5.09E-10 |
| VC0083 | ubiquinone/menaquinone biosynthesis methyltransferase | 2.06 | 3.13E-11 |
| VC0093 | glycerol-3-phosphate acyltransferase | 3.10 | 2.50E-26 |
| VC0109 | ribosome biogenesis GTP-binding protein YsxC | 2.12 | 1.82E-14 |
| VC0113 | methyltransferase-re lated protein | 2.47 | 3.65E-14 |
| VC0114 | conserved hypothetical protein | 2.83 | 3.30E-28 |
| VC0115 | conserved hypothetical protein | 4.93 | 4.84E-46 |
| VC0120 | porphobilinogen deaminase | 2.16 | 7.11E-11 |

| | | | |
|---|---|---|---|
| VC0123 | cyaY protein | 3.10 | 2.39E-21 |
| VC0140 | conserved hypothetical protein | 2.55 | 3.71E-13 |
| VC0142a | hypothetical protein | 5.05 | 9.07E-22 |
| VC0143 | hypothetical protein | 17.24 | 1.92E-73 |
| VC0144 | conserved hypothetical protein | 2.15 | 2.18E-09 |
| VC0154 | tRNA (uracil-5-)-methyltransferase | 2.54 | 4.40E-16 |
| VC0156 | vitamin B12 receptor | 2.45 | 1.88E-14 |
| VC0194 | gamma-glutamyltransp eptidase | 2.16 | 1.41E-10 |
| VC0195 | rarD protein | 2.12 | 1.34E-06 |
| VC0196 | ATP-dependent DNA helicase RecQ | 3.14 | 3.20E-23 |
| VC0197 | gene 3 protein-related protein | 2.02 | 2.69E-05 |
| VC0201 | iron(III) ABC transporter, ATP-binding protein | 2.09 | 1.29E-03 |
| VC0210 | ribonuclease PH | 2.30 | 9.38E-12 |
| VC0218 | ribosomal protein L28 | 6.89 | 4.95E-37 |
| VC0219 | ribosomal protein L33 | 7.07 | 3.44E-55 |
| VC0223 | ADP-heptose--LPS heptosyltransferase II,putative | 2.14 | 4.15E-16 |
| VC0271 | conserved hypothetical protein | 2.62 | 2.06E-14 |
| VC0275 | phosphoribosylamine- -glycine ligase | 4.69 | 1.25E-31 |
| VC0276 | bifunctional phosphoribosylaminoimidazolecarboxamide formyltrans | 4.31 | 9.46E-36 |
| VC0290 | Fis family transcriptional regulator | 7.14 | 8.48E-66 |
| VC0291 | NifR3/Smm1 family protein | 5.49 | 8.62E-64 |
| VC0292 | ribosomal protein L11 methyltransferase | 3.02 | 2.00E-22 |
| VC0302 | conserved hypothetical protein | 2.17 | 2.55E-11 |
| VC0308 | hypothetical protein | 2.12 | 1.20E-11 |
| VC0339 | phosphatidylserine decarboxylase | 2.08 | 2.23E-08 |
| VC0341 | oligoribonuclease | 2.39 | 3.19E-20 |
| VC0342 | iron-sulfur cluster-binding protein | 2.05 | 2.11E-14 |
| VC0359 | ribosomal protein S12 | 5.48 | 5.49E-17 |
| VC0360 | ribosomal protein S7 | 4.94 | 2.98E-13 |
| VC0366 | ribosomal protein S6 | 3.07 | 4.55E-11 |
| VC0367 | primosomal replication protein N | 2.99 | 5.03E-14 |
| VC0368 | ribosomal protein S18 | 2.44 | 1.07E-11 |
| VC0369 | ribosomal protein L9 | 2.16 | 1.41E-08 |
| VC0370 | conserved hypothetical protein | 2.03 | 6.35E-04 |
| VC0390 | B12-dependent methionine synthase | 3.23 | 4.67E-20 |
| VC0397 | single-strand binding protein | 3.52 | 1.17E-24 |
| VC0424 | conserved hypothetical protein | 6.30 | 9.06E-44 |
| VC0425 | hypothetical protein | 2.25 | 6.07E-10 |
| VC0426 | hypothetical protein | 2.59 | 1.24E-06 |
| VC0428 | conserved hypothetical protein | 2.06 | 4.04E-09 |
| VC0431 | arginine repressor | 2.47 | 6.51E-13 |
| VC0435 | ribosomal protein L21 | 5.29 | 4.92E-15 |
| VC0436 | ribosomal protein L27 | 5.45 | 5.54E-40 |
| VC0437 | GTPase ObgE | 3.31 | 4.40E-20 |
| VC0438 | conserved hypothetical protein | 2.39 | 2.62E-13 |
| VC0451 | conserved hypothetical protein | 2.05 | 4.45E-06 |
| VC0453 | tRNA (guanine-N(7)-)-methyltransferase | 2.09 | 3.40E-15 |
| VC0469 | conserved hypothetical protein | 5.72 | 5.79E-48 |
| VC0476 | D-erythrose 4-phosphate dehydrogenase | 3.05 | 1.73E-29 |
| VC0483 | conserved hypothetical protein | 3.87 | 9.40E-26 |
| VC0509 | hypothetical protein | 2.77 | 1.08E-03 |
| VC0516 | phage integrase | 2.85 | 3.26E-21 |
| VC0518 | DNA primase | 2.09 | 2.10E-10 |
| VC0520 | 30S ribosomal protein S21 | 2.25 | 6.57E-21 |
| VC0527 | cell division protein FtsB | 3.23 | 4.81E-20 |
| VC0542 | CinA-related protein | 3.03 | 4.54E-31 |
| VC0543 | recombinase A | 2.19 | 1.20E-08 |
| VC0544 | recombination regulator RecX | 2.19 | 1.19E-06 |
| VC0561 | ribosomal protein S16 | 3.65 | 2.66E-22 |
| VC0562 | 16S rRNA processing protein RimM | 3.22 | 5.83E-14 |
| VC0564 | ribosomal protein L19 | 2.28 | 2.31E-21 |
| VC0576 | stringent starvation protein A | 2.37 | 3.02E-15 |
| VC0582 | conserved hypothetical protein | 2.38 | 1.08E-15 |
| VC0585 | hypoxanthine phosphoribosyltransferase | 2.92 | 9.63E-17 |
| VC0587 | sulfate permease family protein | 2.74 | 4.26E-17 |
| VC0596 | dnaK suppressor protein | 5.20 | 1.12E-31 |
| VC0605 | conserved hypothetical protein | 2.04 | 9.96E-06 |
| VC0635 | conserved hypothetical protein | 4.10 | 8.80E-27 |
| VC0636 | 23S rRNA methyltransferase | 2.13 | 6.95E-09 |
| VC0640 | preprotein translocase subunit SecG | 2.98 | 2.92E-14 |

| | | | |
|---|---|---|---|
| VC0641 | conserved hypothetical protein | 3.27 | 7.38E-40 |
| VC0642 | N utilization substance protein A | 2.67 | 1.88E-18 |
| VC0646 | ribosomal protein S15 | 2.92 | 2.46E-13 |
| VC0647 | polyribonucleotide nucleotidyltransferase | 2.39 | 6.55E-07 |
| VC0649 | transcriptional regulator, MarR family | 4.40 | 4.71E-32 |
| VC0650 | multidrug efflux pump VmrA | 3.17 | 3.97E-28 |
| VC0654 | conserved hypothetical protein | 3.58 | 7.11E-30 |
| VC0660 | ATP-dependent RNA helicase SrmB | 2.45 | 5.62E-15 |
| VC0668 | DNA mismatch repair protein MutH | 2.67 | 2.81E-12 |
| VC0679 | ribosomal protein S20 | 5.40 | 5.62E-23 |
| VC0681 | riboflavin kinase/FMN adenylyltransferase | 2.12 | 3.40E-13 |
| VC0693 | response regulator | 2.65 | 4.13E-13 |
| VC0694 | conserved hypothetical protein | 2.92 | 2.56E-16 |
| VC0695 | phospho-2-dehydro-3- deoxyheptonate aldolase,tyr-sensitive | 7.53 | 2.67E-70 |
| VC0709 | 23S rRNA pseudouridine synthase D | 2.97 | 7.11E-18 |
| VC0710 | conserved hypothetical protein | 2.57 | 4.19E-26 |
| VC0717 | protease, putative | 2.33 | 4.79E-14 |
| VC0718 | conserved hypothetical protein | 3.20 | 1.54E-22 |
| VC0734 | malate synthase A | 5.80 | 9.78E-45 |
| VC0736 | isocitrate lyase | 2.72 | 1.63E-16 |
| VC0738 | S-adenosylmethionine:tRNA ribosyltransferase-isomerase | 4.38 | 7.99E-38 |
| VC0739 | S-adenosylmethionine :tRNAribosyltransferase-isomer ase | 2.27 | 2.30E-20 |
| VC0745 | inositol monophosphate family protein | 4.04 | 1.79E-57 |
| VC0746 | RNA methyltransferase, TrmH family | 3.01 | 5.83E-19 |
| VC0750 | hesB family protein | 2.80 | 1.88E-16 |
| VC0757 | conserved hypothetical protein | 3.34 | 2.60E-27 |
| VC0767 | inosine-5`-monophosp hate dehydrogenase | 2.55 | 3.46E-15 |
| VC0791 | sensor kinase citA | 3.04 | 1.24E-15 |
| VC0804 | ferredoxin | 2.31 | 1.32E-06 |
| VC0806 | conserved hypothetical protein | 2.99 | 8.96E-18 |
| VC0807 | hypothetical protein | 9.61 | 2.26E-69 |
| VC0847 | integrase, phage family | 4.06 | 8.23E-48 |
| VC0854 | heat shock protein GrpE | 2.27 | 2.22E-10 |
| VC0864 | yfhC protein | 2.05 | 5.45E-07 |
| VC0869 | phosphoribosylformyl glycinamidine synthase | 2.81 | 1.86E-15 |
| VC0870 | IS1004 transposase | 6.73 | 8.00E-33 |
| VC0871 | hypothetical protein | 2.65 | 3.19E-11 |
| VC0872 | conserved hypothetical protein | 2.76 | 2.40E-13 |
| VC0873 | conserved hypothetical protein | 2.91 | 1.41E-13 |
| VC0875 | prolyl-tRNA synthetase | 2.59 | 1.88E-17 |
| VC0877 | hypothetical protein | 13.82 | 9.58E-92 |
| VC0879 | 50S ribosomal protein L36 | 6.56 | 1.14E-23 |
| VC0880 | conserved hypothetical protein | 2.30 | 1.97E-14 |
| VC0888 | pseudouridine synthase Rlu family protein | 2.29 | 1.45E-05 |
| VC0891 | exodeoxyribonuclease , small subunit | 2.17 | 6.48E-09 |
| VC0894 | thiamin biosynthesis protein ThiI | 4.60 | 2.05E-28 |
| VC0906 | ABC transporter, permease protein | 2.07 | 1.60E-13 |
| VC0907 | ABC transporter, ATP-binding protein | 3.44 | 1.19E-23 |
| VC0908 | histidinol phosphatase-related protein | 2.40 | 4.65E-21 |
| VC0916 | phosphotyrosine protein phosphatase | 57.66 | 3.04E-41 |
| VC0917 | UDP-N-acetylglucosam ine 2-epimerase | 2.92 | 2.20E-08 |
| VC0934 | capsular polysaccharide biosynthesisglycosyltransferas e, putative | 4.86 | 2.30E-10 |
| VC0945 | conserved hypothetical protein | 7.43 | 3.64E-74 |
| VC0952 | conserved hypothetical protein | 2.02 | 8.23E-13 |
| VC0962 | conserved hypothetical protein | 3.67 | 7.80E-23 |
| VC0963 | VisC-related protein | 2.52 | 2.58E-17 |
| VC0986 | adenylate kinase | 3.02 | 1.10E-14 |
| VC0987 | ferrochelatase | 2.21 | 5.56E-18 |
| VC1002 | dedD protein | 2.13 | 6.38E-09 |
| VC1003 | bacteriocin production protein | 2.97 | 2.01E-22 |
| VC1006 | ribonuclease T | 3.70 | 1.85E-29 |
| VC1017 | RnfA-related protein | 2.04 | 1.32E-07 |
| VC1040 | cob(I)alamin adenosyltransferase | 2.85 | 1.01E-14 |
| VC1046 | 3-ketoacyl-CoA thiolase | 6.40 | 7.46E-46 |
| VC1047 | multifunctional fatty acid oxidation complex subunit alpha | 5.85 | 1.06E-41 |
| VC1058 | conserved hypothetical protein | 2.20 | 3.88E-09 |
| VC1068 | transcriptional regulator, ArsR family | 2.13 | 3.80E-05 |
| VC1078 | hypothetical protein | 2.20 | 1.19E-06 |
| VC1079 | conserved hypothetical protein | 4.11 | 1.07E-29 |
| VC1105 | conserved hypothetical protein | 2.62 | 4.01E-13 |

| | | | |
|---|---|---|---|
| VC1107 | outer-membrane lipoprotein carrier protein | 2.10 | 1.32E-15 |
| VC1111 | adenosylmethionine-8 -amino-7-oxononanoateaminotran sferase | 3.07 | 9.17E-13 |
| VC1112 | biotin synthase | 4.26 | 9.86E-24 |
| VC1113 | 8-amino-7-oxononanoa te synthase | 2.65 | 6.74E-12 |
| VC1114 | biotin synthesis protein BioC | 3.49 | 1.16E-17 |
| VC1115 | dithiobiotin synthetase | 2.05 | 6.07E-16 |
| VC1116 | hypothetical protein | 7.00 | 2.03E-48 |
| VC1117 | heat shock protein HtpX | 2.45 | 5.48E-14 |
| VC1118 | transcriptional regulator, putative | 10.59 | 4.65E-81 |
| VC1119 | oxidoreductase, short-chaindehydrogenase/reduc tase family | 5.00 | 2.05E-37 |
| VC1120 | conserved hypothetical protein | 4.65 | 1.30E-41 |
| VC1121 | conserved hypothetical protein | 3.36 | 2.36E-29 |
| VC1122 | cyclopropane-fatty-a cyl-phospholipid synthase | 5.41 | 2.08E-42 |
| VC1123 | hypothetical protein | 4.88 | 3.80E-40 |
| VC1126 | adenylosuccinate lyase | 2.03 | 1.32E-06 |
| VC1128 | tRNA-specific 2-thiouridylase MnmA | 2.08 | 1.70E-16 |
| VC1132 | ATP phosphoribosyltransferase | 2.64 | 2.46E-15 |
| VC1146 | glutaredoxin 1 | 3.71 | 2.25E-31 |
| VC1152 | hypothetical protein | 2.50 | 1.82E-25 |
| VC1191 | hypothetical protein | 3.29 | 1.20E-24 |
| VC1193 | hypothetical protein | 6.29 | 3.47E-47 |
| VC1196 | hypothetical protein | 2.03 | 4.06E-08 |
| VC1208 | conserved hypothetical protein | 2.01 | 6.68E-09 |
| VC1209 | elongation factor P family protein | 2.12 | 1.66E-10 |
| VC1221 | hypothetical protein | 2.22 | 1.21E-08 |
| VC1228 | phosphoribosylglycinamide formyltransferase 2 | 2.34 | 2.32E-11 |
| VC1237 | nicotinate-nucleotide--dimethylbenzimidazole phosphoribosyltransfe | 3.84 | 1.24E-28 |
| VC1238 | adenosylcobinamide-GDP ribazoletransferase | 3.49 | 5.41E-19 |
| VC1239 | cobinamide kinase/cobinamide phosphateguanylyltransferase | 3.84 | 5.99E-19 |
| VC1240 | alpha-ribazole-5`-ph osphate phosphatase CobC,putative | 2.27 | 2.27E-05 |
| VC1246 | hypothetical protein | 2.28 | 7.73E-18 |
| VC1253 | hypothetical protein | 4.41 | 4.56E-34 |
| VC1257 | 3-demethylubiquinone -9 3-methyltransferase | 2.06 | 7.25E-15 |
| VC1258 | DNA gyrase, subunit A | 2.47 | 2.47E-07 |
| VC1259 | conserved hypothetical protein | 4.67 | 1.38E-19 |
| VC1262 | hypothetical protein | 3.09 | 5.84E-12 |
| VC1263 | GTP cyclohydrolase II | 6.90 | 3.29E-68 |
| VC1271 | hypothetical protein | 3.23 | 8.49E-17 |
| VC1278 | transcriptional regulator, MarR family | 3.97 | 1.96E-45 |
| VC1279 | transporter, BCCT family | 3.07 | 3.79E-18 |
| VC1296 | phosphomethylpyrimid ine kinase | 2.40 | 1.22E-11 |
| VC1299 | 6-pyruvoyl tetrahydrobiopterin synthase,putative | 2.72 | 3.51E-07 |
| VC1300 | L-serine dehydratase 1 | 2.01 | 5.37E-08 |
| VC1306 | conserved hypothetical protein | 2.72 | 3.33E-13 |
| VC1341 | acetyltransferase, putative | 3.78 | 2.66E-27 |
| VC1342 | MutT/nudix family protein | 2.15 | 6.83E-11 |
| VC1350 | antioxidant, putative | 5.52 | 1.64E-53 |
| VC1354 | conserved hypothetical protein | 4.32 | 2.28E-53 |
| VC1358 | conserved hypothetical protein | 3.95 | 8.44E-13 |
| VC1363 | siroheme synthase component enzyme | 2.06 | 2.43E-11 |
| VC1365 | conserved hypothetical protein | 2.94 | 4.69E-16 |
| VC1366 | exsB protein | 3.44 | 5.24E-31 |
| VC1368 | hypothetical protein | 2.68 | 1.36E-10 |
| VC1374 | DnaK-related protein | 2.01 | 5.06E-08 |
| VC1375 | hypothetical protein | 3.55 | 4.97E-14 |
| VC1382 | ATP-dependent helicase HrpA | 2.23 | 7.59E-17 |
| VC1386 | heat shock protein 70 family protein | 2.91 | 6.16E-14 |
| VC1387 | lipoate-protein ligase A | 2.27 | 1.93E-10 |
| VC1389 | hypothetical protein | 19.85 | 2.78E-09 |
| VC1390 | transcriptional regulator, LysR family | 3.44 | 9.15E-15 |
| VC1392 | deoxyribodipyrimidin e photolyase, putative | 6.04 | 3.70E-74 |
| VC1422a | membrane protein | 2.85 | 1.21E-16 |
| VC1423 | hypothetical protein | 2.83 | 1.09E-26 |
| VC1424 | spermidine/putrescin e ABC transporter,periplasmic spermidine/putr | 2.26 | 1.86E-12 |
| VC1428 | putrescine/spermidine ABC transporter ATPase protein | 2.42 | 9.81E-17 |
| VC1432 | conserved hypothetical protein | 2.21 | 1.46E-12 |
| VC1434 | fumarate and nitrate reduction regulatoryprotein | 2.47 | 3.63E-13 |
| VC1449 | hypothetical protein | 3.15 | 9.85E-14 |
| VC1452 | RstC protein | 2.26 | 4.38E-05 |
| VC1454 | RstA1 protein | 770.61 | 4.06E-04 |

| | | | |
|---|---|---|---|
| VC1455 | transcriptional repressor RstR | 3.73 | 1.68E-28 |
| VC1464 | transcriptional repressor RstR | 3.85 | 9.51E-27 |
| VC1465 | hypothetical protein | 2.01 | 5.75E-13 |
| VC1467 | hypothetical protein | 3.02 | 2.82E-30 |
| VC1473 | hypothetical protein | 3.01 | 3.64E-30 |
| VC1486 | ABC transporter, ATP-binding protein | 2.11 | 1.06E-09 |
| VC1487 | conserved hypothetical protein | 3.80 | 4.92E-16 |
| VC1488 | 23S rRNA m(2)G2445 methyltransferase | 2.33 | 1.30E-13 |
| VC1491 | dihydroorotate dehydrogenase | 2.50 | 2.87E-15 |
| VC1502 | 16S rRNA (cytosine(1407)-C(5))-methyltransferase RsmF | 2.82 | 1.62E-25 |
| VC1503 | conserved hypothetical protein | 4.10 | 2.40E-30 |
| VC1505 | hypothetical protein | 3.65 | 4.53E-32 |
| VC1506 | hypothetical protein | 4.84 | 3.48E-41 |
| VC1520 | ABC transporter, ATP-binding protein | 2.47 | 7.60E-13 |
| VC1530 | hypothetical protein | 6.45 | 5.02E-22 |
| VC1531 | hypothetical protein | 2.34 | 7.92E-20 |
| VC1541 | hypothetical protein | 3.20 | 3.12E-12 |
| VC1552 | glycerol-3-phosphate transporter ATP-binding subunit, ugpC | 2.19 | 3.87E-04 |
| VC1605 | sensor kinase citA, putative | 2.26 | 2.04E-18 |
| VC1605a | hypothetical protein | 3.22 | 5.20E-15 |
| VC1610 | hypothetical protein | 3.61 | 1.17E-19 |
| VC1616 | glutaredoxin, putative | 2.73 | 1.13E-12 |
| VC1630 | ABC transporter, ATP-binding protein | 2.14 | 4.35E-16 |
| VC1635 | ribosomal small subunit pseudouridine synthaseA | 4.13 | 1.30E-31 |
| VC1637 | hypothetical protein | 8.21 | 1.62E-55 |
| VC1638 | DNA-binding response regulator | 4.48 | 2.29E-26 |
| VC1639 | sensor histidine kinase | 2.77 | 1.48E-16 |
| VC1645 | conserved hypothetical protein | 3.72 | 4.45E-29 |
| VC1668 | pseudouridine synthase Rlu family protein | 2.34 | 5.31E-12 |
| VC1686 | hypothetical protein | 3.02 | 1.44E-31 |
| VC1687 | conserved hypothetical protein | 2.30 | 6.95E-09 |
| VC1701 | conserved hypothetical protein | 2.81 | 1.19E-09 |
| VC1704 | 5-methyltetrahydropt eroyltriglutamate--homocystein e methyltransf | 13.44 | 8.15E-159 |
| VC1708 | conserved hypothetical protein | 2.64 | 1.11E-13 |
| VC1711 | conserved hypothetical protein | 3.84 | 6.67E-25 |
| VC1719 | DNA-binding response regulator TorR | 8.60 | 1.60E-74 |
| VC1720 | chaperone protein TorD | 2.03 | 2.15E-05 |
| VC1722 | conserved hypothetical protein | 3.26 | 9.28E-28 |
| VC1725 | beta-ketoadipate enol-lactone hydrolase,putative | 2.53 | 1.72E-24 |
| VC1731 | conserved hypothetical protein | 4.03 | 8.27E-26 |
| VC1737 | translation initiation factor IF-1 | 4.10 | 1.61E-27 |
| VC1740 | oxidoreductase, acyl-CoA dehydrogenase family | 7.88 | 3.05E-50 |
| VC1741 | transcriptional regulator, TetR family | 8.10 | 1.40E-109 |
| VC1785 | transcriptional regulator | 4.03 | 2.02E-04 |
| VC1801 | hypothetical protein | 4.00 | 1.14E-08 |
| VC1803 | hypothetical protein | 2.64 | 9.26E-15 |
| VC1804 | hypothetical protein | 2.02 | 3.77E-05 |
| VC1810 | hypothetical protein | 2.83 | 1.03E-19 |
| VC1811 | conserved hypothetical protein | 3.13 | 3.02E-23 |
| VC1812 | conserved hypothetical protein | 4.41 | 1.00E-12 |
| VC1814 | deoxyribodipyrimidin e photolyase | 15.11 | 4.31E-62 |
| VC1815 | C-factor, putative | 10.31 | 1.64E-47 |
| VC1816 | hypothetical protein | 3.70 | 2.43E-12 |
| VC1823 | PTS system, fructose-specific IIB component | 3.03 | 1.67E-11 |
| VC1832 | hypothetical protein | 2.13 | 1.30E-17 |
| VC1833 | quinolinate synthetase A | 2.23 | 2.59E-19 |
| VC1835 | peptidoglycan-associ ated lipoprotein | 2.27 | 1.53E-07 |
| VC1837 | tolA protein | 2.19 | 6.11E-13 |
| VC1838 | tolR membrane protein | 2.29 | 5.77E-13 |
| VC1839 | tolQ protein | 3.17 | 5.40E-22 |
| VC1840 | conserved hypothetical protein | 8.64 | 1.10E-115 |
| VC1845 | Holliday junction DNA helicase RuvB | 2.42 | 2.31E-18 |
| VC1846 | Holliday junction DNA helicase RuvA | 2.67 | 1.19E-16 |
| VC1849 | peptidyl-prolyl cis-trans isomerase B | 3.30 | 4.03E-22 |
| VC1860 | exodeoxyribonuclease III | 2.71 | 4.14E-15 |
| VC1863 | amino acid ABC transporter, periplasmic aminoacid-binding protein | 2.02 | 3.93E-09 |
| VC1864 | amino acid ABC transporter, ATP-binding protein | 2.40 | 3.55E-12 |
| VC1876 | conserved hypothetical protein | 2.39 | 1.99E-09 |
| VC1883 | ABC transporter, ATP-binding protein | 2.20 | 4.56E-10 |

| | | | |
|---|---|---|---|
| VC1890 | NADH dehydrogenase | 2.77 | 8.51E-17 |
| VC1906 | hypothetical protein | 2.08 | 5.15E-08 |
| VC1913 | conserved hypothetical protein | 3.54 | 6.93E-26 |
| VC1915 | ribosomal protein S1 | 2.11 | 1.36E-03 |
| VC1916 | cytidylate kinase | 2.43 | 9.70E-12 |
| VC1924 | hypothetical protein | 2.29 | 2.70E-09 |
| VC1927 | C4-dicarboxylate transport protein | 5.19 | 1.09E-34 |
| VC1929 | C4-dicarboxylate-bin ding periplasmic protein | 7.73 | 2.56E-19 |
| VC1931 | conserved hypothetical protein | 2.76 | 3.76E-24 |
| VC1938 | conserved hypothetical protein | 2.38 | 4.70E-20 |
| VC1942 | methylenetetrahydrof olatedehydrogenase/methenyltet rahydrofolate | 2.03 | 3.22E-12 |
| VC1961 | cell division topological specificity factor MinE | 2.67 | 2.27E-14 |
| VC1963 | conserved hypothetical protein | 2.25 | 7.15E-11 |
| VC1964 | hypothetical protein | 4.83 | 3.03E-20 |
| VC1980 | conserved hypothetical protein | 2.36 | 2.41E-09 |
| VC1983 | peptidase, putative | 2.74 | 7.03E-20 |
| VC1993 | 2,4-dienoyl-CoA reductase | 2.89 | 1.01E-16 |
| VC2020 | acyl carrier protein | 2.42 | 4.75E-12 |
| VC2025 | ribosomal protein L32 | 2.54 | 3.07E-12 |
| VC2026 | conserved hypothetical protein | 2.66 | 6.28E-06 |
| VC2027 | Maf-like protein | 2.79 | 7.64E-27 |
| VC2028 | ribosomal large subunit pseudouridine synthaseC | 2.38 | 1.44E-20 |
| VC2031 | sulfate permease family protein | 2.31 | 1.03E-11 |
| VC2035 | conserved hypothetical protein | 2.25 | 1.01E-16 |
| VC2042 | histone deacetylase/AcuC/AphA family protein | 3.14 | 1.02E-16 |
| VC2044 | conserved hypothetical protein | 7.01 | 1.75E-51 |
| VC2057 | heme exporter protein A | 2.40 | 1.89E-15 |
| VC2073 | conserved hypothetical protein | 2.34 | 9.71E-08 |
| VC2078 | ferrous iron transport protein A | 4.21 | 2.12E-13 |
| VC2081 | zinc ABC transporter, periplasmic zinc-bindingprotein | 2.33 | 8.94E-14 |
| VC2082 | zinc ABC transporter, ATP-binding protein | 4.70 | 3.79E-32 |
| VC2098 | hypothetical protein | 2.02 | 4.47E-14 |
| VC2115 | conserved hypothetical protein | 2.05 | 3.24E-09 |
| VC2134 | flagellar hook-basal body protein FliE | 2.38 | 3.44E-11 |
| VC2146 | conserved hypothetical protein | 2.55 | 7.11E-23 |
| VC2160 | thioredoxin-dependent thiol peroxidase | 2.55 | 4.78E-18 |
| VC2163 | conserved hypothetical protein | 2.64 | 1.53E-14 |
| VC2180 | glutamyl-tRNA reductase | 2.76 | 3.43E-21 |
| VC2184 | peptidyl-tRNA hydrolase | 2.92 | 1.22E-24 |
| VC2185 | GTP-binding protein | 2.17 | 1.14E-10 |
| VC2213 | outer membrane protein OmpA | 2.45 | 1.08E-04 |
| VC2214 | glutamyl-tRNA synthetase | 2.39 | 3.40E-13 |
| VC2215 | cation transport ATPase, E1-E2 family | 2.01 | 3.72E-12 |
| VC2223 | pseudouridine synthase family 1 protein | 2.47 | 9.03E-14 |
| VC2226 | phosphoribosylformyl glycinamidine cyclo-ligase | 2.79 | 4.66E-15 |
| VC2227 | phosphoribosylglycinamide formyltransferase | 2.90 | 6.91E-22 |
| VC2239 | nitrogen regulatory protein P-II | 2.85 | 4.65E-15 |
| VC2259 | elongation factor Ts | 2.89 | 1.26E-08 |
| VC2268 | 6,7-dimethyl-8-ribityllumazine synthase | 2.20 | 1.69E-10 |
| VC2272 | transcriptional regulator NrdR | 2.73 | 1.99E-17 |
| VC2296 | bolA protein | 5.38 | 1.11E-81 |
| VC2301 | transcriptional activator, putative | 52.55 | 3.29E-87 |
| VC2302 | RNA polymerase sigma-70 factor, ECF subfamily | 5.01 | 1.58E-46 |
| VC2303 | conserved hypothetical protein | 17.42 | 2.97E-107 |
| VC2305 | outer membrane protein OmpK | 4.26 | 3.18E-40 |
| VC2311 | HesA/MoeB/ThiF family protein | 2.30 | 6.30E-11 |
| VC2318 | hypothetical protein | 16.03 | 6.47E-111 |
| VC2326 | conserved hypothetical protein | 5.14 | 5.82E-35 |
| VC2342 | elongation factor G | 2.14 | 2.38E-04 |
| VC2346 | smp protein, putative | 2.39 | 8.75E-20 |
| VC2361 | formate acetyl transferase-related protein | 2.30 | 1.43E-04 |
| VC2388 | hypothetical protein | 2.46 | 4.29E-08 |
| VC2389 | carbamoyl-phosphate synthase, large subunit | 9.74 | 4.42E-65 |
| VC2390 | carbamoyl-phosphate synthase, small subunit | 6.18 | 2.83E-47 |
| VC2444 | general secretion pathway protein B, putative | 2.02 | 4.41E-07 |
| VC2478 | conserved hypothetical protein | 2.89 | 7.72E-14 |
| VC2479 | conserved hypothetical protein | 2.08 | 1.56E-08 |
| VC2480 | ribose-5-phosphate isomerase | 2.63 | 1.18E-16 |
| VC2484 | long-chain-fatty-aci d--CoA ligase, putative | 5.57 | 6.68E-61 |
| VC2485 | transcriptional regulator, LysR family | 3.45 | 1.60E-26 |

| | | | |
|---|---|---|---|
| VC2500 | conserved hypothetical protein | 3.23 | 4.03E-39 |
| VC2509 | aspartate carbamoyltransferase | 3.68 | 1.29E-30 |
| VC2511 | aspartate carbamoyltransferase, regulatorysubunit | 5.44 | 1.57E-44 |
| VC2512 | conserved hypothetical protein | 2.51 | 3.74E-14 |
| VC2540 | hypothetical protein | 4.03 | 5.34E-20 |
| VC2541 | 3-octaprenyl-4-hydro xybenzoate carboxy-lyase | 3.21 | 1.25E-30 |
| VC2542 | UDP-N-acetylmuramate :L-alanyl-gamma-D-glutamyl-mes o-diamin | 3.43 | 1.44E-33 |
| VC2551 | conserved hypothetical protein | 2.16 | 2.41E-10 |
| VC2554 | conserved hypothetical protein | 2.06 | 2.45E-08 |
| VC2555 | hypothetical protein | 3.08 | 9.43E-17 |
| VC2562 | bifunctional 2',3'-cyclic nucleotide 2'-phosphodiesterase/3'-nucleotid | 2.04 | 7.76E-08 |
| VC2563 | hypothetical protein | 2.50 | 1.66E-13 |
| VC2564 | ATP-dependent RNA helicase DbpA | 2.41 | 3.00E-10 |
| VC2568 | peptidyl-prolyl cis-trans isomerase, FKBP-type | 2.88 | 1.24E-14 |
| VC2569 | hypothetical protein | 3.51 | 1.76E-21 |
| VC2582 | ribosomal protein S8 | 2.10 | 3.85E-07 |
| VC2583 | ribosomal protein S14 | 2.44 | 1.40E-11 |
| VC2584 | ribosomal protein L5 | 2.89 | 1.76E-07 |
| VC2585 | ribosomal protein L24 | 5.26 | 8.46E-15 |
| VC2586 | ribosomal protein L14 | 4.43 | 5.38E-13 |
| VC2594 | ribosomal protein L23 | 2.31 | 1.68E-10 |
| VC2597 | ribosomal protein S10 | 2.29 | 6.74E-10 |
| VC2600 | conserved hypothetical protein | 3.51 | 1.28E-30 |
| VC2617 | arginine/ornithine succinyltransferase,putative | 2.05 | 5.66E-09 |
| VC2618 | acetylornithine aminotransferase | 2.37 | 1.57E-10 |
| VC2624 | phosphoglycolate phosphatase | 2.04 | 1.42E-08 |
| VC2625 | ribulose-phosphate 3-epimerase | 2.04 | 7.58E-15 |
| VC2629 | shikimate kinase I | 2.61 | 1.69E-22 |
| VC2642 | argininosuccinate synthase | 4.59 | 1.94E-38 |
| VC2643 | acetylglutamate kinase | 7.65 | 2.81E-58 |
| VC2644 | N-acetyl-gamma-glutamyl-phosphate reductase | 13.23 | 4.12E-97 |
| VC2647 | conserved hypothetical protein | 3.07 | 1.84E-22 |
| VC2660 | elongation factor P | 3.23 | 2.14E-24 |
| VC2662 | conserved hypothetical protein | 4.61 | 3.54E-28 |
| VC2670 | triosephosphate isomerase | 2.90 | 1.21E-13 |
| VC2675 | protease HslVU, subunit HslV | 2.92 | 2.92E-24 |
| VC2679 | ribosomal protein L31 | 4.32 | 7.95E-28 |
| VC2682 | met repressor | 3.73 | 8.95E-39 |
| VC2685 | 5,10-methylenetetrahydrofolate reductase | 3.74 | 1.79E-28 |
| VC2687 | glutamyl-tRNA synthetase | 2.70 | 9.20E-16 |
| VC2712 | xanthine/uracil permease family protein | 2.30 | 4.22E-10 |
| VC2715 | transcription elongation factor GreB | 2.05 | 6.74E-06 |
| VC2717 | hypothetical protein | 2.24 | 3.87E-09 |
| VC2718 | bioH protein | 2.83 | 4.23E-13 |
| VC2735 | conserved hypothetical protein | 2.09 | 1.67E-08 |
| VC2744 | GTP-binding protein TypA | 2.07 | 4.77E-11 |
| VC2753 | hypothetical protein | 6.07 | 3.08E-30 |
| VC2757 | conserved hypothetical protein | 3.81 | 4.37E-34 |
| VC2758 | multifunctional fatty acid oxidation complex subunit alpha | 6.02 | 1.48E-29 |
| VC2759 | 3-ketoacyl-CoA thiolase | 5.41 | 8.40E-31 |
| VC2774 | glucose inhibited division protein B | 2.85 | 1.12E-20 |
| VC2775 | glucose inhibited division protein A | 3.11 | 5.61E-39 |
| VCA0005 | hypothetical protein | 2.58 | 2.86E-22 |
| VCA0010 | conserved hypothetical protein | 13.25 | 5.69E-90 |
| VCA0035 | phosphatidylglycerop hosphatase B, putative | 2.92 | 2.74E-18 |
| VCA0040 | conserved hypothetical protein | 2.46 | 5.18E-22 |
| VCA0076 | conserved hypothetical protein | 7.58 | 2.41E-80 |
| VCA0086 | hypothetical protein | 7.44 | 6.08E-80 |
| VCA0087 | hypothetical protein | 8.94 | 5.82E-33 |
| VCA0098 | nicotinate phosphoribosyltransferase | 3.66 | 1.42E-40 |
| VCA0125 | hypothetical protein | 6.81 | 1.70E-94 |
| VCA0133 | pseudogene | 2.48 | 1.32E-11 |
| VCA0139 | hypothetical protein | 6.82 | 2.28E-47 |
| VCA0143 | hypothetical protein | 3.04 | 2.40E-04 |
| VCA0149 | hypothetical protein | 4.18 | 1.98E-21 |
| VCA0158 | hypothetical protein | 3.96 | 9.80E-27 |
| VCA0159 | conserved hypothetical protein | 4.38 | 5.48E-64 |
| VCA0166 | cold shock transcriptional regulator CspA | 3.76 | 1.48E-10 |
| VCA0177 | hypothetical protein | 2.74 | 4.45E-16 |
| VCA0179 | NupC family protein | 2.36 | 5.07E-06 |

| | | | |
|---|---|---|---|
| VCA0184 | cold shock DNA-binding domain protein | 2.67 | 2.31E-11 |
| VCA0185 | conserved hypothetical protein | 3.55 | 9.71E-14 |
| VCA0197 | GMP reductase | 2.66 | 9.55E-13 |
| VCA0198 | site-specific DNA-methyltransferase, putative | 2.93 | 3.70E-26 |
| VCA0204 | ATP-dependent RNA helicase RhlE | 2.89 | 1.99E-19 |
| VCA0215 | multidrug resistance protein D | 8.05 | 2.03E-109 |
| VCA0240 | hypothetical protein | 2.02 | 1.58E-05 |
| VCA0246 | PTS system ascorbate-specific transporter subunits IICB | 2.08 | 7.69E-08 |
| VCA0269 | decarboxylase, group II | 4.59 | 5.30E-39 |
| VCA0279 | transcriptional regulator, HTH_3 family | 8.66 | 5.50E-51 |
| VCA0287 | threonyl-tRNA synthetase | 2.37 | 4.60E-07 |
| VCA0288 | initiation factor IF3 | 3.43 | 1.36E-08 |
| VCA0289 | ribosomal protein L35 | 4.09 | 1.52E-16 |
| VCA0290 | ribosomal protein L20 | 3.74 | 7.27E-15 |
| VCA0332 | conserved hypothetical protein | 2.99 | 4.68E-18 |
| VCA0333 | hypothetical protein | 2.83 | 3.28E-28 |
| VCA0349 | relB protein | 2.03 | 2.57E-07 |
| VCA0369 | hypothetical protein | 5.85 | 4.21E-04 |
| VCA0385 | conserved hypothetical protein | 2.21 | 2.75E-09 |
| VCA0391 | killer protein, putative | 3.03 | 8.92E-26 |
| VCA0392 | antidote protein, putative | 2.44 | 2.64E-14 |
| VCA0422 | conserved hypothetical protein | 3.30 | 4.39E-33 |
| VCA0423 | conserved hypothetical protein | 3.65 | 4.39E-40 |
| VCA0426 | conserved hypothetical protein | 2.40 | 2.65E-10 |
| VCA0444 | relE protein | 3.04 | 1.94E-19 |
| VCA0445 | conserved hypothetical protein | 3.19 | 1.13E-16 |
| VCA0468 | hypothetical protein | 3.09 | 3.72E-24 |
| VCA0481 | conserved hypothetical protein | 2.10 | 1.69E-09 |
| VCA0486 | hypothetical protein | 3.75 | 1.76E-32 |
| VCA0487 | conserved hypothetical protein | 2.39 | 1.40E-22 |
| VCA0488 | conserved hypothetical protein | 2.68 | 5.78E-18 |
| VCA0490 | lipase, GDXG family | 3.59 | 1.96E-22 |
| VCA0491 | hypothetical protein | 3.63 | 4.06E-25 |
| VCA0492 | pseudogene | 3.06 | 2.63E-29 |
| VCA0493 | IS1004 transposase | 3.09 | 4.51E-14 |
| VCA0494 | hypothetical protein, interruption | 3.96 | 5.99E-11 |
| VCA0497 | hypothetical protein | 2.35 | 2.47E-15 |
| VCA0503 | conserved hypothetical protein | 2.35 | 4.24E-11 |
| VCA0504 | relB protein | 2.30 | 3.12E-09 |
| VCA0505 | acetyltransferase, putative | 2.22 | 1.71E-10 |
| VCA0506 | conserved hypothetical protein | 2.14 | 1.26E-08 |
| VCA0518 | PTS system, fructose-specific IIA/FPR component | 4.65 | 1.78E-31 |
| VCA0520 | exonuclease SbcD, putative | 3.19 | 3.63E-31 |
| VCA0531 | sensor histidine kinase | 2.89 | 1.99E-24 |
| VCA0532 | DNA-binding response regulator | 2.24 | 7.24E-09 |
| VCA0535 | hypothetical protein | 5.38 | 5.93E-71 |
| VCA0546 | conserved hypothetical protein | 3.56 | 4.82E-30 |
| VCA0547 | hypothetical protein | 3.17 | 8.12E-17 |
| VCA0549 | phnA protein | 3.04 | 2.09E-35 |
| VCA0575 | transcriptional regulator, LysR family | 2.05 | 2.72E-14 |
| VCA0580 | conserved hypothetical protein | 2.15 | 3.21E-16 |
| VCA0582 | conserved hypothetical protein | 3.21 | 7.18E-20 |
| VCA0586 | conserved hypothetical protein | 3.56 | 1.83E-10 |
| VCA0587 | conserved hypothetical protein | 2.76 | 1.89E-09 |
| VCA0596 | pseudogene | 2.73 | 1.30E-12 |
| VCA0608 | conserved hypothetical protein | 2.78 | 1.62E-27 |
| VCA0614 | formate--tetrahydrof olate ligase | 3.72 | 8.45E-43 |
| VCA0616 | GTP cyclohydrolase I | 2.07 | 4.16E-08 |
| VCA0627 | rRNA methylase, putative | 2.42 | 1.26E-07 |
| VCA0634 | conserved hypothetical protein | 2.54 | 2.24E-11 |
| VCA0635 | transcriptional regulator, LysR family | 2.20 | 3.41E-06 |
| VCA0652 | hypothetical protein | 3.81 | 7.06E-22 |
| VCA0693 | preprotein translocase subunit SecD | 3.80 | 1.48E-21 |
| VCA0694 | hypothetical protein | 3.48 | 3.09E-29 |
| VCA0707 | regulatory protein UhpC, putative | 3.74 | 6.71E-19 |
| VCA0732 | conserved hypothetical protein | 10.18 | 3.60E-41 |
| VCA0733 | hypothetical protein | 16.28 | 2.11E-48 |
| VCA0741 | conserved hypothetical protein | 2.60 | 7.59E-14 |
| VCA0742 | hypothetical protein | 2.27 | 6.48E-07 |
| VCA0753 | conserved hypothetical protein | 2.70 | 6.19E-10 |

| | | | |
|---|---|---|---|
| VCA0754 | lipase-related protein | 2.28 | 1.33E-06 |
| VCA0758 | arginine ABC transporter, permease protein | 2.51 | 8.07E-11 |
| VCA0759 | arginine ABC transporter, periplasmicarginine-binding protein | 4.78 | 8.04E-32 |
| VCA0760 | arginine ABC transporter, ATP-binding protein | 3.07 | 1.93E-21 |
| VCA0768 | ATP-dependent RNA helicase, DEAD box family | 3.13 | 3.00E-34 |
| VCA0770 | hypothetical protein | 2.94 | 4.75E-13 |
| VCA0779 | hypothetical protein | 2.49 | 9.52E-15 |
| VCA0793 | hypothetical protein | 2.96 | 1.83E-11 |
| VCA0794 | hypothetical protein | 4.05 | 1.62E-11 |
| VCA0795 | resolvase, putative | 2.54 | 1.09E-17 |
| VCA0804 | ATP-dependent RNA helicase DeaD | 3.84 | 2.19E-22 |
| VCA0805 | exoribonuclease II | 2.00 | 4.63E-15 |
| VCA0809 | conserved hypothetical protein | 2.88 | 4.30E-19 |
| VCA0821 | hypothetical protein | 2.29 | 4.27E-09 |
| VCA0831 | hypothetical protein | 2.31 | 4.65E-04 |
| VCA0834 | hypothetical protein | 2.01 | 9.21E-10 |
| VCA0837 | hemolysin, putative | 2.13 | 4.68E-10 |
| VCA0838 | conserved hypothetical protein | 6.12 | 9.28E-63 |
| VCA0843 | glyceraldehyde 3-phosphate dehydrogenase | 2.19 | 6.33E-14 |
| VCA0845 | hypothetical protein | 6.81 | 8.45E-49 |
| VCA0846 | conserved hypothetical protein | 4.22 | 7.56E-30 |
| VCA0847 | conserved hypothetical protein | 6.06 | 3.93E-34 |
| VCA0862 | long-chain fatty acid transport protein | 2.37 | 1.29E-10 |
| VCA0863 | lipase, putative | 2.78 | 2.61E-15 |
| VCA0890 | glyoxylase I family protein | 2.26 | 2.83E-07 |
| VCA0893 | hypothetical protein | 4.21 | 1.68E-34 |
| VCA0898 | 6-phosphogluconate dehydrogenase,decarboxylating | 2.19 | 2.41E-10 |
| VCA0899 | hypothetical protein | 9.48 | 5.23E-23 |
| VCA0902 | hypothetical protein | 3.27 | 5.86E-25 |
| VCA0925 | dihydroorotase | 3.17 | 5.99E-24 |
| VCA0926 | transcriptional regulator, AraC/XylS family | 2.68 | 2.14E-15 |
| VCA0927 | conserved hypothetical protein | 2.42 | 4.55E-08 |
| VCA0933 | cold shock domain family protein | 7.50 | 5.01E-44 |
| VCA0938 | pseudogene | 3.50 | 4.82E-24 |
| VCA0953 | peptidyl-prolyl cis-trans isomerase C | 2.19 | 1.59E-09 |
| VCA0989 | conserved hypothetical protein | 2.89 | 5.46E-28 |
| VCA1004 | conserved hypothetical protein | 2.58 | 7.62E-13 |
| VCA1005 | transcriptional regulator, MarR family | 2.45 | 9.77E-08 |
| VCA1012 | conserved hypothetical protein | 2.06 | 6.67E-05 |
| VCA1013 | conserved hypothetical protein | 2.20 | 4.29E-09 |
| VCA1035 | hypothetical protein | 2.01 | 1.85E-07 |
| VCA1044 | hypothetical protein | 3.68 | 2.52E-32 |
| VCA1051 | conserved hypothetical protein | 2.00 | 5.15E-07 |
| VCA1058 | transcriptional regulator, LysR family | 2.26 | 7.28E-07 |
| VCA1073 | bifunctional proline dehydrogenase/pyrroline-5-carboxylate dehydro | 2.30 | 1.88E-09 |
| VCA1078 | transcriptional regulator, LuxR family | 2.02 | 1.08E-13 |
| VCA1111 | thermostable hemolysin | 2.96 | 2.81E-06 |

| 126 differentially-expressed genes common to both Δ*ompU* and Δ*rpoE* compared to the WT | | | | | |
|---|---|---|---|---|---|
| **Name** | **Function** | **FC** | **FDR *p* -value** | **FC Δ*rpoE*** | **FDR *p* -value** |
| 16Sh | 16Sh | -2.33 | 7.18E-07 | -14.86 | 4.22E-24 |
| 5Sb | 5Sb | 4.89 | 4.55E-07 | -134.06 | 1.60E-42 |
| 5Sc | 5Sc | 15.97 | 9.85E-05 | -6749.82 | 5.39E-96 |
| 5Sd | 5Sd | 8.98 | 3.91E-04 | -18963.94 | 9.00E-47 |
| 5Sf | 5Sf | 7.07 | 6.64E-05 | -1328.85 | 2.23E-165 |
| 5Sh | 5Sh | 3.07 | 8.58E-04 | -104.01 | 5.06E-35 |
| tRNA-Arg- | tRNA-Arg-1 | -2.37 | 5.68E-04 | 3.43 | 2.00E-15 |
| tRNA-Gly- | tRNA-Gly-8 | -2.14 | 1.70E-04 | 5.58 | 3.56E-21 |
| tRNA-Met- | tRNA-Met-8 | -2.88 | 8.91E-06 | 2.34 | 1.75E-07 |
| VC0017 | hypothetical protein | 2.66 | 3.25E-07 | 55.57 | 1.21E-126 |
| VC0062 | thiamin-phosphate pyrophosphorylase | 2.85 | 3.43E-04 | 3.28 | 1.12E-12 |
| VC0063 | thiF protein | 3.13 | 4.31E-06 | 2.17 | 5.22E-07 |
| VC0091 | O-methyltransferase- related protein | 3.70 | 8.88E-09 | 2.50 | 7.91E-13 |
| VC0107 | hypothetical protein | 8.41 | 8.16E-19 | 19.52 | 3.08E-55 |
| VC0157 | alkaline serine protease | 4.50 | 2.07E-14 | -3.76 | 1.14E-49 |
| VC0181 | conserved hypothetical protein | -2.19 | 3.52E-04 | -2.94 | 4.51E-25 |
| VC0364 | bacterioferritin-ass ociated ferredoxin | 3.25 | 2.53E-07 | 4.95 | 4.53E-45 |
| VC0379a | phage shock protein G | -2.16 | 1.54E-06 | 3.31 | 1.34E-34 |
| VC0475 | iron-regulated outer membrane virulence protein,TonB receptor fam | 8.15 | 2.02E-40 | 6.82 | 1.95E-52 |
| VC0494 | conserved hypothetical protein | -2.94 | 8.03E-06 | 3.86 | 6.19E-11 |

| | | | | | |
|---|---|---|---|---|---|
| VC0503 | conserved hypothetical protein | -3.27 | 9.60E-14 | 2.05 | 4.62E-09 |
| VC0566 | protease DO | -2.31 | 2.88E-07 | -9.28 | 3.02E-78 |
| VC0608 | iron(III) ABC transporter, periplasmiciron-compound-bindi ng protein | 4.76 | 1.21E-14 | 6.66 | 5.37E-54 |
| VC0676 | nptA protein | 4.54 | 1.54E-17 | 6.60 | 1.36E-36 |
| VC0747 | conserved hypothetical protein | 2.24 | 1.34E-05 | 2.70 | 7.67E-29 |
| VC0748 | aminotransferase NifS, class V | 2.13 | 4.83E-04 | 2.64 | 5.88E-18 |
| VC0749 | NifU-related protein | 2.28 | 3.31E-05 | 2.50 | 1.73E-21 |
| VC0754 | conserved hypothetical protein | 2.20 | 4.81E-05 | 2.24 | 4.17E-06 |
| VC0771 | vibriobactin-specifi c isochorismatase | 8.03 | 3.45E-15 | 2.80 | 5.69E-07 |
| VC0774 | vibriobactin-specifi c2,3-dihydro-2,3-dihydroxybenz oate dehydroge | 5.82 | 1.08E-16 | 2.45 | 2.07E-03 |
| VC0776 | ferric vibriobactin ABC transporter, periplasmicferric vibriobactin-bin | 4.16 | 3.49E-13 | 2.90 | 2.62E-08 |
| VC0818 | pseudogene; within VPI-I; unknown function | -2.18 | 4.36E-04 | 2.17 | 1.20E-06 |
| VC0825 | toxin co-regulated pilus biosynthesis protein I | -2.13 | 3.69E-04 | 2.48 | 1.93E-09 |
| VC0827 | toxin co-regulated pilus biosynthesis protein H | 2.53 | 9.33E-08 | 6.22 | 2.09E-86 |
| VC0828 | toxin co-regulated pilin | 2.54 | 2.88E-06 | 4.21 | 4.00E-34 |
| VC0834 | toxin co-regulated pilus biosynthesis protein S | 2.92 | 4.33E-05 | -4.78 | 2.26E-12 |
| VC0835 | toxin co-regulated pilus biosynthesis protein T | 2.78 | 3.37E-08 | -4.75 | 4.99E-11 |
| VC0838 | TCP pilus virulence regulatory protein | 4.64 | 8.40E-07 | -4.74 | 3.04E-08 |
| VC0885 | hypothetical protein | 3.04 | 2.60E-13 | -6.13 | 1.36E-44 |
| VC0928 | hypothetical protein | 2.20 | 1.60E-04 | 7.33 | 7.48E-43 |
| VC0930 | hemolysin-related protein | 2.66 | 6.13E-04 | 2.38 | 5.54E-12 |
| VC0936 | polysaccharide export-related protein | 2.37 | 9.24E-04 | -3.16 | 1.76E-13 |
| VC0938 | hypothetical protein | 2.52 | 2.37E-05 | -2.12 | 1.49E-06 |
| VC0972 | porin, putative | 2.25 | 1.65E-05 | -2.36 | 1.43E-10 |
| VC1029 | GGDEF family protein | 2.34 | 4.79E-04 | -2.19 | 5.33E-08 |
| VC1051 | hypothetical protein | 2.08 | 9.34E-04 | 2.18 | 1.92E-06 |
| VC1061 | cysteine synthase/cystathionine beta-synthasefamily protein | 3.39 | 1.33E-10 | -4.75 | 1.78E-26 |
| VC1204 | formiminoglutamase | 2.82 | 3.80E-06 | -5.80 | 3.41E-24 |
| VC1318 | outer membrane protein OmpV | 7.89 | 6.00E-24 | -43.63 | 1.39E-99 |
| VC1329 | opacity protein-related protein | 13.58 | 1.78E-40 | -2.62 | 1.55E-07 |
| VC1344 | 4-hydroxyphenylpyruv ate dioxygenase | 3.94 | 2.76E-14 | 3.54 | 5.79E-40 |
| VC1345 | oxidoreductase, putative | 5.22 | 1.54E-23 | 2.83 | 1.78E-19 |
| VC1346 | conserved hypothetical protein | 4.94 | 8.80E-24 | 2.21 | 1.57E-13 |
| VC1422 | sodium/alanine symporter | -2.20 | 8.66E-07 | 3.80 | 1.79E-38 |
| VC1456 | cholera enterotoxin, B subunit | 6.16 | 1.83E-11 | -68.23 | 3.51E-85 |
| VC1457 | cholera enterotoxin, A subunit | 3.61 | 4.52E-09 | -79.05 | 1.69E-110 |
| VC1472 | hypothetical protein | -4.32 | 6.83E-06 | 2.29 | 9.71E-04 |
| VC1548 | hypothetical protein | 3.62 | 1.11E-07 | 2.32 | 4.93E-08 |
| VC1570 | quinol oxidase, subunit II | 3.93 | 3.72E-09 | 6.38 | 1.77E-24 |
| VC1571 | quinol oxidase, subunit I | 3.34 | 1.53E-09 | 8.16 | 3.85E-42 |
| VC1572 | hypothetical protein | 2.74 | 5.92E-06 | 2.60 | 6.65E-05 |
| VC1577 | hypothetical protein | 4.99 | 1.11E-09 | -8.10 | 9.05E-43 |
| VC1578 | hypothetical protein | 5.82 | 7.21E-12 | -4.58 | 3.17E-16 |
| VC1579 | enterobactin synthetase component F-relatedprotein | 5.77 | 4.43E-17 | -8.22 | 7.70E-98 |
| VC1633 | hypothetical protein | 2.63 | 7.14E-07 | 2.36 | 5.54E-20 |
| VC1644 | hypothetical protein | 2.27 | 8.49E-07 | 3.41 | 2.66E-42 |
| VC1675 | multidrug resistance protein, putative | -2.04 | 6.66E-05 | -2.35 | 7.60E-20 |
| VC1678 | phage shock protein A | -2.58 | 1.16E-07 | 3.67 | 1.55E-27 |
| VC1786 | DNA repair protein RadC, putative | -3.45 | 7.95E-05 | 2.19 | 4.11E-03 |
| VC1802 | hypothetical protein | -3.45 | 2.37E-04 | 19.79 | 6.26E-18 |
| VC1807 | pseudogene | 3.04 | 1.31E-04 | 5.57 | 3.57E-05 |
| VC1819 | aldehyde dehydrogenase | 5.72 | 1.75E-24 | 6.09 | 5.03E-37 |
| VC1888 | hemolysin-related protein | 4.06 | 4.04E-14 | 8.63 | 4.61E-61 |
| VC1928 | C4-dicarboxylate transport protein DctQ,putative | 2.21 | 3.24E-07 | 2.92 | 1.56E-16 |
| VC1962 | lipoprotein | 2.10 | 1.40E-06 | 3.27 | 2.23E-20 |
| VC2359 | uracil-DNA glycosylase | 2.70 | 9.86E-07 | 3.19 | 2.03E-16 |
| VC2691 | periplasmic protein cpxP, putative | 2.28 | 5.08E-05 | 9.30 | 2.28E-53 |
| VCA0063 | protease II | 2.76 | 1.03E-06 | 2.29 | 5.66E-10 |
| VCA0070 | phosphate ABC transporter, periplasmicphosphate-binding protein | 3.04 | 1.24E-08 | 4.86 | 5.38E-31 |
| VCA0071 | phosphate ABC transporter, permease protein | 2.79 | 1.04E-07 | 3.11 | 1.43E-07 |
| VCA0072 | phosphate ABC transporter, permease protein | 2.45 | 1.19E-04 | 2.66 | 1.69E-07 |
| VCA0083 | multidrug resistance protein D | 4.84 | 4.39E-07 | 5.85 | 5.39E-35 |
| VCA0094 | conserved hypothetical protein | 14.05 | 4.79E-26 | 2.45 | 1.98E-03 |
| VCA0146 | conserved hypothetical protein | 2.91 | 1.07E-04 | -2.61 | 1.59E-07 |
| VCA0152 | conserved hypothetical protein | 2.04 | 2.58E-04 | -5.85 | 4.33E-33 |
| VCA0161 | tryptophanase | 5.99 | 1.14E-24 | 2.06 | 5.45E-09 |
| VCA0163 | conserved hypothetical protein | 2.72 | 2.20E-06 | 2.72 | 3.05E-10 |
| VCA0221 | lactonizing lipase | 5.71 | 1.80E-17 | -2.02 | 2.90E-06 |
| VCA0227 | iron(III) ABC transporter, periplasmiciron-compound-binding protein | 4.29 | 7.98E-16 | 10.77 | 9.88E-55 |
| VCA0231 | transcriptional regulator, AraC/XylS family | 7.89 | 4.42E-22 | 2.13 | 3.30E-06 |

| | | | | | |
|---|---|---|---|---|---|
| VCA0271 | hypothetical protein | 2.18 | 6.77E-07 | 2.42 | 1.83E-14 |
| VCA0276 | Glycine cleavage system P protein GcvP, authentic frameshif | 2.77 | 9.97E-11 | -4.31 | 5.27E-29 |
| VCA0278 | serine hydroxymethyltransferase | 2.39 | 6.88E-08 | -3.79 | 2.00E-24 |
| VCA0280 | glycine cleavage system protein GcvT; contains authentic frameshi | 2.09 | 3.53E-06 | -6.09 | 3.49E-49 |
| VCA0308 | deoxyguanosinetripho sphatetriphosphohydrolase-rela ted protein | -2.05 | 3.77E-06 | -3.77 | 1.09E-31 |
| VCA0310 | hypothetical protein | -2.14 | 6.61E-05 | -2.71 | 1.56E-21 |
| VCA0316 | acetyltransferase, putative | -2.51 | 2.09E-05 | -2.69 | 8.89E-16 |
| VCA0339 | hypothetical protein | -3.10 | 3.06E-04 | -5.50 | 1.62E-17 |
| VCA0346 | H-REV 107-related protein | -3.12 | 3.56E-12 | 2.58 | 6.75E-21 |
| VCA0355 | conserved hypothetical protein | -2.48 | 2.11E-07 | -3.14 | 4.55E-16 |
| VCA0356 | hypothetical protein | -2.11 | 1.80E-05 | -2.15 | 1.04E-16 |
| VCA0356a | hypothetical protein | -2.25 | 5.64E-04 | -2.53 | 1.77E-10 |
| VCA0386 | hypothetical protein | -2.68 | 3.71E-08 | 3.15 | 5.73E-15 |
| VCA0419 | hypothetical protein | -2.60 | 6.58E-07 | -2.57 | 7.76E-14 |
| VCA0465 | hypothetical protein | -2.10 | 4.67E-04 | -2.03 | 8.28E-08 |
| VCA0480 | hypothetical protein | -2.16 | 1.78E-04 | 2.04 | 1.46E-06 |
| VCA0519 | fructose repressor | -2.04 | 5.56E-05 | 7.90 | 2.08E-64 |
| VCA0576 | heme transport protein HutA | 6.55 | 2.04E-16 | 7.27 | 2.22E-63 |
| VCA0683 | sensor protein UhpB | 2.12 | 2.73E-04 | -2.75 | 1.66E-17 |
| VCA0690 | acetyl-CoA acetyltransferase | -2.05 | 6.52E-06 | -6.07 | 9.62E-25 |
| VCA0691 | acetoacetyl-CoA reductase | -2.18 | 1.61E-07 | -5.35 | 2.47E-36 |
| VCA0849 | hypothetical protein | 2.33 | 2.45E-05 | 2.08 | 3.12E-12 |
| VCA0864 | methyl-accepting chemotaxis protein | 3.00 | 2.23E-07 | -2.23 | 2.27E-13 |
| VCA0874 | hypothetical protein | 2.31 | 2.24E-04 | 3.29 | 7.44E-09 |
| VCA0886 | 2-amino-3-ketobutyra te coenzyme A ligase | 3.12 | 3.86E-10 | -5.41 | 2.20E-48 |
| VCA0907 | conserved hypothetical protein | 5.91 | 5.51E-24 | 2.25 | 1.96E-11 |
| VCA0909 | oxygen-independent coproporphyrinogen IIIoxidase, putative | 9.80 | 7.27E-18 | 3.74 | 7.85E-18 |
| VCA0910 | tonB1 protein | 9.83 | 2.96E-22 | 2.83 | 1.19E-05 |
| VCA0911 | TonB system transport protein ExbB1 | 10.30 | 7.54E-20 | 2.43 | 2.77E-03 |
| VCA0952 | transcriptional regulator, LuxR family | 3.48 | 3.76E-05 | 11.33 | 3.96E-58 |
| VCA0976 | hypothetical protein | 7.46 | 1.72E-15 | 3.12 | 1.02E-08 |
| VCA0977 | ABC transporter, ATP-binding protein | 5.44 | 3.75E-24 | 2.94 | 9.34E-16 |
| VCA1031 | putative methyl accepting chemotaxis protein; authentic frameshift n | 2.26 | 8.10E-06 | 7.13 | 3.58E-51 |
| VCA1041 | phosphomannomutase, putative | 2.18 | 1.07E-05 | -3.62 | 1.21E-30 |
| VCA1115 | ParA family protein | -2.02 | 1.21E-05 | 5.66 | 2.23E-47 |
| VCr025 | 5S ribosomal RNA | 4.65 | 1.50E-05 | -239.23 | 3.68E-98 |
