## Supplementary material for "Modular small RNA drives pathogen emergence": Table S3

**Table S3:** OueS-regulated biofilm genes obtained by comparing expression values of WT and Δ*ompU* -pOueS biofilm grown cells to their expression in the *ompU* mutant.

| Genes repressed by OueS | | | | | |
|---|---|---|---|---|---|
| **Locus Tag** | **Function** | **WT/Δ*ompU*** | | **Δ*ompU* -pOueS/Δ*ompU*** | |
| | | **FC** | **FDR *p* -value** | **FC** | **FDR *p* -value** |
| VC0017 | hypothetical protein | -2.7 | 1.1E-05 | -4.1 | 7.4E-21 |
| VC0063 | thiF protein | -3.1 | 1.0E-04 | -3.0 | 7.6E-05 |
| VC0065 | thiG protein | -3.3 | 1.2E-06 | -3.2 | 3.2E-05 |
| VC0066 | thiH protein | -2.5 | 4.6E-04 | -2.6 | 1.5E-05 |
| VC0091 | O-methyltransferase- related protein | -3.7 | 4.1E-07 | -2.7 | 6.1E-07 |
| VC0107 | RyhB sRNA | -8.4 | 1.6E-16 | -3.8 | 7.0E-08 |
| VC0157 | alkaline serine protease | -4.5 | 2.0E-12 | -2.2 | 4.9E-06 |
| VC0199 | hemolysin secretion ATP-binding protein,putative | -3.4 | 3.1E-05 | -3.1 | 8.9E-06 |
| VC0200 | iron(III) compound receptor | -6.5 | 6.9E-11 | -4.6 | 1.8E-11 |
| VC0202 | iron(III) ABC transporter, periplasmiciron-compound-bindi ng protein | -2.5 | 4.1E-03 | -2.9 | 3.0E-04 |
| VC0284 | putative outer membrane receptor | -3.1 | 1.3E-08 | -2.6 | 6.8E-08 |
| VC0474 | iron-regulated virulence regulatory proteinIrgB | -3.3 | 4.5E-05 | -2.1 | 9.1E-03 |
| VC0475 | iron-regulated outer membrane virulence protein,TonB receptor family | -8.2 | 2.5E-37 | -2.4 | 1.2E-09 |
| VC0608 | iron(III) ABC transporter, periplasmiciron-compound-binding protein | -4.8 | 1.2E-12 | -2.9 | 4.4E-11 |
| VC0771 | vibriobactin-specific isochorismatase | -8.0 | 3.6E-13 | -4.1 | 1.4E-09 |
| VC0773 | vibriobactin-specific isochorismate synthase | -5.0 | 1.5E-05 | -2.4 | 3.2E-04 |
| VC0774 | vibriobactin-specifi c2,3-dihydro-2,3-dihydroxybenz oate dehydrogenase | -5.8 | 1.6E-14 | -5.3 | 3.3E-18 |
| VC0775 | vibriobactin synthesis protein, putative | -2.7 | 6.6E-05 | -3.2 | 7.5E-12 |
| VC0777 | ferric vibriobactin ABC transporter, permeaseprotein | -3.5 | 1.1E-05 | -2.9 | 1.1E-05 |
| VC0820 | ToxR-activated gene A protein | -4.0 | 5.9E-09 | -2.6 | 1.6E-04 |
| VC0821 | hypothetical protein | -3.6 | 3.5E-13 | -2.4 | 2.1E-08 |
| VC0823 | hypothetical protein | -2.8 | 9.0E-09 | -2.1 | 9.8E-06 |
| VC0838 | TCP pilus virulence regulatory protein | -4.6 | 2.4E-05 | -2.3 | 4.5E-03 |
| VC0929 | RbmB | -5.8 | 6.0E-27 | -2.8 | 5.0E-11 |
| VC0930 | hemolysin-related protein, RbmC | -2.7 | 6.8E-03 | -2.9 | 1.0E-11 |
| VC0931 | RbmD | -3.1 | 3.1E-04 | -2.6 | 3.3E-05 |
| VC0932 | RbmE | -4.8 | 4.4E-07 | -2.8 | 2.6E-10 |
| VC0940 | conserved hypothetical protein | -2.1 | 3.7E-03 | -2.6 | 7.6E-05 |
| VC0972 | porin, putative | -2.3 | 3.2E-04 | -2.4 | 7.0E-06 |
| VC1029 | GGDEF family protein | -2.3 | 5.5E-03 | -2.3 | 2.0E-05 |
| VC1061 | cysteine synthase/cystathionine beta-synthasefamily protein | -3.4 | 7.6E-09 | -3.0 | 6.5E-12 |
| VC1265 | hypothetical protein | -2.4 | 3.6E-05 | -2.1 | 3.6E-04 |
| VC1268 | conserved hypothetical protein | -2.4 | 1.0E-03 | -2.5 | 1.4E-05 |
| VC1329 | opacity protein-related protein | -13.6 | 2.5E-37 | -6.7 | 4.0E-23 |
| VC1330 | hypothetical protein | -4.3 | 1.9E-12 | -2.2 | 5.4E-05 |
| VC1346 | conserved hypothetical protein | -4.9 | 2.5E-21 | -2.0 | 3.7E-05 |
| VC1456 | cholera enterotoxin, B subunit | -6.2 | 1.1E-09 | -4.0 | 5.7E-09 |
| VC1457 | cholera enterotoxin, A subunit | -3.6 | 2.2E-07 | -3.6 | 1.0E-14 |
| VC1543 | hypothetical protein | -2.6 | 8.0E-08 | -2.1 | 3.0E-05 |
| VC1544 | tonB2 protein | -2.8 | 1.3E-06 | -2.6 | 1.6E-07 |
| VC1545 | TonB system transport protein ExbD2 | -2.8 | 1.1E-04 | -3.6 | 3.6E-11 |
| VC1546 | TonB system transport protein ExbB2 | -3.7 | 5.5E-09 | -2.7 | 7.0E-08 |
| VC1547 | biopolymer transport protein ExbB-relatedprotein | -3.6 | 2.1E-11 | -2.6 | 2.7E-11 |
| VC1548 | hypothetical protein | -3.6 | 4.3E-06 | -2.2 | 1.8E-03 |
| VC1570 | quinol oxidase, subunit II | -3.9 | 1.9E-07 | -3.1 | 7.4E-06 |
| VC1571 | quinol oxidase, subunit I | -3.3 | 8.0E-08 | -2.2 | 1.0E-04 |
| VC1573 | fumarate hydratase, class II | -4.4 | 2.7E-07 | -3.2 | 7.1E-05 |
| VC1585 | catalase | -5.7 | 2.5E-10 | -3.4 | 9.7E-06 |
| VC1644 | hypothetical protein | -2.3 | 2.4E-05 | -2.4 | 3.5E-08 |
| VC1688 | hypothetical protein | -9.3 | 7.6E-12 | -7.1 | 3.6E-10 |
| VC1888 | hemolysin-related protein, Bap1 | -4.1 | 3.5E-12 | -2.3 | 9.0E-07 |
| VC1945 | FAD monooxygenase, PheA/TfdB family | -2.6 | 2.2E-06 | -3.4 | 1.0E-13 |
| VC1947 | transcriptional regulator, LysR family | -3.2 | 4.8E-06 | -6.9 | 1.4E-21 |
| VC1948 | hypothetical protein | -52.7 | 1.1E-06 | -38.3 | 3.3E-11 |
| VC1949 | pvcA protein | -6.4 | 3.5E-13 | -8.4 | 1.1E-38 |
| VC2004 | conserved hypothetical protein | -3.4 | 1.5E-10 | -2.2 | 4.0E-03 |
| VC2069 | flagellar biosynthetic protein FlhA | -2.2 | 7.7E-05 | -2.0 | 6.8E-04 |
| VC2209 | nonribosomal peptide synthetase VibF | -4.5 | 1.5E-13 | -4.1 | 3.0E-21 |
| VC2210 | vibriobactin utilization protein ViuB | -7.1 | 1.2E-28 | -3.8 | 1.9E-14 |
| VC2211 | ferric vibriobactin receptor | -5.5 | 4.5E-11 | -3.0 | 1.2E-04 |
| VC2212 | hypothetical protein | -3.1 | 5.8E-09 | -2.3 | 2.4E-05 |
| VC2539 | thiamin ABC transporter, periplasmicthiamin-binding protein | -2.7 | 2.8E-05 | -2.1 | 6.7E-06 |
| VC2667 | hypothetical protein | -6.7 | 2.8E-12 | -4.1 | 2.8E-10 |
| VC2691 | periplasmic protein cpxP, putative | -2.3 | 8.7E-04 | 3.1 | 1.2E-12 |
| VC2694 | superoxide dismutase, Mn | -3.5 | 8.2E-14 | -3.0 | 4.5E-14 |

| | | | | | |
|---|---|---|---|---|---|
| VCA0083 | multidrug resistance protein D | -4.8 | 1.4E-05 | -5.1 | 4.6E-07 |
| VCA0084 | soxR protein | -2.2 | 4.2E-03 | -2.5 | 1.3E-04 |
| VCA0094 | conserved hypothetical protein | -14.1 | 2.5E-23 | -13.3 | 3.6E-30 |
| VCA0095 | hypothetical protein | -5.8 | 5.5E-05 | -5.0 | 1.2E-04 |
| VCA0140 | spindolin-related protein | -4.7 | 1.1E-14 | -2.5 | 7.3E-07 |
| VCA0144 | immunogenic protein | -3.0 | 3.1E-03 | -3.4 | 3.2E-05 |
| VCA0146 | conserved hypothetical protein | -2.9 | 1.6E-03 | -3.2 | 8.5E-04 |
| VCA0147 | transcriptional regulator, putative | -3.6 | 1.3E-04 | -3.6 | 5.2E-10 |
| VCA0160 | tryptophan-specific transport protein | -5.2 | 7.8E-13 | -7.0 | 1.2E-21 |
| VCA0161 | tryptophanase | -6.0 | 5.3E-22 | -8.2 | 4.6E-50 |
| VCA0163 | conserved hypothetical protein | -2.7 | 5.5E-05 | -2.1 | 1.2E-04 |
| VCA0221 | lactonizing lipase | -5.7 | 3.0E-15 | -5.1 | 1.0E-18 |
| VCA0222 | lipase activator protein, putative | -3.4 | 6.9E-05 | -2.8 | 8.5E-04 |
| VCA0227 | iron(III) ABC transporter, periplasmiciron-compound-binding protein | -4.3 | 1.1E-13 | -2.1 | 4.2E-05 |
| VCA0231 | transcriptional regulator, AraC/XylS family | -7.9 | 9.6E-20 | -3.8 | 5.5E-10 |
| VCA0232 | enterobactin receptor, VctA | -6.9 | 2.9E-13 | -5.8 | 2.0E-20 |
| VCA0233 | hypothetical protein | -4.6 | 9.3E-08 | -3.4 | 4.8E-05 |
| VCA0280 | glycine cleavage system protein GcvT; contains authentic frameshift | -2.1 | 8.5E-05 | -2.1 | 7.6E-05 |
| VCA0682 | transcriptional regulator UhpA | -2.4 | 2.5E-04 | -2.8 | 3.0E-09 |
| VCA0683 | sensor protein UhpB | -2.1 | 3.6E-03 | -3.1 | 1.6E-05 |
| VCA0885 | threonine 3-dehydrogenase | -3.0 | 2.1E-11 | -3.9 | 3.7E-17 |
| VCA0886 | 2-amino-3-ketobutyrate coenzyme A ligase | -3.1 | 2.1E-08 | -3.4 | 9.4E-17 |
| VCA0907 | conserved hypothetical protein | -5.9 | 1.8E-21 | -3.7 | 4.7E-14 |
| VCA0908 | conserved hypothetical protein | -7.5 | 2.6E-28 | -4.1 | 1.1E-17 |
| VCA0909 | oxygen-independent coproporphyrinogen IIIoxidase, putative | -9.8 | 1.3E-15 | -4.6 | 6.6E-11 |
| VCA0910 | tonB1 protein | -9.8 | 7.3E-20 | -4.9 | 1.3E-16 |
| VCA0911 | TonB system transport protein ExbB1 | -10.3 | 1.5E-17 | -5.5 | 6.1E-16 |
| VCA0912 | TonB system transport protein ExbD1 | -7.3 | 2.2E-11 | -4.2 | 1.6E-16 |
| VCA0913 | hemin ABC transporter, periplasmic hemin-bindingprotein HutB | -6.6 | 8.0E-10 | -5.5 | 1.2E-12 |
| VCA0914 | hemin ABC transporter, permease protein,putative | -5.6 | 6.5E-11 | -5.1 | 4.0E-15 |
| VCA0928 | hypothetical protein | -2.4 | 1.2E-03 | -2.4 | 7.9E-06 |
| VCA0946 | maltose/maltodextrin ABC transporter,ATP-binding protein | -2.3 | 3.5E-05 | -4.0 | 4.8E-21 |
| VCA0976 | hypothetical protein | -7.5 | 2.1E-13 | -3.8 | 2.7E-07 |
| VCA0977 | ABC transporter, ATP-binding protein | -5.4 | 1.4E-21 | -3.5 | 3.1E-14 |
| VCA1041 | phosphomannomutase, putative | -2.2 | 2.2E-04 | -2.2 | 1.4E-06 |
| VCr025 | 5S ribosomal RNA | -4.7 | 3.0E-04 | -5.1 | 3.1E-05 |

| Genes activated by OueS | | | | | |
|---|---|---|---|---|---|
| Locus Tag | Function | WT/∆*ompU* | | ∆*ompU* -pOueS/∆*ompU* | |
| | | FC | FDR *p*-value | FC | FDR *p*-value |
| VC0176 | transcriptional regulator, putative | 2.8 | 7.1E-03 | 3.3 | 3.2E-05 |
| VC0177 | hypothetical protein | 2.6 | 9.1E-05 | 3.6 | 2.3E-12 |
| VC0178 | patatin-related protein | 2.2 | 1.7E-04 | 2.5 | 5.6E-08 |
| VC0179 | hypothetical protein | 2.5 | 9.0E-06 | 2.2 | 4.0E-08 |
| VC0180 | conserved hypothetical protein | 2.3 | 1.2E-06 | 2.1 | 7.4E-05 |
| VC0494 | conserved hypothetical protein | 2.9 | 1.7E-04 | 3.8 | 1.1E-08 |
| VC0502 | type IV pilin, putative | 2.7 | 1.1E-06 | 3.1 | 5.7E-11 |
| VC0503 | conserved hypothetical protein | 3.3 | 7.9E-12 | 2.9 | 1.9E-14 |
| VC0506 | hypothetical protein | 2.6 | 5.0E-04 | 2.3 | 3.6E-06 |
| VC0507 | hypothetical protein | 4.8 | 2.2E-04 | 4.8 | 8.7E-04 |
| VC0633 | outer membrane protein OmpU | 15766.6 | 0.0E+00 | 2471.8 | 0.0E+00 |
| VC0817 | transposase, putative | 2.1 | 2.4E-03 | 2.5 | 1.4E-07 |
| VC0818 | pseudogene; within VPI-I; unknown function | 2.2 | 5.2E-03 | 2.6 | 5.8E-04 |
| VC0825 | toxin co-regulated pilus biosynthesis protein I | 2.1 | 4.5E-03 | 3.4 | 1.4E-10 |
| VC1472 | hypothetical protein | 4.3 | 1.5E-04 | 4.1 | 1.2E-03 |
| VC1762 | hypothetical protein | 2.1 | 1.1E-03 | 2.1 | 2.8E-03 |
| VC1763 | chemotaxis protein MotB-related protein | 2.8 | 1.1E-05 | 2.7 | 2.7E-04 |
| VC1774 | conserved hypothetical protein | 2.0 | 3.7E-03 | 2.3 | 3.3E-05 |
| VC1793 | hypothetical protein | 2.9 | 5.3E-03 | 3.3 | 8.8E-04 |
| VCA0346 | H-REV 107-related protein | 3.1 | 2.4E-10 | 2.1 | 1.4E-05 |
| VCA0367 | hypothetical protein | 2.1 | 2.8E-04 | 2.3 | 9.9E-07 |
| VCA0386 | hypothetical protein | 2.7 | 1.5E-06 | 2.8 | 3.0E-09 |
| VCA0420 | hypothetical protein | 2.5 | 6.1E-04 | 2.1 | 2.4E-05 |
| VCA0443 | lipoprotein Blc | 2.4 | 1.0E-04 | 2.1 | 1.2E-04 |
| VCA0446 | haemagglutinin | 3.6 | 6.6E-11 | 2.0 | 7.1E-04 |
| VCA0449 | hypothetical protein | 4.7 | 9.7E-06 | 3.5 | 1.4E-03 |
| VCA0450 | hypothetical protein | 2.7 | 8.3E-03 | 2.8 | 9.9E-04 |
| VCA0508 | transposase OrfAB, subunit B | 2.8 | 4.6E-07 | 2.4 | 6.5E-04 |
| 16Sh | 16S ribosomal RNA | 2.3 | 2.1E-05 | 12.2 | 1.0E-43 |
| tRNA-Gly-8 | tRNA biosynthesis | 2.1 | 2.4E-03 | 3.0 | 8.4E-08 |
| tRNA-Met-8 | tRNA biosynthesis | 2.9 | 1.8E-04 | 3.1 | 1.5E-05 |
