## Supplementary material for "Modular small RNA drives pathogen emergence": Table S4

**Table S4:** Differentially expressed genes in the *ompU* and *rpoE* mutants compared to WT C6706 when grown in virulence inducing AKI conditions.

789 differentially-expressed genes in AKI-grown Δ*ompU* cells, independent of Δ*rpoE*, compared to WT C6706

| Locus Tag | Function | FC in Δ*ompU* | *p*-value |
|---|---|---|---|
| VC0001 | hypothetical protein | -2.30 | 7.11E-04 |
| VC0002 | mioC protein | 2.17 | 2.19E-07 |
| VC0018 | 16 kDa heat shock protein A | -2.02 | 1.17E-05 |
| VC0032 | ComM-related protein | -2.25 | 2.01E-04 |
| VC0049 | smg protein | -2.73 | 2.77E-12 |
| VC0050 | DNA topoisomerase I-related protein | -2.22 | 1.32E-07 |
| VC0070 | hypothetical protein | 2.02 | 6.54E-07 |
| VC0076 | universal stress protein A | -2.52 | 1.86E-09 |
| VC0078 | ferritin | -2.69 | 6.51E-15 |
| VC0079 | conserved hypothetical protein | -6.23 | 5.29E-52 |
| VC0080 | conserved hypothetical protein | -2.16 | 1.63E-08 |
| VC0089 | cytochrome c551 peroxidase | -20.61 | 3.85E-71 |
| VC0090 | DNA-damage-inducible protein F | -4.44 | 1.57E-26 |
| VC0098 | methyl-accepting chemotaxis protein | -3.40 | 2.04E-26 |
| VC0134 | conserved hypothetical protein | 2.04 | 2.40E-15 |
| VC0139 | DPS family protein | -4.21 | 7.81E-22 |
| VC0142 | hypothetical protein | -2.42 | 2.89E-09 |
| VC0156 | vitamin B12 receptor | 6.26 | 1.15E-26 |
| VC0164 | multidrug resistance protein, putative | 2.68 | 3.83E-15 |
| VC0165 | conserved hypothetical protein | 3.06 | 1.01E-10 |
| VC0166 | transcriptional regulator, TetR family | 2.78 | 9.52E-13 |
| VC0169 | hypothetical protein | -6.72 | 6.99E-25 |
| VC0172 | peptide ABC transporter, permease protein | -2.06 | 3.08E-07 |
| VC0174 | hypothetical protein | -2.03 | 4.01E-15 |
| VC0185 | transposase, putative | -3.19 | 5.71E-16 |
| VC0191 | conserved hypothetical protein | -2.00 | 1.04E-05 |
| VC0194 | gamma-glutamyltransp eptidase | 4.29 | 5.66E-26 |
| VC0228 | hypothetical protein | 2.38 | 7.43E-14 |
| VC0242 | phosphomannomutase | 2.26 | 2.50E-10 |
| VC0243 | GDP-mannose 4,6-dehydratase | 2.44 | 8.73E-11 |
| VC0273 | DNA-binding protein HU | 2.06 | 2.34E-05 |
| VC0281 | lysine decarboxylase, inducible | -3.22 | 5.70E-08 |
| VC0295 | acetyl-CoA carboxylase, biotin carboxylase | 3.38 | 1.97E-20 |
| VC0296 | acetyl-CoA carboxylase, biotin carboxyl carrierprotein | 3.17 | 8.26E-25 |
| VC0297 | 3-dehydroquinate dehydratase | 2.20 | 1.01E-10 |
| VC0298 | acetyl-CoA synthase | -10.02 | 2.66E-43 |
| VC0304 | guanosine-5 | -2.23 | 3.24E-16 |
| VC0308 | hypothetical protein | -3.46 | 5.04E-20 |
| VC0330 | regulator of sigma D | -2.30 | 1.95E-11 |
| VC0354 | peptidyl-prolyl cis-trans isomerase, FKBP-type | 3.20 | 3.10E-23 |
| VC0357 | conserved hypothetical protein | -2.01 | 5.99E-08 |
| VC0360 | ribosomal protein S7 | 3.15 | 2.32E-27 |
| VC0361 | elongation factor G | 2.01 | 9.91E-09 |
| VC0365 | bacterioferritin | -3.46 | 4.82E-22 |
| VC0367 | primosomal replication protein N | 2.48 | 3.09E-14 |
| VC0370 | conserved hypothetical protein | 2.91 | 9.52E-11 |
| VC0377 | conserved hypothetical protein | -2.33 | 9.32E-12 |
| VC0378 | zinc uptake regulation protein, putative | -2.34 | 4.39E-12 |
| VC0383 | hypothetical protein | -4.12 | 4.41E-09 |
| VC0395 | UTP--glucose-1-phosp hate uridylyltransferase | 2.32 | 4.59E-12 |
| VC0397 | single-strand binding protein | 2.22 | 1.13E-09 |
| VC0403 | MSHA biogenesis protein MshM | -3.10 | 4.37E-14 |
| VC0404 | MSHA biogenesis protein MshN | -2.32 | 3.13E-13 |
| VC0405 | MSHA biogenesis protein MshE | -2.01 | 4.65E-10 |
| VC0407 | MSHA biogenesis protein MshF | -2.12 | 1.82E-08 |
| VC0412 | hypothetical protein | -2.23 | 8.91E-14 |
| VC0413 | hypothetical protein | -2.76 | 4.25E-21 |
| VC0414 | hypothetical protein | -2.25 | 2.47E-11 |
| VC0445 | survival protein SurA | 2.34 | 2.91E-09 |
| VC0452 | A/G-specific adenine glycosylase | 2.34 | 3.11E-09 |
| VC0462 | twitching motility protein PilT | -2.39 | 3.20E-09 |
| VC0463 | twitching motility protein PilT | -2.60 | 1.10E-19 |
| VC0465 | tyrosyl-tRNA synthetase | -2.18 | 5.66E-07 |
| VC0472 | S-adenosylmethionine synthase | 2.25 | 1.01E-12 |
| VC0478 | fructose-bisphosphat e aldolase, class II | 2.46 | 4.06E-26 |
| VC0483 | conserved hypothetical protein | 2.73 | 1.21E-18 |
| VC0485 | pyruvate kinase I | 2.11 | 2.97E-05 |
| VC0496 | hypothetical protein | -2.25 | 8.04E-05 |
| VC0501 | pseudogene within VSP-II; potential transposase | -3.27 | 1.16E-15 |
| VC0533 | lipoprotein NlpD | -2.18 | 8.74E-09 |
| VC0534 | RNA polymerase sigma-38 factor | -4.11 | 1.50E-23 |
| VC0549 | hypothetical protein | -2.21 | 1.70E-11 |
| VC0550 | oxaloacetate decarboxylase, alpha subunit | -2.68 | 7.49E-09 |
| VC0551 | oxaloacetate decarboxylase, beta subunit | -2.14 | 2.26E-07 |
| VC0552 | quinone oxidoreductase | -2.14 | 1.95E-13 |
| VC0596 | dnaK suppressor protein | 3.15 | 1.51E-11 |
| VC0613 | beta-N-acetylhexosam inidase | -2.19 | 1.57E-06 |
| VC0614 | conserved hypothetical protein | -2.68 | 3.49E-06 |
| VC0615 | endoglucanase-relate d protein | -2.38 | 1.26E-08 |
| VC0618 | peptide ABC transporter, permease protein | -2.62 | 3.33E-05 |
| VC0619 | peptide ABC transporter, permease protein | -2.03 | 1.39E-03 |
| VC0620 | peptide ABC transporter, periplasmicpeptide-binding protein | -2.60 | 2.26E-13 |
| VC0631 | tyrosyl-tRNA synthetase | 2.57 | 2.72E-12 |
| VC0641 | conserved hypothetical protein | 2.12 | 3.10E-12 |
| VC0647 | polyribonucleotide nucleotidyltransferase | 2.28 | 1.39E-11 |
| VC0649 | transcriptional regulator, MarR family | -2.23 | 1.37E-07 |
| VC0658 | c-di-GMP phosphodiesterase A-related protein | -4.99 | 3.61E-33 |
| VC0661 | conserved hypothetical protein | -2.74 | 1.31E-10 |
| VC0692 | beta-hexosaminidase | 2.31 | 3.47E-10 |
| VC0695 | phospho-2-dehydro-3- deoxyheptonate aldolase,tyr-sensitive | 3.72 | 3.41E-20 |
| VC0697 | hypothetical protein | -3.96 | 2.47E-29 |

| | | | |
|---|---|---|---|
| VC0703 | c-di-GMP phosphodiesterase A-related protein | 2.05 | 4.74E-07 |
| VC0728 | conserved hypothetical protein | -4.83 | 5.72E-28 |
| VC0731 | antioxidant, AhpC/Tsa family | 2.10 | 3.02E-07 |
| VC0736 | isocitrate lyase | -6.45 | 1.48E-44 |
| VC0737 | acetoin utilization protein AcuB, putative | -5.32 | 7.50E-22 |
| VC0749 | NifU-related protein | 2.16 | 6.23E-08 |
| VC0754 | conserved hypothetical protein | 2.11 | 1.18E-05 |
| VC0771 | vibriobactin-specifi c isochorismatase | 3.91 | 1.14E-13 |
| VC0790 | transcriptional regulator CitB | -2.33 | 5.27E-05 |
| VC0804 | ferredoxin | -2.28 | 2.83E-04 |
| VC0812 | helicase-related protein | 2.20 | 1.69E-09 |
| VC0817 | transposase, putative | -2.04 | 5.05E-07 |
| VC0820 | ToxR-activated gene A protein | 2.05 | 1.08E-05 |
| VC0821 | hypothetical protein | 2.09 | 2.61E-08 |
| VC0840 | accessory colonization factor AcfB | 2.28 | 3.37E-07 |
| VC0841 | accessory colonization factor AcfC | 2.02 | 4.50E-04 |
| VC0847 | integrase, phage family | 2.17 | 2.61E-09 |
| VC0854 | heat shock protein GrpE | 2.39 | 7.57E-18 |
| VC0856 | dnaJ protein | 2.04 | 1.56E-10 |
| VC0871 | hypothetical protein | 2.27 | 1.12E-06 |
| VC0872 | conserved hypothetical protein | 2.34 | 1.62E-06 |
| VC0875 | prolyl-tRNA synthetase | 2.42 | 2.66E-12 |
| VC0882 | conserved hypothetical protein | 2.12 | 1.38E-04 |
| VC0884 | acetyltransferase-re lated protein | -2.14 | 1.45E-07 |
| VC0892 | chemotaxis protein PomA | -3.14 | 1.84E-17 |
| VC0894 | thiamin biosynthesis protein ThiI | 2.25 | 3.01E-11 |
| VC0901 | hypothetical protein | 2.32 | 3.02E-16 |
| VC0910 | PTS system, trehalose-specific IIBC component | -2.48 | 4.65E-10 |
| VC0928 | hypothetical protein | 3.08 | 2.46E-15 |
| VC0935 | hypothetical protein | -2.90 | 4.24E-09 |
| VC0936 | polysaccharide export-related protein | -5.54 | 1.29E-19 |
| VC0937 | exopolysaccharide biosynthesis protein,putative | -2.56 | 2.07E-11 |
| VC0938 | hypothetical protein | -4.19 | 4.46E-13 |
| VC0957 | conserved hypothetical protein | -4.36 | 1.31E-27 |
| VC0962 | conserved hypothetical protein | 2.47 | 7.02E-11 |
| VC0985 | heat shock protein HtpG | 3.94 | 5.75E-18 |
| VC0995 | PTS system, N-acetylglucosamine-specific IIABCcomponent | -2.10 | 7.43E-11 |
| VC1008 | sodium-type flagellar protein MotY | -3.67 | 3.75E-25 |
| VC1009 | conserved hypothetical protein | -4.19 | 1.81E-29 |
| VC1029 | GGDEF family protein | -3.63 | 6.64E-16 |
| VC1031 | inosine monophosphate dehydrogenase-relatedprotein | -10.24 | 4.21E-77 |
| VC1039 | asmA protein | 3.09 | 1.46E-21 |
| VC1040 | cob(I)alamin adenosyltransferase | 2.91 | 1.71E-18 |
| VC1043 | long-chain fatty acid transport protein | -2.35 | 2.35E-11 |
| VC1050 | response regulator | -5.11 | 8.29E-36 |
| VC1057 | proteinase inhibitor, putative | -2.30 | 4.60E-05 |
| VC1064 | lipoprotein-related protein | -2.79 | 7.05E-16 |
| VC1066 | hypothetical protein | -3.97 | 3.04E-23 |
| VC1080 | hypothetical protein | -3.79 | 3.64E-25 |
| VC1081 | response regulator | -4.90 | 1.00E-33 |
| VC1082 | response regulator | -11.21 | 7.83E-57 |
| VC1083 | hypothetical protein | -10.25 | 4.88E-67 |
| VC1084 | sensory box sensor histidine kinase | -12.57 | 1.86E-46 |
| VC1085 | sensor histidine kinase | -7.96 | 9.90E-45 |
| VC1086 | response regulator | -5.24 | 2.72E-46 |
| VC1087 | response regulator | -4.39 | 1.48E-31 |
| VC1088 | sensor histidine kinase | -5.87 | 4.64E-35 |
| VC1089 | periplasmic binding protein-related protein | -4.69 | 7.66E-33 |
| VC1093 | oligopeptide ABC transporter, permease protein | 3.04 | 3.37E-14 |
| VC1095 | oligopeptide ABC transporter, ATP-bindingprotein | 2.05 | 1.35E-08 |
| VC1099 | conserved hypothetical protein | -2.11 | 4.01E-07 |
| VC1100 | hypothetical protein | -2.12 | 1.56E-06 |
| VC1118 | transcriptional regulator, putative | -2.10 | 1.95E-07 |
| VC1119 | oxidoreductase, short-chaindehydrogenase/reduc tase family | -2.92 | 4.10E-10 |
| VC1120 | conserved hypothetical protein | -2.19 | 3.39E-09 |
| VC1121 | conserved hypothetical protein | -2.13 | 1.59E-07 |
| VC1124 | conserved hypothetical protein | -4.65 | 1.15E-25 |
| VC1125 | hypothetical protein | -4.90 | 2.76E-27 |
| VC1126 | adenylosuccinate lyase | 2.46 | 7.18E-11 |
| VC1131 | conserved hypothetical protein | -2.27 | 1.46E-08 |
| VC1147 | iron-containing alcohol dehydrogenase | -3.03 | 7.10E-19 |
| VC1153 | conserved hypothetical protein | -2.45 | 1.59E-15 |
| VC1155 | response regulator | -3.76 | 2.97E-29 |
| VC1156 | sensor histidine kinase | -2.47 | 3.84E-13 |
| VC1159 | phosphoserine aminotransferase | 2.47 | 1.71E-14 |
| VC1185 | GGDEF family protein | -2.41 | 2.46E-08 |
| VC1189 | hypothetical protein | -4.50 | 1.89E-20 |
| VC1193 | hypothetical protein | 4.87 | 8.78E-27 |
| VC1195 | lipoprotein, putative | 4.13 | 4.85E-26 |
| VC1208 | conserved hypothetical protein | 2.72 | 1.70E-19 |
| VC1209 | elongation factor P family protein | 3.07 | 1.44E-23 |
| VC1211 | conserved hypothetical protein | -2.11 | 4.68E-07 |
| VC1216 | GGDEF family protein | -2.38 | 1.56E-11 |
| VC1223 | hypothetical protein | -3.89 | 1.28E-18 |
| VC1224 | hypothetical protein | -4.29 | 3.65E-17 |
| VC1235 | sodium/dicarboxylate symporter | 2.28 | 4.21E-08 |
| VC1236 | PilB-related protein | -2.80 | 4.77E-13 |
| VC1240 | alpha-ribazole-5`-ph osphate phosphatase CobC,putative | 2.05 | 6.59E-07 |
| VC1246 | hypothetical protein | 2.61 | 3.39E-15 |
| VC1247 | hypothetical protein | -2.58 | 2.74E-11 |
| VC1248 | methyl-accepting chemotaxis protein | -10.03 | 7.68E-58 |
| VC1249 | conserved hypothetical protein | -5.61 | 6.63E-43 |
| VC1259 | conserved hypothetical protein | 2.67 | 1.29E-08 |
| VC1262 | hypothetical protein | -2.12 | 4.04E-06 |
| VC1269 | conserved hypothetical protein | -2.83 | 5.12E-19 |
| VC1271 | hypothetical protein | -2.03 | 5.12E-04 |
| VC1272 | hypothetical protein | 2.51 | 4.97E-07 |

| | | | |
|---|---|---|---|
| VC1289 | methyl-accepting chemotaxis protein | 2.42 | 1.23E-14 |
| VC1295 | conserved hypothetical protein | -2.87 | 5.54E-14 |
| VC1298 | methyl-accepting chemotaxis protein | -2.11 | 1.36E-05 |
| VC1313 | methyl-accepting chemotaxis protein | -4.18 | 3.37E-21 |
| VC1315 | sensor histidine kinase | -3.50 | 4.44E-10 |
| VC1316 | chemotaxis protein CheY, putative | -4.26 | 1.70E-33 |
| VC1317 | conserved hypothetical protein | 2.87 | 6.48E-19 |
| VC1318 | outer membrane protein OmpV | 4.57 | 2.54E-18 |
| VC1322 | conserved hypothetical protein | -5.75 | 1.38E-29 |
| VC1323 | hypothetical protein | -3.76 | 1.63E-21 |
| VC1340 | prpE protein | -2.14 | 1.53E-08 |
| VC1348 | response regulator | -5.12 | 4.72E-27 |
| VC1349 | sensory box sensor histidine kinase/responseregulator | -4.38 | 1.43E-26 |
| VC1353 | GGDEF family protein | -2.07 | 2.85E-06 |
| VC1359 | amino acid ABC transporter, ATP-binding protein | -2.96 | 1.69E-14 |
| VC1360 | amino acid ABC transporter, permease protein | -4.06 | 1.63E-18 |
| VC1361 | amino acid ABC transporter, permease protein | -3.99 | 2.10E-20 |
| VC1362 | amino acid ABC transporter, periplasmic aminoacid-binding protein | -9.03 | 1.86E-54 |
| VC1368 | hypothetical protein | -3.42 | 2.74E-18 |
| VC1369 | conserved hypothetical protein | -6.14 | 1.98E-32 |
| VC1371 | hypothetical protein | -2.83 | 3.88E-14 |
| VC1374 | DnaK-related protein | 3.14 | 8.27E-15 |
| VC1375 | hypothetical protein | 2.91 | 5.78E-13 |
| VC1376 | GGDEF family protein | -2.76 | 1.52E-17 |
| VC1391 | multidrug transporter, putative | 2.31 | 1.50E-08 |
| VC1393 | sugE protein | -2.27 | 2.95E-08 |
| VC1394 | methyl-accepting chemotaxis protein | -6.06 | 1.57E-47 |
| VC1395 | response regulator cheY1 | -7.29 | 2.58E-57 |
| VC1396 | hypothetical protein | -6.61 | 2.72E-35 |
| VC1397 | chemotaxis protein CheA | -9.00 | 5.14E-57 |
| VC1398 | chemotaxis protein CheY | -9.32 | 8.38E-45 |
| VC1399 | chemotaxis protein methyltransferase CheR | -8.30 | 3.89E-52 |
| VC1400 | hypothetical protein | -7.28 | 1.95E-37 |
| VC1403 | methyl-accepting chemotaxis protein | -4.70 | 1.11E-24 |
| VC1405 | methyl-accepting chemotaxis protein | -3.05 | 1.39E-16 |
| VC1406 | methyl-accepting chemotaxis protein | -3.42 | 8.09E-18 |
| VC1410 | multidrug resistance protein VceA | 2.43 | 5.08E-10 |
| VC1414 | thermostable carboxypeptidase 1 | 2.43 | 2.50E-07 |
| VC1416 | vgrG protein | 4.88 | 7.98E-47 |
| VC1417 | hypothetical protein | 3.15 | 1.25E-16 |
| VC1418 | hypothetical protein | 2.85 | 3.19E-16 |
| VC1419 | hypothetical protein | 2.47 | 1.75E-11 |
| VC1425 | spermidine/putrescin e ABC transporter,periplasmic spermidine/putrescine-binding protein | 2.26 | 4.78E-11 |
| VC1433 | conserved hypothetical protein | -2.61 | 1.36E-13 |
| VC1486 | ABC transporter, ATP-binding protein | 2.01 | 8.91E-15 |
| VC1487 | conserved hypothetical protein | 2.61 | 2.94E-06 |
| VC1498 | conserved hypothetical protein | 2.35 | 1.93E-09 |
| VC1517 | hypothetical protein | -2.08 | 3.43E-08 |
| VC1528 | hypothetical protein | -2.99 | 6.05E-14 |
| VC1535 | methyl-accepting chemotaxis protein | -2.53 | 1.77E-05 |
| VC1538 | hypothetical protein | -2.87 | 2.20E-15 |
| VC1539 | conserved hypothetical protein | -2.61 | 7.02E-12 |
| VC1549 | glycerol-3-phosphate ABC transporter,periplasmic glycerol-3-phosphate-binding protein | -2.02 | 5.97E-04 |
| VC1551 | glycerol-3-phosphate ABC transporter, permeaseprotein | -2.92 | 6.83E-05 |
| VC1577 | hypothetical protein | 2.84 | 2.53E-15 |
| VC1578 | hypothetical protein | 2.74 | 2.11E-10 |
| VC1579 | enterobactin synthetase component F-relatedprotein | 2.63 | 3.77E-19 |
| VC1581 | NADH dehydrogenase, putative | -2.07 | 3.22E-03 |
| VC1601 | hypothetical protein | -3.21 | 1.63E-15 |
| VC1602 | chemotaxis protein CheV | -3.10 | 6.91E-10 |
| VC1603 | hypothetical protein | -3.30 | 2.08E-19 |
| VC1612 | fimbrial biogenesis and twitching motilityprotein, putative | -2.72 | 2.10E-13 |
| VC1613 | hypothetical protein | -4.25 | 1.52E-22 |
| VC1614 | conserved hypothetical protein | -2.34 | 2.20E-09 |
| VC1623 | carboxynorspermidine decarboxylase | 3.25 | 7.45E-24 |
| VC1624 | conserved hypothetical protein | 2.02 | 5.90E-07 |
| VC1625 | Pseudogene; involved in spermidine biosynthesis | 3.58 | 7.16E-25 |
| VC1641 | conserved hypothetical protein | -3.13 | 8.84E-24 |
| VC1643 | methyl-accepting chemotaxis protein | -5.49 | 2.04E-21 |
| VC1659 | conserved hypothetical protein | -2.57 | 6.67E-12 |
| VC1660 | ABC transporter ATP-binding protein | -3.34 | 1.87E-18 |
| VC1660a | ABC transporter permease | -2.90 | 3.10E-13 |
| VC1662 | conserved hypothetical protein | -3.34 | 1.07E-16 |
| VC1663 | heat shock protein HslJ | 3.36 | 2.33E-24 |
| VC1672 | DNA-3-methyladenine glycosidase I | -2.90 | 6.65E-11 |
| VC1673 | transporter, AcrB/D/F family | -2.36 | 9.63E-09 |
| VC1674 | periplasmic linker protein, putative | -3.18 | 6.76E-17 |
| VC1675 | multidrug resistance protein, putative | -2.82 | 8.93E-17 |
| VC1699 | hypothetical protein | -2.29 | 2.05E-16 |
| VC1701 | conserved hypothetical protein | 2.08 | 9.24E-04 |
| VC1704 | 5-methyltetrahydropt eroyltriglutamate--homocystein e methyltransferase | 2.14 | 1.17E-08 |
| VC1710 | conserved hypothetical protein | -2.12 | 3.53E-11 |
| VC1723 | conserved hypothetical protein | -2.43 | 1.26E-08 |
| VC1736 | arginyl-tRNA-protein transferase-relatedprotein | -3.30 | 3.87E-20 |
| VC1738 | hypothetical protein | 2.46 | 1.70E-07 |
| VC1748 | hypothetical protein | 2.03 | 8.37E-05 |
| VC1750 | hypothetical protein | 2.10 | 8.91E-09 |
| VC1758 | integrase, phage family | 2.01 | 1.45E-07 |
| VC1778 | conserved hypothetical protein | -2.76 | 3.18E-16 |
| VC1812 | conserved hypothetical protein | -2.14 | 9.34E-04 |
| VC1820 | PTS system, fructose-specific IIA component | 54.45 | 3.79E-22 |
| VC1821 | PTS system, fructose-specific IIBC component | 2.53 | 5.12E-10 |
| VC1822 | PTS system, fructose-specific IIABC component | 4.36 | 3.68E-25 |
| VC1823 | PTS system, fructose-specific IIB component | 4.85 | 7.59E-27 |
| VC1825 | transcriptional regulator | 3.49 | 9.30E-14 |
| VC1826 | PTS system, fructose-specific IIABC component | 2.99 | 2.36E-10 |
| VC1831 | sensor histidine kinase | -3.42 | 1.11E-22 |

| | | | |
|---|---|---|---|
| VC1835 | peptidoglycan-associ ated lipoprotein | 3.94 | 3.46E-30 |
| VC1841 | conserved hypothetical protein | -2.75 | 4.54E-16 |
| VC1842 | conserved hypothetical protein | -2.13 | 2.77E-04 |
| VC1843 | cytochrome d ubiquinol oxidase, subunit II | -2.85 | 6.25E-16 |
| VC1844 | cytochrome d ubiquinol oxidase, subunit I | -3.07 | 5.60E-17 |
| VC1849 | peptidyl-prolyl cis-trans isomerase B | 2.68 | 2.22E-16 |
| VC1851 | conserved hypothetical protein | -4.78 | 5.95E-33 |
| VC1853 | conserved hypothetical protein | 2.77 | 1.66E-19 |
| VC1867 | conserved hypothetical protein | -2.28 | 2.00E-08 |
| VC1868 | methyl-accepting chemotaxis protein | -6.82 | 4.06E-44 |
| VC1872 | conserved hypothetical protein | -10.36 | 1.02E-61 |
| VC1873 | conserved hypothetical protein | -13.64 | 6.01E-51 |
| VC1874 | conserved hypothetical protein | -12.48 | 3.91E-85 |
| VC1876 | conserved hypothetical protein | 2.61 | 1.10E-11 |
| VC1883 | ABC transporter, ATP-binding protein | 2.03 | 3.46E-08 |
| VC1889 | ribosomal-protein-se rine acetyltransferase,putative | -2.11 | 5.03E-06 |
| VC1898 | methyl-accepting chemotaxis protein | -2.21 | 3.06E-13 |
| VC1905 | alanine dehydrogenase | 3.67 | 1.11E-28 |
| VC1920 | ATP-dependent protease LA | 2.43 | 6.17E-14 |
| VC1929 | C4-dicarboxylate-bin ding periplasmic protein | 3.02 | 5.15E-16 |
| VC1932 | hypothetical protein | -2.56 | 3.48E-07 |
| VC1933 | hypothetical protein | -3.78 | 1.32E-22 |
| VC1934 | GGDEF family protein | -3.82 | 4.49E-22 |
| VC1935 | CDP-diacylglycerol-- glycerol-3-phosphate3-phosphat idyltransferase-related protein | 2.40 | 5.83E-10 |
| VC1936 | phosphatidate cytidylyltransferase, putative | 3.18 | 6.28E-13 |
| VC1937 | conserved hypothetical protein | 2.65 | 3.18E-07 |
| VC1944 | PvcB protein | -2.44 | 2.49E-06 |
| VC1945 | FAD monooxygenase, PheA/TfdB family | -2.11 | 2.38E-06 |
| VC1947 | transcriptional regulator, LysR family | 2.08 | 3.05E-06 |
| VC1950 | biotin sulfoxide reductase | -4.11 | 1.52E-22 |
| VC1953 | NupC family protein | -2.35 | 1.96E-05 |
| VC1962 | lipoprotein | 5.58 | 7.92E-57 |
| VC1967 | methyl-accepting chemotaxis protein | -3.39 | 5.40E-16 |
| VC1987 | outer membrane lipoprotein Slp, putative | -2.09 | 1.09E-08 |
| VC1991 | hypothetical protein | -4.50 | 1.12E-27 |
| VC1994 | protease IV | 2.16 | 3.08E-11 |
| VC1997 | hypothetical protein | -2.47 | 2.84E-14 |
| VC2005 | hypothetical protein | -3.55 | 2.01E-26 |
| VC2008 | pyruvate kinase II | -3.20 | 5.97E-19 |
| VC2009 | conserved hypothetical protein | -2.69 | 1.82E-15 |
| VC2012 | sodium-dependent transporter | -2.09 | 1.10E-08 |
| VC2022 | malonyl Coa-acyl carrier protein transacylase | 3.20 | 2.12E-15 |
| VC2026 | conserved hypothetical protein | 2.45 | 2.30E-11 |
| VC2038 | hypothetical protein | -2.42 | 2.04E-07 |
| VC2044 | conserved hypothetical protein | 3.41 | 7.17E-27 |
| VC2045 | superoxide dismutase, Fe | 2.18 | 1.94E-09 |
| VC2046 | hypothetical protein | -3.44 | 7.98E-19 |
| VC2047 | oxidoreductase, short-chaindehydrogenase/reduc tase family | -3.41 | 8.51E-27 |
| VC2053 | cytochrome c-type biogenesis protein CcmE | 2.20 | 1.14E-08 |
| VC2058 | hypothetical protein | -3.03 | 1.09E-23 |
| VC2059 | purine-binding chemotaxis protein CheW | -2.83 | 9.36E-18 |
| VC2060 | conserved hypothetical protein | -2.85 | 1.98E-09 |
| VC2061 | ParA family protein | -2.44 | 7.06E-14 |
| VC2062 | protein-glutamate methylesterase CheB | -2.48 | 4.01E-13 |
| VC2063 | chemotaxis protein CheA | -2.74 | 2.86E-15 |
| VC2064 | chemotaxis protein CheZ | -2.40 | 2.57E-07 |
| VC2065 | chemotaxis protein CheY | -2.44 | 3.59E-07 |
| VC2067 | MinD-related protein | -2.05 | 6.33E-10 |
| VC2070 | phosphohistidine phosphatase | -2.24 | 1.14E-09 |
| VC2078 | ferrous iron transport protein A | 3.74 | 4.66E-16 |
| VC2103 | transcriptional regulator, LysR family | -2.70 | 5.34E-12 |
| VC2105 | hypothetical protein | -2.10 | 3.13E-06 |
| VC2107 | aspartate-semialdehy de dehydrogenase, putative | 2.24 | 3.48E-19 |
| VC2109 | pseudogene; 3' encodes Vcr076 | 2.12 | 7.20E-09 |
| VC2124 | flagellar protein FliO | -2.75 | 3.06E-15 |
| VC2128 | flagellar hook-length control protein FliK,putative | -4.36 | 1.08E-17 |
| VC2139 | flagellar rod protein FlaI, putative | -2.86 | 5.76E-25 |
| VC2141 | flagellin FlaG | -4.12 | 1.12E-26 |
| VC2142 | flagellin FlaB | -4.56 | 6.73E-27 |
| VC2143 | flagellin FlaD | -3.37 | 2.13E-17 |
| VC2144 | flagellin FlaE | -4.36 | 8.81E-34 |
| VC2161 | methyl-accepting chemotaxis protein | -4.10 | 2.14E-26 |
| VC2171 | uracil permease | 2.29 | 1.98E-09 |
| VC2187 | flagellin FlaC | -3.43 | 6.86E-17 |
| VC2188 | flagellin core protein A | -2.81 | 4.79E-14 |
| VC2201 | chemotaxis protein methyltransferase CheR | -3.41 | 3.77E-21 |
| VC2202 | chemotaxis protein CheV | -3.22 | 3.30E-19 |
| VC2204 | negative regulator of flagellin synthesis FlgM,putative | -3.27 | 6.37E-12 |
| VC2205 | hypothetical protein | -3.62 | 3.54E-22 |
| VC2206 | conserved hypothetical protein | -3.95 | 1.39E-30 |
| VC2207 | hypothetical protein | -3.24 | 5.69E-12 |
| VC2210 | vibriobactin utilization protein ViuB | 2.05 | 4.89E-06 |
| VC2221 | hypothetical protein | -2.38 | 3.12E-08 |
| VC2241 | cytochrome c554 | -3.89 | 1.46E-15 |
| VC2264 | conserved hypothetical protein | -4.33 | 2.04E-31 |
| VC2285 | GGDEF family protein | -2.21 | 1.12E-07 |
| VC2293 | NADH:ubiquinone oxidoreductase, Natranslocating, gamma subunit | 2.11 | 9.58E-11 |
| VC2296 | bolA protein | -2.53 | 1.25E-24 |
| VC2314 | hypothetical protein | -4.50 | 5.31E-15 |
| VC2316 | N-acetylglutamate synthase | -3.46 | 1.01E-15 |
| VC2329 | 2,3,4,5-tetrahydropy ridine-2-carboxylateN-succinyl transferase | 2.25 | 2.56E-08 |
| VC2330 | conserved hypothetical protein | -2.20 | 1.43E-05 |
| VC2340 | conserved hypothetical protein | -7.15 | 1.20E-28 |
| VC2342 | elongation factor G | 3.07 | 3.75E-14 |
| VC2344 | hypothetical protein | -2.00 | 3.43E-09 |
| VC2352 | NupC family protein | 2.02 | 1.55E-07 |
| VC2356 | sodium/alanine symporter | 2.23 | 1.85E-18 |

| | | | |
|---|---|---|---|
| VC2357 | hypothetical protein | -2.10 | 8.02E-09 |
| VC2358 | hypothetical protein | -3.88 | 9.58E-26 |
| VC2360 | endonuclease IV | 2.77 | 5.76E-14 |
| VC2370 | sensory box/GGDEF family protein | -2.84 | 1.01E-12 |
| VC2378 | conserved hypothetical protein | -2.94 | 2.69E-32 |
| VC2383 | transcriptional regulator, LysR family | -2.15 | 5.43E-08 |
| VC2384 | conserved hypothetical protein | -4.04 | 2.11E-18 |
| VC2386 | conserved hypothetical protein | 2.79 | 7.79E-17 |
| VC2439 | methyl-accepting chemotaxis protein | -2.03 | 3.63E-09 |
| VC2454 | GGDEF family protein | -3.68 | 4.31E-24 |
| VC2455 | hypothetical protein | -5.04 | 3.72E-25 |
| VC2456 | hypothetical protein | -4.55 | 3.61E-33 |
| VC2462 | signal peptidase I | 2.22 | 1.08E-18 |
| VC2472 | conserved hypothetical protein | 2.09 | 6.46E-07 |
| VC2487 | conserved hypothetical protein | -2.06 | 1.09E-06 |
| VC2494 | hypothetical protein | -4.95 | 1.30E-27 |
| VC2507 | conserved hypothetical protein | -10.11 | 1.88E-76 |
| VC2508 | ornithine carbamoyltransferase | -5.98 | 1.08E-28 |
| VC2512 | conserved hypothetical protein | 4.11 | 1.56E-29 |
| VC2513 | 1-acyl-sn-glycerol-3 -phosphate acyltransferase | 2.18 | 7.12E-07 |
| VC2517 | conserved hypothetical protein | 2.20 | 1.97E-11 |
| VC2518 | conserved hypothetical protein | 2.03 | 3.38E-11 |
| VC2529 | RNA polymerase sigma-54 factor | -2.52 | 2.87E-14 |
| VC2530 | sigma-54 modulation protein, putative | -2.91 | 9.61E-10 |
| VC2533 | phosphocarrier protein NPr | -2.00 | 6.50E-04 |
| VC2545 | inorganic pyrophosphatase | 2.07 | 2.78E-05 |
| VC2568 | peptidyl-prolyl cis-trans isomerase, FKBP-type | 3.51 | 1.68E-23 |
| VC2571 | DNA-directed RNA polymerase, alpha subunit | 2.07 | 3.17E-07 |
| VC2598 | RNA methyltransferase, TrmH family | -2.64 | 7.60E-12 |
| VC2599 | ribonuclease R | -2.64 | 3.32E-13 |
| VC2601 | sodium-type flagellar protein MotX | -4.40 | 2.18E-25 |
| VC2617 | arginine/ornithine succinyltransferase,putative | 2.28 | 2.68E-10 |
| VC2622 | hypothetical protein | -5.42 | 1.00E-29 |
| VC2646 | phosphoenolpyruvate carboxylase | 2.13 | 1.20E-07 |
| VC2647 | conserved hypothetical protein | -2.13 | 5.09E-08 |
| VC2656 | fumarate reductase, flavoprotein subunit | -2.73 | 2.37E-12 |
| VC2657 | fumarate reductase, iron-sulfur protein | -5.09 | 6.43E-31 |
| VC2658 | fumarate reductase, 15 kDa hydrophobic protein | -5.05 | 3.13E-36 |
| VC2659 | fumarate reductase, 13 kDa hydrophobic protein | -3.92 | 6.47E-29 |
| VC2660 | elongation factor P | 2.54 | 6.84E-11 |
| VC2682 | met repressor | 2.06 | 6.34E-08 |
| VC2699 | C4-dicarboxylate transporter, anaerobic | -2.07 | 2.42E-08 |
| VC2702 | transcriptional regulator, LuxR family | -2.77 | 1.10E-19 |
| VC2704 | hypothetical protein | -12.57 | 2.56E-46 |
| VC2705 | sodium/solute symporter, putative | -13.90 | 3.15E-71 |
| VC2706 | conserved hypothetical protein | 4.01 | 5.94E-19 |
| VC2716 | conserved hypothetical protein | 3.04 | 1.46E-34 |
| VC2717 | hypothetical protein | -2.61 | 2.49E-15 |
| VC2726 | general secretion pathway protein K | -2.12 | 2.94E-08 |
| VC2727 | general secretion pathway protein J | -2.24 | 2.12E-08 |
| VC2738 | phosphoenolpyruvate carboxykinase | -2.19 | 2.33E-07 |
| VC2749 | nitrogen regulation protein NR(I) | -2.68 | 1.70E-19 |
| VC2750 | GGDEF family protein | -3.02 | 1.28E-16 |
| VC2761 | multidrug resistance protein | 2.38 | 5.49E-12 |
| VC2765 | ATP synthase F1, gamma subunit | 2.13 | 7.53E-12 |
| VC2766 | ATP synthase F1, alpha subunit | 2.09 | 4.76E-07 |
| VC2767 | ATP synthase F1, delta subunit | 2.05 | 8.55E-09 |
| VC2768 | ATP synthase F0, B subunit | 2.01 | 3.88E-09 |
| VC2775 | glucose inhibited division protein A | 2.00 | 3.11E-10 |
| VCA0005 | hypothetical protein | 2.70 | 5.26E-13 |
| VCA0006 | conserved hypothetical protein | 2.70 | 6.79E-19 |
| VCA0009 | hypothetical protein | -12.82 | 1.64E-24 |
| VCA0018 | vgrG protein | 3.18 | 1.83E-15 |
| VCA0019 | hypothetical protein | 3.45 | 1.22E-13 |
| VCA0020 | hypothetical protein | 3.03 | 2.93E-34 |
| VCA0026 | conserved hypothetical protein | 3.53 | 7.83E-23 |
| VCA0030 | hypothetical protein | -2.15 | 7.05E-09 |
| VCA0031 | methyl-accepting chemotaxis protein | -3.93 | 4.17E-23 |
| VCA0032 | hypothetical protein | -3.60 | 1.23E-23 |
| VCA0034 | conserved hypothetical protein | -3.90 | 1.12E-27 |
| VCA0037 | conserved hypothetical protein | 2.08 | 4.08E-09 |
| VCA0042 | hypothetical protein | -2.02 | 3.61E-08 |
| VCA0047 | conserved hypothetical protein | -2.26 | 8.65E-13 |
| VCA0049 | GGDEF family protein | -2.47 | 9.05E-14 |
| VCA0051 | hypothetical protein | -2.30 | 3.86E-12 |
| VCA0074 | GGDEF family protein | -3.34 | 8.30E-18 |
| VCA0075 | hypothetical protein | -2.48 | 2.55E-06 |
| VCA0078 | hypothetical protein | -2.37 | 3.77E-12 |
| VCA0088 | proton/glutamate symporter | 6.17 | 5.99E-84 |
| VCA0107 | conserved hypothetical protein | 6.09 | 1.17E-32 |
| VCA0108 | conserved hypothetical protein | 5.78 | 1.21E-45 |
| VCA0109 | hypothetical protein | 2.62 | 3.51E-10 |
| VCA0110 | hypothetical protein | 2.07 | 4.80E-07 |
| VCA0111 | hypothetical protein | 2.05 | 6.80E-07 |
| VCA0112 | hypothetical protein | 2.13 | 6.92E-09 |
| VCA0130 | ribose ABC transporter, periplasmicD-ribose-binding protein | -2.46 | 3.01E-10 |
| VCA0139 | hypothetical protein | 6.20 | 1.58E-38 |
| VCA0141 | C4-dicarboxylate transport sensor protein,putative | -2.10 | 3.10E-11 |
| VCA0144 | immunogenic protein | -2.24 | 5.35E-09 |
| VCA0146 | conserved hypothetical protein | -2.35 | 2.56E-05 |
| VCA0147 | transcriptional regulator, putative | -2.88 | 9.33E-09 |
| VCA0151 | oxidoreductase, putative | -2.26 | 4.48E-06 |
| VCA0152 | conserved hypothetical protein | -10.63 | 8.09E-52 |
| VCA0153 | conserved hypothetical protein | -9.45 | 4.16E-31 |
| VCA0154 | conserved hypothetical protein | -8.01 | 1.05E-34 |
| VCA0155 | NADH dehydrogenase, putative | -6.46 | 8.26E-46 |
| VCA0157 | NADH dehydrogenase, putative | -5.28 | 1.10E-38 |

| | | | |
|---|---|---|---|
| VCA0160 | tryptophan-specific transport protein | -2.45 | 1.04E-05 |
| VCA0184 | cold shock DNA-binding domain protein | 2.08 | 3.55E-08 |
| VCA0186 | hypothetical protein | -3.41 | 2.08E-28 |
| VCA0188 | hypothetical protein | -4.49 | 1.10E-17 |
| VCA0189 | response regulator | -2.78 | 1.37E-15 |
| VCA0191 | conserved hypothetical protein | -4.10 | 1.75E-27 |
| VCA0192 | D-lactate dehydrogenase | 3.57 | 1.07E-25 |
| VCA0195 | hypothetical protein | -3.77 | 8.80E-22 |
| VCA0205 | C4-dicarboxylate transporter, anaerobic | -3.06 | 2.89E-21 |
| VCA0210 | response regulator, putative | -7.27 | 5.10E-57 |
| VCA0211 | sensory box sensor histidine kinase | -6.60 | 1.85E-52 |
| VCA0217 | GGDEF family protein | -2.11 | 6.53E-07 |
| VCA0220 | hemolysin secretion protein HylB | -2.45 | 1.14E-14 |
| VCA0221 | lactonizing lipase | -2.36 | 3.43E-06 |
| VCA0229 | iron(III) ABC transporter, permease protein | 4.15 | 3.94E-07 |
| VCA0230 | iron(III) ABC transporter, ATP-binding protein | 5.12 | 1.96E-27 |
| VCA0241 | hexulose-6-phosphate isomerase SgbU, putative | -3.24 | 1.11E-05 |
| VCA0250 | alpha-amylase | -2.37 | 1.16E-08 |
| VCA0260 | hypothetical protein | -2.50 | 8.19E-10 |
| VCA0277 | glycine cleavage system H protein | 2.18 | 1.22E-09 |
| VCA0293 | hypothetical protein | -2.01 | 3.59E-07 |
| VCA0319 | conserved hypothetical protein | -2.31 | 2.38E-05 |
| VCA0344 | hypothetical protein | -2.34 | 1.94E-09 |
| VCA0363 | hypothetical protein | 2.15 | 1.01E-04 |
| VCA0369 | hypothetical protein | 6.69 | 3.62E-04 |
| VCA0375 | hypothetical protein | -2.01 | 2.45E-03 |
| VCA0428 | hypothetical protein | -2.30 | 2.13E-11 |
| VCA0490 | lipase, GDXG family | 2.01 | 8.59E-11 |
| VCA0518 | PTS system, fructose-specific IIA/FPR component | 6.47 | 1.19E-28 |
| VCA0519 | fructose repressor | 2.69 | 2.73E-10 |
| VCA0524 | conserved hypothetical protein | -2.22 | 6.10E-08 |
| VCA0534 | pseudogene; contains authentic frameshift mutation | -2.53 | 1.84E-08 |
| VCA0539 | conserved hypothetical protein | 2.02 | 1.76E-09 |
| VCA0544 | conserved hypothetical protein | -2.03 | 6.46E-11 |
| VCA0549 | phnA protein | 2.43 | 1.68E-19 |
| VCA0551 | hypothetical protein | -3.69 | 9.28E-42 |
| VCA0556 | hypothetical protein | 2.61 | 1.77E-06 |
| VCA0557 | GGDEF family protein | -2.73 | 4.08E-17 |
| VCA0558 | gamma-glutamyltransp eptidase, putative | -2.69 | 1.01E-13 |
| VCA0570 | Sui1 family protein | -2.46 | 7.75E-12 |
| VCA0574 | conserved hypothetical protein | -3.35 | 4.84E-23 |
| VCA0583 | hypothetical protein | -9.15 | 2.06E-41 |
| VCA0584 | glutathione S-transferase, putative | -2.01 | 1.03E-08 |
| VCA0588 | peptide ABC transporter, ATP-binding protein,putative | -2.47 | 2.37E-11 |
| VCA0590 | peptide ABC transporter, permease protein,putative | -2.61 | 6.00E-10 |
| VCA0591 | peptide ABC transporter, periplasmicpeptide-binding protein, putative | -2.27 | 1.89E-11 |
| VCA0593 | hypothetical protein | -4.90 | 2.30E-33 |
| VCA0603 | ABC transporter, periplasmic substrate-bindingprotein | -2.55 | 1.58E-09 |
| VCA0607 | regulator of nucleoside diphosphate kinase | 2.03 | 9.21E-07 |
| VCA0608 | conserved hypothetical protein | 2.31 | 1.51E-11 |
| VCA0610 | sigma cross-reacting protein 27A | -2.30 | 2.17E-11 |
| VCA0619 | hypothetical protein | -4.55 | 1.32E-39 |
| VCA0620 | thiosulfate sulfurtransferase SseA, putative | -2.15 | 2.89E-11 |
| VCA0638 | transporter, AcrB/D/F family | -2.70 | 1.95E-11 |
| VCA0639 | AcrA/AcrE family protein | -3.09 | 6.07E-21 |
| VCA0643 | conserved hypothetical protein | 2.32 | 9.44E-06 |
| VCA0646 | conserved hypothetical protein/hemolysin,putative | -5.04 | 6.55E-27 |
| VCA0652 | hypothetical protein | 3.72 | 3.45E-19 |
| VCA0658 | methyl-accepting chemotaxis protein | 2.17 | 6.52E-11 |
| VCA0659 | protein F-related protein | -3.90 | 9.31E-39 |
| VCA0666 | L-serine dehydratase 1 | -3.36 | 1.01E-20 |
| VCA0679 | periplasmic nitrate reductase, cytochrome c-typeprotein | -2.74 | 2.30E-07 |
| VCA0680 | periplasmic nitrate reductase, cytochrome c-typeprotein | -3.03 | 1.40E-08 |
| VCA0681 | conserved hypothetical protein | -6.55 | 2.37E-52 |
| VCA0682 | transcriptional regulator UhpA | -2.17 | 1.50E-07 |
| VCA0683 | sensor protein UhpB | -2.59 | 1.69E-12 |
| VCA0691 | acetoacetyl-CoA reductase | -2.13 | 2.34E-07 |
| VCA0692 | pseudogene with authentic frameshift mutation | 2.40 | 1.83E-07 |
| VCA0695 | hypothetical protein | -6.03 | 3.97E-25 |
| VCA0700 | chitodextrinase | -2.19 | 6.84E-07 |
| VCA0719 | sensor histide kinase | -6.32 | 1.54E-31 |
| VCA0720 | guanylate cyclase-related protein | -5.21 | 1.63E-31 |
| VCA0722 | hypothetical protein | -2.17 | 1.49E-05 |
| VCA0732 | conserved hypothetical protein | 2.61 | 2.77E-09 |
| VCA0734 | hypothetical protein | -2.52 | 6.06E-17 |
| VCA0736 | sensor histidine kinase LuxQ | -2.32 | 3.45E-09 |
| VCA0738 | hypothetical protein | -3.87 | 1.42E-23 |
| VCA0748 | anaerobic glycerol-3-phosphate dehydrogenase,subunit B | 11.86 | 1.88E-71 |
| VCA0752 | thioredoxin 2 | 3.20 | 8.66E-18 |
| VCA0766 | cytochrome c554 | -2.42 | 1.71E-06 |
| VCA0787 | hypothetical protein | -4.36 | 5.35E-04 |
| VCA0788 | DnaJ-related protein | -2.39 | 7.80E-12 |
| VCA0789 | conserved hypothetical protein | 3.41 | 2.09E-19 |
| VCA0798 | CbbY family protein | -4.07 | 2.72E-33 |
| VCA0803 | serine protease, putative | -13.51 | 1.07E-59 |
| VCA0806 | hypothetical protein | -2.11 | 1.22E-07 |
| VCA0811 | chitinase, putative | 3.04 | 5.12E-16 |
| VCA0834 | hypothetical protein | -3.35 | 1.79E-21 |
| VCA0843 | glyceraldehyde 3-phosphate dehydrogenase | 2.74 | 5.94E-09 |
| VCA0848 | GGDEF family protein | -4.30 | 7.84E-56 |
| VCA0862 | long-chain fatty acid transport protein | 3.33 | 2.79E-15 |
| VCA0863 | lipase, putative | 4.64 | 1.68E-33 |
| VCA0864 | methyl-accepting chemotaxis protein | -2.62 | 1.13E-11 |
| VCA0867 | outer membrane protein OmpW | -2.23 | 4.34E-06 |
| VCA0868 | hypothetical protein | -4.91 | 3.92E-36 |
| VCA0884 | hypothetical protein | -5.96 | 7.08E-44 |
| VCA0892 | hypothetical protein | -2.95 | 2.98E-14 |

| | | | |
|---|---|---|---|
| VCA0895 | chemotactic transducer-related protein | -4.14 | 2.76E-27 |
| VCA0897 | devB protein | 2.74 | 1.56E-13 |
| VCA0898 | 6-phosphogluconate dehydrogenase,decarboxylating | 3.18 | 2.40E-16 |
| VCA0900 | hypothetical protein | -3.09 | 1.02E-16 |
| VCA0902 | hypothetical protein | -2.08 | 7.54E-07 |
| VCA0906 | methyl-accepting chemotaxis protein | -8.00 | 8.96E-65 |
| VCA0920 | hypothetical protein | -4.50 | 4.76E-28 |
| VCA0923 | methyl-accepting chemotaxis protein | -6.54 | 3.56E-42 |
| VCA0931 | conserved hypothetical protein | -3.22 | 9.52E-18 |
| VCA0939 | sensory box/GGDEF family protein | -2.82 | 9.90E-16 |
| VCA0940 | transcriptional regulator, DeoR family | 2.19 | 1.36E-08 |
| VCA0960 | GGDEF family protein | -2.32 | 1.36E-11 |
| VCA0965 | GGDEF family protein | -5.71 | 1.42E-32 |
| VCA0966 | hypothetical protein | -3.14 | 8.78E-12 |
| VCA0978 | amino acid ABC transporter, periplasmic aminoacid-binding protein, putative | -5.61 | 8.54E-33 |
| VCA0979 | methyl-accepting chemotaxis protein | -4.70 | 3.75E-32 |
| VCA0981 | hypothetical protein | -4.69 | 1.17E-32 |
| VCA0983 | L-lactate permease, putative | 6.86 | 1.46E-58 |
| VCA0985 | oxidoreductase/iron- sulfur cluster-bindingprotein | 4.52 | 3.16E-25 |
| VCA0987 | phosphoenolpyruvate synthase | 2.37 | 4.00E-15 |
| VCA0997 | hypothetical protein | 2.00 | 4.68E-06 |
| VCA1001 | transcriptional regulator, AraC/XylS family | -2.65 | 2.48E-09 |
| VCA1003 | hypothetical protein | -2.31 | 2.70E-05 |
| VCA1008 | outer membrane protein, putative | -2.34 | 3.84E-04 |
| VCA1011 | glutaredoxin-related protein | -2.19 | 3.64E-07 |
| VCA1015 | Na+/H+ antiporter | -5.89 | 3.22E-53 |
| VCA1016 | hypothetical protein | -7.75 | 8.56E-78 |
| VCA1017 | methylated-DNA--prot ein-cysteineS-methyltransferas e | -6.04 | 7.41E-48 |
| VCA1033 | extracellular solute-binding protein, putative | -6.13 | 6.46E-29 |
| VCA1034 | methyl-accepting chemotaxis protein | -6.84 | 2.03E-47 |
| VCA1035 | hypothetical protein | 3.88 | 1.18E-17 |
| VCA1054 | conserved hypothetical protein | -5.07 | 5.59E-30 |
| VCA1055 | transcriptional regulator, LysR family | -2.11 | 2.66E-12 |
| VCA1056 | methyl-accepting chemotaxis protein | -6.37 | 8.71E-32 |
| VCA1066 | hypothetical protein | -2.05 | 1.57E-03 |
| VCA1073 | bifunctional proline dehydrogenase/pyrroline-5-carboxylate dehydrogenase | 4.47 | 5.15E-31 |
| VCA1079 | pyridoxamine 5`-phosphate oxidase | 2.12 | 1.38E-16 |
| VCA1087 | anti-sigma F factor antagonist, putative | -4.60 | 6.45E-26 |
| VCA1088 | methyl-accepting chemotaxis protein | -5.16 | 7.21E-42 |
| VCA1089 | cheB3 methylesterase | -8.12 | 4.37E-76 |
| VCA1090 | chemotaxis protein CheD, putative | -10.11 | 8.89E-64 |
| VCA1094 | purine-binding chemotaxis protein CheW | -8.18 | 3.39E-44 |
| VCA1095 | chemotaxis protein CheA | -8.00 | 1.43E-29 |
| VCA1096 | chemotaxis protein CheY | -8.00 | 1.26E-46 |
| VCA1097 | conserved hypothetical protein | -7.34 | 1.21E-52 |
| VCA1104 | phoP homolog | -2.28 | 1.13E-11 |
| VCA1105 | DNA-binding response regulator | -2.16 | 1.23E-07 |
| VCA1106 | hypothetical protein | -2.25 | 7.14E-09 |
| 16Sf | 16Sf | 3.47 | 8.14E-04 |
| 23Sh | 23Sh | 2.44 | 2.09E-11 |
| tRNA-Asp-4 | **tRNA-Asp-4** | 2.51 | 3.41E-14 |
| tRNA-Gly-3 | **tRNA-Gly-3** | 2.54 | 4.38E-11 |
| tRNA-Met-5 | **tRNA-Met-5** | 3.11 | 1.86E-05 |
| tRNA-Ser-1 | **tRNA-Ser-1** | 3.05 | 8.63E-06 |
| tRNA-Thr-3 | **tRNA-Thr-3** | 2.16 | 1.88E-06 |
| VC0007 | ribosomal protein L34 | 2.42 | 3.02E-10 |
| VC0030 | acetolactate synthase II, small subunit | 2.06 | 9.88E-05 |
| VC0123 | cyaY protein | 2.16 | 3.24E-09 |
| VC0218 | ribosomal protein L28 | 3.46 | 5.63E-13 |
| VC0219 | ribosomal protein L33 | 3.78 | 1.08E-34 |
| VC0321 | elongation factor Tu | 2.41 | 4.12E-09 |
| VC0322 | preprotein translocase, SecE subunit | 2.18 | 2.83E-08 |
| VC0323 | transcription antitermination protein NusG | 2.36 | 2.60E-13 |
| VC0324 | ribosomal protein L11, RplK | 3.34 | 1.85E-19 |
| VC0325 | ribosomal protein L1 | 2.92 | 2.16E-16 |
| VC0326 | ribosomal protein L10 | 3.35 | 1.26E-17 |
| VC0359 | ribosomal protein S12 | 3.21 | 5.80E-17 |
| VC0362 | elongation factor TU | 3.75 | 2.54E-17 |
| VC0366 | ribosomal protein S6 | 2.69 | 1.22E-17 |
| VC0368 | ribosomal protein S18 | 3.01 | 9.56E-20 |
| VC0369 | ribosomal protein L9 | 2.65 | 5.41E-16 |
| VC0374 | glucose-6-phosphate isomerase | 2.76 | 2.41E-16 |
| VC0435 | ribosomal protein L21 | 2.64 | 4.50E-14 |
| VC0436 | ribosomal protein L27 | 2.30 | 2.31E-13 |
| VC0441 | bis(5`-nucleosyl)-te traphosphatase | -2.03 | 7.58E-09 |
| VC0529 | conserved hypothetical protein | 2.48 | 2.48E-10 |
| VC0530 | conserved hypothetical protein | 2.11 | 4.35E-08 |
| VC0561 | ribosomal protein S16 | 4.06 | 2.45E-16 |
| VC0562 | 16S rRNA processing protein RimM | 3.45 | 6.68E-23 |
| VC0563 | tRNA (guanine-N1)-methyltransferase | 3.72 | 5.58E-30 |
| VC0564 | ribosomal protein L19 | 5.28 | 1.77E-46 |
| VC0569 | hypothetical protein | 2.42 | 6.19E-14 |
| VC0570 | ribosomal protein L13 | 2.42 | 4.09E-14 |
| VC0591 | pantoate--beta-alani ne ligase | 2.09 | 4.04E-11 |
| VC0592 | 3-methyl-2-oxobutano atehydroxymethyltransferase | 2.16 | 1.15E-07 |
| VC0634 | transcription elongation factor GreA | 2.09 | 1.00E-06 |
| VC0639 | phosphoglucomutase/p hosphomannomutase familyprotein MrsA | 2.15 | 5.52E-11 |
| VC0642 | N utilization substance protein A | 2.82 | 2.46E-18 |
| VC0646 | ribosomal protein S15 | 3.33 | 5.05E-19 |
| VC0650 | multidrug efflux pump VmrA | -2.42 | 5.01E-09 |
| VC0664 | lysyl-tRNA synthetase, heat inducible | 2.62 | 3.62E-15 |
| VC0679 | ribosomal protein S20 | 2.61 | 6.10E-15 |
| VC0718 | conserved hypothetical protein | 2.21 | 4.94E-08 |
| VC0739 | S-adenosylmethionine :tRNAribosyltransferase-isomer ase | 2.32 | 4.25E-10 |
| VC0877 | hypothetical protein | 2.57 | 1.72E-10 |
| VC0893 | chemotaxis protein PomB | -3.20 | 2.39E-24 |
| VC0905 | lipoprotein YaeC | 3.53 | 8.59E-28 |

| | | | |
|---|---|---|---|
| VC0907 | ABC transporter, ATP-binding protein | 2.40 | 2.26E-10 |
| VC0918 | UDP-N-acetyl-D-manno saminuronic aciddehydrogenase | -2.80 | 1.14E-10 |
| VC0941 | serine hydroxymethyltransferase | 2.64 | 2.38E-08 |
| VC0991 | asparagine synthetase B, glutamine-hydrolyzing | 3.76 | 5.26E-26 |
| VC1036 | methionyl-tRNA synthetase | 2.03 | 7.53E-09 |
| VC1092 | oligopeptide ABC transporter, permease protein | 2.77 | 4.12E-13 |
| VC1094 | oligopeptide ABC transporter, ATP-bindingprotein | 2.71 | 1.94E-17 |
| VC1143 | conserved hypothetical protein | -2.03 | 1.33E-08 |
| VC1144 | ATP-dependent Clp protease, ATP-binding subunitClpA | -3.05 | 4.43E-18 |
| VC1146 | glutaredoxin 1 | 4.37 | 8.24E-24 |
| VC1190 | phosphoribosylaminoi midazole-succinocarboxamidesyn thase, putative | -2.30 | 7.43E-09 |
| VC1221 | hypothetical protein | 3.20 | 3.79E-25 |
| VC1239 | cobinamide kinase/cobinamide phosphateguanylyltransferase | 2.12 | 3.71E-05 |
| VC1312 | alanine racemase, putative | -3.46 | 1.08E-15 |
| VC1627 | Na+/H+ antiporter protein | -2.66 | 3.14E-13 |
| VC1836 | tolB protein | 2.67 | 5.33E-14 |
| VC1914 | integration host factor, beta subunit | -2.88 | 2.33E-17 |
| VC1915 | ribosomal protein S1 | 2.26 | 2.20E-07 |
| VC1923 | trigger factor | 2.39 | 6.52E-10 |
| VC2020 | acyl carrier protein | 2.83 | 2.28E-18 |
| VC2021 | 3-oxoacyl-(acyl-carr ier-protein) reductase | 2.30 | 1.29E-11 |
| VC2025 | ribosomal protein L32 | 2.56 | 2.57E-25 |
| VC2066 | RNA polymerase sigma factor for flagellar operonFliA | -2.11 | 7.61E-10 |
| VC2068 | flagellar biosynthetic protein FlhF, putative | -2.08 | 1.16E-10 |
| VC2069 | flagellar biosynthetic protein FlhA | -2.05 | 7.65E-11 |
| VC2120 | flagellar biosynthetic protein FlhB | -2.42 | 9.06E-13 |
| VC2121 | flagellar biosynthetic protein FliR | -2.24 | 3.06E-09 |
| VC2122 | flagellar biosynthetic protein FliQ | -2.83 | 8.09E-16 |
| VC2123 | flagellar biosynthetic protein FliP | -2.98 | 3.26E-13 |
| VC2125 | flagellar motor switch protein FliN | -2.88 | 3.46E-13 |
| VC2126 | flagellar motor switch protein FliM | -3.14 | 5.10E-20 |
| VC2127 | flagellar protein FliL, putative | -3.22 | 7.43E-20 |
| VC2129 | flagellar protein FliJ, putative | -2.32 | 7.72E-10 |
| VC2130 | flagellum-specific ATP synthase FliI | -2.52 | 1.57E-14 |
| VC2131 | flagellar assembly protein FliH, putative | -2.22 | 3.24E-14 |
| VC2132 | flagellar motor switch protein FliG | -2.24 | 1.76E-10 |
| VC2138 | flagellar protein FliS | -3.20 | 3.31E-21 |
| VC2140 | flagellar hook-associated protein FliD | -4.12 | 2.50E-31 |
| VC2190 | flagellar hook-associated protein FlgL | -3.89 | 5.39E-14 |
| VC2191 | flagellar hook-associated protein FlgM | -2.80 | 1.98E-18 |
| VC2192 | flagellar protein FlgJ | -4.30 | 8.88E-30 |
| VC2193 | flagellar P-ring protein FlgI | -3.18 | 5.45E-21 |
| VC2194 | flagellar L-ring protein FlgH | -2.56 | 1.62E-13 |
| VC2195 | flagellar basal-body rod protein FlgG | -2.82 | 6.74E-17 |
| VC2196 | flagellar basal-body rod protein FlgF | -2.34 | 2.96E-12 |
| VC2197 | flagellar hook protein FlgE | -2.87 | 2.48E-19 |
| VC2198 | basal-body rod modification protein FlgD | -3.26 | 1.07E-21 |
| VC2199 | flagellar basal-body rod protein FlgC | -2.26 | 9.52E-11 |
| VC2214 | glutamyl-tRNA synthetase | 2.42 | 2.72E-11 |
| VC2223 | pseudouridine synthase family 1 protein | 2.18 | 5.49E-18 |
| VC2231 | oxidoreductase, acyl-CoA dehydrogenase family | -2.67 | 7.45E-11 |
| VC2234 | ribonuclease HI | -2.24 | 2.75E-08 |
| VC2257 | ribosome recycling factor | 2.21 | 7.43E-11 |
| VC2258 | uridylate kinase | 2.31 | 9.35E-14 |
| VC2259 | elongation factor Ts | 4.64 | 2.08E-48 |
| VC2260 | ribosomal protein S2 | 3.46 | 1.78E-19 |
| VC2400 | UDP-N-acetylmuramate --alanine ligase | 2.28 | 1.46E-13 |
| VC2414 | pyruvate dehydrogenase, E1 component | 2.54 | 1.97E-14 |
| VC2485 | transcriptional regulator, LysR family | 2.19 | 6.39E-16 |
| VC2503 | valyl-tRNA synthetase | 2.42 | 2.23E-07 |
| VC2570 | ribosomal protein L17 | 2.56 | 3.49E-08 |
| VC2572 | ribosomal protein S4 | 2.25 | 4.04E-13 |
| VC2578 | ribosomal protein L30 | 2.42 | 3.53E-07 |
| VC2579 | ribosomal protein S5 | 2.30 | 1.14E-22 |
| VC2580 | ribosomal protein L18 | 2.50 | 6.49E-13 |
| VC2581 | ribosomal protein L6 | 2.50 | 1.73E-12 |
| VC2582 | ribosomal protein S8 | 2.84 | 1.62E-15 |
| VC2583 | ribosomal protein S14 | 2.56 | 5.09E-14 |
| VC2584 | ribosomal protein L5 | 2.35 | 6.74E-13 |
| VC2585 | ribosomal protein L24 | 2.55 | 4.40E-15 |
| VC2586 | ribosomal protein L14 | 2.46 | 5.91E-14 |
| VC2587 | ribosomal protein S17 | 2.58 | 5.97E-14 |
| VC2588 | ribosomal protein L29 | 2.48 | 9.54E-08 |
| VC2589 | ribosomal protein L16 | 2.23 | 4.40E-21 |
| VC2590 | ribosomal protein S3 | 2.04 | 2.81E-07 |
| VC2591 | ribosomal protein L22 | 2.73 | 2.30E-14 |
| VC2592 | ribosomal protein S19 | 2.61 | 1.11E-14 |
| VC2593 | ribosomal protein L2 | 3.30 | 3.93E-17 |
| VC2594 | ribosomal protein L23 | 4.00 | 1.50E-15 |
| VC2595 | ribosomal protein L4 | 3.39 | 1.20E-23 |
| VC2596 | ribosomal protein L3 | 3.03 | 1.44E-20 |
| VC2597 | ribosomal protein S10 | 3.01 | 2.44E-17 |
| VC2618 | acetylornithine aminotransferase | 2.75 | 5.24E-14 |
| VC2663 | molecular chaperone groEL_1 | 3.09 | 4.53E-16 |
| VC2664 | chaperonin, 60 Kd subunit, groES_1 | 3.41 | 1.78E-12 |
| VC2670 | triosephosphate isomerase | 3.45 | 2.23E-24 |
| VC2679 | ribosomal protein L31 | 5.91 | 2.02E-49 |
| VC2691 | periplasmic protein cpxP, putative | 2.80 | 1.89E-10 |
| VC2715 | transcription elongation factor GreB | 2.28 | 1.54E-06 |
| VC2748 | nitrogen regulation protein | -2.39 | 6.91E-10 |
| VC2762 | UDP-N-acetylglucosam ine pyrophosphorylase | 3.19 | 3.81E-21 |
| VC2774 | glucose inhibited division protein B | 2.43 | 1.98E-12 |
| VCA0014 | 4-alpha-glucanotrans ferase | -2.80 | 3.79E-15 |
| VCA0149 | hypothetical protein | 2.53 | 4.83E-09 |
| VCA0246 | SgaT protein | -2.53 | 1.97E-06 |
| VCA0289 | ribosomal protein L35 | 2.89 | 1.35E-16 |
| VCA0290 | ribosomal protein L20 | 2.91 | 2.33E-17 |

| | | | |
|---|---|---|---|
| VCA0512 | anaerobic ribonucleoside-triphosphate reductaseactivating protein | -2.75 | 4.13E-14 |
| VCA0563 | NAD(P) transhydrogenase, alpha subunit | 3.43 | 1.11E-12 |
| VCA0564 | NAD(P) transhydrogenase, beta subunit | 3.74 | 1.21E-27 |
| VCA0698 | hypothetical protein | -7.35 | 8.18E-50 |
| VCA0747 | anaerobic glycerol-3-phosphate dehydrogenase,subunit A | 21.97 | 2.79E-119 |
| VCA0749 | anaerobic glycerol-3-phosphate dehydrogenase,subunit C | 11.92 | 4.52E-61 |
| VCA0757 | arginine ABC transporter, permease protein | -2.57 | 1.04E-13 |
| VCA0760 | arginine ABC transporter, ATP-binding protein | -3.49 | 3.83E-19 |
| VCA0819 | chaperonin, 10 Kd subunit | 3.41 | 2.70E-15 |
| VCA0820 | chaperonin, 60 Kd subunit | 2.40 | 3.16E-11 |
| VCA0870 | D-alanyl-D-alanine endopeptidase | -2.04 | 1.54E-07 |
| VCA0880 | hypothetical protein | -5.95 | 4.13E-54 |
| VCA0881 | hypothetical protein | -4.84 | 2.55E-42 |
| VCA0882 | hypothetical protein | -4.92 | 1.20E-35 |
| VCA0883 | hypothetical protein | -2.02 | 8.81E-09 |
| VCA0943 | maltose ABC transporter, permease protein | -2.32 | 1.30E-11 |
| VCA0945 | maltose ABC transporter, periplasmicmaltose-binding protein | -3.16 | 4.64E-14 |
| VCA0984 | L-lactate dehydrogenase | 11.24 | 1.86E-97 |
| VCA1060 | 3,4-dihydroxy-2-buta none 4-phosphate synthase | 3.70 | 1.25E-20 |

| 190 differentially-expressed genes in AKI-grown Δ*rpoE* cells, independent of Δ*ompU* , compared to WT C6706 | | | |
|---|---|---|---|
| **Locus Tag** | **Function** | **FC in Δ*rpoE*** | ***p*-value** |
| 16Sa | 16Sa | -20.22 | 1.48E-28 |
| 16Sb | 16Sb | -20.79 | 6.49E-32 |
| 16Sc | 16Sc | -13.13 | 1.37E-22 |
| 16Sd | 16Sd | -2.44 | 8.07E-04 |
| 16Se | 16Se | -10.18 | 1.19E-16 |
| 16Sg | 16Sg | -34.94 | 2.73E-45 |
| 16Sh | 16Sh | -6.03 | 3.87E-08 |
| 23Sa | 23Sa | -7.43 | 2.45E-15 |
| 23Sb | 23Sb | 3.68 | 1.05E-07 |
| 23Sc | 23Sc | -4.56 | 5.27E-06 |
| 23Sd | 23Sd | -7.85 | 8.81E-17 |
| 23Se | 23Se | -85.13 | 2.53E-65 |
| 23Sf | 23Sf | -4.10 | 2.62E-08 |
| 23Sg | 23Sg | -24.58 | 4.35E-43 |
| 5Sa | 5Sa | -79.61 | 6.43E-47 |
| 5Sb | 5Sb | -156.53 | 3.58E-15 |
| 5Se | 5Se | -207.22 | 2.11E-16 |
| 5Sf | 5Sf | ######## | 6.29E-08 |
| 5Sh | 5Sh | -159.57 | 2.83E-50 |
| tRNA-Ala-3 | tRNA-Ala-3 | 5.00 | 4.53E-05 |
| tRNA-Ala-5 | tRNA-Ala-5 | 10.78 | 5.22E-14 |
| tRNA-Asn-3 | tRNA-Asn-3 | 2.97 | 5.03E-05 |
| tRNA-Gly-5 | tRNA-Gly-5 | -5.09 | 1.64E-07 |
| tRNA-Ile-2 | tRNA-Ile-2 | 6.55 | 3.83E-09 |
| tRNA-Met-8 | tRNA-Met-8 | -3.60 | 1.31E-04 |
| tRNA-Val-4 | tRNA-Val-4 | 3.43 | 1.34E-05 |
| VC0005 | conserved hypothetical protein | -2.39 | 4.60E-04 |
| VC0021 | glycyl-tRNA synthetase, alpha chain; glyQ | -2.35 | 7.69E-07 |
| VC0026 | zinc-binding alcohol dehydrogenase | -2.21 | 9.13E-05 |
| VC0027 | threonine dehydratase | -2.37 | 4.51E-06 |
| VC0034 | thiol:disulfide interchange protein | -2.92 | 2.68E-11 |
| VC0038 | hypothetical protein | 2.27 | 5.13E-04 |
| VC0127 | conserved hypothetical protein | -2.17 | 2.79E-04 |
| VC0161 | transcriptional activator IlvY | -2.07 | 2.01E-04 |
| VC0171 | peptide ABC transporter, periplasmicpeptide-binding protein | 2.12 | 7.27E-04 |
| VC0175 | deoxycytidylate deaminase-related protein | -2.56 | 1.64E-08 |
| VC0176 | transcriptional regulator, putative | -2.68 | 2.54E-05 |
| VC0178 | patatin-related protein | -2.30 | 8.34E-06 |
| VC0181 | conserved hypothetical protein | -3.09 | 5.22E-10 |
| VC0184 | hypothetical protein | -3.24 | 5.26E-07 |
| VC0202 | iron(III) ABC transporter, periplasmiciron-compound-bindi ng protein | 2.79 | 2.55E-04 |
| VC0204 | conserved hypothetical protein | -2.44 | 2.38E-04 |
| VC0217 | DNA repair protein RadC | -2.12 | 1.13E-03 |
| VC0274 | hypothetical protein | -2.50 | 5.80E-06 |
| VC0286 | gluconate permease, putative | -2.95 | 4.46E-09 |
| VC0353 | conserved hypothetical protein | -2.39 | 3.69E-07 |
| VC0411 | MSHA pilin protein MshD | -2.23 | 2.15E-04 |
| VC0443 | dimethyladenosine transferase | -2.09 | 4.40E-06 |
| VC0490 | conserved hypothetical protein | -2.46 | 6.39E-05 |
| VC0491 | hypothetical protein | -3.01 | 7.96E-11 |
| VC0492 | hypothetical protein | -2.38 | 1.23E-04 |
| VC0493 | hypothetical protein | -2.76 | 2.83E-09 |
| VC0566 | protease DO | -2.85 | 4.15E-10 |
| VC0583 | quorum sensing regulator HapR | 2.43 | 7.87E-04 |
| VC0676 | nptA protein | 8.30 | 1.29E-11 |
| VC0708 | conserved hypothetical protein | -2.47 | 7.94E-08 |
| VC0720 | histidine protein kinase PhoR | 2.51 | 4.55E-04 |
| VC0743 | protein-export membrane protein SecD | -2.02 | 3.16E-05 |
| VC0761 | conserved hypothetical protein | 2.10 | 3.80E-04 |
| VC0770 | conserved hypothetical protein | -2.37 | 9.06E-04 |
| VC0774 | vibriobactin-specifi c2,3-dihydro-2,3-dihydroxybenz oate dehydrogenase | 6.91 | 1.15E-06 |
| VC0775 | vibriobactin synthesis protein, putative | 6.03 | 5.29E-07 |
| VC0776 | ferric vibriobactin ABC transporter, periplasmicferric vibriobactin-binding protein | 3.44 | 5.97E-06 |
| VC0777 | ferric vibriobactin ABC transporter, permeaseprotein | 3.76 | 2.75E-05 |
| VC0811 | hypothetical protein | -2.18 | 4.89E-05 |
| VC0823 | hypothetical protein | -2.11 | 2.69E-05 |
| VC0827 | toxin co-regulated pilus biosynthesis protein H | 2.99 | 2.34E-09 |
| VC0829 | toxin co-regulated pilus biosynthesis protein B | 3.00 | 5.06E-06 |
| VC0830 | toxin co-regulated pilus biosynthesis protein Q | 2.96 | 2.15E-05 |
| VC0831 | toxin co-regulated pilus biosynthesis outermembrane protein C | 3.09 | 1.96E-06 |
| VC0832 | toxin co-regulated pilus biosynthesis protein R | 2.73 | 3.55E-04 |
| VC0862 | hypothetical protein | -3.01 | 6.38E-06 |
| VC0869 | phosphoribosylformyl glycinamidine synthase | 2.28 | 1.55E-06 |
| VC0916 | phosphotyrosine protein phosphatase | 6.69 | 4.30E-04 |

| | | | |
|---|---|---|---|
| VC0947 | D-alanyl-D-alanine carboxypeptidase | -2.16 | 2.84E-05 |
| VC1051 | hypothetical protein | 2.18 | 1.59E-04 |
| VC1059 | oxidoreductase, short-chaindehydrogenase/reduc tase family | -2.03 | 7.75E-05 |
| VC1184 | NifS-related protein | 2.10 | 6.98E-06 |
| VC1186 | sanA protein | -2.35 | 7.78E-07 |
| VC1188 | malate oxidoreductase | -7.83 | 1.08E-25 |
| VC1205 | imidazolonepropionas e | -2.38 | 6.54E-05 |
| VC1206 | histidine utilization repressor | -2.90 | 7.98E-07 |
| VC1255 | ribonucleoside-dipho sphate reductase, betasubunit | 2.44 | 1.61E-05 |
| VC1267 | hypothetical protein | 2.49 | 3.28E-06 |
| VC1307 | pseudogene with authentic frameshift mutation | 2.01 | 4.61E-04 |
| VC1319 | sensor histidine kinase | -2.33 | 2.41E-05 |
| VC1329 | opacity protein-related protein | -3.65 | 5.91E-07 |
| VC1332 | conserved hypothetical protein | 2.07 | 5.40E-05 |
| VC1333 | hypothetical protein | 3.60 | 2.35E-14 |
| VC1334 | conserved hypothetical protein | 3.10 | 6.54E-13 |
| VC1336 | carboxyphosphonoenol pyruvate phosphonomutase | 2.55 | 1.77E-04 |
| VC1337 | methylcitrate synthase | 2.97 | 2.18E-06 |
| VC1338 | aconitate hydratase 1 | 2.27 | 1.85E-05 |
| VC1341 | acetyltransferase, putative | 2.04 | 5.13E-05 |
| VC1344 | 4-hydroxyphenylpyruv ate dioxygenase | 3.22 | 3.51E-12 |
| VC1345 | oxidoreductase, putative | 3.07 | 9.30E-11 |
| VC1346 | conserved hypothetical protein | 3.47 | 1.91E-11 |
| VC1367 | GGDEF family protein | -2.08 | 6.19E-06 |
| VC1449 | hypothetical protein | 2.47 | 3.33E-04 |
| VC1463 | RstA2 protein | -2.33 | 1.79E-04 |
| VC1470 | XRE family transcriptional regulator | -3.28 | 2.89E-06 |
| VC1524 | ABC transporter, permease protein | -2.16 | 3.64E-04 |
| VC1543 | hypothetical protein | 2.48 | 4.94E-05 |
| VC1544 | tonB2 protein | 3.29 | 2.20E-06 |
| VC1556 | conserved hypothetical protein | 2.12 | 3.26E-04 |
| VC1566 | conserved hypothetical protein | -2.21 | 7.62E-04 |
| VC1567 | conserved hypothetical protein | -2.46 | 1.18E-04 |
| VC1568 | ABC transporter, ATP-binding protein | -3.11 | 2.09E-06 |
| VC1583 | superoxide dismutase, Cu-Zn | 2.15 | 1.13E-03 |
| VC1587 | conserved hypothetical protein | 16.79 | 4.96E-06 |
| VC1598 | ABC transporter, periplasmic substrate-bindingprotein-relat ed protein | -2.12 | 2.87E-05 |
| VC1610 | hypothetical protein | 2.02 | 1.13E-03 |
| VC1678 | phage shock protein A | 11.08 | 1.01E-15 |
| VC1741 | transcriptional regulator, TetR family | 2.48 | 1.79E-06 |
| VC1764 | hypothetical protein | -2.20 | 2.64E-06 |
| VC1777 | conserved hypothetical protein | -2.39 | 1.07E-07 |
| VC1781 | conserved hypothetical protein | -2.96 | 4.34E-06 |
| VC1819 | aldehyde dehydrogenase | -6.43 | 1.31E-18 |
| VC1871 | conserved hypothetical protein | -2.43 | 1.83E-04 |
| VC1972 | o-succinylbenzoate-C oA synthase | -2.14 | 3.89E-04 |
| VC1973 | naphthoate synthase | -2.37 | 6.26E-06 |
| VC1974 | conserved hypothetical protein | -2.77 | 4.67E-06 |
| VC2019 | 3-oxoacyl-(acyl-carr ier-protein) synthase II | -2.10 | 1.81E-05 |
| VC2032 | hypothetical protein | -2.16 | 6.39E-04 |
| VC2076 | hypothetical protein | 4.36 | 1.66E-12 |
| VC2077 | ferrous iron transport protein B | 2.69 | 1.28E-07 |
| VC2156 | lipoprotein-34 NlpB | -2.02 | 3.93E-04 |
| VC2209 | nonribosomal peptide synthetase VibF | 2.44 | 5.37E-04 |
| VC2275 | transcriptional regulator Crl | -2.73 | 9.66E-06 |
| VC2371 | conserved hypothetical protein | -2.42 | 2.93E-04 |
| VC2510 | aspartate carbamoyltransferase, catalyticsubunit | 3.05 | 1.74E-10 |
| VC2511 | aspartate carbamoyltransferase, regulatorysubunit | 3.16 | 1.60E-09 |
| VC2636 | transcriptional regulator, LysR family | -2.85 | 1.96E-08 |
| VC2652 | conserved hypothetical protein | -3.13 | 2.84E-04 |
| VC2653 | protein-transport protein SecB | -2.85 | 3.89E-06 |
| VC2711 | ATP-dependent DNA helicase RecG | -2.48 | 9.17E-09 |
| VC2720 | conserved hypothetical protein | -2.10 | 4.43E-05 |
| VCA0004 | hypothetical protein | 2.19 | 2.76E-04 |
| VCA0033 | hypothetical protein | 2.05 | 3.38E-04 |
| VCA0059 | major outer membrane lipoprotein | 3.25 | 3.45E-06 |
| VCA0070 | phosphate ABC transporter, periplasmicphosphate-binding protein | 21.64 | 2.45E-13 |
| VCA0071 | phosphate ABC transporter, permease protein | 8.84 | 6.71E-08 |
| VCA0072 | phosphate ABC transporter, permease protein | 12.70 | 1.17E-08 |
| VCA0073 | phosphate ABC transporter, ATP-binding protein | 8.42 | 1.24E-05 |
| VCA0106 | hypothetical protein | -2.13 | 8.16E-04 |
| VCA0119 | hypothetical protein | -2.43 | 1.04E-03 |
| VCA0140 | spindolin-related protein | 2.65 | 7.86E-04 |
| VCA0165 | GGDEF family protein | -2.49 | 9.55E-09 |
| VCA0166 | cold shock transcriptional regulator CspA | -2.26 | 1.01E-03 |
| VCA0168 | hypothetical protein | -2.87 | 1.47E-07 |
| VCA0172 | conserved hypothetical protein | -2.04 | 1.70E-04 |
| VCA0175 | MoxR-related protein | -4.72 | 2.14E-14 |
| VCA0202 | IS1004 transposase | -28.66 | 8.81E-06 |
| VCA0215 | hypothetical protein | 2.91 | 2.68E-09 |
| VCA0256 | transcriptional regulator | 2.07 | 8.91E-04 |
| VCA0281 | integrase, putative | -2.34 | 4.59E-06 |
| VCA0283 | hypothetical protein | -3.54 | 8.79E-12 |
| VCA0284 | hypothetical protein | -2.68 | 2.77E-05 |
| VCA0306 | hypothetical protein | -2.12 | 4.62E-04 |
| VCA0336 | hypothetical protein | -6.31 | 2.11E-12 |
| VCA0337 | microcin immunity protein MccF | -2.07 | 7.72E-05 |
| VCA0340 | hypothetical protein | -2.36 | 1.29E-05 |
| VCA0354 | hypothetical protein | -2.14 | 4.89E-05 |
| VCA0355 | conserved hypothetical protein | -2.10 | 5.90E-04 |
| VCA0468 | hypothetical protein | 2.03 | 1.62E-04 |
| VCA0486 | hypothetical protein | 3.49 | 8.38E-09 |
| VCA0526 | conserved hypothetical protein | -2.10 | 1.36E-04 |
| VCA0532 | DNA-binding response regulator | 2.07 | 1.15E-03 |
| VCA0533 | tatA protein | 2.51 | 4.92E-06 |
| VCA0546 | conserved hypothetical protein | 2.31 | 2.72E-04 |
| VCA0547 | hypothetical protein | 2.02 | 8.18E-05 |

| | | | |
|---|---|---|---|
| VCA0635 | transcriptional regulator, LysR family | 2.19 | 7.00E-04 |
| VCA0676 | iron-sulfur cluster-binding protein NapF | 4.73 | 3.15E-04 |
| VCA0677 | napD protein | 4.11 | 2.40E-04 |
| VCA0686 | iron(III) ABC transporter, permease protein | -2.98 | 1.11E-05 |
| VCA0704 | phosphoglycerate transport systemtranscriptional regulatory protein PgtA | 2.31 | 2.19E-06 |
| VCA0707 | regulatory protein UhpC, putative | 17.03 | 1.60E-27 |
| VCA0708 | pyruvate kinase II | 4.67 | 7.48E-08 |
| VCA0831 | hypothetical protein | 5.11 | 3.91E-10 |
| VCA0849 | hypothetical protein | 2.85 | 4.33E-09 |
| VCA0874 | hypothetical protein | -4.09 | 4.35E-06 |
| VCA0887 | pseudogene with authentic frameshift mutation | 3.14 | 2.39E-06 |
| VCA0888 | transcriptional regulator, LuxR family | 2.24 | 1.39E-04 |
| VCA0933 | cold shock domain family protein | -7.03 | 3.55E-14 |
| VCA0951 | conserved hypothetical protein | -21.68 | 1.60E-20 |
| VCA0952 | transcriptional regulator, LuxR family | 4.02 | 1.03E-05 |
| VCA0994 | hypothetical protein | 2.57 | 1.41E-08 |
| VCA1021 | conserved hypothetical protein | 2.46 | 1.50E-07 |
| VCA1037 | amino acid ABC transporter, ATP-binding protein | -2.03 | 2.04E-04 |
| VCr025 | 5S ribosomal RNA | -567.55 | 5.36E-57 |

158 differentially-expressed genes common to both Δ*ompU* and Δ*rpoE*, compared to AKI-grown WT cells

| Name | Function | FC Δ*rpoE* | *p*-value | FC Δ*ompU* | *p*-value |
|---|---|---|---|---|---|
| 5Sc | 5Sc | -3453.75 | 2.04E-98 | -2.04 | 4.94E-09 |
| 5Sd | 5Sd | -4720.44 | 6.75E-88 | -2.04 | 5.09E-09 |
| 5Sg | 5Sg | -17365.49 | 5.77E-06 | -142.21 | 3.27E-110 |
| tRNA-Ala-2 | tRNA-Ala-2 | 10.50 | 1.71E-10 | 3.09 | 2.68E-06 |
| tRNA-Lys-2 | tRNA-Lys-2 | 2.88 | 3.60E-04 | 2.01 | 9.21E-05 |
| tRNA-Tyr-3 | tRNA-Tyr-3 | 3.25 | 7.15E-05 | 4.39 | 4.74E-12 |
| VC0015a | hypothetical protein | 2.30 | 7.86E-07 | -2.15 | 1.01E-09 |
| VC0017 | hypothetical protein | 2.68 | 1.08E-04 | -2.94 | 2.18E-05 |
| VC0040 | hemolysin, putative | -2.04 | 9.12E-06 | -2.19 | 4.65E-18 |
| VC0069 | multidrug resistance protein, putative | 2.45 | 1.16E-06 | -2.85 | 7.90E-14 |
| VC0075 | MadN protein | -2.20 | 1.63E-06 | -2.56 | 3.15E-19 |
| VC0107 | hypothetical protein | 15.91 | 7.64E-14 | 2.53 | 2.05E-10 |
| VC0280 | cadaverine/lysine antiporter CadB, putative | -2.98 | 7.28E-05 | -2.69 | 5.87E-05 |
| VC0287 | thermoresistant gluconokinase | -2.49 | 2.38E-04 | 2.21 | 3.49E-09 |
| VC0299 | DNA polymerase III, epsilon subunit, putative | 2.47 | 7.66E-05 | -5.44 | 2.00E-23 |
| VC0300 | conserved hypothetical protein | 2.76 | 1.60E-04 | -7.61 | 1.06E-35 |
| VC0327 | ribosomal protein L7/L12 | -2.01 | 5.82E-05 | 3.24 | 1.00E-23 |
| VC0364 | bacterioferritin-ass ociated ferredoxin | 11.80 | 6.48E-24 | 3.52 | 2.57E-15 |
| VC0379a | phage shock protein G | 5.22 | 2.49E-07 | -2.10 | 1.38E-06 |
| VC0553 | conserved hypothetical protein | -2.00 | 8.07E-04 | -2.11 | 1.63E-06 |
| VC0608 | iron(III) ABC transporter, periplasmiciron-compound-bindi ng protein | 5.27 | 2.72E-12 | 9.28 | 2.55E-80 |
| VC0633 | outer membrane protein OmpU | -4.62 | 2.06E-07 | -3191116.29 | 4.23E-14 |
| VC0734 | malate synthase A | 2.23 | 6.39E-04 | -4.94 | 1.71E-22 |
| VC0863 | conserved hypothetical protein | -2.94 | 5.76E-06 | -2.27 | 3.19E-08 |
| VC0930 | hemolysin-related protein | 3.17 | 8.42E-08 | 2.66 | 1.16E-15 |
| VC0945 | conserved hypothetical protein | 3.19 | 1.38E-05 | 2.26 | 4.57E-08 |
| VC1032 | zinc/cadmium/mercury/lead-transporting ATPase | 3.67 | 1.28E-04 | -3.26 | 7.57E-06 |
| VC1061 | cysteine synthase/cystathionine beta-synthasefamily protein | -2.73 | 1.04E-05 | -4.04 | 2.31E-14 |
| VC1116 | hypothetical protein | 3.65 | 9.06E-12 | -5.03 | 5.97E-23 |
| VC1187 | hypothetical protein | -4.48 | 6.63E-10 | -2.04 | 3.98E-05 |
| VC1207 | hypothetical protein | 2.00 | 8.97E-04 | -4.68 | 8.53E-23 |
| VC1222 | integration host factor, alpha subunit | 2.93 | 4.67E-05 | -3.69 | 3.05E-13 |
| VC1264 | iron-regulated protein A, putative | 3.80 | 1.72E-14 | 2.58 | 4.53E-24 |
| VC1265 | hypothetical protein | 2.70 | 9.29E-06 | 2.02 | 5.96E-07 |
| VC1266 | hypothetical protein | 4.26 | 3.08E-08 | 2.00 | 1.98E-07 |
| VC1280 | hypothetical protein | -2.46 | 2.23E-06 | 2.12 | 4.52E-08 |
| VC1325 | galactoside ABC transporter, periplasmicD-galactose/D-gluco se-binding protein | -5.61 | 2.10E-10 | -5.16 | 4.85E-31 |
| VC1327 | galactoside ABC transporter, ATP-bindingprotein | -6.83 | 5.27E-24 | -3.55 | 2.73E-23 |
| VC1328 | galactoside ABC transporter, permease protein | -6.33 | 2.28E-28 | -3.14 | 6.08E-19 |
| VC1339 | conserved hypothetical protein | 2.11 | 3.10E-05 | -2.13 | 1.39E-06 |
| VC1355 | acylphosphatase | 2.04 | 9.87E-05 | -2.92 | 3.76E-14 |
| VC1370 | GGDEF family protein | 2.06 | 4.56E-05 | -7.21 | 1.61E-45 |
| VC1401 | protein-glutamate methylesterase CheB | 2.52 | 1.05E-05 | -5.28 | 3.40E-31 |
| VC1402 | purine-binding chemotaxis protein Chew,putative | 2.25 | 9.75E-04 | -4.64 | 3.77E-19 |
| VC1415 | hcp protein | -4.56 | 5.20E-09 | 9.53 | 1.25E-48 |
| VC1456 | cholera enterotoxin, B subunit | 4.05 | 3.30E-06 | -3.08 | 3.18E-03 |
| VC1457 | cholera enterotoxin, A subunit | 4.12 | 8.65E-06 | -3.83 | 3.52E-04 |
| VC1484 | ribosome modulation factor | 3.22 | 3.72E-06 | -5.91 | 2.99E-22 |
| VC1495 | hypothetical protein | 2.44 | 2.92E-04 | -2.82 | 5.53E-06 |
| VC1509 | nicotinatemononucleo tide:5,6-dimethylbenzimidazole phosphoribosyltransferase | 2.44 | 9.97E-04 | -3.76 | 4.54E-18 |
| VC1545 | TonB system transport protein ExbD2 | 5.63 | 8.68E-10 | 2.20 | 3.41E-05 |
| VC1546 | TonB system transport protein ExbB2 | 4.13 | 9.34E-08 | 4.04 | 1.39E-18 |
| VC1547 | biopolymer transport protein ExbB-relatedprotein | 4.16 | 5.15E-08 | 5.45 | 4.20E-31 |
| VC1548 | hypothetical protein | 2.60 | 2.69E-04 | 3.82 | 1.92E-17 |
| VC1560 | catalase/peroxidase | 2.78 | 6.31E-04 | -5.03 | 2.90E-29 |
| VC1563 | conserved hypothetical protein | -3.77 | 2.54E-08 | 2.07 | 1.41E-06 |
| VC1565 | outer membrane protein TolC, putative | -4.17 | 1.48E-10 | 2.49 | 1.78E-10 |
| VC1593 | GGDEF family protein | -4.81 | 3.96E-15 | -2.87 | 3.16E-17 |
| VC1594 | aldose 1-epimerase | -19.71 | 6.47E-70 | -3.72 | 1.55E-26 |
| VC1595 | galactokinase | -26.28 | 1.39E-85 | -3.60 | 6.83E-26 |
| VC1596 | galactose-1-phosphat e uridylyltransferase | -33.92 | 6.31E-60 | -2.86 | 2.44E-17 |
| VC1605a | hypothetical protein | 2.38 | 4.83E-04 | -2.80 | 3.18E-18 |
| VC1618 | multidrug resistance protein, putative | 2.45 | 1.22E-07 | -2.80 | 2.24E-05 |
| VC1640 | ribosomal protein L25 | -4.96 | 4.61E-19 | 3.64 | 6.24E-25 |
| VC1655 | magnesium transporter | 9.75 | 1.08E-10 | 3.05 | 4.82E-18 |
| VC1661 | hypothetical protein | 2.18 | 7.18E-06 | -2.80 | 2.43E-12 |
| VC1676 | phage shock protein C | 6.27 | 1.52E-11 | -2.31 | 1.67E-08 |
| VC1677 | phage shock protein B | 8.74 | 6.09E-17 | -2.02 | 1.90E-06 |
| VC1706 | transcriptional activator MetR | 2.11 | 6.34E-05 | -2.15 | 9.18E-07 |
| VC1707 | hypothetical protein | 3.53 | 2.32E-06 | -5.78 | 6.11E-30 |
| VC1743 | hypothetical protein | -4.13 | 2.88E-09 | -2.35 | 1.35E-07 |
| VC1854 | porin, putative | 12.73 | 5.64E-22 | 2.37 | 3.17E-07 |
| VC1888 | hemolysin-related protein | 2.91 | 2.75E-07 | 2.85 | 3.31E-25 |
| VC2213 | outer membrane protein OmpA | 5.56 | 2.72E-10 | 6.97 | 1.20E-34 |

| | | | | | |
|---|---|---|---|---|---|
| VC2240 | decarboxylase | -3.31 | 4.67E-09 | -3.35 | 8.81E-17 |
| VC2337 | transcriptional regulator, LacI family | -2.48 | 6.00E-07 | -2.24 | 4.90E-10 |
| VC2338 | ß-lactamase lacZ | -12.10 | 3.81E-28 | -2.91 | 1.40E-17 |
| VC2361 | formate acetyl transferase-related protein | 4.14 | 4.05E-08 | -2.67 | 1.94E-17 |
| VC2385 | RNA-directed DNA polymerase | -2.07 | 8.01E-05 | 2.11 | 2.88E-07 |
| VC2387 | conserved hypothetical protein | -2.81 | 2.04E-04 | 2.50 | 9.87E-11 |
| VC2389 | carbamoyl-phosphate synthase, large subunit | 2.19 | 6.87E-04 | 2.23 | 2.62E-12 |
| VC2464 | sigma-E factor regulatory protein RseC | -11.00 | 4.83E-43 | -3.96 | 1.58E-33 |
| VC2465 | sigma-E factor regulatory protein RseB | -14.73 | 2.43E-43 | -3.58 | 5.91E-26 |
| VC2466 | sigma-E factor negative regulatory protein RseA | -19.88 | 4.50E-68 | -4.14 | 1.26E-31 |
| VC2467 | RNA polymerase sigma-E factor | -71453.09 | 2.13E-07 | -3.70 | 2.47E-14 |
| VC2565 | elaA protein | -5.79 | 2.29E-12 | -6.72 | 1.33E-40 |
| VC2566 | conserved hypothetical protein | -6.70 | 1.71E-18 | -3.78 | 1.10E-37 |
| VC2662 | conserved hypothetical protein | 3.56 | 1.70E-08 | 5.18 | 2.38E-39 |
| VC2667 | hypothetical protein | 2.58 | 2.97E-04 | -2.17 | 3.54E-05 |
| VC2697 | GGDEF family protein | 2.00 | 4.11E-04 | -2.66 | 5.11E-11 |
| VC2753 | hypothetical protein | 5.14 | 6.70E-08 | 2.48 | 9.55E-10 |
| VCA0008 | methyl-accepting chemotaxis protein | 2.16 | 6.90E-04 | -13.72 | 2.04E-57 |
| VCA0013 | maltodextrin phosphorylase | -2.07 | 2.09E-04 | -2.92 | 2.19E-11 |
| VCA0017 | hcp protein | -4.19 | 4.72E-09 | 12.02 | 3.29E-57 |
| VCA0035 | phosphatidylglycerop hosphatase B, putative | 5.19 | 5.60E-24 | 3.19 | 5.28E-19 |
| VCA0052 | hypothetical protein | -2.09 | 4.17E-05 | -2.07 | 3.00E-08 |
| VCA0080 | GGDEF family protein | 2.76 | 2.10E-05 | -4.35 | 9.21E-31 |
| VCA0087 | hypothetical protein | 3.29 | 1.59E-05 | 2.70 | 9.55E-06 |
| VCA0136 | glycerophosphoryl diester phosphodiesterase | -2.71 | 1.06E-06 | 11.57 | 1.35E-42 |
| VCA0137 | glycerol-3-phosphate transporter | -7.45 | 3.84E-30 | 10.54 | 1.65E-73 |
| VCA0156 | conserved hypothetical protein | 2.22 | 1.15E-05 | -6.93 | 1.13E-30 |
| VCA0158 | hypothetical protein | 4.00 | 4.34E-10 | -2.72 | 4.52E-10 |
| VCA0159 | conserved hypothetical protein | 4.39 | 3.44E-09 | -2.27 | 2.67E-16 |
| VCA0161 | tryptophanase | 4.48 | 8.61E-14 | -2.95 | 3.88E-11 |
| VCA0190 | hypothetical protein | 2.04 | 2.84E-04 | -3.92 | 1.45E-25 |
| VCA0212 | hypothetical protein | 2.06 | 2.86E-04 | -4.39 | 3.64E-26 |
| VCA0219 | haemolysin | 2.51 | 1.80E-05 | -2.09 | 4.01E-08 |
| VCA0223 | protease | 2.69 | 3.92E-05 | 3.11 | 7.09E-08 |
| VCA0227 | iron(III) ABC transporter, periplasmiciron-compound-binding protein | 7.08 | 1.35E-11 | 19.29 | 1.75E-88 |
| VCA0228 | iron(III) ABC transporter, permease protein | 2.45 | 8.32E-04 | 2.99 | 2.18E-09 |
| VCA0268 | methyl-accepting chemotaxis protein | 2.19 | 5.80E-04 | -5.91 | 5.83E-28 |
| VCA0269 | decarboxylase, group II | 2.57 | 3.15E-05 | -4.12 | 1.67E-19 |
| VCA0282 | IS5 transposase | -2.16 | 4.40E-04 | -2.45 | 1.93E-08 |
| VCA0288 | initiation factor IF3 | 7.94 | 2.02E-09 | 2.29 | 1.04E-08 |
| VCA0308 | deoxyguanosinetripho sphatetriphosphohydrolase-rela ted protein | -2.15 | 1.34E-05 | 2.11 | 1.16E-05 |
| VCA0441 | hypothetical protein | -2.16 | 1.59E-05 | 2.83 | 2.26E-22 |
| VCA0507 | transposase OrfAB, subunit A | -2.50 | 2.67E-07 | -2.54 | 2.06E-17 |
| VCA0508 | transposase OrfAB, subunit B | -2.90 | 5.62E-12 | -2.61 | 7.82E-11 |
| VCA0517 | 1-phosphofructokinas e | -2.61 | 4.97E-05 | 2.47 | 2.70E-08 |
| VCA0540 | formate transporter 1, putative | -2.55 | 1.04E-05 | 2.37 | 2.63E-06 |
| VCA0576 | heme transport protein HutA | 2.33 | 8.47E-05 | 5.15 | 1.04E-29 |
| VCA0615 | peptide methionine sulfoxide reductase | 11.68 | 1.47E-06 | -2.28 | 3.45E-07 |
| VCA0647 | hypothetical protein | 4.07 | 4.68E-05 | -3.61 | 3.13E-10 |
| VCA0648 | hypothetical protein | 3.53 | 2.14E-04 | -5.07 | 6.90E-12 |
| VCA0649 | hypothetical protein | 4.07 | 1.12E-04 | -3.48 | 1.50E-08 |
| VCA0650 | hypothetical protein | 5.14 | 3.16E-05 | -4.66 | 1.55E-12 |
| VCA0651 | conserved hypothetical protein | 7.46 | 8.48E-06 | -4.80 | 2.08E-05 |
| VCA0657 | aerobic glycerol-3-phosphate dehydrogenase | -3.08 | 1.75E-05 | 55.01 | 3.66E-159 |
| VCA0678 | periplasmic nitrate reductase | 3.18 | 8.96E-04 | -2.37 | 2.89E-08 |
| VCA0684 | regulatory protein UhpC | -3.08 | 1.29E-04 | -2.47 | 8.11E-14 |
| VCA0721 | hypothetical protein | 2.77 | 4.00E-05 | 4.34 | 1.11E-20 |
| VCA0731 | hypothetical protein | 2.98 | 1.69E-04 | -6.51 | 9.17E-18 |
| VCA0744 | glycerol kinase | -2.15 | 2.10E-05 | 12.12 | 3.52E-69 |
| VCA0745 | pseudogene with authentic frameshift mutation | -2.80 | 3.72E-06 | 12.67 | 3.63E-82 |
| VCA0758 | arginine ABC transporter, permease protein | 2.01 | 4.68E-05 | -2.40 | 1.02E-07 |
| VCA0828 | phenylalanine-4-hydr oxylase | 3.40 | 4.09E-12 | 2.38 | 7.71E-09 |
| VCA0845 | hypothetical protein | 2.44 | 1.21E-03 | 2.02 | 1.48E-07 |
| VCA0860 | alpha-amylase | -2.11 | 2.95E-04 | -3.42 | 1.45E-12 |
| VCA0865 | hemagglutinin/protea se | 5.87 | 3.93E-10 | -9.40 | 5.40E-35 |
| VCA0907 | conserved hypothetical protein | 3.12 | 3.05E-04 | 2.46 | 9.55E-09 |
| VCA0908 | conserved hypothetical protein | 3.71 | 3.81E-05 | 2.51 | 6.72E-09 |
| VCA0909 | oxygen-independent coproporphyrinogen IIIoxidase, putative | 6.26 | 4.41E-11 | 2.99 | 3.95E-10 |
| VCA0910 | tonB1 protein | 6.77 | 1.91E-09 | 2.58 | 3.79E-05 |
| VCA0911 | TonB system transport protein ExbB1 | 6.04 | 1.98E-07 | 3.78 | 3.70E-08 |
| VCA0912 | TonB system transport protein ExbD1 | 4.53 | 1.35E-05 | 3.35 | 5.33E-07 |
| VCA0913 | hemin ABC transporter, periplasmic hemin-bindingprotein HutB | 4.01 | 7.10E-06 | 2.42 | 4.57E-05 |
| VCA0935 | hypothetical protein | 4.63 | 2.25E-08 | -6.55 | 4.68E-37 |
| VCA0944 | maltose ABC transporter, permease protein | -2.36 | 1.24E-04 | -4.30 | 4.21E-15 |
| VCA0946 | maltose/maltodextrin ABC transporter,ATP-binding protein | -2.38 | 1.43E-04 | -2.46 | 1.93E-07 |
| VCA0957 | malate synthase-related protein | 3.92 | 2.17E-16 | -2.18 | 4.50E-08 |
| VCA0988 | methyl-accepting chemotaxis protein | 2.28 | 4.11E-06 | -2.77 | 2.89E-13 |
| VCA1024 | hypothetical protein | 3.54 | 9.46E-06 | -13.60 | 5.70E-85 |
| VCA1028 | maltoporin | -3.98 | 1.50E-07 | -2.79 | 2.80E-06 |
| VCA1086 | response regulator | 2.23 | 4.99E-04 | -5.77 | 1.04E-41 |
| VCA1091 | chemotaxis protein methyltransferase CheR | 2.42 | 8.38E-07 | -7.93 | 1.10E-37 |
| VCA1092 | methyl-accepting chemotaxis protein | 2.34 | 3.94E-04 | -7.96 | 4.56E-52 |
| VCA1093 | purine-binding chemotaxis protein CheW | 2.14 | 3.27E-06 | -7.87 | 1.24E-40 |
| VCA1108 | oxidoreductase, short-chaindehydrogenase/reduc tase family | 2.74 | 1.18E-05 | -3.38 | 5.12E-09 |
