## Supplementary material for "Modular small RNA drives pathogen emergence": Table S5

**Table S5**: Differentially-expressed 453 genes in the *ompU* mutant whose expression is rescued by OueS, compared to WT C6706, when grown under virulence inducing AKI conditions

| Name | Function | ∆*ompU*/WT | | ∆*ompU*-pOueS/WT | |
|---|---|---|---|---|---|
| | | FC | FDR p-value | FC | FDR p-value |
| 5Sc | 5S ribosomal RNA | -2.0 | 2.5E-08 | -34.2 | 3.3E-38 |
| 5Sd | 5S ribosomal RNA | -2.0 | 2.6E-08 | -35.6 | 1.4E-39 |
| 5Sg | 5S ribosomal RNA | -142.2 | 4.0E-107 | -46.2 | 8.9E-22 |
| VC0001 | hypothetical protein | -2.3 | 1.8E-03 | -3.0 | 3.2E-03 |
| VC0018 | 16 kDa heat shock protein A | -2.0 | 3.8E-05 | -2.4 | 9.1E-04 |
| VC0032 | ComM-related protein | -2.3 | 5.5E-04 | -2.4 | 2.2E-03 |
| VC0069 | multidrug resistance protein, putative | -2.9 | 6.1E-13 | -2.7 | 2.8E-05 |
| VC0079 | conserved hypothetical protein | -6.2 | 4.4E-50 | -3.5 | 4.5E-11 |
| VC0089 | cytochrome c551 peroxidase | -20.6 | 7.9E-69 | -5.2 | 1.7E-06 |
| VC0090 | DNA-damage-inducible protein F | -4.4 | 3.2E-25 | -3.1 | 3.5E-10 |
| VC0098 | methyl-accepting chemotaxis protein | -3.4 | 4.1E-25 | -2.3 | 1.5E-04 |
| VC0139 | DPS family protein | -4.2 | 1.1E-20 | -2.7 | 2.0E-06 |
| VC0142 | hypothetical protein | -2.4 | 1.5E-08 | -2.2 | 6.4E-05 |
| VC0169 | hypothetical protein | -6.7 | 1.2E-23 | -4.1 | 2.8E-06 |
| VC0172 | peptide ABC transporter, permease protein | -2.1 | 1.2E-06 | -2.2 | 1.6E-03 |
| VC0185 | transposase, putative | -3.2 | 5.3E-15 | -3.4 | 2.6E-12 |
| VC0300 | conserved hypothetical protein | -7.6 | 4.2E-34 | -3.3 | 2.1E-03 |
| VC0308 | hypothetical protein | -3.5 | 6.6E-19 | -2.7 | 1.1E-05 |
| VC0365 | bacterioferritin | -3.5 | 7.0E-21 | -3.2 | 5.2E-08 |
| VC0383 | hypothetical protein | -4.1 | 2.3E-08 | -3.2 | 2.3E-04 |
| VC0403 | MSHA biogenesis protein MshM | -3.1 | 3.5E-13 | -2.5 | 4.6E-05 |
| VC0404 | MSHA biogenesis protein MshN | -2.3 | 2.3E-12 | -2.1 | 1.4E-04 |
| VC0407 | MSHA biogenesis protein MshF | -2.1 | 8.6E-08 | -2.4 | 1.2E-06 |
| VC0412 | hypothetical protein | -2.2 | 6.8E-13 | -2.6 | 1.9E-07 |
| VC0413 | MshP | -2.8 | 5.9E-20 | -3.3 | 4.6E-10 |
| VC0414 | MshQ | -2.3 | 1.6E-10 | -2.2 | 5.6E-03 |
| VC0501 | pseudogene within VSP-II; potential transposase | -3.3 | 1.0E-14 | -2.2 | 4.0E-03 |
| VC0633 | outer membrane protein OmpU | -3191116.3 | 3.4E-13 | -21.8 | 3.7E-07 |
| VC0658 | c-di-GMP phosphodiesterase A-related protein | -5.0 | 1.2E-31 | -3.0 | 8.7E-05 |
| VC0697 | hypothetical protein | -4.0 | 6.2E-28 | -2.7 | 7.7E-06 |
| VC0728 | conserved hypothetical protein | -4.8 | 1.3E-26 | -2.4 | 4.2E-06 |
| VC0734 | malate synthase A | -4.9 | 2.6E-21 | -4.2 | 1.0E-11 |
| VC0736 | isocitrate lyase | -6.5 | 8.4E-43 | -2.2 | 6.5E-04 |
| VC0817 | transposase, putative | -2.0 | 2.0E-06 | -2.1 | 1.5E-04 |
| VC0893 | chemotaxis protein PomB | -3.2 | 4.0E-23 | -2.3 | 9.8E-04 |
| VC0935 | hypothetical protein | -2.9 | 2.2E-08 | -2.3 | 6.5E-03 |
| VC0936 | polysaccharide export-related protein | -5.5 | 1.7E-18 | -5.3 | 1.2E-08 |
| VC0937 | exopolysaccharide biosynthesis protein,putative | -2.6 | 1.3E-10 | -2.4 | 2.6E-03 |
| VC0938 | hypothetical protein | -4.2 | 3.2E-12 | -3.6 | 1.2E-06 |
| VC1008 | sodium-type flagellar protein MotY | -3.7 | 6.7E-24 | -2.2 | 1.0E-03 |
| VC1009 | conserved hypothetical protein | -4.2 | 4.6E-28 | -2.6 | 1.3E-06 |
| VC1029 | GGDEF family protein | -3.6 | 6.1E-15 | -2.5 | 3.1E-03 |
| VC1031 | inosine monophosphate dehydrogenase-relatedprotein | -10.2 | 1.3E-74 | -5.8 | 9.2E-18 |
| VC1043 | long-chain fatty acid transport protein | -2.4 | 1.5E-10 | -2.7 | 1.0E-03 |
| VC1050 | response regulator | -5.1 | 3.4E-34 | -3.1 | 1.5E-06 |
| VC1066 | hypothetical protein | -4.0 | 4.8E-22 | -2.7 | 6.3E-08 |
| VC1080 | hypothetical protein | -3.8 | 6.6E-24 | -2.7 | 2.1E-06 |
| VC1082 | response regulator | -11.2 | 8.6E-55 | -3.0 | 7.9E-09 |
| VC1083 | hypothetical protein | -10.3 | 9.0E-65 | -3.1 | 4.1E-11 |
| VC1084 | sensory box sensor histidine kinase | -12.6 | 1.2E-44 | -4.3 | 5.5E-07 |
| VC1085 | sensor histidine kinase | -8.0 | 5.7E-43 | -4.0 | 1.5E-13 |
| VC1086 | response regulator | -5.2 | 1.7E-44 | -2.9 | 2.9E-07 |
| VC1087 | response regulator | -4.4 | 4.5E-30 | -2.4 | 4.7E-03 |
| VC1088 | sensor histidine kinase | -5.9 | 1.8E-33 | -4.1 | 4.3E-10 |
| VC1089 | periplasmic binding protein-related protein | -4.7 | 2.5E-31 | -2.6 | 1.6E-03 |
| VC1124 | conserved hypothetical protein | -4.7 | 2.2E-24 | -2.4 | 4.3E-06 |
| VC1125 | hypothetical protein | -4.9 | 6.0E-26 | -3.2 | 3.7E-05 |
| VC1131 | conserved hypothetical protein | -2.3 | 7.0E-08 | -2.1 | 1.5E-04 |
| VC1187 | hypothetical protein | -2.0 | 1.2E-04 | -3.0 | 2.9E-05 |
| VC1189 | hypothetical protein | -4.5 | 2.5E-19 | -2.5 | 2.5E-03 |
| VC1207 | hypothetical protein | -4.7 | 1.3E-21 | -2.4 | 2.0E-03 |
| VC1211 | conserved hypothetical protein | -2.1 | 1.8E-06 | -2.4 | 1.6E-03 |
| VC1222 | integration host factor, alpha subunit | -3.7 | 2.2E-12 | -2.2 | 8.0E-03 |
| VC1223 | hypothetical protein | -3.9 | 1.5E-17 | -2.6 | 6.2E-04 |
| VC1247 | hypothetical protein | -2.6 | 1.7E-10 | -2.5 | 8.8E-04 |
| VC1248 | methyl-accepting chemotaxis protein | -10.0 | 1.1E-55 | -6.1 | 1.4E-08 |
| VC1262 | hypothetical protein | -2.1 | 1.4E-05 | -2.5 | 2.4E-03 |
| VC1295 | conserved hypothetical protein | -2.9 | 4.3E-13 | -3.4 | 1.2E-05 |

| | | | | | |
|---|---|---|---|---|---|
| VC1298 | methyl-accepting chemotaxis protein | -2.1 | 4.4E-05 | -2.4 | 1.6E-03 |
| VC1312 | alanine racemase, putative | -3.5 | 9.8E-15 | -2.7 | 5.7E-06 |
| VC1313 | methyl-accepting chemotaxis protein | -4.2 | 4.7E-20 | -3.0 | 4.9E-05 |
| VC1316 | chemotaxis protein CheY, putative | -4.3 | 5.9E-32 | -2.1 | 2.2E-04 |
| VC1322 | conserved hypothetical protein | -5.8 | 3.5E-28 | -3.8 | 6.7E-06 |
| VC1323 | hypothetical protein | -3.8 | 2.3E-20 | -2.3 | 8.1E-04 |
| VC1328 | galactoside ABC transporter, permease protein | -3.1 | 7.2E-18 | -2.2 | 4.6E-04 |
| VC1348 | response regulator | -5.1 | 1.0E-25 | -4.4 | 1.8E-12 |
| VC1349 | sensory box sensor histidine kinase/responseregulator | -4.4 | 2.9E-25 | -3.3 | 4.6E-08 |
| VC1353 | GGDEF family protein | -2.1 | 1.0E-05 | -2.1 | 1.8E-03 |
| VC1355 | acylphosphatase | -2.9 | 3.0E-13 | -2.2 | 1.9E-04 |
| VC1359 | amino acid ABC transporter, ATP-binding protein | -3.0 | 1.4E-13 | -2.1 | 1.6E-03 |
| VC1360 | amino acid ABC transporter, permease protein | -4.1 | 1.9E-17 | -2.9 | 3.9E-04 |
| VC1361 | amino acid ABC transporter, permease protein | -4.0 | 2.8E-19 | -2.9 | 9.3E-06 |
| VC1362 | amino acid ABC transporter, periplasmic aminoacid-binding protein | -9.0 | 1.9E-52 | -4.8 | 2.5E-05 |
| VC1368 | hypothetical protein | -3.4 | 3.1E-17 | -2.2 | 1.6E-03 |
| VC1369 | conserved hypothetical protein | -6.1 | 6.2E-31 | -4.4 | 2.5E-08 |
| VC1370 | GGDEF family protein | -7.2 | 9.6E-44 | -5.6 | 1.1E-13 |
| VC1371 | hypothetical protein | -2.8 | 3.1E-13 | -2.6 | 9.6E-08 |
| VC1376 | GGDEF family protein | -2.8 | 1.6E-16 | -2.6 | 5.5E-05 |
| VC1394 | methyl-accepting chemotaxis protein | -6.1 | 1.1E-45 | -5.4 | 3.1E-10 |
| VC1395 | response regulator cheY1 | -7.3 | 3.3E-55 | -5.9 | 5.0E-12 |
| VC1396 | hypothetical protein | -6.6 | 1.1E-33 | -4.0 | 2.8E-07 |
| VC1397 | chemotaxis protein CheA | -9.0 | 5.9E-55 | -5.3 | 9.1E-12 |
| VC1398 | chemotaxis protein CheY | -9.3 | 4.9E-43 | -5.4 | 1.5E-11 |
| VC1399 | chemotaxis protein methyltransferase CheR | -8.3 | 3.4E-50 | -6.4 | 9.9E-12 |
| VC1400 | hypothetical protein | -7.3 | 8.3E-36 | -3.9 | 4.6E-06 |
| VC1401 | protein-glutamate methylesterase CheB | -5.3 | 9.9E-30 | -2.9 | 1.4E-05 |
| VC1403 | methyl-accepting chemotaxis protein | -4.7 | 1.9E-23 | -2.6 | 1.0E-03 |
| VC1405 | methyl-accepting chemotaxis protein | -3.1 | 1.4E-15 | -3.5 | 7.9E-08 |
| VC1406 | methyl-accepting chemotaxis protein | -3.4 | 8.9E-17 | -2.7 | 7.3E-05 |
| VC1495 | hypothetical protein | -2.8 | 1.9E-05 | -5.7 | 7.4E-09 |
| VC1509 | nicotinatemononucleo tide:5,6-dimethylbenzimidazole phosphoribosyltransferase | -3.8 | 5.1E-17 | -2.1 | 4.6E-03 |
| VC1528 | hypothetical protein | -3.0 | 4.7E-13 | -2.6 | 1.5E-03 |
| VC1538 | hypothetical protein | -2.9 | 1.9E-14 | -2.4 | 6.8E-05 |
| VC1560 | catalase/peroxidase | -5.0 | 7.2E-28 | -2.9 | 5.9E-03 |
| VC1593 | GGDEF family protein | -2.9 | 3.2E-16 | -2.2 | 5.2E-04 |
| VC1594 | aldose 1-epimerase | -3.7 | 3.1E-25 | -2.6 | 1.3E-03 |
| VC1601 | hypothetical protein | -3.2 | 1.5E-14 | -2.5 | 9.2E-07 |
| VC1602 | chemotaxis protein CheV | -3.1 | 3.8E-09 | -2.7 | 1.5E-03 |
| VC1603 | hypothetical protein | -3.3 | 2.6E-18 | -3.3 | 1.4E-06 |
| VC1605a | hypothetical protein | -2.8 | 3.6E-17 | -2.1 | 7.3E-03 |
| VC1641 | conserved hypothetical protein | -3.1 | 1.5E-22 | -3.0 | 1.7E-08 |
| VC1643 | methyl-accepting chemotaxis protein | -5.5 | 2.9E-20 | -2.7 | 2.5E-04 |
| VC1661 | hypothetical protein | -2.8 | 1.7E-11 | -2.2 | 1.4E-03 |
| VC1662 | conserved hypothetical protein | -3.3 | 1.1E-15 | -2.6 | 8.7E-04 |
| VC1672 | DNA-3-methyladenine glycosidase I | -2.9 | 4.0E-10 | -2.5 | 1.2E-03 |
| VC1674 | periplasmic linker protein, putative | -3.2 | 6.8E-16 | -2.4 | 1.6E-03 |
| VC1675 | multidrug resistance protein, putative | -2.8 | 8.8E-16 | -2.1 | 2.4E-04 |
| VC1707 | hypothetical protein | -5.8 | 1.6E-28 | -3.1 | 5.3E-06 |
| VC1736 | arginyl-tRNA-protein transferase-relatedprotein | -3.3 | 5.1E-19 | -2.5 | 1.5E-04 |
| VC1831 | sensor histidine kinase | -3.4 | 1.7E-21 | -4.0 | 1.7E-12 |
| VC1851 | conserved hypothetical protein | -4.8 | 2.0E-31 | -3.6 | 1.1E-07 |
| VC1868 | methyl-accepting chemotaxis protein | -6.8 | 2.2E-42 | -4.3 | 2.0E-08 |
| VC1873 | conserved hypothetical protein | -13.6 | 4.8E-49 | -4.6 | 5.8E-08 |
| VC1874 | conserved hypothetical protein | -12.5 | 2.4E-82 | -5.9 | 5.3E-09 |
| VC1914 | integration host factor, beta subunit | -2.9 | 2.4E-16 | -2.1 | 7.5E-04 |
| VC1932 | hypothetical protein | -2.6 | 1.4E-06 | -2.3 | 5.3E-03 |
| VC1933 | hypothetical protein | -3.8 | 2.0E-21 | -3.8 | 3.2E-07 |
| VC1934 | GGDEF family protein | -3.8 | 6.6E-21 | -4.0 | 1.8E-06 |
| VC1967 | methyl-accepting chemotaxis protein | -3.4 | 5.0E-15 | -3.5 | 4.3E-08 |
| VC1997 | hypothetical protein | -2.5 | 2.3E-13 | -2.1 | 5.2E-04 |
| VC2038 | hypothetical protein | -2.4 | 8.5E-07 | -2.4 | 6.8E-04 |
| VC2046 | hypothetical protein | -3.4 | 9.4E-18 | -2.8 | 5.5E-09 |
| VC2047 | oxidoreductase, short-chaindehydrogenase/reduc tase family | -3.4 | 1.8E-25 | -2.4 | 4.3E-03 |
| VC2058 | hypothetical protein | -3.0 | 1.8E-22 | -2.5 | 2.1E-05 |
| VC2060 | conserved hypothetical protein | -2.9 | 1.1E-08 | -2.8 | 5.2E-04 |
| VC2061 | ParA family protein | -2.4 | 5.4E-13 | -2.1 | 3.7E-04 |
| VC2120 | flagellar biosynthetic protein FlhB | -2.4 | 6.4E-12 | -2.1 | 4.0E-03 |
| VC2121 | flagellar biosynthetic protein FliR | -2.2 | 1.6E-08 | -2.2 | 2.8E-03 |
| VC2122 | flagellar biosynthetic protein FliQ | -2.8 | 7.4E-15 | -2.8 | 1.5E-04 |
| VC2123 | flagellar biosynthetic protein FliP | -3.0 | 2.4E-12 | -2.9 | 8.5E-06 |

| | | | | | |
|---|---|---|---|---|---|
| VC2124 | flagellar protein FliO | -2.8 | 2.6E-14 | -2.1 | 2.6E-04 |
| VC2128 | flagellar hook-length control protein FliK,putative | -4.4 | 1.2E-16 | -2.4 | 1.6E-03 |
| VC2138 | flagellar protein FliS | -3.2 | 4.6E-20 | -2.5 | 8.4E-06 |
| VC2139 | flagellar rod protein FlaI, putative | -2.9 | 1.0E-23 | -2.4 | 2.8E-05 |
| VC2140 | flagellar hook-associated protein FliD | -4.1 | 7.3E-30 | -2.9 | 7.7E-04 |
| VC2141 | flagellin FlaG | -4.1 | 2.3E-25 | -2.8 | 1.0E-03 |
| VC2144 | flagellin FlaE | -4.4 | 3.2E-32 | -3.3 | 1.5E-08 |
| VC2161 | methyl-accepting chemotaxis protein | -4.1 | 4.3E-25 | -2.9 | 5.7E-04 |
| **VC2190** | flagellar hook-associated protein FlgL | -3.9 | 4.2E-13 | -2.8 | 3.2E-04 |
| **VC2192** | flagellar protein FlgJ | -4.3 | 2.3E-28 | -2.2 | 2.1E-04 |
| VC2201 | chemotaxis protein methyltransferase CheR | -3.4 | 5.2E-20 | -2.7 | 4.1E-03 |
| VC2204 | negative regulator of flagellin synthesis FlgM,putative | -3.3 | 4.3E-11 | -2.7 | 2.9E-04 |
| VC2205 | hypothetical protein | -3.6 | 5.2E-21 | -3.1 | 2.0E-08 |
| VC2206 | conserved hypothetical protein | -4.0 | 3.8E-29 | -2.6 | 5.9E-06 |
| VC2207 | hypothetical protein | -3.2 | 3.8E-11 | -2.2 | 3.1E-03 |
| VC2240 | decarboxylase | -3.4 | 8.7E-16 | -2.6 | 1.2E-06 |
| VC2264 | conserved hypothetical protein | -4.3 | 6.0E-30 | -2.9 | 2.1E-04 |
| VC2285 | GGDEF family protein | -2.2 | 4.8E-07 | -2.5 | 4.3E-05 |
| VC2314 | hypothetical protein | -4.5 | 4.5E-14 | -3.9 | 3.5E-06 |
| VC2316 | N-acetylglutamate synthase | -3.5 | 9.1E-15 | -2.9 | 2.7E-04 |
| VC2340 | conserved hypothetical protein | -7.2 | 2.8E-27 | -4.8 | 4.8E-07 |
| VC2357 | hypothetical protein | -2.1 | 4.0E-08 | -2.1 | 6.1E-04 |
| VC2358 | hypothetical protein | -3.9 | 1.8E-24 | -2.6 | 1.3E-03 |
| VC2370 | sensory box/GGDEF family protein | -2.8 | 7.1E-12 | -2.7 | 5.1E-07 |
| VC2384 | conserved hypothetical protein | -4.0 | 2.4E-17 | -3.9 | 1.1E-05 |
| VC2454 | GGDEF family protein | -3.7 | 7.2E-23 | -3.0 | 4.2E-06 |
| VC2455 | hypothetical protein | -5.0 | 6.7E-24 | -3.0 | 2.2E-05 |
| VC2456 | hypothetical protein | -4.6 | 1.2E-31 | -2.8 | 2.6E-05 |
| VC2464 | sigma-E factor regulatory protein RseC | -4.0 | 5.6E-32 | -3.2 | 1.1E-04 |
| VC2465 | sigma-E factor regulatory protein RseB | -3.6 | 1.1E-24 | -2.9 | 3.2E-08 |
| VC2466 | sigma-E factor negative regulatory protein RseA | -4.1 | 3.8E-30 | -2.3 | 2.7E-04 |
| VC2467 | RNA polymerase sigma-E factor | -3.7 | 2.0E-13 | -3.0 | 5.7E-04 |
| VC2507 | conserved hypothetical protein | -10.1 | 5.3E-74 | -3.0 | 7.1E-06 |
| VC2565 | elaA protein | -6.7 | 6.2E-39 | -4.6 | 6.0E-16 |
| VC2566 | conserved hypothetical protein | -3.8 | 4.7E-36 | -3.6 | 5.9E-11 |
| VC2601 | sodium-type flagellar protein MotX | -4.4 | 4.1E-24 | -2.4 | 3.6E-06 |
| VC2622 | hypothetical protein | -5.4 | 2.6E-28 | -4.2 | 4.1E-10 |
| VC2647 | conserved hypothetical protein | -2.1 | 2.3E-07 | -2.8 | 2.0E-08 |
| VC2702 | transcriptional regulator, LuxR family | -2.8 | 1.4E-18 | -2.2 | 4.8E-05 |
| VC2704 | hypothetical protein | -12.6 | 1.6E-44 | -7.7 | 3.3E-08 |
| VC2705 | sodium/solute symporter, putative | -13.9 | 6.8E-69 | -11.7 | 1.0E-10 |
| VC2717 | hypothetical protein | -2.6 | 2.2E-14 | -2.9 | 3.8E-09 |
| VC2726 | general secretion pathway protein K | -2.1 | 1.4E-07 | -2.1 | 1.9E-03 |
| VC2727 | general secretion pathway protein J | -2.2 | 9.9E-08 | -2.7 | 7.8E-04 |
| VC2750 | GGDEF family protein | -3.0 | 1.3E-15 | -2.9 | 1.1E-04 |
| VCA0008 | methyl-accepting chemotaxis protein | -13.7 | 2.7E-55 | -7.3 | 8.6E-14 |
| VCA0009 | hypothetical protein | -12.8 | 2.8E-23 | -7.6 | 2.4E-12 |
| VCA0030 | hypothetical protein | -2.2 | 3.6E-08 | -2.3 | 9.8E-03 |
| VCA0031 | methyl-accepting chemotaxis protein | -3.9 | 6.5E-22 | -2.5 | 1.1E-03 |
| VCA0032 | hypothetical protein | -3.6 | 2.0E-22 | -2.4 | 1.0E-04 |
| VCA0074 | GGDEF family protein | -3.3 | 9.1E-17 | -4.2 | 4.5E-10 |
| VCA0075 | hypothetical protein | -2.5 | 9.2E-06 | -2.6 | 1.4E-03 |
| VCA0078 | hypothetical protein | -2.4 | 2.6E-11 | -2.1 | 4.6E-03 |
| VCA0080 | GGDEF family protein | -4.4 | 2.6E-29 | -3.0 | 3.6E-06 |
| VCA0152 | conserved hypothetical protein | -10.6 | 6.6E-50 | -4.5 | 4.6E-07 |
| VCA0153 | conserved hypothetical protein | -9.5 | 1.2E-29 | -3.5 | 7.5E-04 |
| VCA0154 | conserved hypothetical protein | -8.0 | 4.0E-33 | -4.2 | 2.0E-07 |
| VCA0155 | NADH dehydrogenase, putative | -6.5 | 5.1E-44 | -3.8 | 6.3E-07 |
| VCA0156 | conserved hypothetical protein | -6.9 | 3.1E-29 | -3.3 | 8.3E-07 |
| VCA0157 | NADH dehydrogenase, putative | -5.3 | 4.9E-37 | -3.4 | 1.1E-05 |
| VCA0188 | hypothetical protein | -4.5 | 1.2E-16 | -3.9 | 7.7E-06 |
| VCA0191 | conserved hypothetical protein | -4.1 | 3.9E-26 | -2.5 | 4.0E-04 |
| VCA0195 | hypothetical protein | -3.8 | 1.3E-20 | -2.4 | 9.6E-05 |
| VCA0210 | response regulator, putative | -7.3 | 5.9E-55 | -4.2 | 7.9E-09 |
| VCA0211 | sensory box sensor histidine kinase | -6.6 | 1.7E-50 | -4.7 | 3.3E-11 |
| VCA0212 | hypothetical protein | -4.4 | 7.2E-25 | -3.5 | 6.5E-10 |
| VCA0217 | GGDEF family protein | -2.1 | 2.5E-06 | -2.7 | 9.6E-05 |
| VCA0220 | hemolysin secretion protein HylB | -2.5 | 9.5E-14 | -2.1 | 1.5E-03 |
| VCA0268 | methyl-accepting chemotaxis protein | -5.9 | 1.4E-26 | -4.9 | 3.4E-10 |
| VCA0269 | decarboxylase, group II | -4.1 | 2.1E-18 | -3.4 | 8.3E-06 |
| VCA0319 | conserved hypothetical protein | -2.3 | 7.5E-05 | -4.8 | 1.0E-08 |
| VCA0534 | pseudogene; contains authentic frameshift mutation | -2.5 | 8.7E-08 | -2.9 | 9.0E-05 |

| | | | | | |
|---|---|---|---|---|---|
| VCA0551 | hypothetical protein | -3.7 | 4.6E-40 | -2.9 | 1.8E-08 |
| VCA0557 | GGDEF family protein | -2.7 | 4.1E-16 | -3.0 | 1.1E-07 |
| VCA0570 | Sui1 family protein | -2.5 | 5.1E-11 | -2.2 | 6.5E-04 |
| VCA0583 | hypothetical protein | -9.2 | 9.8E-40 | -4.7 | 3.7E-10 |
| VCA0593 | hypothetical protein | -4.9 | 7.9E-32 | -3.4 | 1.8E-08 |
| VCA0619 | hypothetical protein | -4.6 | 6.1E-38 | -2.3 | 5.0E-06 |
| VCA0646 | conserved hypothetical protein/hemolysin,putative | -5.0 | 1.4E-25 | -3.4 | 1.5E-06 |
| VCA0647 | hypothetical protein | -3.6 | 1.8E-09 | -2.9 | 1.1E-04 |
| VCA0648 | hypothetical protein | -5.1 | 4.6E-11 | -3.2 | 1.5E-04 |
| VCA0649 | hypothetical protein | -3.5 | 7.2E-08 | -2.6 | 6.5E-03 |
| VCA0650 | hypothetical protein | -4.7 | 1.1E-11 | -3.6 | 1.5E-04 |
| VCA0651 | conserved hypothetical protein | -4.8 | 6.6E-05 | -2.8 | 8.2E-03 |
| VCA0659 | protein F-related protein | -3.9 | 4.2E-37 | -3.6 | 5.2E-10 |
| VCA0678 | periplasmic nitrate reductase | -2.4 | 1.3E-07 | 2.9 | 1.7E-03 |
| VCA0679 | periplasmic nitrate reductase, cytochrome c-typeprotein | -2.7 | 9.5E-07 | 2.7 | 9.3E-04 |
| VCA0681 | conserved hypothetical protein | -6.6 | 2.1E-50 | -4.1 | 6.9E-09 |
| VCA0683 | sensor protein UhpB | -2.6 | 1.2E-11 | -2.3 | 1.6E-03 |
| VCA0695 | hypothetical protein | -6.0 | 7.1E-24 | -3.6 | 7.8E-06 |
| VCA0698 | hypothetical protein | -7.4 | 6.4E-48 | -4.3 | 6.8E-10 |
| VCA0719 | sensor histide kinase | -6.3 | 4.6E-30 | -3.7 | 4.7E-05 |
| VCA0720 | guanylate cyclase-related protein | -5.2 | 4.8E-30 | -3.5 | 1.3E-10 |
| VCA0731 | hypothetical protein | -6.5 | 1.0E-16 | -6.4 | 4.8E-11 |
| VCA0736 | sensor histidine kinase LuxQ | -2.3 | 1.8E-08 | -2.4 | 7.5E-04 |
| VCA0757 | arginine ABC transporter, permease protein | -2.6 | 7.8E-13 | -2.1 | 8.3E-03 |
| VCA0760 | arginine ABC transporter, ATP-binding protein | -3.5 | 4.7E-18 | -2.9 | 2.0E-08 |
| VCA0788 | DnaJ-related protein | -2.4 | 5.1E-11 | -2.3 | 4.5E-04 |
| VCA0803 | serine protease, putative | -13.5 | 1.6E-57 | -6.2 | 1.7E-20 |
| VCA0834 | hypothetical protein | -3.4 | 2.5E-20 | -2.2 | 2.0E-04 |
| VCA0848 | GGDEF family protein | -4.3 | 8.2E-54 | -2.9 | 3.8E-05 |
| VCA0864 | methyl-accepting chemotaxis protein | -2.6 | 7.4E-11 | -2.6 | 2.1E-04 |
| VCA0865 | hemagglutinin/protea se | -9.4 | 2.1E-33 | -8.9 | 8.6E-15 |
| VCA0868 | hypothetical protein | -4.9 | 1.6E-34 | -2.2 | 9.4E-03 |
| VCA0880 | makD | -6.0 | 4.1E-52 | -5.8 | 1.6E-18 |
| VCA0881 | makC | -4.8 | 1.3E-40 | -3.8 | 8.0E-07 |
| VCA0882 | makB | -4.9 | 4.8E-34 | -4.2 | 7.6E-07 |
| VCA0884 | hypothetical protein | -6.0 | 3.8E-42 | -4.2 | 1.5E-06 |
| VCA0892 | hypothetical protein | -3.0 | 2.4E-13 | -3.1 | 7.4E-09 |
| VCA0895 | chemotactic transducer-related protein | -4.1 | 6.0E-26 | -3.4 | 3.0E-07 |
| VCA0900 | hypothetical protein | -3.1 | 1.0E-15 | -2.2 | 4.6E-04 |
| VCA0902 | hypothetical protein | -2.1 | 2.9E-06 | -2.4 | 4.4E-04 |
| VCA0906 | methyl-accepting chemotaxis protein | -8.0 | 1.6E-62 | -4.8 | 1.4E-11 |
| VCA0920 | hypothetical protein | -4.5 | 1.1E-26 | -3.3 | 9.6E-11 |
| VCA0923 | methyl-accepting chemotaxis protein | -6.5 | 1.8E-40 | -4.3 | 2.0E-14 |
| VCA0931 | conserved hypothetical protein | -3.2 | 1.0E-16 | -2.5 | 1.1E-04 |
| VCA0935 | hypothetical protein | -6.6 | 2.0E-35 | -7.0 | 2.1E-07 |
| VCA0965 | GGDEF family protein | -5.7 | 4.5E-31 | -4.7 | 9.1E-09 |
| VCA0978 | amino acid ABC transporter, periplasmic aminoacid-binding protein, putative | -5.6 | 2.8E-31 | -2.9 | 1.3E-08 |
| VCA0979 | methyl-accepting chemotaxis protein | -4.7 | 1.2E-30 | -2.8 | 3.3E-06 |
| VCA0981 | hypothetical protein | -4.7 | 3.8E-31 | -3.3 | 2.2E-05 |
| VCA0988 | methyl-accepting chemotaxis protein | -2.8 | 2.1E-12 | -3.0 | 9.7E-08 |
| VCA1015 | Na+/H+ antiporter | -5.9 | 3.1E-51 | -2.5 | 4.9E-05 |
| VCA1016 | hypothetical protein | -7.8 | 2.9E-75 | -3.7 | 1.1E-08 |
| VCA1017 | methylated-DNA--prot ein-cysteineS-methyltransferas e | -6.0 | 5.4E-46 | -4.3 | 1.1E-10 |
| VCA1024 | hypothetical protein | -13.6 | 3.0E-82 | -4.9 | 3.3E-12 |
| VCA1033 | extracellular solute-binding protein, putative | -6.1 | 1.6E-27 | -3.2 | 8.6E-07 |
| VCA1034 | methyl-accepting chemotaxis protein | -6.8 | 1.4E-45 | -3.7 | 8.6E-11 |
| VCA1054 | conserved hypothetical protein | -5.1 | 1.5E-28 | -2.8 | 1.2E-06 |
| VCA1056 | methyl-accepting chemotaxis protein | -6.4 | 2.7E-30 | -5.1 | 8.2E-13 |
| VCA1086 | response regulator | -5.8 | 5.0E-40 | -3.8 | 1.9E-07 |
| VCA1087 | anti-sigma F factor antagonist, putative | -4.6 | 1.2E-24 | -3.8 | 6.3E-05 |
| VCA1088 | methyl-accepting chemotaxis protein | -5.2 | 3.6E-40 | -4.4 | 4.6E-09 |
| VCA1089 | cheB3 methylesterase | -8.1 | 1.2E-73 | -4.6 | 1.1E-07 |
| VCA1090 | chemotaxis protein CheD, putative | -10.1 | 1.5E-61 | -5.4 | 2.4E-09 |
| VCA1091 | chemotaxis protein methyltransferase CheR | -7.9 | 4.7E-36 | -4.5 | 1.5E-06 |
| VCA1092 | methyl-accepting chemotaxis protein | -8.0 | 3.9E-50 | -3.4 | 1.0E-06 |
| VCA1093 | purine-binding chemotaxis protein CheW | -7.9 | 5.9E-39 | -2.8 | 1.4E-04 |
| VCA1094 | purine-binding chemotaxis protein CheW | -8.2 | 1.9E-42 | -3.9 | 1.1E-05 |
| VCA1095 | chemotaxis protein CheA | -8.0 | 3.7E-28 | -4.2 | 9.4E-07 |
| VCA1096 | chemotaxis protein CheY | -8.0 | 8.4E-45 | -3.8 | 2.3E-05 |
| VCA1097 | conserved hypothetical protein | -7.3 | 1.1E-50 | -3.9 | 7.4E-09 |
| VCA1108 | oxidoreductase, short-chaindehydrogenase/reduc tase family | -3.4 | 2.6E-08 | -2.4 | 2.0E-03 |

| | | | | | |
|---|---|---|---|---|---|
| VC0134 | conserved hypothetical protein | 2.0 | 2.1E-14 | 2.4 | 9.5E-04 |
| VC0156 | vitamin B12 receptor | 6.3 | 2.4E-25 | 3.3 | 6.9E-09 |
| VC0164 | multidrug resistance protein, putative | 2.7 | 3.3E-14 | 3.1 | 1.2E-04 |
| VC0165 | conserved hypothetical protein | 3.1 | 6.0E-10 | 2.6 | 8.5E-06 |
| VC0194 | gamma-glutamyltransp eptidase | 4.3 | 1.1E-24 | 3.2 | 1.2E-05 |
| VC0219 | ribosomal protein L33 | 3.8 | 4.1E-33 | 2.0 | 1.3E-03 |
| VC0295 | acetyl-CoA carboxylase, biotin carboxylase | 3.4 | 2.6E-19 | 2.0 | 1.6E-03 |
| VC0296 | acetyl-CoA carboxylase, biotin carboxyl carrierprotein | 3.2 | 1.4E-23 | 2.5 | 1.1E-04 |
| VC0324 | ribosomal protein L11, RplK | 3.3 | 2.3E-18 | 2.8 | 4.0E-03 |
| VC0327 | ribosomal protein L7/L12 | 3.2 | 1.7E-22 | 2.6 | 3.2E-03 |
| VC0360 | ribosomal protein S7 | 3.2 | 5.1E-26 | 2.3 | 1.1E-03 |
| VC0368 | ribosomal protein S18 | 3.0 | 1.2E-18 | 2.3 | 2.1E-03 |
| VC0374 | glucose-6-phosphate isomerase | 2.8 | 2.3E-15 | 3.1 | 9.7E-05 |
| VC0395 | UTP--glucose-1-phosp hate uridylyltransferase | 2.3 | 3.1E-11 | 2.6 | 3.6E-05 |
| VC0478 | fructose-bisphosphat e aldolase, class II | 2.5 | 8.0E-25 | 2.4 | 7.1E-03 |
| VC0483 | conserved hypothetical protein | 2.7 | 1.4E-17 | 2.2 | 9.7E-05 |
| VC0485 | pyruvate kinase I | 2.1 | 9.2E-05 | 2.2 | 5.0E-03 |
| VC0529 | conserved hypothetical protein | 2.5 | 1.4E-09 | 2.2 | 4.8E-05 |
| VC0530 | conserved hypothetical protein | 2.1 | 2.0E-07 | 2.1 | 2.5E-03 |
| VC0564 | ribosomal protein L19 | 5.3 | 1.2E-44 | 2.1 | 4.9E-04 |
| VC0591 | pantoate--beta-alani ne ligase | 2.1 | 2.5E-10 | 2.4 | 3.8E-05 |
| VC0592 | 3-methyl-2-oxobutano atehydroxymethyltransferase | 2.2 | 5.0E-07 | 2.5 | 9.6E-05 |
| VC0596 | dnaK suppressor protein | 3.2 | 9.7E-11 | 2.1 | 3.4E-04 |
| VC0608 | iron(III) ABC transporter, periplasmiciron-compound-binding protein | 9.3 | 9.4E-78 | 3.5 | 1.5E-04 |
| VC0642 | N utilization substance protein A | 2.8 | 2.8E-17 | 2.8 | 1.3E-03 |
| VC0664 | lysyl-tRNA synthetase, heat inducible | 2.6 | 3.1E-14 | 3.2 | 2.1E-09 |
| VC0695 | phospho-2-dehydro-3- deoxyheptonate aldolase,tyr-sensitive | 3.7 | 4.5E-19 | 3.6 | 2.6E-14 |
| VC0718 | conserved hypothetical protein | 2.2 | 2.2E-07 | 2.5 | 9.1E-05 |
| VC0739 | S-adenosylmethionine :tRNAribosyltransferase-isomer ase | 2.3 | 2.4E-09 | 2.3 | 2.6E-04 |
| VC0749 | NifU-related protein | 2.2 | 2.7E-07 | 2.3 | 2.2E-03 |
| VC0754 | conserved hypothetical protein | 2.1 | 3.8E-05 | 2.3 | 2.8E-03 |
| VC0771 | vibriobactin-specifi c isochorismatase | 3.9 | 8.6E-13 | 2.9 | 8.9E-04 |
| VC0847 | integrase, phage family | 2.2 | 1.4E-08 | 2.4 | 1.6E-03 |
| VC0871 | hypothetical protein | 2.3 | 4.2E-06 | 3.2 | 2.6E-07 |
| VC0872 | conserved hypothetical protein | 2.3 | 6.0E-06 | 3.2 | 6.7E-06 |
| VC0905 | lipoprotein YaeC | 3.5 | 2.0E-26 | 2.6 | 9.3E-05 |
| VC0928 | hypothetical protein | 3.1 | 2.2E-14 | 2.7 | 1.4E-04 |
| VC0930 | hemolysin-related protein | 2.7 | 1.1E-14 | 2.4 | 3.4E-07 |
| VC0941 | serine hydroxymethyltransferase | 2.6 | 1.1E-07 | 2.4 | 2.4E-03 |
| VC0945 | conserved hypothetical protein | 2.3 | 2.1E-07 | 2.3 | 2.4E-03 |
| VC0962 | conserved hypothetical protein | 2.5 | 4.3E-10 | 2.1 | 1.6E-04 |
| VC0985 | heat shock protein HtpG | 3.9 | 6.3E-17 | 3.5 | 3.6E-06 |
| VC0991 | asparagine synthetase B, glutamine-hydrolyzing | 3.8 | 1.0E-24 | 3.6 | 2.9E-09 |
| VC1039 | asmA protein | 3.1 | 2.1E-20 | 3.3 | 5.4E-10 |
| VC1040 | cob(I)alamin adenosyltransferase | 2.9 | 2.0E-17 | 2.5 | 2.7E-04 |
| VC1092 | oligopeptide ABC transporter, permease protein | 2.8 | 3.0E-12 | 2.6 | 2.8E-05 |
| VC1093 | oligopeptide ABC transporter, permease protein | 3.0 | 2.7E-13 | 2.7 | 6.7E-06 |
| VC1094 | oligopeptide ABC transporter, ATP-bindingprotein | 2.7 | 2.0E-16 | 2.4 | 4.9E-05 |
| VC1095 | oligopeptide ABC transporter, ATP-bindingprotein | 2.1 | 6.5E-08 | 2.2 | 1.6E-04 |
| VC1159 | phosphoserine aminotransferase | 2.5 | 1.4E-13 | 2.5 | 4.4E-05 |
| VC1193 | hypothetical protein | 4.9 | 1.8E-25 | 2.4 | 3.2E-03 |
| VC1195 | lipoprotein, putative | 4.1 | 9.5E-25 | 2.2 | 8.8E-03 |
| VC1209 | elongation factor P family protein | 3.1 | 2.3E-22 | 2.2 | 3.6E-04 |
| VC1221 | hypothetical protein | 3.2 | 6.8E-24 | 2.5 | 3.9E-05 |
| VC1259 | conserved hypothetical protein | 2.7 | 6.3E-08 | 2.5 | 1.9E-03 |
| VC1280 | hypothetical protein | 2.1 | 2.0E-07 | 2.7 | 6.0E-05 |
| VC1317 | conserved hypothetical protein | 2.9 | 7.7E-18 | 2.8 | 2.9E-08 |
| VC1318 | outer membrane protein OmpV | 4.6 | 2.9E-17 | 5.0 | 8.8E-03 |
| VC1374 | DnaK-related protein | 3.1 | 6.9E-14 | 2.1 | 7.5E-04 |
| VC1375 | hypothetical protein | 2.9 | 4.1E-12 | 2.5 | 2.2E-04 |
| VC1410 | multidrug resistance protein VceA | 2.4 | 2.8E-09 | 2.8 | 7.9E-08 |
| VC1414 | thermostable carboxypeptidase 1 | 2.4 | 1.0E-06 | 2.6 | 4.4E-04 |
| VC1415 | hcp protein | 9.5 | 9.4E-47 | 7.4 | 6.4E-17 |
| VC1416 | vgrG protein | 4.9 | 5.4E-45 | 3.7 | 2.9E-08 |
| VC1417 | hypothetical protein | 3.2 | 1.2E-15 | 2.6 | 3.2E-04 |
| VC1418 | hypothetical protein | 2.9 | 3.0E-15 | 2.4 | 3.1E-04 |
| VC1419 | hypothetical protein | 2.5 | 1.1E-10 | 2.2 | 1.8E-03 |
| VC1563 | conserved hypothetical protein | 2.1 | 5.2E-06 | 2.2 | 3.5E-03 |
| VC1565 | outer membrane protein TolC, putative | 2.5 | 1.0E-09 | 2.4 | 2.3E-03 |
| VC1578 | hypothetical protein | 2.7 | 1.2E-09 | 2.2 | 4.6E-03 |
| VC1623 | carboxynorspermidine decarboxylase | 3.3 | 1.2E-22 | 2.8 | 1.9E-07 |
| VC1624 | conserved hypothetical protein | 2.0 | 2.3E-06 | 2.1 | 9.7E-05 |

| | | | | | |
|---|---|---|---|---|---|
| VC1625 | Pseudogene; involved in spermidine biosynthesis | 3.6 | 1.3E-23 | 2.7 | 1.8E-07 |
| VC1655 | magnesium transporter | 3.1 | 5.3E-17 | 2.3 | 3.7E-05 |
| VC1738 | hypothetical protein | 2.5 | 7.1E-07 | 2.5 | 5.0E-03 |
| VC1750 | hypothetical protein | 2.1 | 4.4E-08 | 2.7 | 2.9E-03 |
| VC1820 | PTS system, fructose-specific IIA component | 54.5 | 5.6E-21 | 112.3 | 7.7E-16 |
| VC1821 | PTS system, fructose-specific IIBC component | 2.5 | 2.9E-09 | 13.6 | 1.5E-10 |
| VC1822 | PTS system, fructose-specific IIABC component | 4.4 | 6.7E-24 | 4.3 | 2.0E-10 |
| VC1823 | PTS system, fructose-specific IIB component | 4.9 | 1.6E-25 | 6.5 | 5.8E-21 |
| VC1825 | transcriptional regulator | 3.5 | 7.1E-13 | 4.7 | 4.0E-13 |
| VC1826 | PTS system, fructose-specific IIABC component | 3.0 | 1.4E-09 | 41.9 | 5.3E-14 |
| VC1849 | peptidyl-prolyl cis-trans isomerase B | 2.7 | 2.1E-15 | 2.4 | 1.8E-05 |
| VC1853 | conserved hypothetical protein | 2.8 | 2.1E-18 | 2.3 | 3.4E-04 |
| VC1883 | ABC transporter, ATP-binding protein | 2.0 | 1.6E-07 | 2.2 | 1.1E-04 |
| VC1888 | hemolysin-related protein | 2.9 | 6.1E-24 | 2.2 | 1.1E-04 |
| VC1962 | lipoprotein | 5.6 | 8.6E-55 | 3.6 | 1.3E-10 |
| VC2022 | malonyl Coa-acyl carrier protein transacylase | 3.2 | 1.9E-14 | 3.4 | 1.7E-06 |
| VC2053 | cytochrome c-type biogenesis protein CcmE | 2.2 | 5.6E-08 | 2.3 | 8.8E-04 |
| VC2213 | outer membrane protein OmpA | 7.0 | 4.5E-33 | 5.5 | 6.5E-03 |
| VC2223 | pseudouridine synthase family 1 protein | 2.2 | 6.1E-17 | 2.1 | 2.5E-03 |
| VC2259 | elongation factor Ts | 4.6 | 1.5E-46 | 3.9 | 1.2E-04 |
| VC2329 | 2,3,4,5-tetrahydropy ridine-2-carboxylateN-succinyl transferase | 2.3 | 1.2E-07 | 2.1 | 6.5E-04 |
| VC2386 | conserved hypothetical protein | 2.8 | 7.8E-16 | 2.2 | 7.0E-05 |
| VC2389 | carbamoyl-phosphate synthase, large subunit | 2.2 | 1.8E-11 | 3.2 | 6.8E-10 |
| VC2400 | UDP-N-acetylmuramate --alanine ligase | 2.3 | 1.1E-12 | 2.0 | 3.6E-04 |
| VC2414 | pyruvate dehydrogenase, E1 component | 2.5 | 1.6E-13 | 2.6 | 3.7E-04 |
| VC2485 | transcriptional regulator, LysR family | 2.2 | 5.9E-15 | 2.9 | 7.4E-09 |
| VC2503 | valyl-tRNA synthetase | 2.4 | 9.2E-07 | 2.2 | 9.4E-03 |
| VC2512 | conserved hypothetical protein | 4.1 | 4.0E-28 | 2.1 | 6.1E-04 |
| VC2568 | peptidyl-prolyl cis-trans isomerase, FKBP-type | 3.5 | 2.7E-22 | 3.1 | 4.2E-06 |
| VC2578 | ribosomal protein L30 | 2.4 | 1.4E-06 | 2.1 | 1.6E-03 |
| VC2582 | ribosomal protein S8 | 2.8 | 1.4E-14 | 2.6 | 1.7E-03 |
| VC2583 | ribosomal protein S14 | 2.6 | 4.0E-13 | 2.3 | 8.2E-03 |
| VC2587 | ribosomal protein S17 | 2.6 | 4.6E-13 | 2.5 | 6.5E-06 |
| VC2588 | ribosomal protein L29 | 2.5 | 4.2E-07 | 2.3 | 5.5E-03 |
| VC2591 | ribosomal protein L22 | 2.7 | 1.9E-13 | 2.7 | 8.2E-04 |
| VC2592 | ribosomal protein S19 | 2.6 | 9.3E-14 | 2.5 | 1.3E-03 |
| VC2594 | ribosomal protein L23 | 4.0 | 1.3E-14 | 3.2 | 2.1E-07 |
| VC2595 | ribosomal protein L4 | 3.4 | 2.0E-22 | 2.4 | 4.4E-03 |
| VC2646 | phosphoenolpyruvate carboxylase | 2.1 | 5.2E-07 | 2.4 | 1.0E-05 |
| VC2662 | conserved hypothetical protein | 5.2 | 1.1E-37 | 6.0 | 3.2E-09 |
| VC2664 | chaperonin, 60 Kd subunit, groES_1 | 3.4 | 1.2E-11 | 3.0 | 1.7E-04 |
| VC2670 | triosephosphate isomerase | 3.5 | 3.8E-23 | 3.5 | 8.7E-05 |
| VC2679 | ribosomal protein L31 | 5.9 | 1.6E-47 | 2.2 | 1.3E-03 |
| VC2691 | periplasmic protein cpxP, putative | 2.8 | 1.1E-09 | 4.5 | 1.5E-04 |
| VC2706 | conserved hypothetical protein | 4.0 | 7.1E-18 | 2.1 | 8.1E-04 |
| VC2715 | transcription elongation factor GreB | 2.3 | 5.7E-06 | 3.0 | 3.3E-05 |
| VC2716 | conserved hypothetical protein | 3.0 | 5.4E-33 | 2.6 | 3.6E-07 |
| VC2761 | multidrug resistance protein | 2.4 | 3.7E-11 | 3.8 | 1.7E-06 |
| VC2762 | UDP-N-acetylglucosam ine pyrophosphorylase | 3.2 | 5.3E-20 | 2.3 | 4.3E-04 |
| VC2774 | glucose inhibited division protein B | 2.4 | 1.4E-11 | 2.0 | 4.5E-05 |
| VCA0017 | hcp-2 protein | 12.0 | 4.0E-55 | 8.6 | 3.9E-18 |
| VCA0018 | vgrG protein | 3.2 | 1.6E-14 | 2.8 | 3.4E-04 |
| VCA0019 | hypothetical protein | 3.5 | 9.1E-13 | 3.8 | 4.0E-06 |
| VCA0020 | hypothetical protein | 3.0 | 1.1E-32 | 3.0 | 1.4E-07 |
| VCA0026 | conserved hypothetical protein | 3.5 | 1.2E-21 | 2.4 | 2.2E-06 |
| VCA0035 | phosphatidylglycerop hosphatase B, putative | 3.2 | 6.4E-18 | 5.0 | 5.6E-09 |
| VCA0087 | hypothetical protein | 2.7 | 3.2E-05 | 2.6 | 3.4E-04 |
| VCA0088 | proton/glutamate symporter | 6.2 | 2.8E-81 | 3.5 | 3.1E-14 |
| VCA0107 | conserved hypothetical protein | 6.1 | 3.8E-31 | 4.3 | 1.1E-08 |
| VCA0108 | conserved hypothetical protein | 5.8 | 7.3E-44 | 4.5 | 5.8E-17 |
| VCA0109 | hypothetical protein | 2.6 | 2.0E-09 | 2.3 | 2.0E-03 |
| VCA0112 | hypothetical protein | 2.1 | 3.5E-08 | 3.0 | 2.1E-05 |
| VCA0136 | glycerophosphoryl diester phosphodiesterase | 11.6 | 7.0E-41 | 18.5 | 2.2E-29 |
| VCA0137 | glycerol-3-phosphate transporter | 10.5 | 4.1E-71 | 14.5 | 2.0E-20 |
| VCA0139 | hypothetical protein | 6.2 | 6.9E-37 | 4.3 | 6.8E-05 |
| VCA0192 | D-lactate dehydrogenase | 3.6 | 2.0E-24 | 3.3 | 9.2E-10 |
| VCA0223 | protease | 3.1 | 3.1E-07 | 2.8 | 1.5E-04 |
| VCA0227 | iron(III) ABC transporter, periplasmiciron-compound-binding protein | 19.3 | 1.3E-85 | 5.6 | 4.8E-04 |
| VCA0277 | glycine cleavage system H protein | 2.2 | 6.6E-09 | 2.0 | 1.2E-03 |
| VCA0517 | 1-phosphofructokinas e | 2.5 | 1.3E-07 | 3.8 | 1.6E-06 |
| VCA0518 | PTS system, fructose-specific IIA/FPR component | 6.5 | 2.8E-27 | 10.2 | 2.4E-22 |
| VCA0519 | fructose repressor | 2.7 | 1.6E-09 | 3.8 | 1.0E-09 |

| | | | | | |
|---|---|---|---|---|---|
| VCA0540 | formate transporter 1, putative | 2.4 | 9.4E-06 | 13.3 | 3.7E-14 |
| VCA0563 | NAD(P) transhydrogenase, alpha subunit | 3.4 | 7.8E-12 | 3.5 | 1.8E-05 |
| VCA0564 | NAD(P) transhydrogenase, beta subunit | 3.7 | 2.7E-26 | 3.6 | 7.6E-10 |
| VCA0652 | hypothetical protein | 3.7 | 4.3E-18 | 2.7 | 5.5E-07 |
| VCA0657 | aerobic glycerol-3-phosphate dehydrogenase | 55.0 | 1.4E-155 | 70.9 | 5.3E-46 |
| VCA0692 | pseudogene with authentic frameshift mutation | 2.4 | 7.7E-07 | 2.5 | 1.8E-03 |
| VCA0721 | hypothetical protein | 4.3 | 1.5E-19 | 3.3 | 3.3E-03 |
| VCA0744 | glycerol kinase | 12.1 | 6.8E-67 | 15.6 | 4.7E-19 |
| VCA0745 | pseudogene with authentic frameshift mutation | 12.7 | 1.5E-79 | 14.6 | 7.6E-33 |
| VCA0747 | anaerobic glycerol-3-phosphate dehydrogenase,subunit A | 22.0 | 5.1E-116 | 43.5 | 1.6E-34 |
| VCA0748 | anaerobic glycerol-3-phosphate dehydrogenase,subunit B | 11.9 | 4.3E-69 | 28.0 | 4.2E-34 |
| VCA0749 | anaerobic glycerol-3-phosphate dehydrogenase,subunit C | 11.9 | 6.9E-59 | 33.0 | 4.0E-45 |
| VCA0789 | conserved hypothetical protein | 3.4 | 2.6E-18 | 3.4 | 5.3E-07 |
| VCA0811 | chitinase, putative | 3.0 | 4.8E-15 | 3.2 | 2.8E-08 |
| VCA0819 | chaperonin, 10 Kd subunit | 3.4 | 2.3E-14 | 2.7 | 1.2E-05 |
| VCA0820 | chaperonin, 60 Kd subunit | 2.4 | 2.0E-10 | 2.2 | 1.9E-03 |
| VCA0862 | long-chain fatty acid transport protein | 3.3 | 2.4E-14 | 3.6 | 2.8E-04 |
| VCA0863 | lipase, putative | 4.6 | 5.9E-32 | 3.4 | 1.7E-07 |
| VCA0897 | devB protein | 2.7 | 1.2E-12 | 2.2 | 2.6E-03 |
| VCA0898 | 6-phosphogluconate dehydrogenase,decarboxylating | 3.2 | 2.3E-15 | 3.2 | 6.9E-09 |
| VCA0983 | L-lactate permease, putative | 6.9 | 2.1E-56 | 3.1 | 6.9E-08 |
| VCA0984 | L-lactate dehydrogenase | 11.2 | 1.7E-94 | 9.2 | 4.6E-25 |
| VCA0985 | oxidoreductase/iron- sulfur cluster-bindingprotein | 4.5 | 5.8E-24 | 4.0 | 2.8E-10 |
| VCA0987 | phosphoenolpyruvate synthase | 2.4 | 3.4E-14 | 2.9 | 3.4E-07 |
| VCA1035 | hypothetical protein | 3.9 | 1.3E-16 | 2.5 | 3.0E-04 |
| VCA1073 | bifunctional proline dehydrogenase/pyrroline-5-carboxylate dehydrogenase | 4.5 | 1.5E-29 | 4.1 | 2.3E-06 |
