## Supplementary material for "Modular small RNA drives pathogen emergence": Table S6

**Table S6:** Gene expression in WT and Δ*toxR* -pOueS strains from biofilm growth. Absolute fold changes (> 2.0) are relative to expression in the Δ*toxR* strain.

| Name | Function | WT/Δ*toxR* | | Δ*toxR* -pOueS/Δ*toxR* | |
|---|---|---|---|---|---|
| | | FC | FDR *p* | FC | FDR *p* |
| 16Sc | ribosomal RNA | 2.1 | 2.31E-06 | -2.1 | 2.34E-09 |
| 16Sd | ribosomal RNA | -8.4 | 4.87E-39 | -2.3 | 9.29E-09 |
| 16Se | ribosomal RNA | 2.1 | 2.44E-06 | -2.1 | 1.50E-08 |
| 16Sh | ribosomal RNA | -10.2 | 2.89E-50 | -2.3 | 2.02E-08 |
| 23Sa | ribosomal RNA | -3.0 | 8.29E-14 | -2.3 | 1.68E-06 |
| 23Sc | ribosomal RNA | -98905.7 | 3.26E-37 | -5.3 | 6.61E-11 |
| 23Sd | ribosomal RNA | 2.2 | 1.07E-07 | -2.3 | 9.75E-09 |
| 5Sf | ribosomal RNA | -6.9 | 1.89E-16 | -2.0 | 1.09E-11 |
| VC0063 | thiF protein | -3.0 | 1.38E-06 | -3.6 | 2.14E-30 |
| VC0065 | thiG protein | -2.5 | 3.18E-05 | -3.0 | 3.46E-17 |
| VC0066 | thiH protein | -2.7 | 1.69E-15 | -3.7 | 3.10E-34 |
| VC0107 | hypothetical protein | -8.3 | 3.93E-43 | -13.3 | 5.10E-100 |
| VC0134 | conserved hypothetical protein | -2.1 | 1.10E-11 | -2.0 | 4.52E-15 |
| VC0199 | hemolysin secretion ATP-binding protein,putative | -2.7 | 1.08E-13 | -3.2 | 2.81E-18 |
| VC0200 | iron(III) compound receptor | -3.3 | 1.38E-11 | -3.6 | 5.50E-39 |
| VC0201 | iron(III) ABC transporter, ATP-binding protein | -2.8 | 1.05E-09 | -2.8 | 9.67E-07 |
| VC0202 | iron(III) ABC transporter, periplasmiciron-compound-bindi ng protein | -2.4 | 3.35E-06 | -4.1 | 9.48E-13 |
| VC0284 | putative outer membrane receptor | -2.7 | 1.68E-10 | -2.6 | 8.14E-20 |
| VC0364 | bacterioferritin-ass ociated ferredoxin | -3.2 | 4.03E-21 | -2.4 | 2.88E-22 |
| VC0608 | iron(III) ABC transporter, periplasmiciron-compound-bindi ng protein | -2.8 | 7.44E-10 | -2.2 | 5.78E-21 |
| VC0676 | nptA protein | -6.6 | 1.26E-28 | -2.4 | 2.20E-23 |
| VC0771 | vibriobactin-specific isochorismatase | -4.1 | 6.81E-11 | -3.6 | 9.11E-33 |
| VC0774 | vibriobactin-specific2,3-dihydro-2,3-dihydroxybenzoate dehydrogenase | -3.0 | 5.43E-09 | -4.3 | 5.39E-13 |
| VC0775 | vibriobactin synthesis protein, putative | -2.4 | 2.38E-07 | -3.0 | 5.25E-14 |
| VC0776 | ferric vibriobactin ABC transporter, periplasmicferric vibriobactin-binding protein | -2.9 | 1.70E-07 | -2.5 | 6.46E-12 |
| VC0777 | ferric vibriobactin ABC transporter, permeaseprotein | -2.6 | 1.10E-07 | -2.3 | 1.38E-05 |
| VC0778 | ferric vibriobactin ABC transporter, permeaseprotein | -3.0 | 1.16E-10 | -3.4 | 1.48E-09 |
| VC0916 | phosphotyrosine protein phosphatase | -4.4 | 4.17E-09 | -2.7 | 1.49E-29 |
| VC0917 | UDP-N-acetylglucosam ine 2-epimerase | -2.7 | 3.40E-07 | -2.3 | 3.26E-16 |
| VC0918 | UDP-N-acetyl-D-manno saminuronic aciddehydrogenase | -2.2 | 1.12E-05 | -2.3 | 9.31E-21 |
| VC0919 | serine acetyltransferase-related protein | -2.6 | 9.95E-08 | -2.7 | 5.62E-23 |
| VC0920 | exopolysaccharide biosynthesis protein EpsF,putative | -2.6 | 7.67E-11 | -2.2 | 3.78E-16 |
| VC0927 | UDP-N-acetyl-D-manno samine transferase | -2.1 | 1.22E-05 | -2.6 | 1.06E-21 |
| VC0930 | hemolysin-related protein | -2.7 | 2.58E-04 | -4.8 | 2.66E-75 |
| VC0931 | conserved hypothetical protein | -3.3 | 5.82E-13 | -3.6 | 2.63E-34 |
| VC0932 | hypothetical protein | -5.3 | 7.96E-10 | -6.3 | 1.33E-95 |
| VC0934 | capsular polysaccharide biosynthesisglycosyltransferas e, putative | -4.2 | 6.89E-06 | -3.8 | 5.31E-52 |
| VC0935 | hypothetical protein | -4.3 | 1.23E-08 | -3.6 | 1.02E-49 |
| VC0936 | polysaccharide export-related protein | -3.5 | 8.37E-09 | -3.0 | 5.41E-33 |
| VC0937 | exopolysaccharide biosynthesis protein,putative | -2.9 | 3.77E-09 | -3.2 | 4.42E-80 |
| VC0938 | hypothetical protein | -3.6 | 1.45E-17 | -3.6 | 1.66E-17 |
| VC0940 | conserved hypothetical protein | -2.0 | 2.64E-07 | -2.0 | 1.25E-13 |
| VC1121 | conserved hypothetical protein | -2.8 | 1.05E-14 | -2.2 | 3.83E-19 |
| VC1168 | proton/glutamate symporter | -2.6 | 1.03E-08 | -2.1 | 1.32E-16 |
| VC1264 | iron-regulated protein A, putative | -2.3 | 2.42E-12 | -2.2 | 2.02E-18 |
| VC1265 | hypothetical protein | -2.5 | 4.12E-10 | -2.8 | 7.72E-21 |
| VC1266 | hypothetical protein | -2.1 | 4.43E-06 | -2.9 | 1.68E-29 |
| VC1267 | hypothetical protein | -2.2 | 9.97E-09 | -2.7 | 1.88E-23 |
| VC1312 | alanine racemase, putative | -3.1 | 5.02E-15 | -2.9 | 1.16E-25 |
| VC1313 | methyl-accepting chemotaxis protein | -3.0 | 1.00E-14 | -2.4 | 2.01E-23 |
| VC1511 | formate dehydrogenase, cytochrome B556 subunit | -2.4 | 2.18E-11 | -3.4 | 2.59E-44 |
| VC1518 | hypothetical protein | -2.2 | 3.95E-07 | -2.4 | 2.92E-12 |
| VC1543 | hypothetical protein | -2.6 | 6.98E-15 | -2.8 | 2.82E-28 |
| VC1544 | tonB2 protein | -2.9 | 1.71E-13 | -2.6 | 1.49E-18 |
| VC1545 | TonB system transport protein ExbD2 | -2.2 | 5.34E-07 | -2.9 | 1.21E-10 |
| VC1546 | TonB system transport protein ExbB2 | -3.0 | 4.86E-11 | -2.7 | 5.02E-22 |
| VC1547 | biopolymer transport protein ExbB-relatedprotein | -2.4 | 7.45E-10 | -2.6 | 9.28E-24 |
| VC1570 | quinol oxidase, subunit II | -4.9 | 3.40E-30 | -4.7 | 1.57E-58 |
| VC1571 | quinol oxidase, subunit I | -4.0 | 3.94E-25 | -3.6 | 2.15E-87 |
| VC1572 | hypothetical protein | -2.1 | 9.38E-05 | -2.6 | 1.12E-06 |
| VC1573 | fumarate hydratase, class II | -4.2 | 5.03E-25 | -4.6 | 1.83E-23 |
| VC1633 | hypothetical protein | -2.6 | 3.35E-11 | -2.2 | 1.35E-19 |
| VC1688 | hypothetical protein | -3.3 | 3.12E-12 | -4.0 | 9.82E-11 |
| VC1819 | aldehyde dehydrogenase | -5.6 | 6.54E-38 | -2.6 | 8.33E-28 |
| VC1888 | hemolysin-related protein | -2.4 | 5.77E-08 | -2.2 | 1.75E-19 |

| | | | | | |
|---|---|---|---|---|---|
| VC2035 | conserved hypothetical protein | -2.5 | 3.56E-14 | -2.1 | 1.00E-11 |
| VC2044 | conserved hypothetical protein | -2.8 | 2.04E-16 | -2.1 | 6.50E-38 |
| VC2209 | nonribosomal peptide synthetase VibF | -3.1 | 8.69E-23 | -3.2 | 9.82E-37 |
| VC2210 | vibriobactin utilization protein ViuB | -4.5 | 1.72E-27 | -3.5 | 1.74E-37 |
| VC2211 | ferric vibriobactin receptor | -7.7 | 1.34E-60 | -8.0 | 9.58E-112 |
| VC2416 | 2`,3`-cyclic-nucleot ide 2`-phosphodiesterase,putative | -2.5 | 4.50E-09 | -2.5 | 4.86E-26 |
| VC2537 | thiamine ABC transporter, ATP-binding protein,putative | -2.1 | 9.27E-04 | -2.3 | 2.08E-12 |
| VC2667 | hypothetical protein | -3.2 | 1.01E-20 | -3.2 | 9.63E-29 |
| VC2694 | superoxide dismutase, Mn | -3.2 | 9.25E-24 | -3.2 | 6.02E-28 |
| VC2705 | sodium/solute symporter, putative | -2.6 | 8.35E-14 | -2.0 | 1.73E-16 |
| VCA0063 | protease II | -3.2 | 1.84E-12 | -2.6 | 1.66E-15 |
| VCA0064 | TonB system receptor, putative | -2.3 | 3.54E-04 | -2.5 | 3.58E-17 |
| VCA0068 | methyl-accepting chemotaxis protein | -2.5 | 2.08E-09 | -2.2 | 4.20E-13 |
| VCA0083 | multidrug resistance protein D | -3.3 | 2.78E-15 | -3.4 | 6.65E-42 |
| VCA0084 | soxR protein | -2.1 | 5.41E-08 | -2.3 | 7.51E-17 |
| VCA0127 | ribose ABC transporter protein | -2.1 | 2.43E-06 | -2.4 | 1.04E-05 |
| VCA0146 | conserved hypothetical protein | -4.3 | 1.88E-13 | -3.8 | 2.84E-13 |
| VCA0147 | transcriptional regulator, putative | -6.5 | 5.61E-35 | -6.5 | 3.49E-67 |
| VCA0160 | tryptophan-specific transport protein | -3.4 | 3.11E-18 | -4.6 | 6.22E-54 |
| VCA0161 | tryptophanase | -2.8 | 3.70E-17 | -2.7 | 1.72E-16 |
| VCA0184 | cold shock DNA-binding domain protein | -2.7 | 5.40E-11 | -2.2 | 1.14E-09 |
| VCA0218 | thermolabile hemolysin | -3.9 | 7.48E-25 | -2.2 | 1.88E-17 |
| VCA0227 | iron(III) ABC transporter, periplasmiciron-compound-binding protein | -3.1 | 8.33E-15 | -2.7 | 7.59E-59 |
| VCA0230 | iron(III) ABC transporter, ATP-binding protein | -2.9 | 2.11E-14 | -2.8 | 1.54E-22 |
| VCA0231 | transcriptional regulator, AraC/XylS family | -4.2 | 9.42E-11 | -2.9 | 8.26E-15 |
| VCA0232 | enterobactin receptor, VctA | -6.6 | 2.53E-50 | -11.2 | 1.64E-141 |
| VCA0233 | hypothetical protein | -5.0 | 1.38E-16 | -6.7 | 2.89E-16 |
| VCA0276 | Glycine cleavage system P protein GcvP, authentic frameshift | -3.5 | 1.02E-21 | -3.0 | 4.04E-79 |
| VCA0277 | glycine cleavage system H protein | -2.7 | 1.77E-15 | -2.7 | 1.98E-30 |
| VCA0576 | heme transport protein HutA | -11.6 | 1.45E-29 | -11.6 | 1.35E-103 |
| VCA0875 | D-serine dehydratase | -2.7 | 1.42E-13 | -2.9 | 2.81E-29 |
| VCA0885 | threonine 3-dehydrogenase | -2.6 | 2.36E-13 | -3.8 | 1.39E-49 |
| VCA0886 | 2-amino-3-ketobutyra te coenzyme A ligase | -2.8 | 2.84E-14 | -3.3 | 5.42E-40 |
| VCA0907 | conserved hypothetical protein | -8.7 | 1.51E-48 | -12.3 | 8.12E-116 |
| VCA0908 | conserved hypothetical protein | -8.6 | 3.03E-43 | -8.9 | 7.01E-127 |
| VCA0909 | oxygen-independent coproporphyrinogen IIIoxidase, putative | -7.4 | 3.15E-28 | -8.0 | 3.47E-115 |
| VCA0910 | tonB1 protein | -6.2 | 2.08E-19 | -5.9 | 2.91E-51 |
| VCA0911 | TonB system transport protein ExbB1 | -8.6 | 3.85E-19 | -9.6 | 7.44E-74 |
| VCA0912 | TonB system transport protein ExbD1 | -8.9 | 4.64E-22 | -10.2 | 6.39E-77 |
| VCA0913 | hemin ABC transporter, periplasmic hemin-bindingprotein HutB | -6.7 | 2.43E-39 | -14.4 | 1.26E-109 |
| VCA0914 | hemin ABC transporter, permease protein,putative | -7.4 | 4.08E-39 | -10.2 | 1.68E-93 |
| VCA0915 | hemin ABC transporter, ATP-binding protein HutD | -2.8 | 3.78E-18 | -2.6 | 2.19E-17 |
| VCA0928 | hypothetical protein | -3.0 | 1.62E-08 | -3.8 | 2.12E-14 |
| VCA0952 | transcriptional regulator, LuxR family | -3.0 | 1.26E-04 | -3.7 | 2.27E-51 |
| VCA0976 | hypothetical protein | -3.6 | 6.71E-12 | -3.0 | 2.93E-17 |
| VCA0977 | ABC transporter, ATP-binding protein | -3.7 | 1.74E-21 | -4.0 | 1.02E-51 |
| VCA1070 | hypothetical protein | -2.6 | 2.32E-09 | -2.9 | 3.67E-26 |
| 16Sc | ribosomal RNA | 2.1 | 2.31E-06 | -2.1 | 2.34E-09 |
| 16Sd | ribosomal RNA | -8.4 | 4.87E-39 | -2.3 | 9.29E-09 |
| 16Se | ribosomal RNA | 2.1 | 2.44E-06 | -2.1 | 1.50E-08 |
| 16Sh | ribosomal RNA | -10.2 | 2.89E-50 | -2.3 | 2.02E-08 |
| 23Sa | ribosomal RNA | -3.0 | 8.29E-14 | -2.3 | 1.68E-06 |
| 23Sc | ribosomal RNA | -98905.7 | 3.26E-37 | -5.3 | 6.61E-11 |
| 23Sd | ribosomal RNA | 2.2 | 1.07E-07 | -2.3 | 9.75E-09 |
| 5Sf | ribosomal RNA | -6.9 | 1.89E-16 | -2.0 | 1.09E-11 |
| VC0076 | universal stress protein A | 4.9 | 5.07E-35 | 2.5 | 2.71E-25 |
| VC0143 | hypothetical protein | 2.7 | 1.61E-08 | 2.0 | 7.25E-04 |
| VC0188 | oligopeptidase A | 3.0 | 8.41E-12 | 2.1 | 1.09E-33 |
| VC0534 | RNA polymerase sigma-38 factor | 2.2 | 1.01E-08 | 2.4 | 4.27E-52 |
| VC0706 | sigma-54 modulation protein, putative | 2.6 | 1.40E-08 | 2.0 | 4.22E-16 |
| VC0885 | hypothetical protein | 2.4 | 4.57E-07 | 2.1 | 1.83E-05 |
| VC0886 | hypothetical protein | 2.5 | 9.44E-13 | 2.2 | 5.53E-16 |
| VC0977 | conserved hypothetical protein | 2.1 | 9.55E-09 | 2.1 | 3.69E-09 |
| VC0985 | heat shock protein HtpG | 3.4 | 4.82E-23 | 2.2 | 9.06E-37 |
| VC1080 | hypothetical protein | 3.2 | 6.57E-13 | 3.7 | 7.64E-30 |
| VC1081 | response regulator | 3.0 | 2.87E-20 | 2.9 | 1.46E-29 |
| VC1249 | conserved hypothetical protein | 3.5 | 2.32E-16 | 2.0 | 2.87E-15 |
| VC1318 | outer membrane protein OmpV | -3.5 | 1.05E-15 | 2.4 | 1.08E-22 |
| VC1325 | galactoside ABC transporter, periplasmicD-galactose/D-gluco se-binding protein | -2.3 | 2.26E-10 | 2.1 | 5.57E-18 |

| | | | | | |
|---|---|---|---|---|---|
| VC1403 | methyl-accepting chemotaxis protein | 3.0 | 1.93E-16 | 2.1 | 3.71E-11 |
| VC1472 | hypothetical protein | 3.3 | 1.13E-04 | 4.2 | 7.48E-07 |
| VC1871 | conserved hypothetical protein | 3.4 | 6.10E-16 | 2.1 | 1.33E-10 |
| VC2361 | formate acetyl transferase-related protein | 4.4 | 9.16E-11 | 2.3 | 1.84E-09 |
| VC2643 | acetylglutamate kinase | 2.7 | 1.31E-09 | 2.1 | 8.87E-06 |
| VC2675 | protease HslVU, subunit HslV | 2.2 | 8.34E-08 | 2.3 | 6.61E-22 |
| VCA0139 | hypothetical protein | -2.1 | 8.08E-08 | 2.7 | 9.16E-27 |
| VCA0328 | biphenyl-2,3-diol 1,2-dioxygenase III-relatedprotein | 2.3 | 5.20E-05 | 2.2 | 1.15E-10 |
| VCA0343 | hypothetical protein | 3.0 | 1.84E-06 | 3.2 | 6.55E-06 |
| VCA0345 | conserved hypothetical protein | 2.0 | 6.02E-14 | 2.2 | 2.11E-18 |
| VCA0346 | H-REV 107-related protein | 3.5 | 4.83E-22 | 2.3 | 6.48E-19 |
| VCA0357 | hypothetical protein | 3.3 | 1.14E-05 | 2.7 | 2.07E-04 |
| VCA0420 | hypothetical protein | 2.0 | 1.17E-03 | 2.1 | 3.41E-10 |
| VCA0446 | haemagglutinin | 4.3 | 5.21E-25 | 2.2 | 1.11E-10 |
| VCA0449 | hypothetical protein | 2.3 | 5.24E-04 | 2.0 | 4.43E-04 |
| VCA0451 | hypothetical protein | 2.8 | 5.51E-05 | 2.2 | 3.12E-04 |
| VCA0507 | transposase OrfAB, subunit A | 2.2 | 5.18E-10 | 2.0 | 1.83E-14 |
| VCA0508 | transposase OrfAB, subunit B | 2.9 | 6.44E-14 | 4.0 | 6.40E-24 |
| VCA0688 | polyhydroxyalkanoic acid synthase | 3.3 | 6.14E-21 | 2.2 | 1.35E-19 |
| VCA0689 | conserved hypothetical protein | 3.4 | 1.67E-14 | 2.1 | 2.79E-11 |
| VCA0690 | acetyl-CoA acetyltransferase | 5.0 | 6.79E-38 | 2.4 | 4.14E-23 |
| VCA0720 | guanylate cyclase-related protein | 3.0 | 1.18E-08 | 2.8 | 3.72E-08 |
| VCA0743 | conserved hypothetical protein | 2.2 | 4.38E-05 | 2.5 | 9.66E-08 |
| VCA0749 | anaerobic glycerol-3-phosphate dehydrogenase,subunit C | 2.4 | 1.96E-09 | 3.6 | 3.21E-23 |
| VCA0880 | hypothetical protein | 2.5 | 1.10E-07 | 3.8 | 2.17E-24 |
| VCA0881 | hypothetical protein | 4.2 | 1.68E-16 | 4.0 | 7.38E-51 |
| VCA0882 | hypothetical protein | 3.3 | 4.62E-10 | 3.0 | 2.16E-75 |
| VCA0981 | hypothetical protein | 3.5 | 7.08E-21 | 3.0 | 5.67E-32 |
| VCr025 | ribosomal RNA 5S | 6.3 | 4.34E-18 | 2.0 | 5.60E-04 |
