## Supplementary material for "Modular small RNA drives pathogen emergence": Table S8

**Table S8**. Overlap between the ToxR and OueS virulence regulons. Subset of the ToxR AKI regulon that comprises the OueS regulated genes (yellow highlight) identified by an absolute ratio of > 2.0 between the two datasets. Expression values are relative to the WT strain.

| **Locus tag** | **Function** | **Δ*toxR* / WT** | | **Δ*toxR*-pOueS / WT** | | **Ratio: *toxR* / *toxR*-pOueS** |
|---|---|---|---|---|---|---|
| | | **FC** | **FDR *p*-value** | **FC** | **FDR *p*-value** | |
| VC0032 | ComM-related protein | -2.8 | 1.01E-06 | 1.1 | 0.72 | -2.6 |
| VC0069 | multidrug resistance protein, putative | -2.5 | 3.34E-12 | 2.4 | 0.00 | -1.0 |
| VC0078 | ferritin | -2.4 | 3.28E-04 | 1.2 | 0.17 | -1.9 |
| VC0079 | conserved hypothetical protein | -3.0 | 4.18E-20 | 1.4 | 0.01 | -2.2 |
| VC0089 | cytochrome c551 peroxidase | -9.4 | 6.86E-48 | 1.3 | 0.09 | -7.3 |
| VC0090 | DNA-damage-inducible protein F | -3.0 | 7.64E-17 | 1.1 | 0.48 | -2.7 |
| VC0096 | hypothetical protein | -2.4 | 3.15E-06 | 1.0 | 0.91 | -2.4 |
| VC0098 | methyl-accepting chemotaxis protein | -2.8 | 2.47E-20 | 1.2 | 0.04 | -2.3 |
| VC0121 | hypothetical protein | -2.7 | 4.50E-07 | 1.1 | 0.38 | -2.4 |
| VC0139 | DPS family protein | -2.6 | 7.28E-16 | 1.3 | 0.06 | -2.1 |
| VC0142 | hypothetical protein | -2.1 | 1.06E-16 | -1.0 | 0.74 | 2.0 |
| VC0169 | hypothetical protein | -3.7 | 4.57E-12 | 1.6 | 0.00 | -2.3 |
| VC0170 | peptide ABC transporter, ATP-binding protein | -2.7 | 2.51E-14 | 1.2 | 0.18 | -2.3 |
| VC0171 | peptide ABC transporter, periplasmicpeptide-binding protein | -2.8 | 1.26E-13 | 1.1 | 0.60 | -2.7 |
| VC0172 | peptide ABC transporter, permease protein | -2.4 | 4.99E-11 | 1.2 | 0.05 | -2.0 |
| VC0298 | acetyl-CoA synthase | -11.9 | 5.93E-28 | 1.2 | 0.20 | -9.9 |
| VC0299 | DNA polymerase III, epsilon subunit, putative | -4.8 | 8.30E-24 | 1.4 | 0.01 | -3.5 |
| VC0300 | conserved hypothetical protein | -7.2 | 3.47E-27 | 1.5 | 0.01 | -4.8 |
| VC0365 | bacterioferritin | -2.6 | 5.10E-13 | 1.6 | 0.00 | -1.6 |
| VC0383 | hypothetical protein | -2.3 | 6.21E-04 | 1.6 | 0.01 | -1.5 |
| VC0392 | aminotransferase, class V | -2.1 | 2.89E-16 | 1.0 | 0.71 | -2.0 |
| VC0403 | MSHA biogenesis protein MshM | -2.0 | 1.66E-08 | -1.0 | 0.95 | 2.0 |
| VC0413 | MshP | -2.3 | 1.17E-12 | 1.1 | 0.40 | -2.0 |
| VC0463 | twitching motility protein PilT | -2.5 | 3.02E-10 | -1.0 | 0.73 | 2.4 |
| VC0533 | lipoprotein NlpD | -2.0 | 2.74E-03 | -1.1 | 0.60 | 1.9 |
| VC0534 | RNA polymerase sigma-38 factor | -2.3 | 3.55E-04 | 1.0 | 0.95 | -2.3 |
| VC0552 | quinone oxidoreductase | -2.0 | 3.71E-12 | -1.0 | 0.87 | 2.0 |
| VC0613 | beta-N-acetylhexosam inidase | -2.1 | 3.84E-07 | 1.1 | 0.58 | -1.9 |
| VC0614 | conserved hypothetical protein | -2.6 | 2.68E-06 | 1.2 | 0.31 | -2.2 |
| VC0615 | endoglucanase-related protein | -2.3 | 6.33E-09 | 1.1 | 0.45 | -2.1 |
| VC0616 | peptide ABC transporter, ATP-binding protein | -2.0 | 1.01E-03 | 1.3 | 0.10 | -1.5 |
| VC0617 | peptide ABC transporter, ATP-binding protein | -2.0 | 1.30E-04 | 1.1 | 0.29 | -1.8 |
| VC0618 | peptide ABC transporter, permease protein | -2.9 | 1.38E-06 | -1.0 | 0.86 | 2.8 |
| VC0620 | peptide ABC transporter, periplasmicpeptide-binding protein | -3.1 | 1.32E-17 | -1.4 | 0.00 | 2.2 |
| VC0633 | outer membrane protein OmpU | -2.4 | 1.11E-06 | -23.1 | 0.00 | 0.1 |
| VC0658 | c-di-GMP phosphodiesterase A-related protein | -2.7 | 3.73E-14 | 1.2 | 0.03 | -2.2 |
| VC0665 | sigma-54 dependent transcriptional regulator | -2.0 | 9.40E-08 | 1.0 | 0.87 | -2.0 |
| VC0697 | hypothetical protein | -2.7 | 2.25E-18 | 1.3 | 0.04 | -2.1 |
| VC0728 | conserved hypothetical protein | -3.0 | 5.86E-18 | 1.1 | 0.41 | -2.8 |
| VC0734 | malate synthase A | -5.7 | 5.92E-34 | 1.2 | 0.11 | -4.7 |
| VC0736 | isocitrate lyase | -2.9 | 3.97E-14 | 1.1 | 0.35 | -2.7 |
| VC0737 | acetoin utilization protein AcuB, putative | -2.5 | 1.84E-12 | 1.3 | 0.01 | -2.0 |
| VC0828 | toxin co-regulated pilin, TcpA | -24.9 | 1.09E-37 | -61.3 | 0.00 | 0.4 |
| VC0829 | toxin co-regulated pilus biosynthesis protein B | -2.0 | 4.00E-04 | -3.4 | 0.00 | 0.6 |
| VC0831 | toxin co-regulated pilus biosynthesis outermembrane protein C | -2.5 | 1.11E-08 | -5.0 | 0.00 | 0.5 |
| VC0833 | toxin co-regulated pilus biosynthesis protein D | -2.5 | 4.26E-04 | -7.5 | 0.00 | 0.3 |
| VC0834 | toxin co-regulated pilus biosynthesis protein S | -2.3 | 4.78E-05 | -8.9 | 0.00 | 0.3 |
| VC0835 | toxin co-regulated pilus biosynthesis protein T | -2.3 | 4.87E-05 | -5.6 | 0.00 | 0.4 |
| VC0836 | toxin co-regulated pilus biosynthesis protein E | -2.1 | 1.12E-03 | -3.7 | 0.00 | 0.6 |
| VC0837 | toxin co-regulated pilus biosynthesis protein F | -2.1 | 4.22E-05 | -5.3 | 0.00 | 0.4 |
| VC0838 | TCP pilus virulence regulatory protein | 1.2 | 0.49 | -2.3 | 0.00 | -0.5 |
| VC0844 | accessory colonization factor AcfA | -3.1 | 5.23E-08 | -9.8 | 0.00 | 0.3 |
| VC0858 | type IV pilin, putative | -2.2 | 1.66E-03 | -1.2 | 0.31 | 1.7 |
| VC0893 | chemotaxis protein PomB | -2.1 | 1.23E-11 | 1.1 | 0.51 | -2.0 |
| VC0935 | hypothetical protein | -2.2 | 3.89E-06 | 1.1 | 0.54 | -2.0 |
| VC0936 | polysaccharide export-related protein | -3.8 | 8.39E-16 | 1.4 | 0.01 | -2.7 |
| VC0937 | exopolysaccharide biosynthesis protein,putative | -2.4 | 5.30E-10 | 1.1 | 0.15 | -2.1 |
| VC0938 | hypothetical protein | -3.5 | 1.74E-11 | 1.3 | 0.07 | -2.6 |
| VC0957 | conserved hypothetical protein | -2.3 | 3.45E-04 | 1.1 | 0.46 | -2.1 |
| VC1008 | sodium-type flagellar protein MotY | -2.2 | 3.76E-09 | -1.0 | 0.88 | 2.2 |
| VC1009 | conserved hypothetical protein | -2.6 | 1.43E-11 | -1.2 | 0.20 | 2.2 |
| VC1029 | GGDEF family protein | -2.1 | 2.59E-06 | 1.2 | 0.02 | -1.7 |
| VC1031 | inosine monophosphate dehydrogenase-relatedprotein | -6.4 | 1.89E-59 | 1.3 | 0.00 | -4.9 |
| VC1032 | zinc/cadmium/mercury/lead-transporting ATPase | -3.3 | 2.45E-04 | -1.3 | 0.39 | 2.6 |
| VC1043 | long-chain fatty acid transport protein | -2.4 | 1.94E-04 | 1.3 | 0.11 | -1.9 |
| VC1050 | response regulator | -3.6 | 3.12E-24 | 1.3 | 0.00 | -2.7 |
| VC1064 | lipoprotein-related protein | -2.1 | 9.83E-09 | 1.1 | 0.37 | -1.9 |
| VC1080 | hypothetical protein | -2.3 | 2.80E-11 | 1.2 | 0.27 | -2.0 |
| VC1082 | response regulator | -2.6 | 1.21E-19 | 1.2 | 0.20 | -2.2 |
| VC1083 | hypothetical protein | -2.9 | 1.10E-13 | 1.2 | 0.21 | -2.5 |
| VC1084 | sensory box sensor histidine kinase | -4.0 | 9.19E-23 | 1.1 | 0.52 | -3.7 |
| VC1085 | sensor histidine kinase | -4.1 | 3.69E-31 | 1.0 | 0.69 | -3.9 |
| VC1086 | response regulator | -3.0 | 5.37E-21 | 1.1 | 0.27 | -2.7 |
| VC1087 | response regulator | -2.8 | 2.96E-14 | 1.2 | 0.17 | -2.4 |
| VC1088 | sensor histidine kinase | -3.5 | 1.69E-19 | 1.1 | 0.49 | -3.2 |
| VC1089 | periplasmic binding protein-related protein | -2.6 | 3.07E-12 | 1.5 | 0.00 | -1.7 |
| VC1101 | conserved hypothetical protein | -2.5 | 7.55E-12 | 1.0 | 0.94 | -2.4 |
| VC1103 | ABC transporter, ATP-binding protein | -2.1 | 7.33E-10 | -1.1 | 0.65 | 1.9 |

| | | | | | | |
|---|---|---|---|---|---|---|
| VC1112 | biotin synthase | -2.3 | 7.57E-05 | -1.3 | 0.10 | 1.7 |
| VC1116 | hypothetical protein | -4.9 | 1.15E-13 | 1.7 | 0.00 | -2.9 |
| VC1124 | conserved hypothetical protein | -2.7 | 1.09E-23 | 1.5 | 0.00 | -1.8 |
| VC1125 | hypothetical protein | -2.7 | 5.39E-13 | 1.4 | 0.01 | -1.9 |
| VC1142 | cold shock-like protein CspD | -2.5 | 3.53E-09 | 1.3 | 0.05 | -1.9 |
| VC1153 | conserved hypothetical protein | -2.2 | 3.39E-12 | 1.0 | 0.73 | -2.1 |
| VC1187 | hypothetical protein | -3.2 | 2.42E-11 | -2.1 | 0.00 | 1.5 |
| VC1188 | malate oxidoreductase | -3.8 | 6.65E-09 | -1.8 | 0.00 | 2.1 |
| VC1207 | hypothetical protein | -2.6 | 2.97E-12 | 1.5 | 0.00 | -1.7 |
| VC1222 | integration host factor, alpha subunit | -2.5 | 1.37E-11 | 1.3 | 0.04 | -1.9 |
| VC1223 | hypothetical protein | -3.3 | 5.33E-24 | 1.1 | 0.58 | -3.1 |
| VC1224 | hypothetical protein | -2.3 | 1.80E-08 | -1.2 | 0.15 | 1.9 |
| VC1236 | PilB-related protein | -2.3 | 5.07E-10 | 1.1 | 0.49 | -2.1 |
| VC1247 | hypothetical protein | -2.2 | 2.81E-07 | 1.2 | 0.10 | -1.7 |
| VC1248 | methyl-accepting chemotaxis protein | -7.0 | 1.62E-14 | 1.5 | 0.01 | -4.7 |
| VC1249 | conserved hypothetical protein | -2.2 | 1.27E-03 | -1.1 | 0.45 | 2.0 |
| VC1262 | hypothetical protein | -2.9 | 9.67E-10 | -1.2 | 0.21 | 2.5 |
| VC1269 | conserved hypothetical protein | -2.4 | 1.79E-23 | 1.3 | 0.05 | -1.9 |
| VC1295 | conserved hypothetical protein | -2.2 | 6.43E-09 | 1.4 | 0.00 | -1.6 |
| VC1298 | methyl-accepting chemotaxis protein | -3.0 | 1.75E-17 | 1.2 | 0.07 | -2.6 |
| VC1312 | alanine racemase, putative | -2.7 | 1.64E-10 | 1.5 | 0.01 | -1.8 |
| VC1313 | methyl-accepting chemotaxis protein | -2.8 | 1.67E-13 | 1.7 | 0.00 | -1.7 |
| VC1316 | chemotaxis protein CheY, putative | -2.6 | 4.94E-14 | 1.0 | 0.97 | -2.6 |
| VC1322 | conserved hypothetical protein | -3.9 | 3.19E-21 | 1.3 | 0.10 | -3.0 |
| VC1323 | hypothetical protein | -2.8 | 1.66E-16 | 1.5 | 0.00 | -1.9 |
| VC1325 | galactoside ABC transporter, periplasmicD-galactose/D-gluco se-binding protein | -4.6 | 1.32E-20 | 1.1 | 0.61 | -4.2 |
| VC1327 | galactoside ABC transporter, ATP-bindingprotein | -3.5 | 2.54E-17 | 1.1 | 0.54 | -3.2 |
| VC1328 | galactoside ABC transporter, permease protein | -4.0 | 5.67E-25 | 1.0 | 0.92 | -4.0 |
| VC1334 | conserved hypothetical protein | -3.1 | 4.33E-17 | 1.1 | 0.23 | -2.7 |
| VC1339 | conserved hypothetical protein | -2.1 | 1.46E-06 | -1.1 | 0.62 | 1.9 |
| VC1340 | prpE protein | -2.1 | 2.31E-08 | -1.0 | 0.98 | 2.1 |
| VC1348 | response regulator | -3.3 | 7.56E-21 | 1.5 | 0.00 | -2.2 |
| VC1349 | sensory box sensor histidine kinase/responseregulator | -3.1 | 6.93E-16 | 1.2 | 0.14 | -2.6 |
| VC1355 | acylphosphatase | -2.2 | 3.67E-08 | 1.4 | 0.02 | -1.6 |
| VC1359 | amino acid ABC transporter, ATP-binding protein | -2.5 | 1.24E-12 | 1.3 | 0.04 | -1.9 |
| VC1360 | amino acid ABC transporter, permease protein | -2.9 | 2.28E-14 | 1.2 | 0.21 | -2.4 |
| VC1361 | amino acid ABC transporter, permease protein | -3.0 | 4.30E-16 | 1.2 | 0.10 | -2.4 |
| VC1362 | amino acid ABC transporter, periplasmic aminoacid-binding protein | -5.6 | 2.98E-12 | 1.3 | 0.15 | -4.5 |
| VC1368 | hypothetical protein | -2.2 | 1.28E-08 | 1.4 | 0.01 | -1.6 |
| VC1369 | conserved hypothetical protein | -3.7 | 1.49E-22 | 1.5 | 0.00 | -2.4 |
| VC1370 | GGDEF family protein | -4.9 | 9.95E-25 | 1.3 | 0.09 | -3.9 |
| VC1376 | GGDEF family protein | -2.2 | 1.43E-09 | 1.2 | 0.03 | -1.8 |
| VC1394 | methyl-accepting chemotaxis protein | -4.1 | 1.36E-29 | 1.3 | 0.03 | -3.2 |
| VC1395 | response regulator cheY1 | -4.6 | 3.55E-40 | 1.2 | 0.05 | -3.8 |
| VC1396 | hypothetical protein | -5.2 | 3.64E-43 | 1.2 | 0.03 | -4.2 |
| VC1397 | chemotaxis protein CheA | -4.9 | 1.22E-34 | 1.2 | 0.26 | -4.3 |
| VC1398 | chemotaxis protein CheY | -5.8 | 1.64E-27 | 1.1 | 0.26 | -5.1 |
| VC1399 | chemotaxis protein methyltransferase CheR | -5.6 | 8.47E-35 | 1.2 | 0.16 | -4.5 |
| VC1400 | hypothetical protein | -4.2 | 4.51E-24 | 1.3 | 0.05 | -3.2 |
| VC1401 | protein-glutamate methylesterase CheB | -3.3 | 5.82E-17 | 1.2 | 0.22 | -2.8 |
| VC1402 | purine-binding chemotaxis protein Chew,putative | -2.9 | 3.66E-12 | 1.2 | 0.27 | -2.5 |
| VC1403 | methyl-accepting chemotaxis protein | -3.5 | 3.64E-19 | 1.4 | 0.00 | -2.4 |
| VC1405 | methyl-accepting chemotaxis protein | -2.6 | 2.69E-13 | 1.4 | 0.01 | -1.9 |
| VC1406 | methyl-accepting chemotaxis protein | -2.7 | 3.06E-18 | 1.3 | 0.00 | -2.1 |
| VC1456 | cholera enterotoxin, B subunit | -18.4 | 3.71E-46 | -21.9 | 0.00 | 0.8 |
| VC1457 | cholera enterotoxin, A subunit | -20.2 | 1.83E-59 | -18.6 | 0.00 | 1.1 |
| VC1484 | ribosome modulation factor | -4.9 | 1.00E-27 | 1.6 | 0.00 | -3.0 |
| VC1506 | hypothetical protein | -3.2 | 1.35E-14 | -1.0 | 0.92 | 3.2 |
| VC1509 | nicotinatemononucleo tide:5,6-dimethylbenzimidazole phosphoribosyltransferase | -3.1 | 6.41E-14 | 1.5 | 0.00 | -2.0 |
| VC1528 | hypothetical protein | -2.3 | 4.16E-11 | 1.5 | 0.01 | -1.6 |
| VC1538 | hypothetical protein | -2.3 | 7.41E-10 | 1.3 | 0.05 | -1.8 |
| VC1560 | catalase/peroxidase | -4.3 | 6.51E-09 | 1.2 | 0.23 | -3.6 |
| VC1594 | aldose 1-epimerase | -2.6 | 1.05E-13 | -1.1 | 0.13 | 2.3 |
| VC1595 | galactokinase | -2.7 | 1.91E-13 | -1.1 | 0.30 | 2.3 |
| VC1601 | hypothetical protein | -2.1 | 2.85E-07 | 1.3 | 0.04 | -1.6 |
| VC1602 | chemotaxis protein CheV | -2.7 | 3.95E-12 | 1.3 | 0.05 | -2.1 |
| VC1603 | hypothetical protein | -2.7 | 2.61E-16 | 1.6 | 0.00 | -1.6 |
| VC1612 | fimbrial biogenesis and twitching motilityprotein, putative | -2.1 | 3.39E-08 | 1.2 | 0.23 | -1.8 |
| VC1618 | multidrug resistance protein, putative | -3.3 | 1.21E-32 | 1.1 | 0.49 | -3.1 |
| VC1641 | conserved hypothetical protein | -2.2 | 6.55E-09 | 1.3 | 0.02 | -1.6 |
| VC1643 | methyl-accepting chemotaxis protein | -3.6 | 1.77E-20 | 1.1 | 0.37 | -3.3 |
| VC1662 | conserved hypothetical protein | -2.8 | 8.03E-13 | 1.2 | 0.02 | -2.2 |
| VC1672 | DNA-3-methyladenine glycosidase I | -2.2 | 3.99E-07 | 1.5 | 0.00 | -1.4 |
| VC1673 | transporter, AcrB/D/F family | -2.5 | 3.84E-12 | 1.0 | 0.70 | -2.4 |
| VC1674 | periplasmic linker protein, putative | -3.5 | 1.62E-17 | 1.0 | 0.61 | -3.3 |
| VC1675 | multidrug resistance protein, putative | -2.7 | 5.16E-19 | 1.2 | 0.29 | -2.3 |
| VC1677 | phage shock protein B | -2.0 | 1.09E-04 | -1.5 | 0.01 | 1.4 |
| VC1699 | hypothetical protein | -2.3 | 1.59E-19 | 1.2 | 0.19 | -1.9 |
| VC1706 | transcriptional activator MetR | -2.1 | 7.45E-07 | 1.4 | 0.01 | -1.5 |
| VC1707 | hypothetical protein | -3.8 | 1.08E-24 | 1.4 | 0.00 | -2.6 |
| VC1736 | arginyl-tRNA-protein transferase-relatedprotein | -3.1 | 1.37E-20 | 1.4 | 0.00 | -2.2 |
| VC1741 | transcriptional regulator, TetR family | -2.2 | 7.63E-08 | 1.5 | 0.00 | -1.5 |
| VC1778 | conserved hypothetical protein | -2.2 | 3.15E-12 | 1.4 | 0.00 | -1.5 |
| VC1819 | aldehyde dehydrogenase | -2.1 | 2.71E-06 | -2.2 | 0.00 | 1.0 |

| | | | | | | |
|---|---|---|---|---|---|---|
| VC1831 | sensor histidine kinase | -3.1 | 4.82E-27 | 1.1 | 0.47 | -2.9 |
| VC1842 | conserved hypothetical protein | -2.1 | 5.72E-04 | 1.1 | 0.59 | -1.9 |
| VC1851 | conserved hypothetical protein | -2.8 | 8.48E-16 | 1.2 | 0.10 | -2.2 |
| VC1868 | methyl-accepting chemotaxis protein | -4.5 | 5.84E-28 | 1.5 | 0.00 | -3.0 |
| VC1872 | conserved hypothetical protein | -5.1 | 5.83E-25 | 1.1 | 0.62 | -4.7 |
| VC1873 | conserved hypothetical protein | -6.1 | 1.14E-42 | 1.0 | 0.96 | -6.1 |
| VC1874 | conserved hypothetical protein | -6.9 | 1.33E-15 | 1.2 | 0.30 | -5.9 |
| VC1914 | integration host factor, beta subunit | -2.7 | 6.48E-19 | 1.3 | 0.01 | -2.2 |
| VC1932 | hypothetical protein | -2.6 | 7.44E-07 | 1.5 | 0.01 | -1.7 |
| VC1933 | hypothetical protein | -3.7 | 4.51E-25 | 1.5 | 0.00 | -2.4 |
| VC1934 | GGDEF family protein | -2.9 | 5.86E-12 | 1.2 | 0.27 | -2.5 |
| VC1950 | biotin sulfoxide reductase | -2.2 | 5.12E-08 | -1.3 | 0.00 | 1.7 |
| VC1952 | chitinase | -2.0 | 5.17E-08 | -1.0 | 0.64 | 1.9 |
| VC1967 | methyl-accepting chemotaxis protein | -2.5 | 1.08E-12 | 1.4 | 0.00 | -1.7 |
| VC1993 | 2,4-dienoyl-CoA reductase | -2.1 | 1.48E-08 | -1.0 | 0.87 | 2.1 |
| VC2008 | pyruvate kinase II | -2.2 | 2.65E-10 | 1.1 | 0.29 | -2.0 |
| VC2047 | oxidoreductase, short-chaindehydrogenase/reduc tase family | -2.5 | 1.94E-17 | 1.3 | 0.01 | -2.0 |
| VC2059 | purine-binding chemotaxis protein CheW | -2.4 | 1.41E-04 | 1.2 | 0.18 | -2.0 |
| VC2060 | conserved hypothetical protein | -2.3 | 2.15E-09 | 1.0 | 0.97 | -2.2 |
| VC2063 | chemotaxis protein CheA | -2.1 | 3.32E-07 | 1.0 | 0.82 | -2.0 |
| VC2103 | transcriptional regulator, LysR family | -2.1 | 5.22E-14 | 1.2 | 0.04 | -1.7 |
| VC2120 | flagellar biosynthetic protein FlhB | -2.1 | 1.31E-08 | -1.1 | 0.52 | 2.0 |
| VC2121 | flagellar biosynthetic protein FliR | -2.0 | 6.15E-07 | -1.2 | 0.07 | 1.7 |
| VC2122 | flagellar biosynthetic protein FliQ | -2.9 | 5.20E-16 | -1.1 | 0.54 | 2.8 |
| VC2123 | flagellar biosynthetic protein FliP | -2.2 | 2.39E-10 | -1.2 | 0.19 | 1.9 |
| VC2124 | flagellar protein FliO | -2.2 | 6.57E-13 | -1.1 | 0.36 | 2.0 |
| VC2125 | flagellar motor switch protein FliN | -2.0 | 1.03E-09 | -1.1 | 0.66 | 1.9 |
| VC2126 | flagellar motor switch protein FliM | -2.2 | 4.77E-09 | -1.1 | 0.27 | 2.0 |
| VC2127 | flagellar protein FliL, putative | -2.4 | 1.46E-10 | -1.0 | 0.70 | 2.3 |
| VC2128 | flagellar hook-length control protein FliK,putative | -2.7 | 2.75E-14 | -1.2 | 0.09 | 2.3 |
| VC2129 | flagellar protein FliJ, putative | -2.0 | 1.81E-12 | 1.1 | 0.54 | -1.9 |
| VC2132 | flagellar motor switch protein FliG | -2.0 | 2.00E-09 | -1.0 | 0.76 | 1.9 |
| VC2138 | flagellar protein FliS | -2.1 | 1.64E-09 | 1.0 | 0.82 | -2.1 |
| VC2139 | flagellar rod protein FlaI, putative | -2.5 | 2.94E-15 | 1.0 | 0.71 | -2.4 |
| VC2140 | flagellar hook-associated protein FliD | -2.9 | 8.95E-06 | -1.0 | 0.79 | 2.8 |
| VC2141 | flagellin FlaG | -3.3 | 1.09E-06 | 1.1 | 0.65 | -3.1 |
| VC2142 | flagellin FlaB | -3.5 | 8.48E-16 | 1.1 | 0.59 | -3.2 |
| VC2143 | flagellin FlaD | -2.4 | 8.21E-08 | 1.2 | 0.32 | -2.0 |
| VC2144 | flagellin FlaE | -2.7 | 9.51E-15 | 1.3 | 0.01 | -2.1 |
| VC2161 | methyl-accepting chemotaxis protein | -3.5 | 2.37E-07 | 1.2 | 0.25 | -3.0 |
| VC2187 | flagellin FlaC | -3.4 | 3.17E-14 | 1.1 | 0.44 | -3.0 |
| VC2190 | flagellar hook-associated protein FlgL | -2.5 | 4.88E-10 | -1.4 | 0.01 | 1.8 |
| VC2192 | flagellar protein FlgJ | -2.4 | 4.22E-11 | -1.3 | 0.07 | 1.9 |
| VC2194 | flagellar L-ring protein FlgH | -2.6 | 1.28E-14 | -1.2 | 0.16 | 2.1 |
| VC2195 | flagellar basal-body rod protein FlgG | -2.3 | 1.99E-14 | -1.1 | 0.34 | 2.0 |
| VC2198 | basal-body rod modification protein FlgD | -2.1 | 1.98E-08 | -1.2 | 0.12 | 1.7 |
| VC2201 | chemotaxis protein methyltransferase CheR | -2.7 | 2.26E-05 | -1.0 | 0.76 | 2.6 |
| VC2202 | chemotaxis protein CheV | -2.3 | 5.91E-04 | 1.2 | 0.21 | -1.9 |
| VC2204 | negative regulator of flagellin synthesis FlgM,putative | -2.2 | 4.33E-10 | 1.2 | 0.07 | -1.9 |
| VC2205 | hypothetical protein | -2.4 | 4.06E-11 | -1.1 | 0.46 | 2.2 |
| VC2206 | conserved hypothetical protein | -2.4 | 3.83E-10 | -1.1 | 0.44 | 2.2 |
| VC2207 | hypothetical protein | -2.2 | 2.19E-10 | 1.0 | 0.99 | -2.2 |
| VC2221 | hypothetical protein | -3.5 | 2.35E-14 | 1.2 | 0.31 | -2.9 |
| VC2231 | oxidoreductase, acyl-CoA dehydrogenase family | -2.4 | 2.10E-11 | 1.2 | 0.09 | -2.1 |
| VC2240 | decarboxylase | -2.1 | 2.59E-10 | -1.2 | 0.03 | 1.7 |
| VC2241 | cytochrome c554 | -2.7 | 3.75E-16 | 1.4 | 0.02 | -2.0 |
| VC2264 | conserved hypothetical protein | -2.8 | 9.69E-16 | 1.1 | 0.51 | -2.6 |
| VC2314 | hypothetical protein | -3.5 | 2.91E-13 | 1.3 | 0.10 | -2.8 |
| VC2316 | N-acetylglutamate synthase | -2.7 | 3.54E-12 | 1.4 | 0.00 | -2.0 |
| VC2338 | ß-galactosidase, LacZ | -2.8 | 2.20E-05 | -1.1 | 0.35 | 2.4 |
| VC2340 | conserved hypothetical protein | -3.5 | 7.33E-22 | 1.4 | 0.01 | -2.5 |
| VC2357 | hypothetical protein | -2.5 | 9.02E-10 | -1.2 | 0.16 | 2.1 |
| VC2358 | hypothetical protein | -3.0 | 8.14E-16 | 1.3 | 0.03 | -2.4 |
| VC2370 | sensory box/GGDEF family protein | -2.1 | 1.34E-08 | 1.2 | 0.21 | -1.8 |
| VC2383 | transcriptional regulator, LysR family | -2.0 | 9.54E-06 | 1.5 | 0.00 | -1.3 |
| VC2384 | conserved hypothetical protein | -3.7 | 7.79E-18 | 1.8 | 0.00 | -2.1 |
| VC2454 | GGDEF family protein | -2.7 | 7.59E-14 | 1.1 | 0.53 | -2.5 |
| VC2455 | hypothetical protein | -2.9 | 3.28E-13 | 1.1 | 0.32 | -2.6 |
| VC2456 | hypothetical protein | -2.8 | 1.09E-18 | 1.2 | 0.12 | -2.3 |
| VC2464 | sigma-E factor regulatory protein RseC | -2.3 | 9.37E-10 | -1.5 | 0.00 | 1.5 |
| VC2465 | sigma-E factor regulatory protein RseB | -2.3 | 1.01E-11 | -1.4 | 0.01 | 1.7 |
| VC2466 | sigma-E factor negative regulatory protein RseA | -2.2 | 4.21E-09 | -1.3 | 0.02 | 1.7 |
| VC2467 | RNA polymerase sigma-E factor | -2.3 | 3.21E-10 | -1.3 | 0.00 | 1.7 |
| VC2507 | conserved hypothetical protein | -3.9 | 6.58E-22 | 1.2 | 0.02 | -3.2 |
| VC2508 | ornithine carbamoyltransferase | -2.2 | 1.25E-10 | 1.2 | 0.15 | -1.9 |
| VC2529 | RNA polymerase sigma-54 factor | -2.1 | 8.76E-08 | 1.0 | 0.76 | -2.0 |
| VC2530 | sigma-54 modulation protein, putative | -2.7 | 4.54E-14 | 1.0 | 0.67 | -2.6 |
| VC2565 | elaA protein | -2.5 | 1.88E-25 | 2.1 | 0.00 | -1.2 |
| VC2601 | sodium-type flagellar protein MotX | -2.7 | 3.78E-17 | 1.2 | 0.06 | -2.2 |
| VC2622 | hypothetical protein | -2.8 | 8.22E-17 | 1.4 | 0.00 | -1.9 |
| VC2697 | GGDEF family protein | -2.7 | 4.61E-11 | 1.4 | 0.00 | -1.9 |
| VC2702 | transcriptional regulator, LuxR family | -2.1 | 4.58E-09 | -1.2 | 0.08 | 1.7 |
| VC2704 | hypothetical protein | -23.5 | 9.84E-52 | 1.3 | 0.00 | -17.7 |
| VC2705 | sodium/solute symporter, putative | -29.0 | 1.28E-135 | 1.2 | 0.08 | -23.2 |

| | | | | | | |
|---|---|---|---|---|---|---|
| VC2717 | hypothetical protein | -3.4 | 5.59E-20 | 1.1 | 0.29 | -3.1 |
| VC2726 | general secretion pathway protein K | -2.0 | 1.24E-07 | -1.4 | 0.00 | 1.5 |
| VC2727 | general secretion pathway protein J | -2.2 | 2.97E-08 | -1.1 | 0.46 | 2.0 |
| VC2750 | GGDEF family protein | -2.7 | 1.64E-14 | 1.2 | 0.16 | -2.2 |
| VCA0008 | methyl-accepting chemotaxis protein | -8.4 | 1.03E-56 | 1.4 | 0.01 | -5.9 |
| VCA0009 | hypothetical protein | -7.6 | 1.46E-20 | 1.3 | 0.12 | -6.1 |
| VCA0013 | maltodextrin phosphorylase | -3.2 | 2.46E-14 | -1.3 | 0.04 | 2.4 |
| VCA0014 | 4-alpha-glucanotrans ferase | -3.0 | 1.42E-23 | -1.3 | 0.00 | 2.2 |
| VCA0031 | methyl-accepting chemotaxis protein | -2.3 | 2.82E-09 | 1.2 | 0.12 | -1.9 |
| VCA0032 | hypothetical protein | -2.6 | 2.01E-12 | 1.4 | 0.00 | -1.8 |
| VCA0034 | conserved hypothetical protein | -2.8 | 2.03E-14 | 1.1 | 0.35 | -2.5 |
| VCA0047 | conserved hypothetical protein | -2.0 | 2.39E-11 | -1.0 | 0.97 | 2.0 |
| VCA0049 | GGDEF family protein | -2.1 | 3.14E-09 | -1.1 | 0.35 | 1.9 |
| VCA0074 | GGDEF family protein | -2.9 | 4.08E-17 | 1.3 | 0.00 | -2.2 |
| VCA0075 | hypothetical protein | -2.6 | 2.83E-06 | 1.2 | 0.18 | -2.1 |
| VCA0078 | hypothetical protein | -2.4 | 1.39E-14 | 1.1 | 0.45 | -2.2 |
| VCA0080 | GGDEF family protein | -2.7 | 2.72E-15 | 1.6 | 0.00 | -1.7 |
| VCA0129 | ribose ABC transporter, permease protein | -2.1 | 2.98E-04 | 1.1 | 0.48 | -1.8 |
| VCA0130 | ribose ABC transporter, periplasmicD-ribose-binding protein | -2.7 | 1.22E-12 | 1.2 | 0.03 | -2.2 |
| VCA0131 | ribokinase | -2.2 | 9.74E-05 | -1.0 | 0.97 | 2.2 |
| VCA0144 | immunogenic protein | -2.1 | 1.81E-06 | 1.1 | 0.21 | -1.8 |
| VCA0146 | conserved hypothetical protein | -2.6 | 2.40E-06 | -1.1 | 0.38 | 2.3 |
| VCA0147 | transcriptional regulator, putative | -2.5 | 5.74E-07 | -1.3 | 0.07 | 1.9 |
| VCA0152 | conserved hypothetical protein | -5.6 | 1.09E-25 | 1.1 | 0.61 | -5.3 |
| VCA0153 | conserved hypothetical protein | -5.5 | 2.54E-23 | 1.2 | 0.28 | -4.8 |
| VCA0154 | conserved hypothetical protein | -5.5 | 2.46E-31 | 1.1 | 0.66 | -5.2 |
| VCA0155 | NADH dehydrogenase, putative | -3.8 | 5.45E-24 | 1.2 | 0.04 | -3.1 |
| VCA0156 | conserved hypothetical protein | -4.0 | 5.45E-22 | 1.2 | 0.13 | -3.5 |
| VCA0157 | NADH dehydrogenase, putative | -3.6 | 3.49E-24 | 1.2 | 0.12 | -2.9 |
| VCA0161 | tryptophanase | -2.7 | 1.48E-11 | 1.3 | 0.00 | -2.0 |
| VCA0176 | methyl-accepting chemotaxis protein | -2.3 | 4.75E-10 | 1.3 | 0.02 | -1.7 |
| VCA0186 | hypothetical protein | -2.1 | 1.13E-06 | 1.0 | 0.98 | -2.1 |
| VCA0188 | hypothetical protein | -3.1 | 1.73E-12 | 1.7 | 0.00 | -1.8 |
| VCA0189 | response regulator | -2.1 | 1.13E-16 | 1.3 | 0.03 | -1.6 |
| VCA0191 | conserved hypothetical protein | -2.1 | 1.41E-09 | -1.0 | 0.86 | 2.1 |
| VCA0210 | response regulator, putative | -4.0 | 1.63E-29 | 1.4 | 0.00 | -2.8 |
| VCA0211 | sensory box sensor histidine kinase | -4.0 | 6.20E-48 | 1.2 | 0.13 | -3.3 |
| VCA0212 | hypothetical protein | -3.2 | 8.82E-19 | 1.4 | 0.03 | -2.4 |
| VCA0220 | hemolysin secretion protein HylB | -2.3 | 1.44E-20 | 1.2 | 0.19 | -2.0 |
| VCA0221 | lactonizing lipase | -3.8 | 1.72E-11 | 1.3 | 0.10 | -2.9 |
| VCA0222 | lipase activator protein, putative | -2.2 | 4.05E-04 | 1.2 | 0.40 | -1.9 |
| VCA0268 | methyl-accepting chemotaxis protein | -3.5 | 1.84E-18 | 2.0 | 0.00 | -1.8 |
| VCA0269 | decarboxylase, group II | -2.9 | 3.41E-12 | 1.9 | 0.00 | -1.5 |
| VCA0282 | IS5 transposase | -2.0 | 1.05E-05 | -1.6 | 0.00 | 1.2 |
| VCA0534 | pseudogene; contains authentic frameshift mutation | -2.4 | 7.95E-10 | -1.4 | 0.03 | 1.7 |
| VCA0551 | hypothetical protein | -2.5 | 4.01E-17 | 1.3 | 0.03 | -1.9 |
| VCA0557 | GGDEF family protein | -2.2 | 2.81E-09 | 1.0 | 0.81 | -2.1 |
| VCA0558 | gamma-glutamyltransp eptidase, putative | -2.5 | 1.68E-12 | 1.3 | 0.13 | -2.0 |
| VCA0574 | conserved hypothetical protein | -2.3 | 2.42E-09 | -1.0 | 0.82 | 2.3 |
| VCA0583 | hypothetical protein | -4.2 | 2.32E-27 | 1.5 | 0.00 | -2.8 |
| VCA0588 | peptide ABC transporter, ATP-binding protein,putative | -2.3 | 1.80E-09 | -1.1 | 0.32 | 2.1 |
| VCA0593 | hypothetical protein | -3.1 | 8.74E-19 | 1.5 | 0.00 | -2.1 |
| VCA0619 | hypothetical protein | -2.5 | 2.40E-15 | 1.0 | 0.70 | -2.4 |
| VCA0638 | transporter, AcrB/D/F family | -2.0 | 1.35E-11 | -1.0 | 0.83 | 2.0 |
| VCA0639 | AcrA/AcrE family protein | -2.2 | 4.50E-17 | 1.3 | 0.06 | -1.7 |
| VCA0645 | hypothetical protein | -2.1 | 6.69E-05 | 1.5 | 0.02 | -1.4 |
| VCA0646 | conserved hypothetical protein/hemolysin,putative | -3.9 | 9.23E-16 | 1.3 | 0.07 | -3.1 |
| VCA0647 | hypothetical protein | -4.1 | 7.58E-12 | 1.3 | 0.10 | -3.1 |
| VCA0648 | hypothetical protein | -2.5 | 1.99E-05 | 1.9 | 0.00 | -1.3 |
| VCA0649 | hypothetical protein | -2.5 | 1.19E-05 | 1.7 | 0.01 | -1.5 |
| VCA0650 | hypothetical protein | -4.4 | 9.33E-11 | 2.2 | 0.00 | -1.9 |
| VCA0659 | protein F-related protein | -3.0 | 7.33E-20 | 1.5 | 0.00 | -1.9 |
| VCA0666 | L-serine dehydratase 1 | -2.4 | 1.37E-12 | 1.1 | 0.34 | -2.1 |
| VCA0681 | conserved hypothetical protein | -3.8 | 1.04E-27 | 1.3 | 0.01 | -2.9 |
| VCA0683 | sensor protein UhpB | -2.3 | 4.26E-09 | -1.0 | 0.66 | 2.2 |
| VCA0684 | regulatory protein UhpC | -2.2 | 1.09E-12 | -1.2 | 0.05 | 1.9 |
| VCA0695 | hypothetical protein | -5.0 | 2.22E-24 | 1.3 | 0.02 | -3.7 |
| VCA0698 | hypothetical protein | -4.4 | 1.89E-28 | 1.3 | 0.02 | -3.4 |
| VCA0707 | regulatory protein UhpC, putative | -2.1 | 1.03E-03 | 2.4 | 0.00 | -0.8 |
| VCA0719 | sensor histide kinase | -3.5 | 2.97E-18 | 1.3 | 0.05 | -2.6 |
| VCA0720 | guanylate cyclase-related protein | -3.5 | 5.42E-25 | 1.4 | 0.00 | -2.5 |
| VCA0722 | hypothetical protein | -2.3 | 9.02E-08 | 1.2 | 0.15 | -1.8 |
| VCA0731 | hypothetical protein | -3.1 | 6.01E-08 | 1.5 | 0.01 | -2.1 |
| VCA0738 | hypothetical protein | -3.0 | 6.63E-15 | 1.9 | 0.00 | -1.6 |
| VCA0757 | arginine ABC transporter, permease protein | -2.4 | 1.16E-07 | 1.1 | 0.49 | -2.3 |
| VCA0758 | arginine ABC transporter, permease protein | -2.3 | 3.84E-07 | 1.2 | 0.18 | -1.9 |
| VCA0760 | arginine ABC transporter, ATP-binding protein | -2.7 | 2.20E-21 | 1.1 | 0.31 | -2.4 |
| VCA0788 | DnaJ-related protein | -2.3 | 6.99E-14 | 1.0 | 0.80 | -2.2 |
| VCA0798 | CbbY family protein | -2.1 | 1.91E-07 | 1.2 | 0.09 | -1.7 |
| VCA0803 | serine protease, putative | -4.5 | 7.01E-29 | 1.5 | 0.00 | -2.9 |
| VCA0813 | aminopeptidase | -2.2 | 5.72E-11 | -1.1 | 0.38 | 2.0 |
| VCA0831 | hypothetical protein | -3.7 | 1.05E-03 | 1.3 | 0.26 | -2.8 |
| VCA0848 | GGDEF family protein | -2.6 | 9.34E-26 | 1.6 | 0.00 | -1.6 |
| VCA0860 | alpha-amylase | -3.5 | 7.67E-16 | -1.2 | 0.24 | 3.0 |

| | | | | | | |
|---|---|---|---|---|---|---|
| VCA0864 | methyl-accepting chemotaxis protein | -2.1 | 1.21E-07 | 1.6 | 0.00 | -1.3 |
| VCA0865 | hemagglutinin/protease | -9.9 | 1.12E-44 | 1.6 | 0.00 | -6.2 |
| VCA0867 | outer membrane protein OmpW | -2.6 | 5.94E-07 | 1.1 | 0.69 | -2.5 |
| VCA0868 | hypothetical protein | -2.3 | 6.45E-09 | 1.1 | 0.33 | -2.1 |
| VCA0880 | motility-associated killing, *makD* | -3.9 | 6.10E-23 | 1.1 | 0.14 | -3.5 |
| VCA0881 | motility-associated killing, *makC* | -4.2 | 1.06E-26 | 1.1 | 0.33 | -3.7 |
| VCA0882 | motility-associated killing, *makB* | -4.2 | 3.21E-22 | 1.0 | 0.86 | -4.1 |
| VCA0883 | motility-associated killing, *makA* | -2.9 | 1.67E-13 | 1.1 | 0.58 | -2.7 |
| VCA0884 | hypothetical protein | -3.8 | 3.67E-21 | 1.0 | 0.79 | -3.6 |
| VCA0892 | hypothetical protein | -3.4 | 3.49E-23 | -1.0 | 0.95 | 3.4 |
| VCA0895 | chemotactic transducer-related protein | -3.0 | 2.49E-20 | 1.2 | 0.06 | -2.5 |
| VCA0900 | hypothetical protein | -2.6 | 9.54E-16 | 1.1 | 0.18 | -2.3 |
| VCA0901 | hypothetical protein | -2.6 | 1.81E-14 | 1.2 | 0.27 | -2.3 |
| VCA0906 | methyl-accepting chemotaxis protein | -5.3 | 5.66E-45 | 1.2 | 0.11 | -4.4 |
| VCA0920 | hypothetical protein | -2.4 | 2.18E-14 | 1.2 | 0.15 | -2.1 |
| VCA0923 | methyl-accepting chemotaxis protein | -4.0 | 2.82E-29 | 1.3 | 0.03 | -3.0 |
| VCA0931 | conserved hypothetical protein | -2.2 | 1.29E-11 | 1.2 | 0.15 | -1.8 |
| VCA0935 | hypothetical protein | -5.9 | 2.86E-12 | 1.9 | 0.00 | -3.2 |
| VCA0944 | maltose ABC transporter, permease protein | -3.4 | 2.28E-13 | -1.4 | 0.07 | 2.5 |
| VCA0945 | maltose ABC transporter, periplasmicmaltose-binding protein | -3.3 | 1.34E-13 | -1.2 | 0.19 | 2.7 |
| VCA0946 | maltose/maltodextrin ABC transporter,ATP-binding protein | -2.5 | 8.38E-08 | -1.1 | 0.61 | 2.3 |
| VCA0957 | malate synthase-related protein | -2.7 | 2.19E-16 | 1.1 | 0.34 | -2.4 |
| VCA0965 | GGDEF family protein | -4.1 | 7.72E-28 | 1.2 | 0.05 | -3.4 |
| VCA0978 | amino acid ABC transporter, periplasmic aminoacid-binding protein, putative | -4.1 | 1.78E-32 | 1.2 | 0.17 | -3.4 |
| VCA0979 | methyl-accepting chemotaxis protein | -2.7 | 9.39E-13 | 1.0 | 0.91 | -2.6 |
| VCA0981 | hypothetical protein | -3.6 | 1.09E-20 | 1.3 | 0.06 | -2.7 |
| VCA0982 | transcriptional regulator, LysR family | -2.1 | 1.23E-08 | -1.2 | 0.14 | 1.8 |
| VCA0988 | methyl-accepting chemotaxis protein | -3.2 | 3.93E-23 | 1.0 | 0.73 | -3.0 |
| VCA1015 | Na+/H+ antiporter | -2.9 | 8.35E-21 | 1.0 | 0.73 | -2.8 |
| VCA1016 | hypothetical protein | -3.9 | 2.23E-23 | 1.5 | 0.00 | -2.7 |
| VCA1017 | methylated-DNA--prot ein-cysteineS-methyltransferas e | -3.9 | 8.30E-22 | 1.4 | 0.00 | -2.9 |
| VCA1024 | hypothetical protein | -6.0 | 6.23E-41 | 1.6 | 0.00 | -3.8 |
| VCA1028 | maltoporin | -3.5 | 6.21E-14 | -1.2 | 0.25 | 2.9 |
| VCA1033 | extracellular solute-binding protein, putative | -4.4 | 8.12E-23 | 1.1 | 0.64 | -4.1 |
| VCA1034 | methyl-accepting chemotaxis protein | -5.5 | 2.28E-40 | 1.2 | 0.08 | -4.6 |
| VCA1042 | Ccm2-related protein | -2.1 | 1.84E-11 | 1.2 | 0.08 | -1.7 |
| VCA1054 | conserved hypothetical protein | -3.7 | 1.81E-29 | 1.3 | 0.00 | -2.9 |
| VCA1056 | methyl-accepting chemotaxis protein | -4.3 | 3.41E-27 | 1.2 | 0.14 | -3.5 |
| VCA1086 | response regulator | -3.4 | 6.64E-19 | 1.4 | 0.01 | -2.5 |
| VCA1087 | anti-sigma F factor antagonist, putative | -3.3 | 5.19E-17 | 1.3 | 0.02 | -2.5 |
| VCA1088 | methyl-accepting chemotaxis protein | -4.0 | 1.67E-33 | 1.3 | 0.03 | -3.1 |
| VCA1089 | cheB3 methylesterase | -5.4 | 1.96E-33 | 1.5 | 0.00 | -3.7 |
| VCA1090 | chemotaxis protein CheD, putative | -6.3 | 4.13E-46 | 1.4 | 0.01 | -4.6 |
| VCA1091 | chemotaxis protein methyltransferase CheR | -4.8 | 2.16E-29 | 1.3 | 0.03 | -3.7 |
| VCA1092 | methyl-accepting chemotaxis protein | -4.3 | 9.46E-25 | 1.2 | 0.09 | -3.5 |
| VCA1093 | purine-binding chemotaxis protein CheW | -4.0 | 6.14E-22 | 1.2 | 0.03 | -3.3 |
| VCA1094 | purine-binding chemotaxis protein CheW | -4.2 | 6.33E-24 | 1.1 | 0.50 | -3.8 |
| VCA1095 | chemotaxis protein CheA | -4.2 | 6.28E-23 | 1.3 | 0.08 | -3.3 |
| VCA1096 | chemotaxis protein CheY | -3.7 | 9.73E-32 | 1.3 | 0.02 | -2.9 |
| VCA1097 | conserved hypothetical protein | -3.6 | 1.32E-30 | 1.4 | 0.00 | -2.5 |
| VCA1108 | oxidoreductase, short-chaindehydrogenase/reduc tase family | -2.5 | 4.90E-05 | 1.4 | 0.03 | -1.7 |
| VCr025 | 5S ribosomal RNA | -36.4 | 8.81E-61 | -22.8 | 0.00 | 1.6 |
| 16Sa | 16S ribosomal RNA | -3.1 | 7.93E-12 | -1.3 | 0.19 | 2.5 |
| 16Sb | 16S ribosomal RNA | -2.7 | 2.27E-08 | -1.2 | 0.20 | 2.1 |
| 16Sc | 16S ribosomal RNA | -3.4 | 7.84E-12 | -1.4 | 0.07 | 2.5 |
| 16Se | 16S ribosomal RNA | -2.5 | 4.14E-08 | -1.3 | 0.12 | 2.0 |
| 16Sg | 16S ribosomal RNA | -2.2 | 9.44E-07 | -1.3 | 0.12 | 1.7 |
| 23Sa | 23S ribosomal RNA | -3.1 | 2.03E-09 | -1.3 | 0.10 | 2.3 |
| 23Sc | 23S ribosomal RNA | -5.6 | 3.27E-08 | -1.7 | 0.11 | 3.2 |
| 23Sd | 23S ribosomal RNA | -2.3 | 2.07E-06 | -1.3 | 0.10 | 1.7 |
| 23Se | 23S ribosomal RNA | -3.4 | 8.63E-13 | -1.3 | 0.08 | 2.5 |
| 23Sf | 23S ribosomal RNA | -2.6 | 3.24E-07 | -1.3 | 0.09 | 1.9 |
| 23Sg | 23S ribosomal RNA | -3.1 | 2.61E-17 | -1.6 | 0.00 | 1.9 |
| 5Sa | 5S ribosomal RNA | -3.9 | 1.46E-08 | -2.2 | 0.00 | 1.8 |
| 5Sb | 5S ribosomal RNA | -5.1 | 1.72E-06 | -4.5 | 0.00 | 1.1 |
| 5Sc | 5S ribosomal RNA | -25.8 | 1.04E-07 | -5.1 | 0.00 | 5.0 |
| 5Sd | 5S ribosomal RNA | -24.7 | 7.30E-08 | -5.0 | 0.00 | 5.0 |
| 5Se | 5S ribosomal RNA | -3.6 | 2.42E-05 | -2.7 | 0.00 | 1.4 |
| 5Sf | 5S ribosomal RNA | -9.2 | 3.89E-53 | -2.8 | 0.00 | 3.2 |
| 5Sg | 5S ribosomal RNA | -19124.1 | 1.84E-06 | -9.7 | 0.00 | 1970.1 |
| 5Sh | 5S ribosomal RNA | -5.5 | 1.18E-12 | -3.1 | 0.00 | 1.8 |
| **Genes repressed by ToxR** | | | | | | |
| **Locus tag** | **Function** | **FC ∆*toxR* / WT** | ***p*-value** | **Function** | **FC WT/ *toxR*** | **Ratio: *toxR* / *toxR*-pOueS** |
| VC0134 | conserved hypothetical protein | 2.6 | 6.63E-12 | 1.1 | 0.56 | 2.4 |
| VC0156 | vitamin B12 receptor | 3.5 | 1.40E-20 | -1.1 | 0.38 | -3.2 |
| VC0164 | multidrug resistance protein, putative | 2.2 | 1.03E-03 | -1.5 | 0.00 | -1.5 |
| VC0218 | ribosomal protein L28 | 2.1 | 1.83E-08 | 1.3 | 0.00 | 1.6 |
| VC0324 | ribosomal protein L11, RplK | 3.2 | 1.35E-14 | -1.2 | 0.28 | -2.8 |
| VC0325 | ribosomal protein L1 | 2.8 | 3.17E-12 | -1.3 | 0.06 | -2.2 |
| VC0326 | ribosomal protein L10 | 3.0 | 3.02E-12 | -1.3 | 0.05 | -2.3 |
| VC0327 | ribosomal protein L7/L12 | 2.7 | 1.08E-12 | -1.4 | 0.00 | -1.9 |

| | | | | | | |
|---|---|---|---|---|---|---|
| VC0359 | ribosomal protein S12 | 2.5 | 8.89E-10 | -1.1 | 0.34 | -2.2 |
| VC0360 | ribosomal protein S7 | 2.5 | 1.81E-11 | -1.3 | 0.01 | -2.0 |
| VC0362 | elongation factor TU | 2.1 | 2.70E-06 | -1.3 | 0.10 | -1.7 |
| VC0367 | primosomal replication protein N | 2.1 | 1.13E-03 | -1.3 | 0.02 | -1.6 |
| VC0368 | ribosomal protein S18 | 2.2 | 6.50E-09 | -1.4 | 0.00 | -1.5 |
| VC0369 | ribosomal protein L9 | 2.2 | 9.63E-09 | -1.4 | 0.00 | -1.5 |
| VC0477 | phosphoglycerate kinase | 2.3 | 2.68E-04 | -1.1 | 0.35 | -2.1 |
| VC0485 | pyruvate kinase I | 2.5 | 2.75E-12 | 1.1 | 0.27 | 2.2 |
| VC0561 | ribosomal protein S16 | 2.4 | 2.32E-10 | -1.0 | 0.97 | -2.3 |
| VC0562 | 16S rRNA processing protein RimM | 2.1 | 1.00E-03 | -1.1 | 0.29 | -1.9 |
| VC0563 | tRNA (guanine-N1)-methyltransferase | 2.2 | 8.78E-04 | -1.2 | 0.01 | -1.8 |
| VC0564 | ribosomal protein L19 | 2.8 | 1.24E-14 | -1.3 | 0.02 | -2.1 |
| VC0596 | dnaK suppressor protein | 2.0 | 3.95E-07 | 1.1 | 0.36 | 1.9 |
| VC0606 | nitrogen regulatory protein P-II | 4.4 | 2.95E-05 | -1.1 | 0.66 | -4.0 |
| VC0607 | Pseudogene | 2.8 | 2.46E-03 | -1.1 | 0.48 | -2.4 |
| VC0608 | iron(III) ABC transporter, periplasmiciron-compound-binding protein | 2.3 | 1.13E-10 | -1.2 | 0.17 | -1.9 |
| VC0642 | N utilization substance protein A | 2.6 | 3.92E-14 | 1.0 | 0.73 | 2.5 |
| VC0643 | initiation factor IF-2 | 2.3 | 1.76E-09 | -1.6 | 0.00 | -1.4 |
| VC0646 | ribosomal protein S15 | 2.5 | 9.00E-05 | -1.1 | 0.30 | -2.2 |
| VC0664 | lysyl-tRNA synthetase, heat inducible | 2.4 | 2.38E-16 | -1.0 | 0.90 | -2.3 |
| VC0687 | carbon starvation protein A, putative | 8.7 | 2.48E-45 | -2.6 | 0.00 | -3.4 |
| VC0695 | phospho-2-dehydro-3- deoxyheptonate aldolase,tyr-sensitive | 2.7 | 9.50E-26 | -1.3 | 0.00 | -2.0 |
| VC0739 | S-adenosylmethionine :tRNAribosyltransferase-isomer ase | 2.2 | 1.46E-13 | 1.4 | 0.00 | 1.6 |
| VC0770 | conserved hypothetical protein | 3.7 | 3.30E-08 | -1.5 | 0.03 | -2.4 |
| VC0771 | vibriobactin-specifi c isochorismatase | 2.1 | 8.52E-04 | -1.1 | 0.78 | -1.9 |
| VC0810 | hypothetical protein | 2.0 | 5.59E-04 | 1.0 | 0.94 | 2.0 |
| VC0847 | integrase, phage family | 2.1 | 9.15E-09 | -1.0 | 0.92 | -2.1 |
| VC0871 | hypothetical protein | 2.6 | 3.45E-12 | -1.4 | 0.09 | -1.9 |
| VC0872 | conserved hypothetical protein | 2.9 | 1.23E-11 | -1.2 | 0.40 | -2.5 |
| VC0905 | lipoprotein YaeC | 2.0 | 9.51E-09 | -1.5 | 0.00 | -1.4 |
| VC0916 | phosphotyrosine protein phosphatase | 4.3 | 3.63E-04 | 1.5 | 0.46 | 2.9 |
| VC0928 | hypothetical protein | 2.2 | 3.59E-07 | 1.0 | 0.83 | 2.1 |
| VC0945 | conserved hypothetical protein | 2.1 | 1.64E-06 | 1.1 | 0.38 | 1.8 |
| VC0985 | heat shock protein HtpG | 2.5 | 3.22E-11 | 1.0 | 0.61 | 2.4 |
| VC0991 | asparagine synthetase B, glutamine-hydrolyzing | 2.5 | 2.35E-17 | -1.0 | 0.72 | -2.4 |
| VC1040 | cob(I)alamin adenosyltransferase | 2.1 | 1.41E-08 | -1.0 | 0.89 | -2.1 |
| VC1193 | hypothetical protein | 2.1 | 8.15E-07 | -1.1 | 0.69 | -1.9 |
| VC1317 | conserved hypothetical protein | 2.5 | 9.72E-15 | 1.3 | 0.01 | 2.0 |
| VC1318 | outer membrane protein OmpV | 4.1 | 6.21E-15 | 1.1 | 0.48 | 3.6 |
| VC1375 | hypothetical protein | 2.1 | 8.24E-06 | 1.4 | 0.01 | 1.5 |
| VC1391 | multidrug transporter, putative | 2.7 | 3.66E-13 | 1.2 | 0.06 | 2.2 |
| VC1409 | multidrug resistance protein, putative | 2.2 | 2.40E-10 | 1.2 | 0.11 | 1.9 |
| VC1410 | multidrug resistance protein VceA | 2.6 | 9.25E-13 | 1.1 | 0.70 | 2.4 |
| VC1415 | hcp protein | 6.5 | 7.40E-36 | -1.8 | 0.00 | -3.7 |
| VC1416 | vgrG protein | 3.2 | 1.48E-21 | -1.1 | 0.51 | -3.0 |
| VC1417 | hypothetical protein | 3.1 | 3.58E-28 | -1.2 | 0.20 | -2.7 |
| VC1418 | hypothetical protein | 2.3 | 6.96E-16 | -1.4 | 0.00 | -1.6 |
| VC1419 | hypothetical protein | 2.2 | 8.38E-15 | -1.2 | 0.24 | -1.8 |
| VC1420 | hypothetical protein | 2.4 | 1.66E-09 | -1.2 | 0.23 | -2.0 |
| VC1487 | conserved hypothetical protein | 2.3 | 1.32E-04 | 1.0 | 1.00 | 2.2 |
| VC1589 | alpha-acetolactate decarboxylase | 2.5 | 3.02E-06 | -1.0 | 0.87 | -2.4 |
| VC1590 | acetolactate synthase | 3.3 | 1.04E-16 | 1.0 | 0.81 | 3.3 |
| VC1591 | oxidoreductase, short-chaindehydrogenase/reduc tase family | 3.7 | 8.01E-22 | -1.2 | 0.07 | -3.0 |
| VC1623 | carboxynorspermidine decarboxylase | 3.0 | 3.83E-33 | -1.2 | 0.12 | -2.6 |
| VC1624 | conserved hypothetical protein | 2.2 | 3.54E-18 | -1.4 | 0.02 | -1.6 |
| VC1625 | Pseudogene; involved in spermidine biosynthesis | 2.3 | 1.05E-12 | -1.1 | 0.66 | -2.2 |
| VC1628 | conserved hypothetical protein | 2.5 | 1.04E-19 | 1.0 | 0.92 | 2.5 |
| VC1629 | conserved hypothetical protein | 2.1 | 3.44E-08 | 1.2 | 0.07 | 1.7 |
| VC1701 | conserved hypothetical protein | 2.0 | 4.99E-04 | 1.1 | 0.58 | 1.8 |
| VC1721 | transcriptional regulator, LacI family | 2.0 | 1.31E-11 | 1.2 | 0.03 | 1.7 |
| VC1748 | hypothetical protein | 2.5 | 7.44E-08 | 1.6 | 0.01 | 1.6 |
| VC1749 | hypothetical protein | 2.1 | 5.51E-07 | 1.3 | 0.04 | 1.5 |
| VC1750 | hypothetical protein | 2.3 | 5.86E-10 | 1.2 | 0.06 | 1.9 |
| VC1790 | transposase OrfAB, subunit A | 2.5 | 1.59E-04 | 1.2 | 0.50 | 2.1 |
| VC1823 | PTS system, fructose-specific IIB component | 2.2 | 3.29E-07 | 1.2 | 0.15 | 1.8 |
| VC1835 | peptidoglycan-associ ated lipoprotein | 2.1 | 1.44E-03 | -1.1 | 0.19 | -1.9 |
| VC1865 | hypothetical protein | 2.9 | 1.68E-18 | -1.2 | 0.02 | -2.4 |
| VC2022 | malonyl Coa-acyl carrier protein transacylase | 2.2 | 7.09E-19 | -1.1 | 0.19 | -2.0 |
| VC2033 | alcohol dehydrogenase/acetaldehydedehy drogenase | 3.1 | 1.75E-13 | -1.3 | 0.04 | -2.4 |
| VC2045 | superoxide dismutase, Fe | 2.2 | 8.82E-09 | -1.2 | 0.12 | -1.8 |
| VC2078 | ferrous iron transport protein A | 2.2 | 1.36E-05 | -1.4 | 0.10 | -1.6 |
| VC2213 | outer membrane protein OmpA | 4.1 | 3.04E-17 | 1.1 | 0.55 | 3.7 |
| VC2259 | elongation factor Ts | 3.4 | 1.00E-18 | -1.3 | 0.00 | -2.7 |
| VC2260 | ribosomal protein S2 | 3.0 | 2.34E-13 | -1.1 | 0.48 | -2.8 |
| VC2371 | conserved hypothetical protein | 2.5 | 9.91E-07 | 1.0 | 0.97 | 2.5 |
| VC2385 | RNA-directed DNA polymerase | 2.1 | 2.70E-16 | 1.2 | 0.08 | 1.8 |
| VC2386 | conserved hypothetical protein | 2.7 | 5.02E-15 | 1.1 | 0.46 | 2.5 |
| VC2387 | conserved hypothetical protein | 2.4 | 1.59E-09 | 1.1 | 0.38 | 2.1 |
| VC2389 | carbamoyl-phosphate synthase, large subunit | 2.2 | 1.31E-13 | 1.2 | 0.23 | 1.9 |
| VC2413 | pyruvate dehydrogenase, E2 component,dihydrolipoamide acetyltransferase | 2.2 | 2.36E-03 | -1.4 | 0.00 | -1.6 |
| VC2414 | pyruvate dehydrogenase, E1 component | 2.1 | 1.52E-03 | -1.0 | 0.87 | -2.1 |
| VC2480 | ribose-5-phosphate isomerase | 2.0 | 4.00E-14 | 1.3 | 0.01 | 1.5 |
| VC2568 | peptidyl-prolyl cis-trans isomerase, FKBP-type | 2.7 | 2.20E-21 | -1.3 | 0.01 | -2.2 |
| VC2569 | hypothetical protein | 2.2 | 6.50E-10 | -1.3 | 0.08 | -1.7 |

| | | | | | | |
|---|---|---|---|---|---|---|
| VC2579 | ribosomal protein S5 | 2.1 | 1.33E-08 | -1.5 | 0.00 | -1.4 |
| VC2580 | ribosomal protein L18 | 2.3 | 4.53E-04 | -1.3 | 0.02 | -1.8 |
| VC2581 | ribosomal protein L6 | 2.1 | 2.37E-07 | -1.3 | 0.06 | -1.6 |
| VC2582 | ribosomal protein S8 | 2.3 | 5.82E-04 | -1.1 | 0.20 | -2.0 |
| VC2583 | ribosomal protein S14 | 2.1 | 6.38E-08 | -1.2 | 0.25 | -1.8 |
| VC2585 | ribosomal protein L24 | 2.1 | 3.15E-07 | -1.1 | 0.13 | -1.8 |
| VC2587 | ribosomal protein S17 | 2.2 | 4.92E-12 | -1.6 | 0.00 | -1.4 |
| VC2588 | ribosomal protein L29 | 2.2 | 6.64E-10 | -1.5 | 0.00 | -1.4 |
| VC2589 | ribosomal protein L16 | 2.2 | 1.07E-09 | -1.6 | 0.00 | -1.4 |
| VC2590 | ribosomal protein S3 | 2.1 | 9.09E-07 | -1.6 | 0.00 | -1.3 |
| VC2591 | ribosomal protein L22 | 2.6 | 6.69E-05 | -1.5 | 0.00 | -1.7 |
| VC2592 | ribosomal protein S19 | 2.3 | 2.77E-04 | -1.5 | 0.00 | -1.6 |
| VC2593 | ribosomal protein L2 | 2.3 | 1.53E-08 | -1.4 | 0.02 | -1.7 |
| VC2594 | ribosomal protein L23 | 2.5 | 1.43E-12 | -1.2 | 0.06 | -2.1 |
| VC2595 | ribosomal protein L4 | 2.0 | 2.77E-03 | -1.1 | 0.14 | -1.8 |
| VC2662 | conserved hypothetical protein | 3.4 | 4.58E-19 | 1.1 | 0.52 | 3.2 |
| VC2663 | molecular chaperone groEL_1 | 2.2 | 1.19E-07 | -1.4 | 0.02 | -1.6 |
| VC2664 | chaperonin, 60 Kd subunit, groES_1 | 2.2 | 1.49E-08 | 1.0 | 0.95 | 2.2 |
| VC2670 | triosephosphate isomerase | 2.6 | 9.46E-14 | 1.0 | 0.71 | 2.5 |
| VC2679 | ribosomal protein L31 | 2.6 | 3.02E-17 | 1.4 | 0.00 | 1.9 |
| VC2687 | hypothetical protein | 2.1 | 1.39E-03 | 1.1 | 0.55 | 2.0 |
| VC2691 | periplasmic protein cpxP, putative | 3.8 | 5.72E-15 | 1.0 | 0.84 | 3.6 |
| VC2715 | transcription elongation factor GreB | 2.0 | 1.58E-04 | 1.5 | 0.01 | 1.3 |
| VC2746 | glutamate--ammonia ligase | 2.2 | 5.23E-06 | -1.4 | 0.01 | -1.5 |
| VC2753 | hypothetical protein | 2.0 | 1.49E-04 | 1.1 | 0.51 | 1.8 |
| VC2761 | multidrug resistance protein | 3.6 | 1.89E-19 | -1.3 | 0.02 | -2.7 |
| VCA0017 | hcp-2 protein | 7.4 | 2.06E-41 | -1.7 | 0.00 | -4.4 |
| VCA0018 | vgrG protein | 2.1 | 3.35E-06 | -1.1 | 0.54 | -1.9 |
| VCA0019 | hypothetical protein | 3.3 | 7.85E-14 | -1.1 | 0.76 | -3.1 |
| VCA0020 | hypothetical protein | 2.7 | 1.40E-27 | -1.2 | 0.07 | -2.2 |
| VCA0021 | hypothetical protein | 2.0 | 8.31E-07 | -1.1 | 0.43 | -1.9 |
| VCA0023 | hypothetical protein | 2.1 | 1.28E-04 | 1.2 | 0.28 | 1.7 |
| VCA0035 | phosphatidylglycerop hosphatase B, putative | 3.0 | 8.88E-17 | 1.7 | 0.00 | 1.8 |
| VCA0087 | hypothetical protein | 3.0 | 2.25E-07 | 1.5 | 0.10 | 2.1 |
| VCA0088 | proton/glutamate symporter | 2.1 | 1.25E-10 | 1.1 | 0.23 | 1.8 |
| VCA0107 | conserved hypothetical protein | 2.7 | 1.39E-16 | -1.3 | 0.03 | -2.1 |
| VCA0108 | conserved hypothetical protein | 2.9 | 1.58E-15 | -1.1 | 0.16 | -2.5 |
| VCA0112 | hypothetical protein | 2.2 | 4.79E-11 | -1.5 | 0.00 | -1.4 |
| VCA0136 | glycerophosphoryl diester phosphodiesterase | 8.9 | 1.84E-52 | -1.3 | 0.00 | -6.8 |
| VCA0137 | glycerol-3-phosphate transporter | 8.8 | 1.31E-19 | -2.0 | 0.00 | -4.5 |
| VCA0139 | hypothetical protein | 3.0 | 1.10E-12 | 1.4 | 0.02 | 2.2 |
| VCA0227 | iron(III) ABC transporter, periplasmiciron-compound-binding protein | 3.3 | 1.42E-06 | -1.1 | 0.62 | -2.9 |
| VCA0230 | iron(III) ABC transporter, ATP-binding protein | 2.9 | 5.74E-09 | -1.3 | 0.19 | -2.3 |
| VCA0363 | hypothetical protein | 2.3 | 1.07E-04 | 1.5 | 0.09 | 1.5 |
| VCA0370 | hypothetical protein | 4.4 | 2.67E-03 | 1.6 | 0.61 | 2.8 |
| VCA0407 | hypothetical protein | 2.6 | 4.02E-04 | -1.3 | 0.48 | -1.9 |
| VCA0517 | 1-phosphofructokinase | 2.9 | 2.75E-11 | -1.6 | 0.01 | -1.9 |
| VCA0518 | PTS system, fructose-specific IIA/FPR component | 5.7 | 3.31E-31 | -1.3 | 0.14 | -4.5 |
| VCA0519 | fructose repressor | 2.8 | 2.04E-15 | -1.1 | 0.53 | -2.6 |
| VCA0540 | formate transporter 1, putative | 9.0 | 2.76E-85 | -1.3 | 0.03 | -7.1 |
| VCA0542 | transcriptional regulator, LysR family | 2.5 | 7.06E-16 | 1.2 | 0.22 | 2.2 |
| VCA0563 | NAD(P) transhydrogenase, alpha subunit | 2.4 | 1.10E-11 | -1.3 | 0.01 | -1.9 |
| VCA0564 | NAD(P) transhydrogenase, beta subunit | 2.5 | 9.03E-14 | -1.2 | 0.03 | -2.1 |
| VCA0576 | heme transport protein HutA | 2.5 | 3.15E-11 | -1.0 | 0.84 | -2.4 |
| VCA0657 | aerobic glycerol-3-phosphate dehydrogenase | 39.8 | 1.99E-121 | -1.8 | 0.00 | -21.7 |
| VCA0732 | conserved hypothetical protein | 2.6 | 6.47E-10 | 1.0 | 0.76 | 2.5 |
| VCA0742 | hypothetical protein | 2.1 | 2.26E-04 | -1.1 | 0.51 | -1.9 |
| VCA0744 | glycerol kinase | 6.5 | 1.89E-39 | -1.4 | 0.01 | -4.5 |
| VCA0745 | pseudogene with authentic frameshift mutation | 6.1 | 9.74E-37 | -1.7 | 0.00 | -3.5 |
| VCA0747 | anaerobic glycerol-3-phosphate dehydrogenase,subunit A | 28.6 | 6.35E-125 | -1.5 | 0.00 | -19.4 |
| VCA0748 | anaerobic glycerol-3-phosphate dehydrogenase,subunit B | 17.1 | 3.70E-94 | -1.3 | 0.02 | -12.7 |
| VCA0749 | anaerobic glycerol-3-phosphate dehydrogenase,subunit C | 19.1 | 8.32E-91 | -1.3 | 0.10 | -15.0 |
| VCA0789 | conserved hypothetical protein | 2.9 | 2.96E-14 | 2.4 | 0.00 | 1.2 |
| VCA0819 | chaperonin, 10 Kd subunit | 2.1 | 4.20E-06 | -1.8 | 0.00 | -1.2 |
| VCA0862 | long-chain fatty acid transport protein | 2.5 | 2.92E-11 | -1.1 | 0.59 | -2.3 |
| VCA0863 | lipase, putative | 2.6 | 4.75E-17 | -1.2 | 0.18 | -2.2 |
| VCA0874 | hypothetical protein | 2.4 | 1.00E-05 | 1.6 | 0.00 | 1.5 |
| VCA0898 | 6-phosphogluconate dehydrogenase,decarboxylating | 2.9 | 2.97E-22 | -1.3 | 0.00 | -2.2 |
| VCA0933 | cold shock domain family protein | 2.4 | 1.49E-10 | 1.5 | 0.00 | 1.6 |
| VCA0984 | L-lactate dehydrogenase | 5.7 | 1.09E-33 | 1.4 | 0.04 | 4.1 |
| VCA0985 | oxidoreductase/iron- sulfur cluster-bindingprotein | 3.0 | 3.71E-16 | 1.1 | 0.41 | 2.8 |
| VCA1013 | conserved hypothetical protein | 2.1 | 2.59E-07 | 1.1 | 0.45 | 1.9 |
| VCA1035 | hypothetical protein | 2.5 | 9.63E-10 | 1.4 | 0.02 | 1.7 |
| VCA1060 | 3,4-dihydroxy-2-buta none 4-phosphate synthase | 2.2 | 5.69E-11 | -1.2 | 0.32 | -1.8 |
| VCA1063 | ornithine decarboxylase, inducible | 3.0 | 4.17E-12 | -2.0 | 0.00 | -1.5 |
| tRNA-Ala-2 | tRNA biosynthesis | 3.1 | 1.70E-05 | 1.2 | 0.59 | 2.5 |
| tRNA-Gln-1 | tRNA biosynthesis | 2.4 | 9.80E-05 | 1.6 | 0.07 | 1.5 |
| tRNA-Gly-3 | tRNA biosynthesis | 2.1 | 4.67E-08 | 1.1 | 0.48 | 1.9 |
| tRNA-Gly-7 | tRNA biosynthesis | 2.5 | 3.48E-04 | -3.3 | 0.01 | -0.8 |
| tRNA-Leu-2 | tRNA biosynthesis | 2.6 | 1.11E-04 | 1.5 | 0.08 | 1.7 |
| tRNA-Leu-9 | tRNA biosynthesis | 2.9 | 7.67E-04 | 2.0 | 0.09 | 1.5 |
| tRNA-Met-4 | tRNA biosynthesis | 2.2 | 1.53E-04 | -1.0 | 0.97 | -2.1 |
| tRNA-Met-5 | tRNA biosynthesis | 4.4 | 8.60E-09 | 2.1 | 0.01 | 2.0 |
| tRNA-Met-6 | tRNA biosynthesis | 2.7 | 1.33E-03 | 1.5 | 0.33 | 1.8 |

| | | | | | | |
|---|---|---|---|---|---|---|
| tRNA-Phe-1 | tRNA biosynthesis | 2.2 | 2.73E-07 | 1.3 | 0.07 | 1.7 |
| tRNA-Thr-6 | tRNA biosynthesis | 2.4 | 6.99E-05 | 1.1 | 0.69 | 2.2 |
| tRNA-Tyr-3 | tRNA biosynthesis | 3.4 | 8.81E-07 | 1.8 | 0.04 | 1.9 |
| tRNA-Tyr-4 | tRNA biosynthesis | 2.2 | 6.29E-06 | 1.2 | 0.25 | 1.9 |
| tRNA-Val-3 | tRNA biosynthesis | 2.3 | 2.39E-05 | -1.3 | 0.39 | -1.8 |
