## Supplementary material for "Modular small RNA drives pathogen emergence": Table S9

**Table S9.** Strains, vectors and primers used in this study.

| | | Strains | | |
|---|---|---|---|---|
| **Code** | **Species** | **Genotype** | **Selection** | **Source** |
| DBS3 | *V. cholerae* | El Tor C6706 wild type (WT; background for all *V. cholerae* strains) | Sm | Lab collection |
| DBS25 | *V. cholerae* | Δ*toxR* | Sm | This study |
| DBS27 | *V. cholerae* | Δ*ompU* | Sm | This study |
| DBS517 | *V. cholerae* | Δ*lacZ* | Sm | This study |
| DBS455 | *V. cholerae* | ΔP$_{ompU}$ | Sm | This study |
| DBS72 | *V. cholerae* | C6706-pBAD22 | Sm | This study |
| DBS84 | *V. cholerae* | Δ*ompU*-pBAD22 | Cb | This study |
| DBS394 | *V. cholerae* | Δ*toxR*-pBAD22 | Cb | This study |
| DBS444 | *V. cholerae* | Δ*ompU*-pBAD-OueS$^{C6706}$ | Cb | This study |
| DBS448 | *V. cholerae* | Δ*ompU*-pBAD-OueS$^{GBE1114}$ | Cb | This study |
| DBS396 | *V. cholerae* | Δ*ompU*-pOmpU$^{C6706}$ | Cb | This study |
| DBS465 | *V. cholerae* | Δ*ompU*-pOmpU$^{GBE1114}$ | Sm | This study |
| DBS310 | *V. cholerae* | Δ*ompU*::OmpU$^{GBE1114}$ | Sm | This study |
| DBS391 | *V. cholerae* | Δ*ompU*-pNS | Cb | This study |
| DBS593 | *V. cholerae* | Δ*ompU*-pOueS 5'-mut | Cb | This study |
| DBS595 | *V. cholerae* | Δ*ompU*-pOueS 3'-mut | Cb | This study |
| DBS590 | *V. cholerae* | Δ*ompU*-pOueS FL-mut | Cb | This study |
| DBS394 | *V. cholerae* | Δ*toxR*-pBAD22 | Cb | This study |
| DBS410 | *V. cholerae* | Δ*toxR*-pOmpU$^{C6706}$ | Cb | This study |
| DBS457 | *V. cholerae* | Δ*toxR*-pOueS$^{C6706}$ | Cb | This study |
| DBS408 | *V. cholerae* | Δ*toxR*-pNS | Cb | This study |
| DBS601 | *V. cholerae* | Δ*toxR*-pOueS 5'-mut | Cb | This study |
| DBS603 | *V. cholerae* | Δ*toxR*-pOueS 3'-mut | Cb | This study |
| DBS599 | *V. cholerae* | Δ*toxR*-pOueS FL-mut | Cb | This study |
| DBS539 | *V. cholerae* | Δ*ompU*-P$_{ompU}$::P$_0$-OueS$^{C6706}$ | Sm | This study |
| DBS560 | *V. cholerae* | Δ*ompU*-P$_{ompU}$::P$_0$-OueS 5'-mut | Sm | This study |
| DBS582 | *V. cholerae* | Δ*ompU*-P$_{ompU}$::P$_0$-OueS 3'-mut | Sm | This study |
| DBS584 | *V. cholerae* | Δ*ompU*-P$_{ompU}$::P$_0$-OueS FL-mut | Sm | This study |
| DBS550 | *V. cholerae* | Δ*ompU*-P$_{ompU}$::P$_0$-OueS$^{GBE1114}$ | Sm | This study |
| DBS571 | *V. cholerae* | Δ*ompU*-P$_{ompU}$::P$_0$-OueS$^{GBE0428}$ | Sm | This study |
| DBS572 | *V. cholerae* | Δ*ompU*-P$_{ompU}$::P$_0$-OueS$^{IRL0181}$ | Sm | This study |
| DBS596 | *V. cholerae* | Δ*ompU*-P$_{ompU}$::P$_0$-OueS$^{GBE1116}$ | Sm | This study |
| DBS586 | *V. cholerae* | Δ*ompU*-P$_{ompU}$::P$_0$-OueS$^{GBE1194}$ | Sm | This study |
| DBS588 | *V. cholerae* | Δ*ompU*-P$_{ompU}$::P$_0$-OueS$^{IRL0074}$ | Sm | This study |
| DBS488 | *V. cholerae* | Δ*ompU*-P$_{ompU}$::P$_{10}$-OueS$^{C6706}$ | Sm | This study |
| DBS489 | *V. cholerae* | Δ*ompU*-P$_{ompU}$::P$_{20}$-OueS$^{C6706}$ | Sm | This study |
| DBS491 | *V. cholerae* | Δ*ompU*-P$_{ompU}$::P$_{30}$-OueS$^{C6706}$ | Sm | This study |
| DBS493 | *V. cholerae* | Δ*ompU*-P$_{ompU}$::P$_{40}$-OueS$^{C6706}$ | Sm | This study |
| DBS495 | *V. cholerae* | Δ*ompU*-P$_{ompU}$::P$_{50}$-OueS$^{C6706}$ | Sm | This study |
| DBS509 | *V. cholerae* | Δ*ompU*-P$_{ompU}$::P$_{60}$-OueS$^{C6706}$ | Sm | This study |
| DBS499 | *V. cholerae* | Δ*ompU*-P$_{ompU}$::P$_{80}$-OueS$^{C6706}$ | Sm | This study |
| DBS512 | *V. cholerae* | Δ*ompU*-P$_{ompU}$::P$_{90}$-OueS$^{C6706}$ | Sm | This study |
| DBS501 | *V. cholerae* | Δ*ompU*-P$_{ompU}$::P$_{100}$-OueS$^{C6706}$ | Sm | This study |
| DBS16 | *E. coli* | *E. coli* S17-1 **λ***pir* | -- | Lab collection |
| DBS18 | *E. coli* | *E. coli* S17-1 **λ***pir*-pKAS154 | Kan | Lab collection |
| DBS19 | *E. coli* | *E. coli* S17-1 **λ***pir*-pBAD22 | Cb | Lab collection |
| SM80 | *E. coli* | pKAS154Δ*toxR* | Kan | Lab collection |
| SM125 | *E. coli* | pKAS154Δ*ompU* | Kan | Lab collection |
| DBS513 | *E. coli* | pKAS154Δ*lacZ* | Kan | This study |
| DBS440 | *E. coli* | pKAS154ΔP$_{ompU}$ | Kan | This study |
| DBS393 | *E. coli* | pBAD-OmpU$^{C6706}$ | Cb | This study |
| DBS454 | *E. coli* | pOmpU$^{GBE1114}$ | Cb | This study |
| DBS388 | *E. coli* | pBAD-NS | Cb | This study |
| DBS521 | *E. coli* | pKAS154Δ*ompU*- P$_{ompU}$-P$_0$OueS$^{C6706}$ | Kan | This study |
| DBS470 | *E. coli* | pKAS154Δ*ompU*- P$_{ompU}$-P$_{10}$OueS$^{C6706}$ | Kan | This study |

| | | | | |
|---|---|---|---|---|
| DBS471 | *E. coli* | pKAS154Δ*ompU*- $P_{ompU}$-$P_{20}$OueS$^{C6706}$ | Kan | This study |
| DBS472 | *E. coli* | pKAS154Δ*ompU*- $P_{ompU}$-$P_{30}$OueS$^{C6706}$ | Kan | This study |
| DBS473 | *E. coli* | pKAS154Δ*ompU*- $P_{ompU}$-$P_{40}$OueS$^{C6706}$ | Kan | This study |
| DBS474 | *E. coli* | pKAS154Δ*ompU*- $P_{ompU}$-$P_{50}$OueS$^{C6706}$ | Kan | This study |
| DBS475 | *E. coli* | pKAS154Δ*ompU*- $P_{ompU}$-$P_{70}$OueS$^{C6706}$ | Kan | This study |
| DBS476 | *E. coli* | pKAS154Δ*ompU*- $P_{ompU}$-$P_{80}$OueS$^{C6706}$ | Kan | This study |
| DBS505 | *E. coli* | pKAS154Δ*ompU*- $P_{ompU}$-$P_{90}$OueS$^{C6706}$ | Kan | This study |
| DBS477 | *E. coli* | pKAS154Δ*ompU*- $P_{ompU}$-$P_{100}$OueS$^{C6706}$ | Kan | This study |
| DBS546 | *E. coli* | pKAS154Δ*ompU*- $P_{ompU}$-$P_{0}$OueS-5' mut | Kan | This study |
| DBS577 | *E. coli* | pKAS154Δ*ompU*- $P_{ompU}$-$P_{0}$OueS-3' mut | Kan | This study |
| DBS579 | *E. coli* | pKAS154Δ*ompU*- $P_{ompU}$-$P_{0}$OueS-FL mut | Kan | This study |
| DBS540 | *E. coli* | pKAS154Δ*ompU*- $P_{ompU}$-$P_{0}$OueS$^{GBE1114}$ | Kan | This study |
| DBS561 | *E. coli* | pKAS154Δ*ompU*- $P_{ompU}$-$P_{0}$OueS$^{GBE0428}$ | Kan | This study |
| DBS562 | *E. coli* | pKAS154Δ*ompU*- $P_{ompU}$-$P_{0}$OueS$^{IRL0181}$ | Kan | This study |
| DBS564 | *E. coli* | pKAS154Δ*ompU*- $P_{ompU}$-$P_{0}$OueS$^{GBE1116}$ | Kan | This study |
| DBS565 | *E. coli* | pKAS154Δ*ompU*- $P_{ompU}$-$P_{0}$OueS$^{GBE1194}$ | Kan | This study |
| DBS566 | *E. coli* | pKAS154Δ*ompU*- $P_{ompU}$-$P_{0}$OueS$^{IRL0074}$ | Kan | This study |

| | **Plasmids** | |
|---|---|---|
| **Plasmid** | **Notes** | **Selection** |
| pBAD22 | Ectopic expression | Cb |
| pBAD-OmpU$^{C6706}$ | Ectopic expression | Cb |
| pBAD-OmpU$^{GBE1114}$ | Ectopic expression | Cb |
| pBAD-Non-specific (NS) | Ectopic expression | Cb |
| pBAD-OueS$^{C6706}$ | Ectopic expression | Cb |
| pBAD-OueS 5'-mut | Ectopic expression | Cb |
| pBAD-OueS 3'-mut | Ectopic expression | Cb |
| pBAD-OueS FL-mut | Ectopic expression | Cb |
| pKAS154 | Allelic exchange | Kan |
| pKAS-OueS$^{C6706}$ | Allelic exchange | Kan |
| pKAS-Δ*ompU* | Allelic exchange | Kan |
| pKAS-Δ*toxR* | Allelic exchange | Kan |
| pKAS-Δ*lacZ* | Allelic exchange | Kan |
| pKAS-Δ$P_{ompU}$ | Allelic exchange | Kan |
| pKAS-Δ$P_{ompU}$ | Allelic exchange | Kan |
| pKAS-$P_{10}$OueS$^{C6706}$ | Allelic exchange | Kan |
| pKAS-$P_{20}$OueS$^{C6706}$ | Allelic exchange | Kan |
| pKAS-$P_{30}$OueS$^{C6706}$ | Allelic exchange | Kan |
| pKAS-$P_{40}$OueS$^{C6706}$ | Allelic exchange | Kan |
| pKAS-$P_{50}$OueS$^{C6706}$ | Allelic exchange | Kan |
| pKAS-$P_{60}$OueS$^{C6706}$ | Allelic exchange | Kan |
| pKAS-$P_{80}$OueS$^{C6706}$ | Allelic exchange | Kan |
| pKAS-$P_{90}$OueS$^{C6706}$ | Allelic exchange | Kan |
| pKAS-$P_{100}$OueS$^{C6706}$ | Allelic exchange | Kan |
| pKAS-OueS$^{GBE1114}$ | Allelic exchange | Kan |
| pKAS-OueS$^{GBE0428}$ | Allelic exchange | Kan |
| pKAS-OueS$^{IRL0181}$ | Allelic exchange | Kan |
| pKAS-OueS$^{GBE1116}$ | Allelic exchange | Kan |
| pKAS-OueS$^{GBE1194}$ | Allelic exchange | Kan |
| pKAS-OueS$^{IRL0074}$ | Allelic exchange | Kan |

| | Primers | |
|---|---|---|
| **Name** | **Sequence** | **Use** |
| DB215 | **ctagggaAGATCT**aaaatgattggctaatttgcgaa | PompU USF |
| DB216 | **ataagaatGCGGCCGC**agattgagcaaaatgcacgcaatc | PompU USR |
| DB217 | **ataagaatGCGGCCGC**atggacaataaattaggacttaa | PompU DSF |

| | | |
|---|---|---|
| DB218 | **ccgccgGAATTC**gaaacgttgctgcagtataacga | PompU DSR |
| DB219 | tcagccaatggatgaaaaaaggtcg | PompU FlkF |
| TG28 | GCGGCAGTCAATGGTGTTTG | OmpU Flk Rev |
| DB220 | **ctagggaAGATCT**acttggctagaaccggggccaggaa | ΔlacZ US-F |
| DB221 | **ataagaatGCGGCCGC**ccctcaagccgaggagtaaagaagt | ΔlacZ US-R |
| DB222 | **ataagaatGCGGCCGC**taagccagagagccttaaggctctcttttttgt | ΔlacZ DS-F |
| DB223 | **ccgccgGAATTC**gtcaaggacatagaaacattgcttg | ΔlacZ DS-R |
| DB224 | gcacggagggaagggtaaaaccgag | ΔlacZ Flk F |
| DB225 | acgccaacgaggtaaaaacgctggt | ΔlacZ Flk R |
| DB226 | **ctagcGAATTC**gccaaacttccgctcttacatctc | OueS-C6706 Fwd |
| DB228 | **ctagcAAGCTT**TAGAAAAACGCCTGCTAGTGAGCA | OueS-C6706 Rev |
| DB249 | **ccatGAGCTC**gccaaacttccgctcttaca | OueS-0 |
| DB233 | **ccatGAGCTC**actacttcaagccaaacttc | OueS-10 |
| DB234 | **ccatGAGCTC**gacgcaacttactacttcaa | OueS-20 |
| DB235 | **ccatGAGCTC**ttttgctatcgacgcaactt | OueS-30 |
| DB236 | **ccatGAGCTC**cagcagataattttgctatc | OueS-40 |
| DB237 | **ccatGAGCTC**aaagaaacttcagcagataa | OueS-50 |
| DB239 | **ccatGAGCTC**acaacaatgcagaaacagcg | OueS-70 |
| DB240 | **ccatGAGCTC**actgcgacatacaacaatgc | OueS-80 |
| DB241 | **ccatGAGCTC**agctgcgtttactgcgacat | OueS-90 |
| DB242 | **ccatGAGCTC**aactaggtcaagctgcgttt | OueS-100 |
| DB243 | **ccgccgGAATTC**gggtggggtcgcctgaaaagcg | Reverse primer for OueS |
| DB256 | **ccatGAGCTC**ACCTAATTTTCGGTCATATA | OueS-$P_0$-5'mut |
| DB199 | **ctagcGAATTC**agcgttacggtcagcaaaac | Non-specific locus Fwd |
| DB200 | **ctagcAAGCTT**tggtatcggtggcacttac | Non-specific locus Rev |
| DB271 | **ccgccgGAATTC**ACCTAATTTTCGGTCATATA | OueSFL-mutF |
| DB272 | **ctagcAAGCTT**TAGTAATACTCGAGCAAGAG | OueSFL-mutR |
| DB273 | **ccgccgGAATTC**GCCAAACTTCCGCTCTTACA | OueS3'-mutF |
| DB274 | **ctagcAAGCTT**TAGAACTAGTCAAGTAGGAT | OueS3'-mutR |
| DB195 | **GATTACGCCAAGCTT**ccgctcttacatctcttaccagttcaatctgctag | for 3' RACE |
| DB196 | **GATTACGCCAAGCTT**tcgtaacgtagaccgatagccagttcgtct | For 5' RACE |
| DB164 | GGGGGGGGAATTCTTTTTTTTTTTT | RACE primer |
| DB133 | ctgaggtaattataacccgg | pKAS32 MCS Flk F |
| DB134 | GGACAACAAGCCAGGGATGT | pKAS32 MCS Flk R |
| GR13 | atgccatagcatttttatcc | pBAS MCS flank |
| GR14 | gatttaatctgtatcagg | pBAS MCS flank |
